# Analysis of Sequences of Bases of Human DNA Using the Square Wave Method (SWM)

**DOI:** 10.1101/2023.11.13.566944

**Authors:** Osvaldo Skliar, Sherry Gapper, Ricardo E. Monge

## Abstract

A description is provided of how the Square Wave Method (SWM) can be applied to analyze sequences of bases of human DNA. The results obtained are displayed using the Square Wave Transform (SWT), an SWM tool. It is hypothesized that the results of this analysis can be of interest, in certain cases, to understand the functional role of some of those sequences. Preliminary data which make this hypothesis plausible are presented.

## 1 Introduction

The Square Wave Method (SWM) is a new algebraic method for the analysis of signals and images [1], [2], [3], [4], and [5]. The Square Wave Transform (SWT) [3] is a tool belonging to the SWM which makes it possible to easily visualize the results obtained when using the method.

Mention is made here of contributions of authors other than those who introduced the SWM, in which reference is made to that method. One of them [6] states: “The idea behind our approach builds upon existing classical results that use *exact* square waves to approximate a function (39)”. The only reference to those “existing classical results” is (39), which is the same as [5] in the present article. Reference to that article is also made in [7].

Other sources, [8], [9], [10],[11] and [12], also refer to that article. The SWM has been mentioned frequently in patents: [13], [14], [15], [16], [17], [18], [19], [20], [21], [22], [23], [24], [25], [26], [27], [28], [29], [30], [31], [32], [33], [34], [35], [36], [37], [38], [39], [40], [41], [42], [43], [44], [45], [46], [47], [48], [49], [50], [51], [52], [53], [54], [55], [56], [57], [58], [59], [60], [61], [62], [63], [64], [65], [66], [67], [68], [69], [70], [71], [72], [73], [74], [75], [76], [77], [78], [79], [80], [81], [82], [83], [84], [85], [86], [87], [88], [89], [90], [91], [92], [93], [94], [95], [96], [97], [98], [99], [100], [101], [102], [103], [104], [105], [106], [107], [108], [109], [110], [111], [112], [113], [114], [115], [116], [117], [118], [119], [120], [121], [122], [123], [124], [125], [126], [127], [128], [129] and [130].

The objectives of this article are the following:

1. specify how sequences of bases of DNA can be analyzed using the SWM and the results thus obtained, presented using the SWT; and
2. formulate the hypothesis that at least in certain cases considering the sequences of bases of the nucleic acids – of DNA in particular – as signals susceptible to being analyzed with a method characteristic of signal analysis, such as the SWM, can be useful in genomics. Preliminary data which make this hypothesis plausible are presented.

## 2 Brief Review of the Use of the SWM for the Analysis of Numerical Sequences

As seen in the following section of this article, the first operation to be carried out to analyze a sequence of bases of DNA with the SWM is the codification of that sequence as a numerical sequence. Therefore, this section will be useful for those who are not familiar with articles on the SWM.

The SWM makes it possible to approximate numerical sequences using a certain sum, to be specified, of parts of trains of digitalized square waves. The concept of “digitalized square wave” is justified because the wave is not treated as a function with a sole discontinuity (in the transition from positive semiwave to negative semiwave, or vice versa) but rather as a sequence of numerical values. Another concept which should be characterized in the treatment of numerical sequences using the SWM is that of frequency.

Usually, in the case of a phenomenon that is repeated with the same characteristics, by *frequency* of that phenomenon one understands the number – *n* – of occasions that it occurs in each unit of time. If the second – *s* – is taken as the unit of time, then the *frequency* of this phenomenon is defined as: *<u>^n^</u>* = *ns^−^*^1^ = *n* Hz. When analyzing a numerical sequence using the SWM, the unit of time is replaced by a “numerical unit” composed of as many elemental numerical subunits as the number of numerical values that the sequence has. Thus, for example, consider the following numerical sequence:

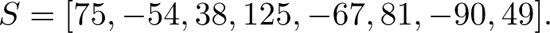

The above numerical sequence is composed of 8 elemental subunits which are the numerical values that constitute that sequence.

The elements corresponding to the phenomena that are repeated with the same characteristics are square waves – the elements comprising each train of square waves. Consider a train of square waves symbolized as follows:

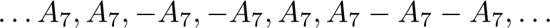

(A justification will be provided below for the numerical values of the subscripts used in this section for the symbols *A_i_* for *i* = 1, 2, 3*, …,* 8.) It can be admitted that this train is prolonged to infinity in both possible directions. Each square wave composing that train of square waves can be symbolized as *A*_7_*, A*_7_*, −A*_7_*, −A*_7_. The frequency corresponding to that train of square waves regarding the numerical unit specified above can be computed by dividing the number of elemental numerical subunits constituting that unit (8) by the number of elemental numerical subunits constituting the square wave considered (4): *f*_7_ = ^8^ = 2. The numerical unit of reference could be thought to “cover” 2 square waves of the type considered; in other words, the event which is repeated – the square wave – occurs twice in the numerical unit of reference.

Consider another train of square waves:

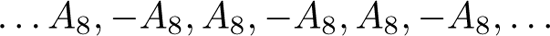

In this case, the square wave constituting the above train of square waves can be symbolized as *A*_8_*, −A*_8_. The frequency corresponding to that train of square waves with respect to the numerical unit of reference considered is computed as: *f*_8_ = ^8^ = 4. In this case, the numerical unit of reference (composed of 8 elemental numerical subunits) “covers” 4 square waves of the type considered. In other words, 4 square waves of this type “occupy a place” in the numerical unit of reference.

Consider another train of square waves:

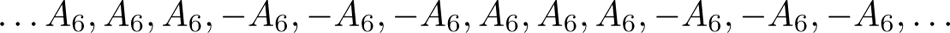

In this case, the square wave repeated in the train of square waves specified is the following: *A*_6_*, A*_6_*, A*_6_*, −A*_6_*, −A*_6_*, −A*_6_. This square wave is composed of 6 elemental units. The frequency of the last train of square waves regarding the numerical unit of reference is: *f*_6_ = ^8^ = ^4^. Here the numerical unit of reference, composed of 8 elemental numerical subunits, “covers” only one square wave and a fraction of a square wave of the same type.

A matrix is presented below whose rows represent certain intervals (or parts) of trains of square waves used for the analysis of the numerical sequence specified. This sequence is indicated above the matrix:

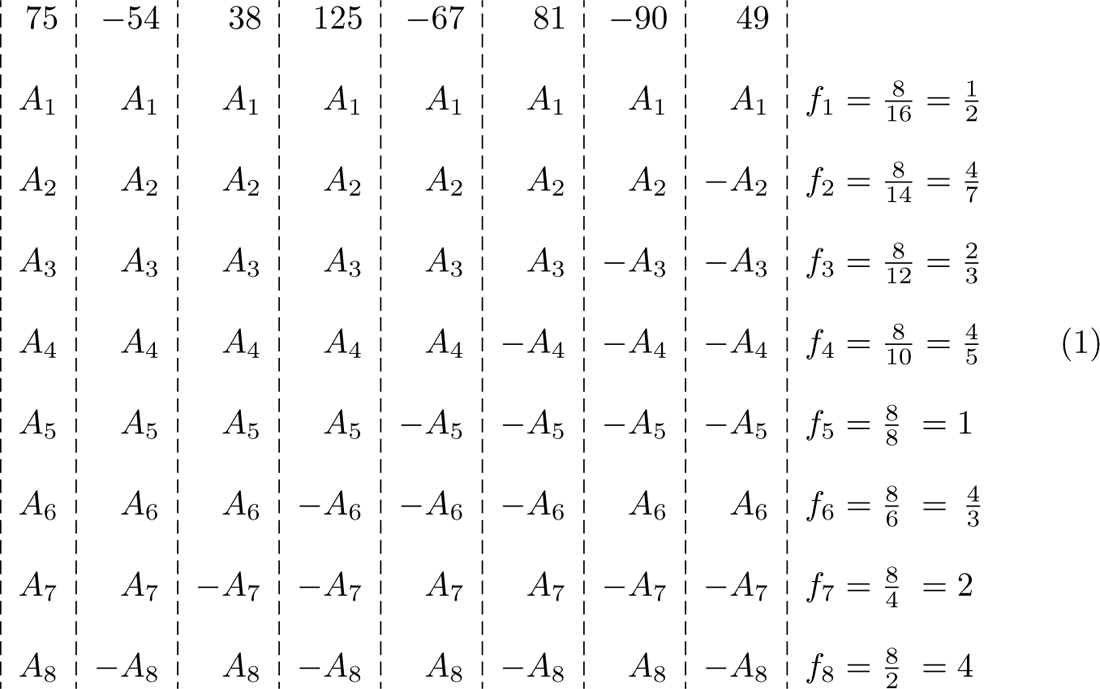

At the right of the matrix represented in (1), the corresponding frequencies for the different trains of square waves have been specified for the case considered. It can be seen that subscript 1 is used in the symbols corresponding to the numerical values of the first train of square waves and in the symbol of the corresponding frequency. Subscript 2 is used in the symbols corresponding to the numerical values of the second train of square waves and in the symbol of the respective frequency; and so on, until finally, subscript 8 is used in the symbols of the numerical values of the eighth train of square waves and in the symbol of the respective frequency. From each of these 8 trains of square waves, only 8 elemental subunits are given in (1), that is, the same number of elemental units that comprise the numerical sequence to be analyzed.

In the first row of the matrix presented in (1), one semiwave of the square wave has been symbolized. The semiwave not symbolized is composed of another 8 symbols: *−A*_1_*, −A*_1_*, −A*_1_*, −A*_1_*, −A*_1_*, −A*_1_*, −A*_1_*, −A*_1_. Each of the square waves that are constituents of the first train of square waves is composed of 16 elemental numerical subunits. In other words, the length of wave *λ*_1_ of each square wave in that train can be considered equal to 16: *λ*_1_ = 16. As indicated above, *f*_1_ = 8/16 = ½.

Note that the number of elemental subunits of each of the square waves comprising the train of square waves corresponding to the second row of the matrix represented in (1) is 14; that is, there are 2 elemental subunits less than in each square wave of the first train of square waves. Given that *f*_2_ = 8/14, = 4/7. The number of elemental subunits of each square wave of the third train of square waves – the one corresponding to the third row in the matrix represented in (1) – is 12; or that is, 2 subunits less in each square wave than in the second train of square waves. Thus, *f*_3_ = 8/12 = 2/3, and so on. Therefore, the following results are obtained: *f*_4_ = 8/10 = 4/5, *f*_5_ = 8/8 = 1, *f*_6_ = 8/6 = 4/3, *f*_7_ = 8/4 = 2 and finally *f*_8_ = ^8^ = 4. The above results are given on the right of the matrix presented in (1).

To obtain the system of linear algebraic equations whose solution makes it possible to compute the numerical values of *A*_1_*, A*_2_*, A*_3_*, A*_4_*, A*_5_*, A*_6_*, A*_7_, and *A*_8_, the following procedure is carried out. The equality is established of a) the sum of all the elements in the first column of the matrix in (1) and b) the numerical value indicated immediately above that column: 75. This numerical value is the first in the numerical sequence to be analyzed. Hence, the following equation is obtained:

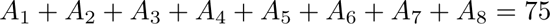

The equality is established below of a) the sum of all the elements composing the second column of the matrix represented in (1), and b) the numerical value indicated immediately above that column: *−*54. This numerical value is the second in the numerical sequence analyzed. Thus the following equation is obtained:

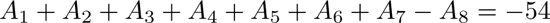

The same procedure is carried out with each of the following columns of the matrix represented in (1): For each of the columns the equality is established of a) the sum of all the elements that compose it, and b) the numerical value indicated immediately above the respective column. Thus the following system of linear algebraic equations is obtained:

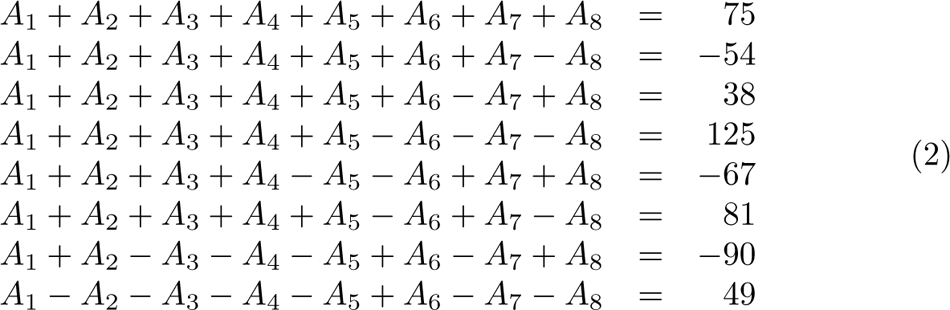

When solving the above system of algebraic linear equations (2), the following numerical values are obtained for the unknowns.

If in (2) the unknowns are replaced by the numerical values computed for them, it can be seen that the diverse equations of the system of equations (2) are satisfied.

If the order of the numerical values is inverted, a different numerical sequence is obtained which will be known as “inverted numerical sequence” with respect to the first sequence:

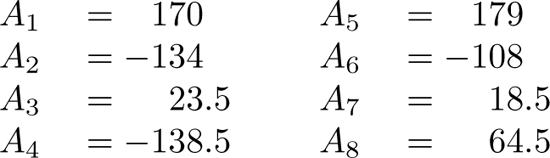

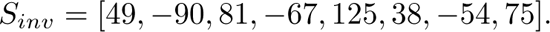

When applying the SWM to the analysis of this last numerical sequence, another sequence of linear algebraic equations is obtained. If the unknowns are computed in this last system of equations, the following numerical values are obtained for them:

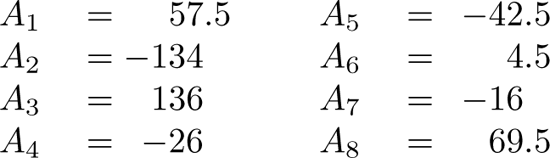

If in the last system of linear algebraic equations mentioned the unknowns are replaced by the numerical values computed for them, it can be seen that all of the equations in that system of equations are also satisfied.

When applying the SWM to the analysis of a numerical sequence, there is an important dependence of the results obtained on the order of the numerical values of the sequence considered. Because the first operation to be carried out when analyzing a sequence of bases of DNA will be to assign each base a numerical value, and given that the order of the bases is very relevant for the functional role of each sequence of bases of DNA considered, this method appears adequate for the analysis of that type of sequences.

The Square Wave Transform (SWT) – a tool of the SWM – makes it possible to present, in a system of orthogonal Cartesian coordinates, the results obtained when analyzing any numerical sequence with that method. In effect, on the *x*-axis it is possible to plot the numerical values of the different frequencies *f_i_*, for *i* = 1, 2, 3*, …, n*, corresponding to the different trains of square waves, and on the *y*-axis, the numerical values of the different *A*_1_, using dots or vertical lines.

In figure 1a, by using the SWT, the results are displayed of the analysis with the SWM of the first numerical sequence considered in this section.

**Figure 1a:**
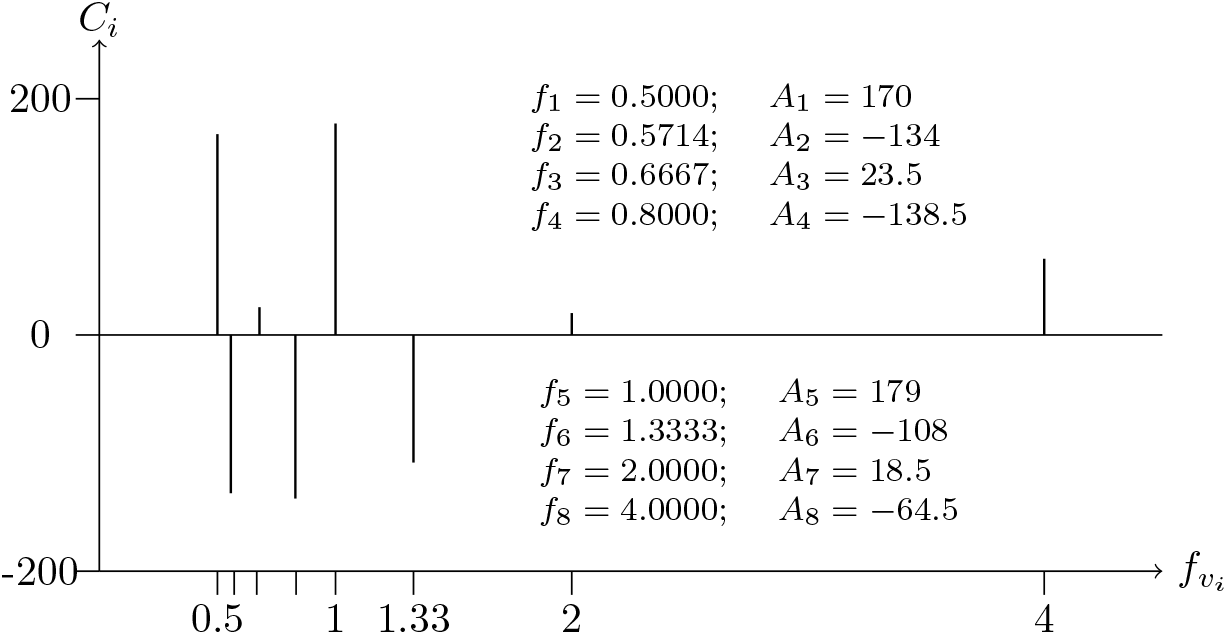
SWT which presents the results of the analysis with the SWM of the first numerical sequence considered in this section

In figure 1b, by using the SWT, the results are displayed of the analysis with the SWM of the second numerical sequence considered in this section – that is, the inverted numerical sequence of the sequence previously considered.

**Figure 1b:**
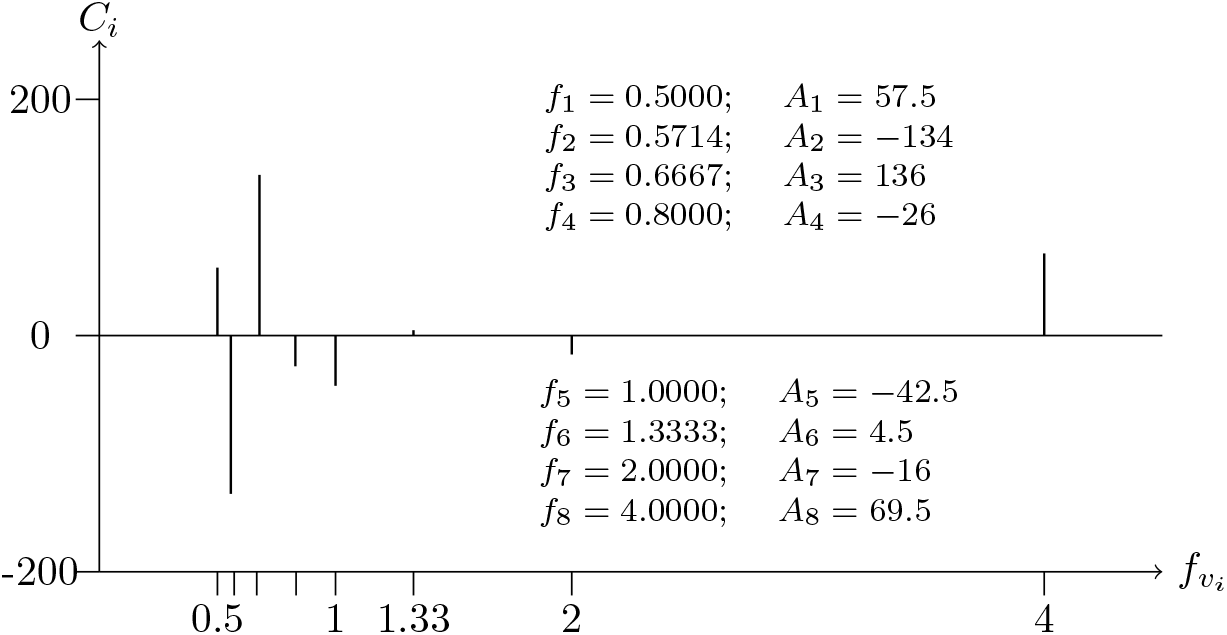
SWT which presents the results of the analysis with the SWM of the inverted numerical sequence of the numerical sequence previously considered

## 3 Presentation—Using the SWT—of the Results of the Analysis with the SWM, of Diverse Sequences of 1200 bases of Human DNA

In this section, consideration will be given to sequences of 1200 bases of human DNA.

To use the SWM to analyze diverse sequences of these bases, the first operation to be carried out is to assign numerical values to the different bases constituting those sequences. Researchers who use the SWM to analyze sequences of bases of DNA with that objective must assign the same numerical values to those bases so that their results will be easily comparable. The assignment of numerical values to the bases of DNA used in this article is the following:

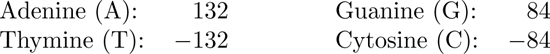

In these cases, given that the interval of reference is composed of 1200 elemental subunits, the first train of square waves considered has square waves of 2400 elemental subunits. Only half a square wave of this type “fills” that interval. Therefore, *f*_1200_ = 1200/2400 = ½.

Each square wave of the second train of square waves considered has 2 elemental subunits less than each square wave in the first train of square waves.

The third train of square waves has the following frequency: *f*_3_ = 1200/2398-2; and so on, until finally, *f*_1199_ = 1200/4 = 300, and *f*_1200_ = <u>^1200^</u> = 600. Of course, this last result specifies that for the train of square waves with the greatest frequency, 600 square waves, each of which is composed of 2 elemental subunits, “fit” in the interval of reference composed of 1200 elemental subunits. In other words, the “repetitive event” – the square wave – “occurs” 600 times in the interval of reference: *f*_1200_ = <u>^1200^</u> = 600.

Before presenting the figures corresponding to the results of the analyses of diverse sequences of 1200 bases of human DNA, certain terminological items should be clarified.

Every square wave of the type considered is composed of 2 semiwaves: one positive and the other negative. Suppose that a numerical value of 125 corresponds to the positive semiwave. In this case, the number *−*125 corresponds to the negative semiwave. It is accepted that the amplitude of that square wave is equal to the absolute value of *−*125: |125| = *|−* 125| = 125. It is also acceptable to state that the amplitude of the train of square waves composed of a sequence of that type of square waves is equal to 125.

The unknowns of the systems of linear algebraic equations which should be solved are denominated *A_i_*, for *i* = 1, 2, 3*, …,* 1200. Regardless of whichever of these *A_i_* turns out to be a positive number or a negative number, the amplitude of each square wave of the *i^th^*train of square waves is *|A_i_|*, the modulus of the numerical value of *|A_i_|*.

For clarity, in the graphs presented in this section using the SWT, the results will be displayed only partially. In effect, only the *A_i_* and the *f_i_* corresponding to the 10 greatest amplitudes of all of those *A_i_*, for *i* = 1, 2, 3*, …,* 1200, will be shown. (Of course, there is no problem in obtaining the complete results.)

For those *A_i_*and *f_i_* which will be displayed in the graphs in this section, they will be denominated “prominent *A_i_*” and “prominent *f_i_*” respectively.

In the caption of each figure in this section, beginning with 2a and 2b, inclusive, the following will be indicated: 1) the source of the information of the data analyzed with the SWM and graphed with the SWT; 2) the number of chromosome to which the sequence of bases considered belongs; 3) the characterization of the chromosomic region considered; and 4) the specification of the sequence of bases analyzed.

**Figure 2a:**
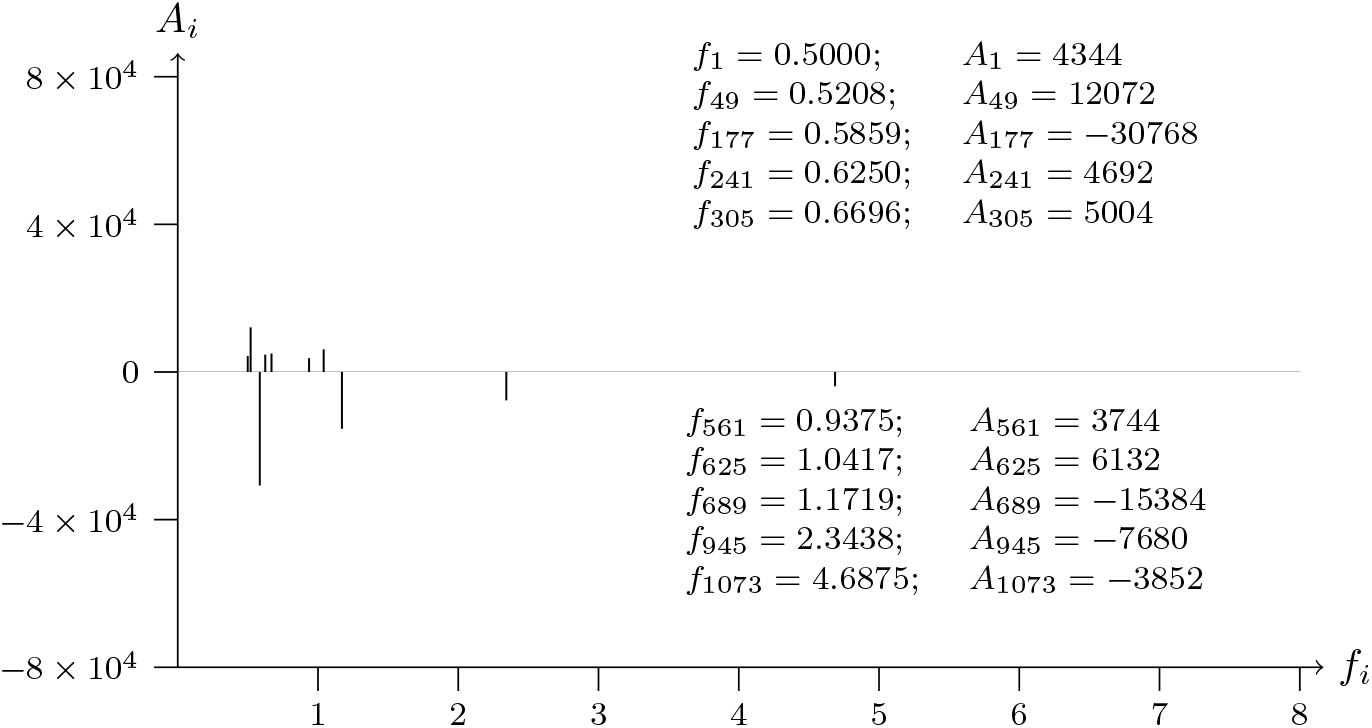
T2T-CHM13v2.0 chromosome 1; p telomere; analyzed data from 1 to 1200

**Figure 2b:**
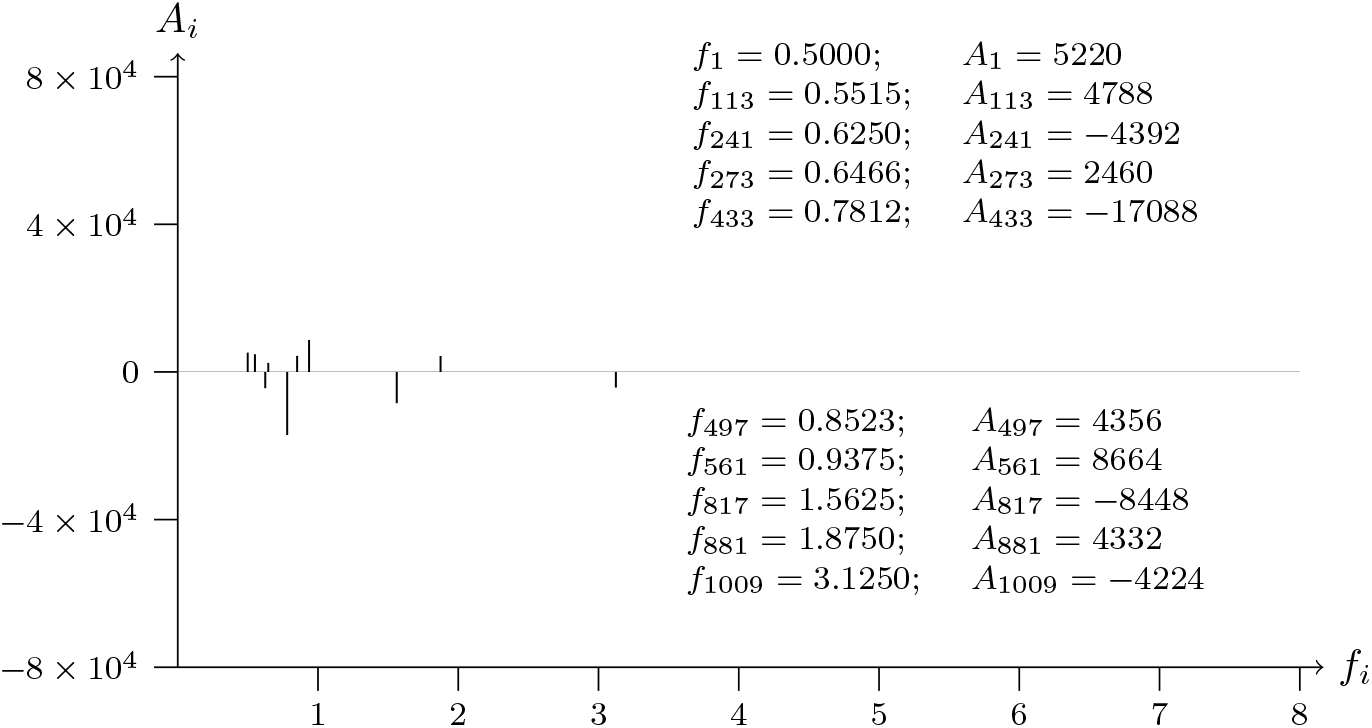
T2T-CHM13v2.0 chromosome 1; p telomere; analyzed data from 1200 to 1

**Figure 3a:**
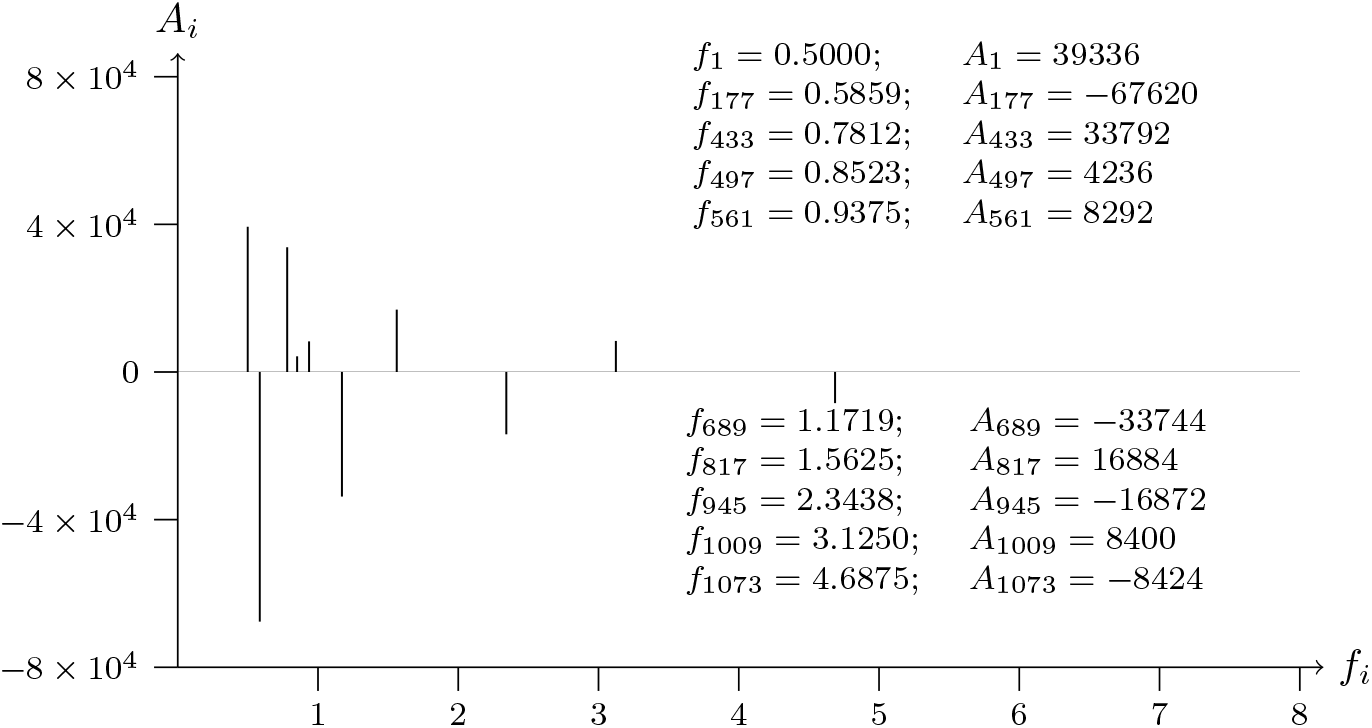
T2T-CHM13v2.0 chromosome 2; p telomere; analyzed data from 1 to 1200

**Figure 3b:**
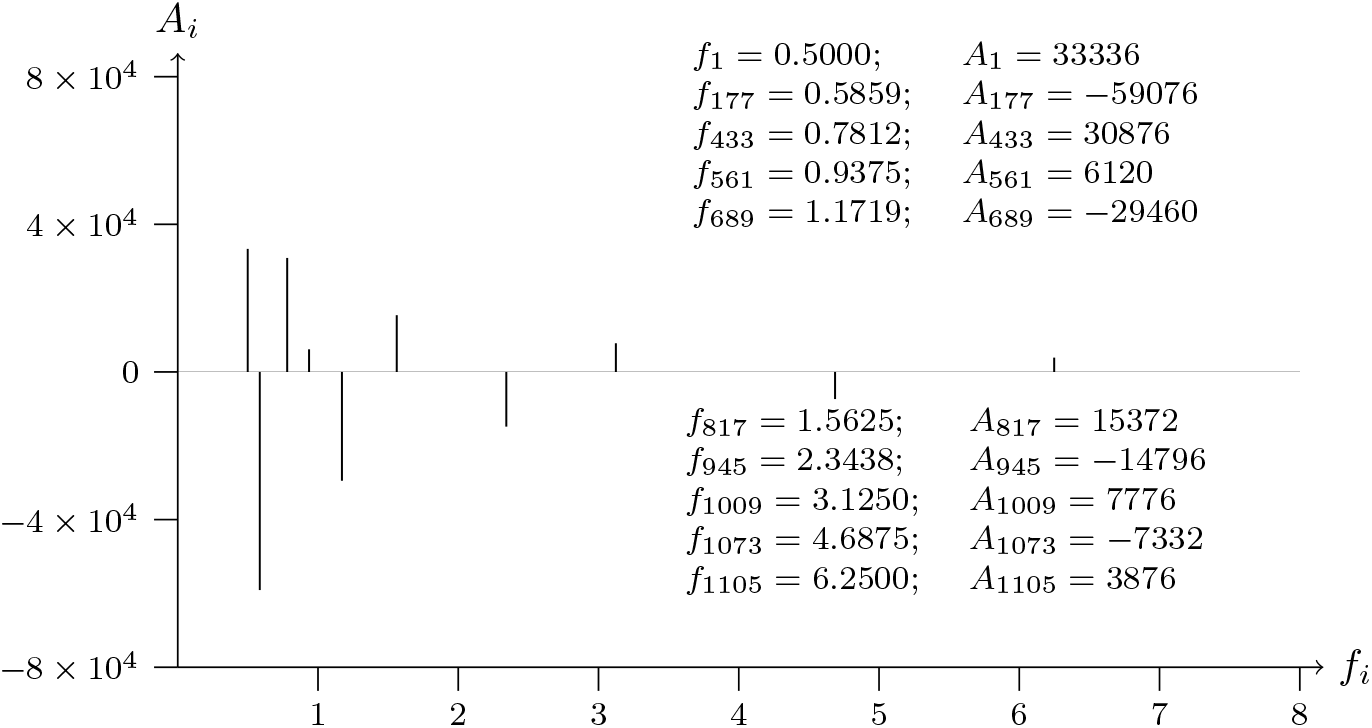
T2T-CHM13v2.0 chromosome 2; p telomere; analyzed data from 1200 to 1

**Figure 4a:**
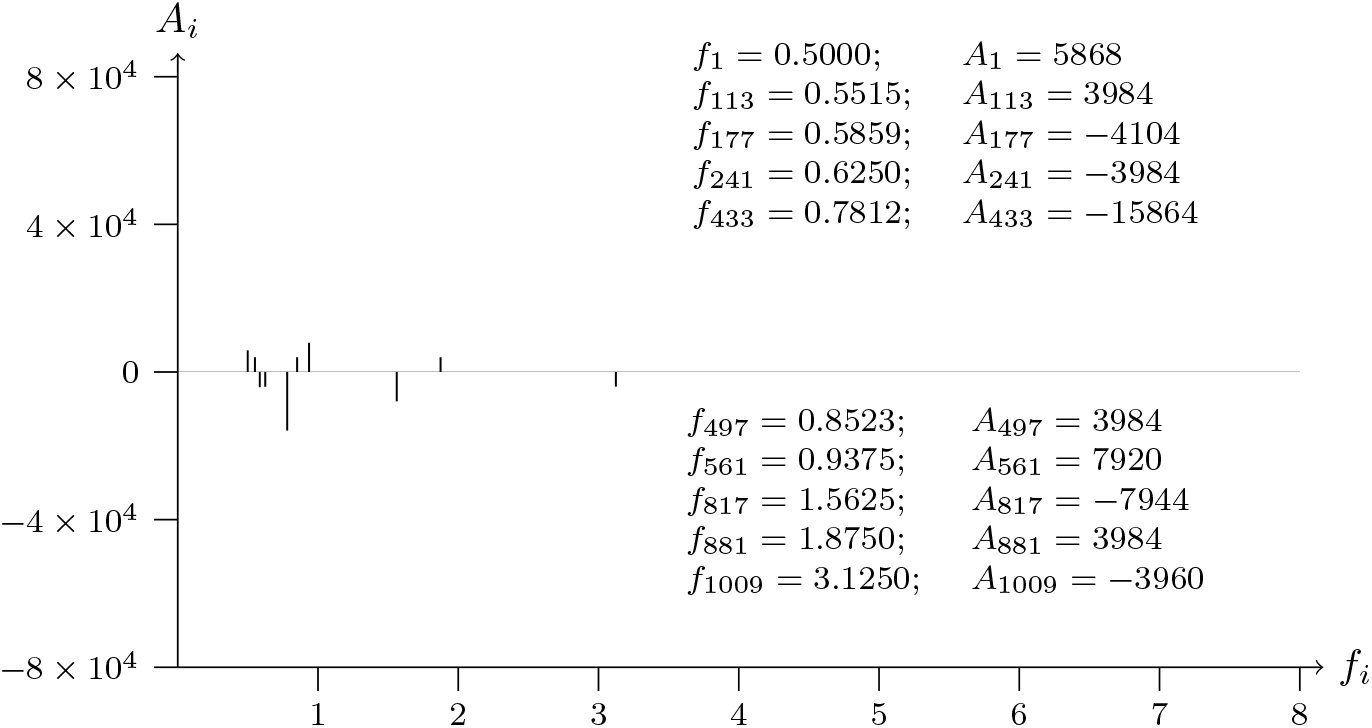
T2T-CHM13v2.0 chromosome 3; p telomere; analyzed data from 1 to 1200

**Figure 4b:**
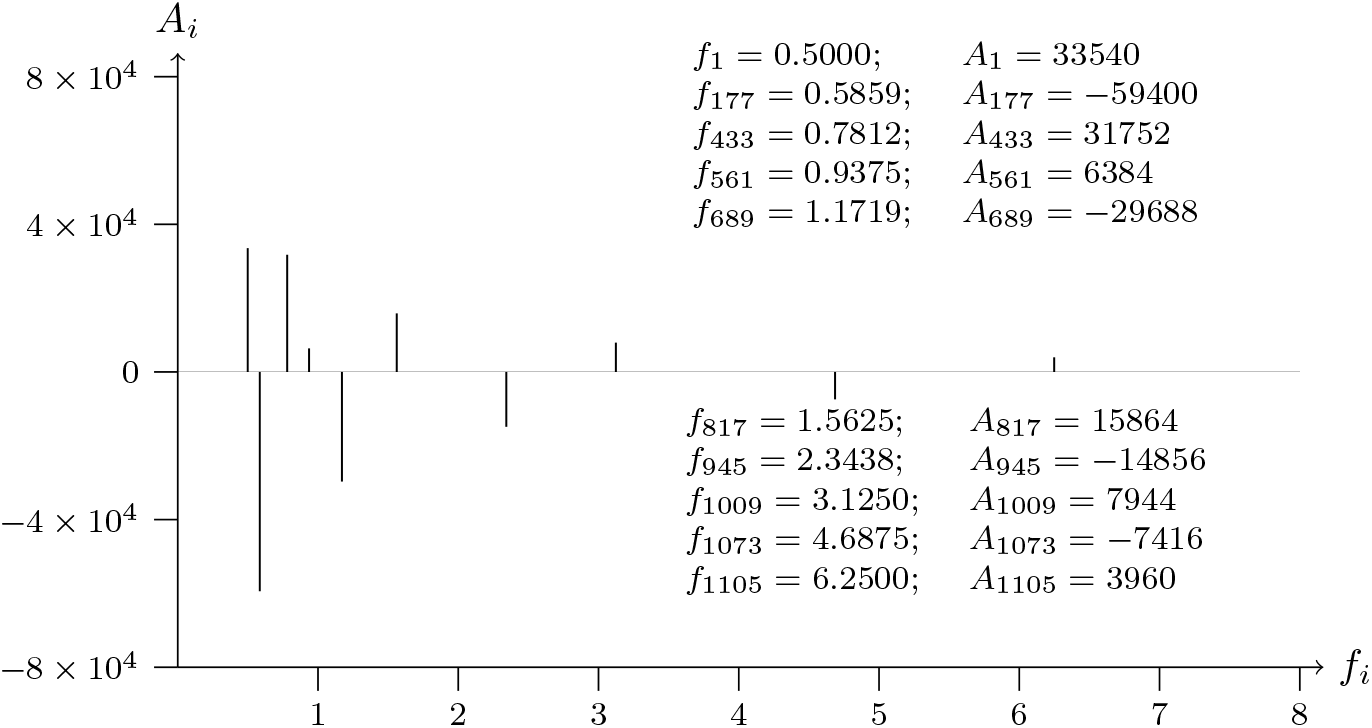
T2T-CHM13v2.0 chromosome 3; p telomere; analyzed data from 1200 to 1

**Figure 5a:**
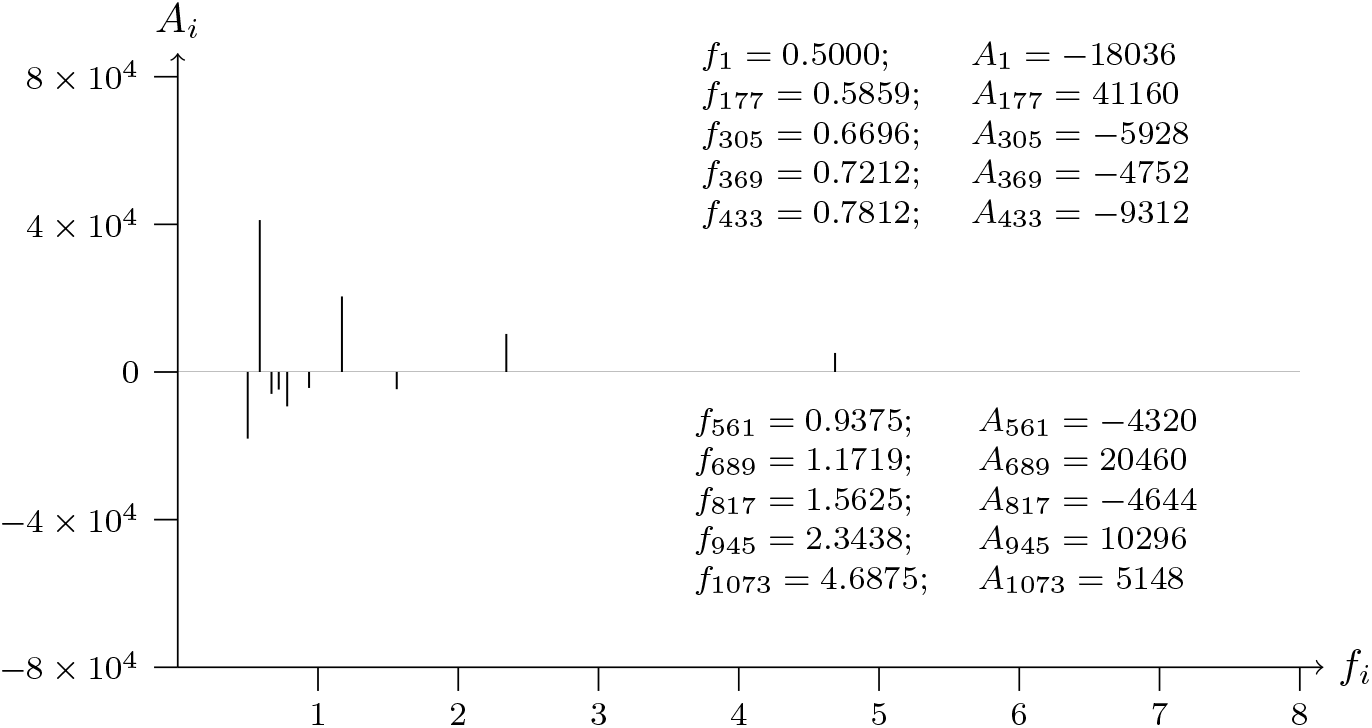
T2T-CHM13v2.0 chromosome 4; p telomere; analyzed data from 1 to 1200

**Figure 5b:**
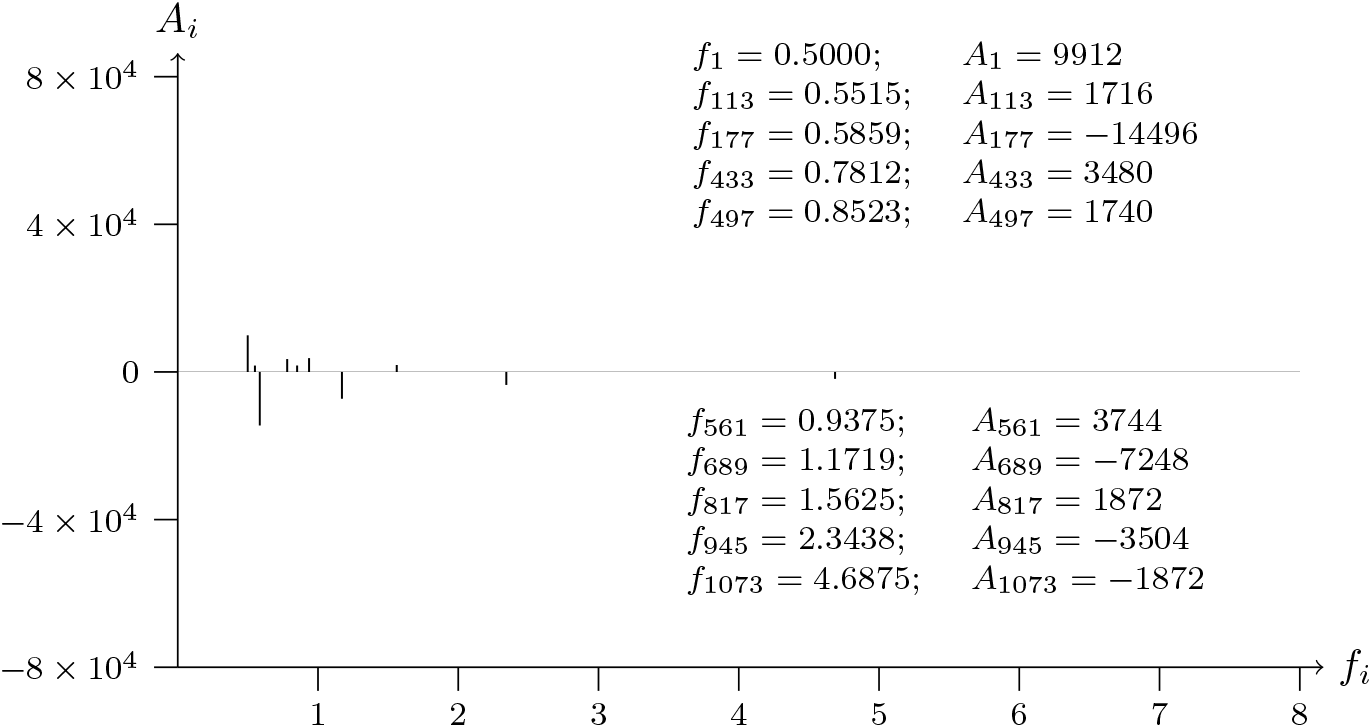
T2T-CHM13v2.0 chromosome 4; p telomere; analyzed data from 1200 to 1

**Figure 6a:**
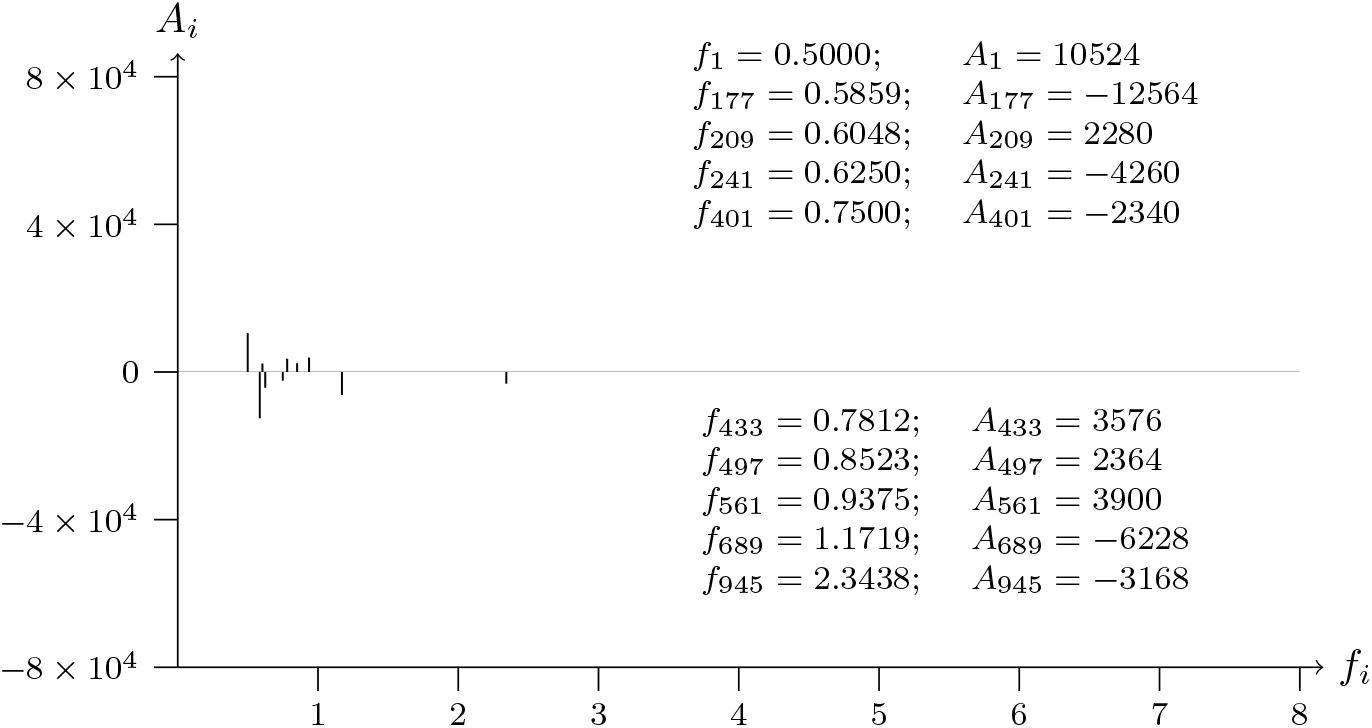
T2T-CHM13v2.0 chromosome 5; p telomere; analyzed data from 1 to 1200

**Figure 6b:**
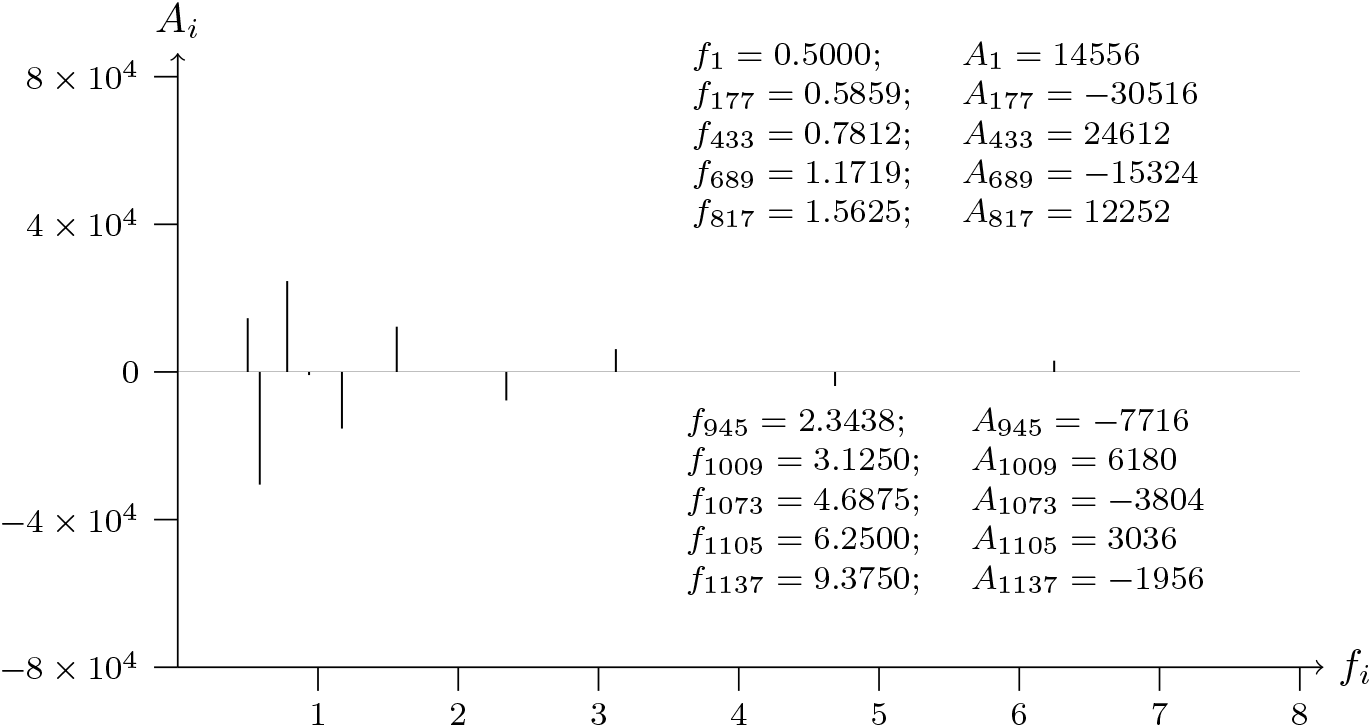
T2T-CHM13v2.0 chromosome 5; p telomere; analyzed data from 1200 to 1

**Figure 7a:**
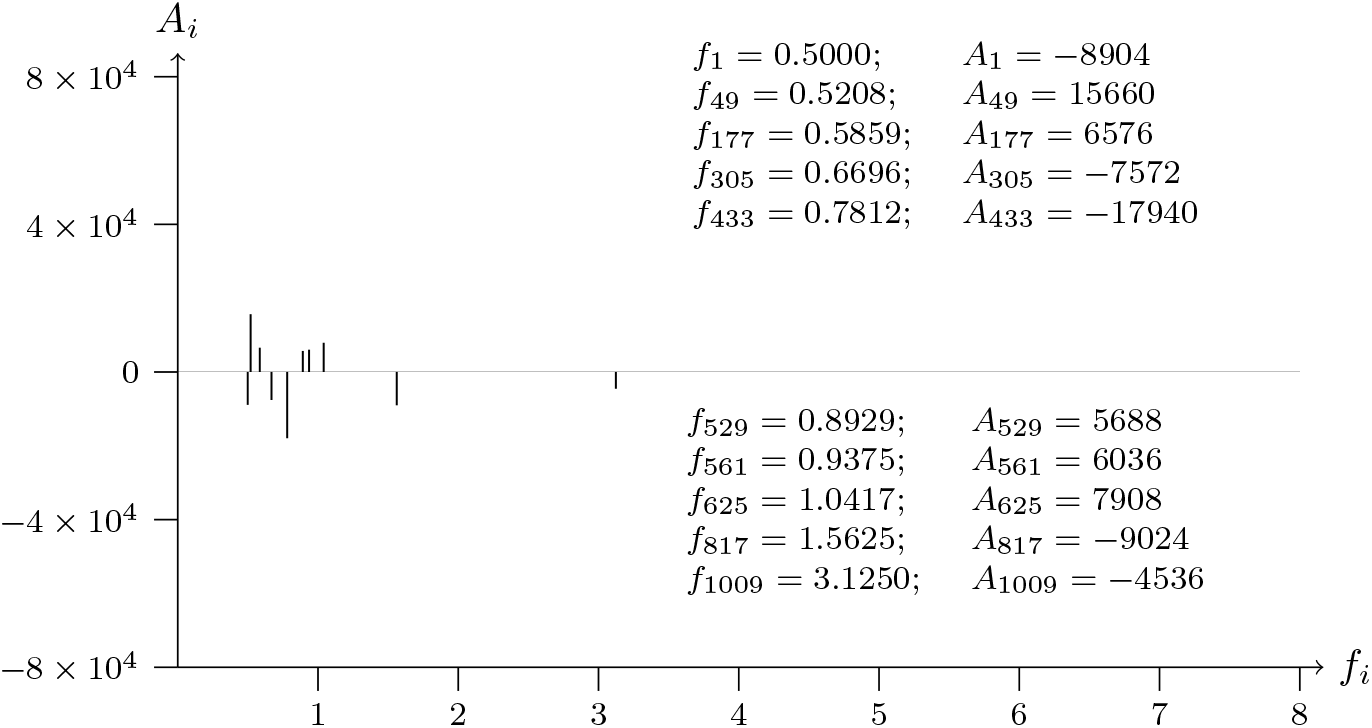
T2T-CHM13v2.0 chromosome 6; p telomere; analyzed data from 1 to 1200

**Figure 7b:**
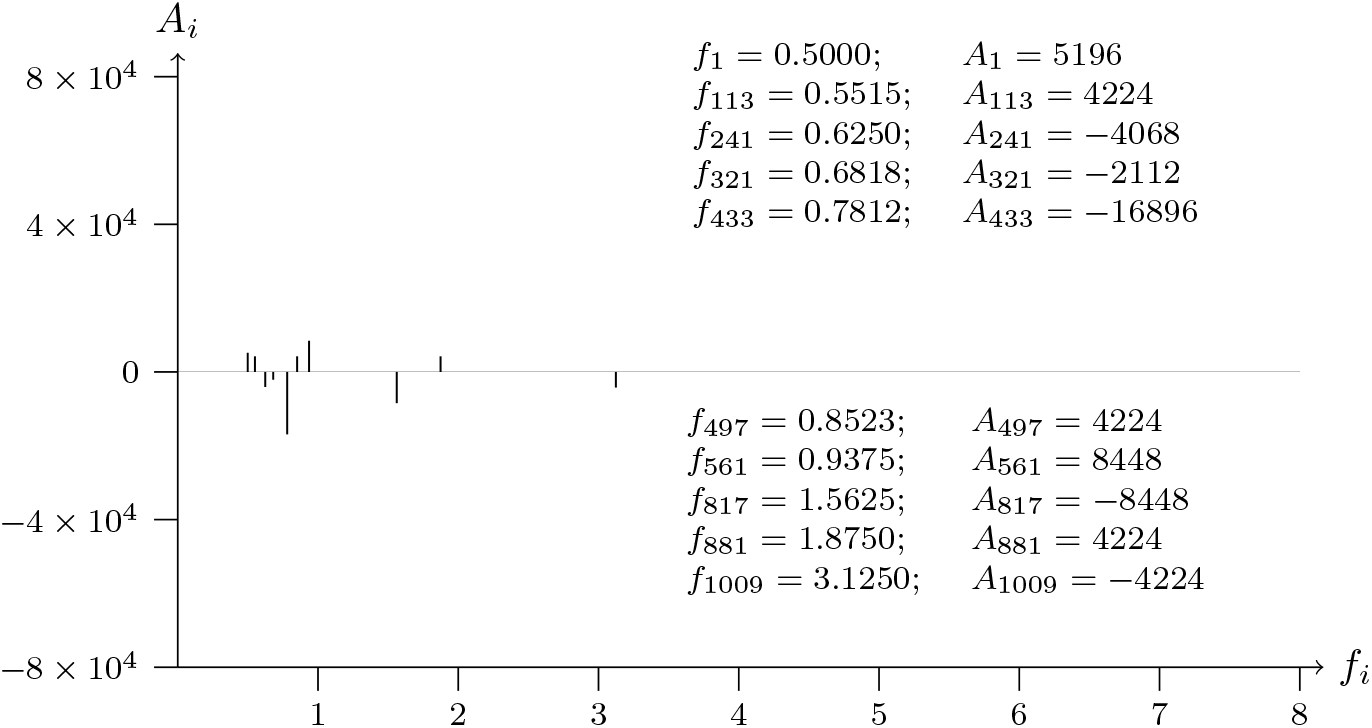
T2T-CHM13v2.0 chromosome 6; p telomere; analyzed data from 1200 to 1

**Figure 8a:**
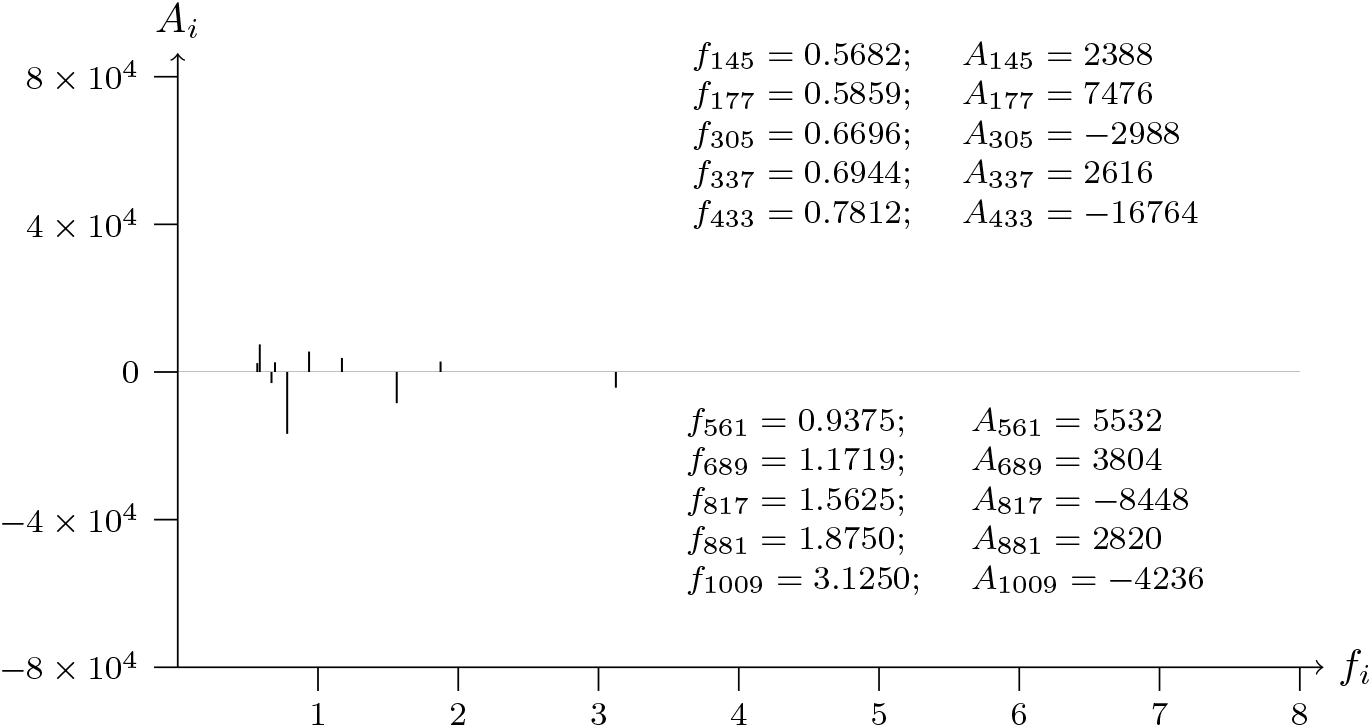
T2T-CHM13v2.0 chromosome 7; p telomere; analyzed data from 1 to 1200

**Figure 8b:**
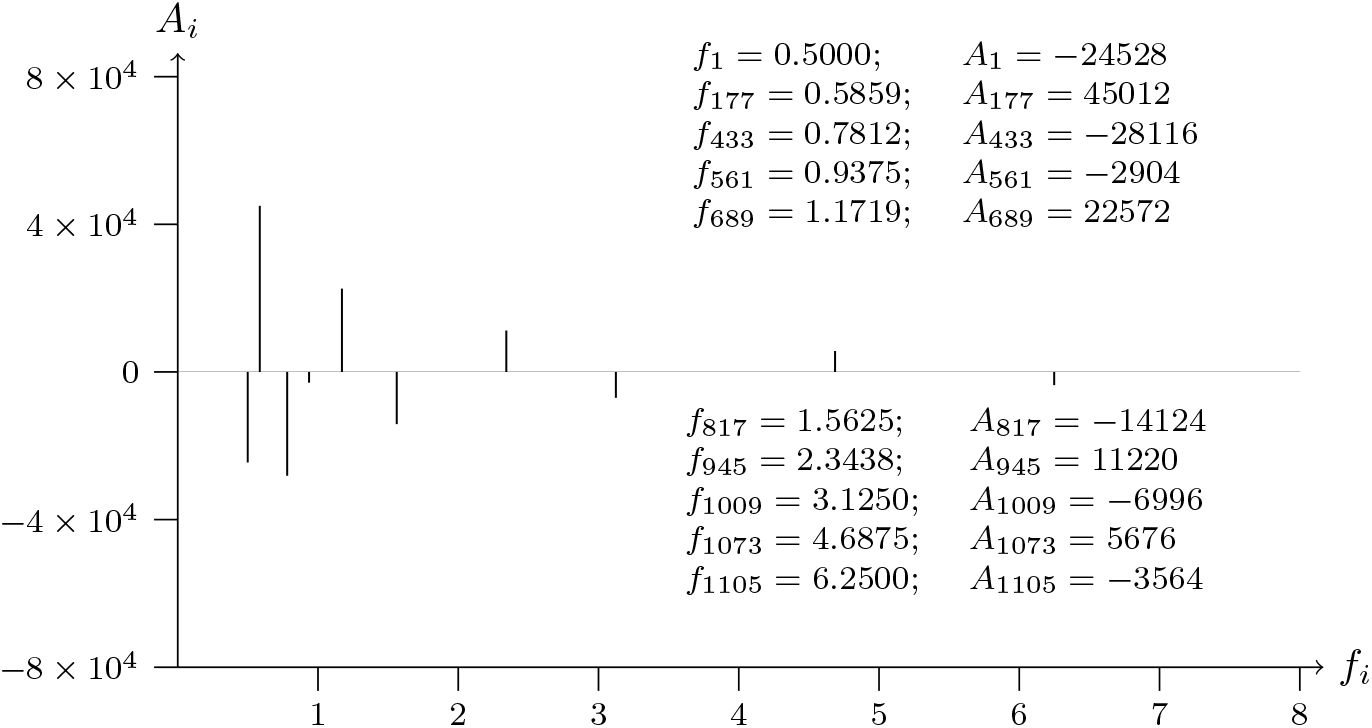
T2T-CHM13v2.0 chromosome 7; p telomere; analyzed data from 1200 to 1

**Figure 9a:**
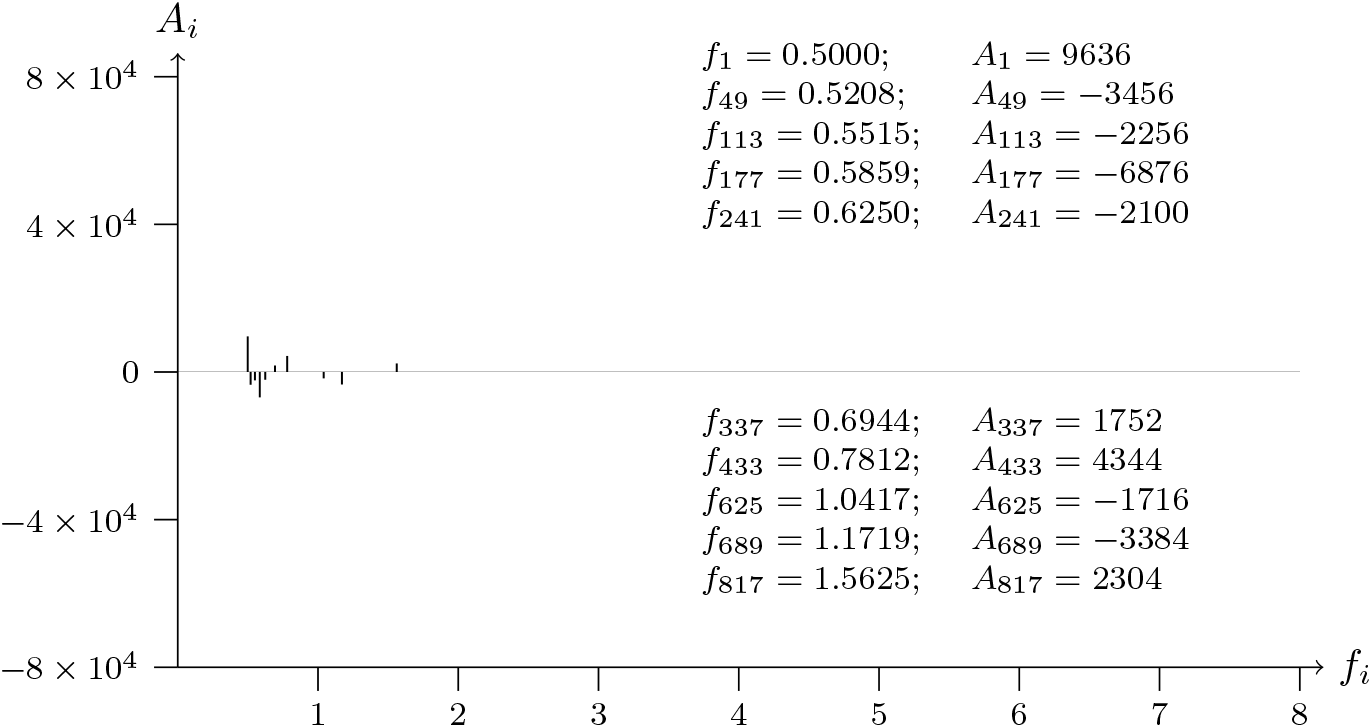
T2T-CHM13v2.0 chromosome 8; p telomere; analyzed data from 1 to 1200

**Figure 9b:**
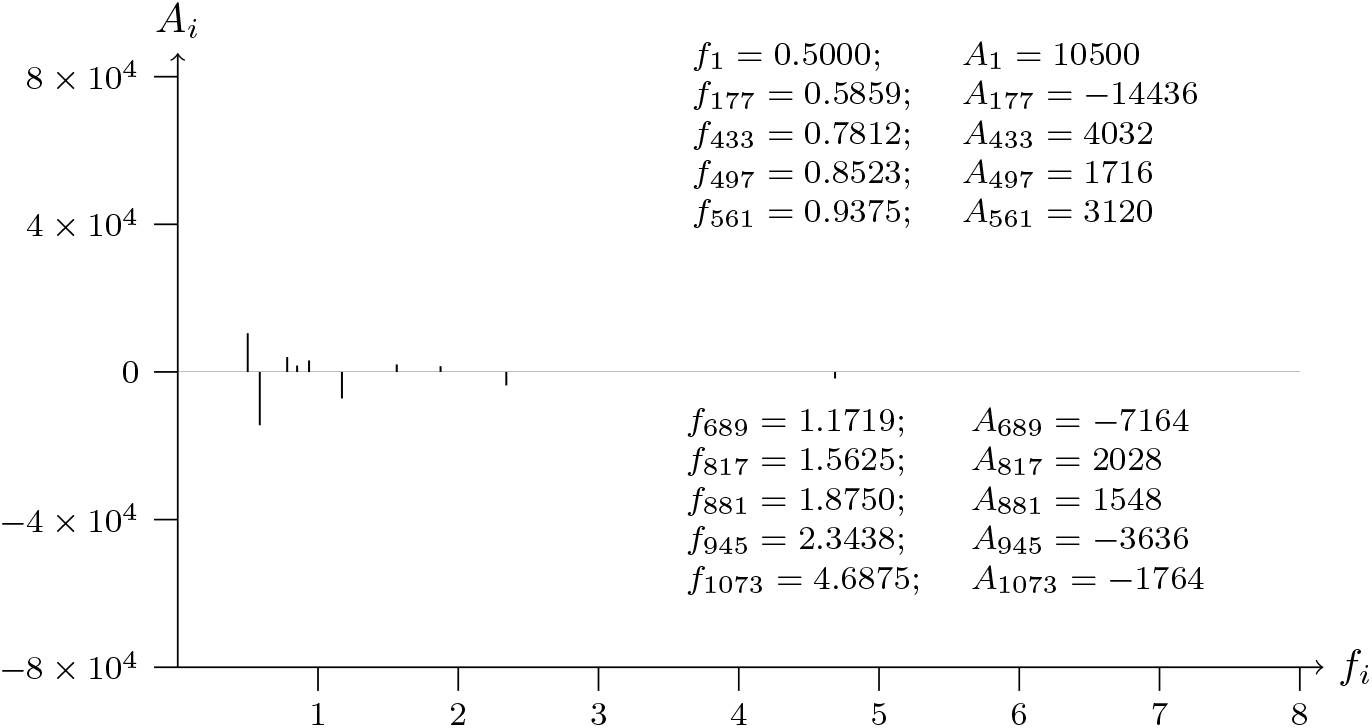
T2T-CHM13v2.0 chromosome 8; p telomere; analyzed data from 1200 to 1

**Figure 10a:**
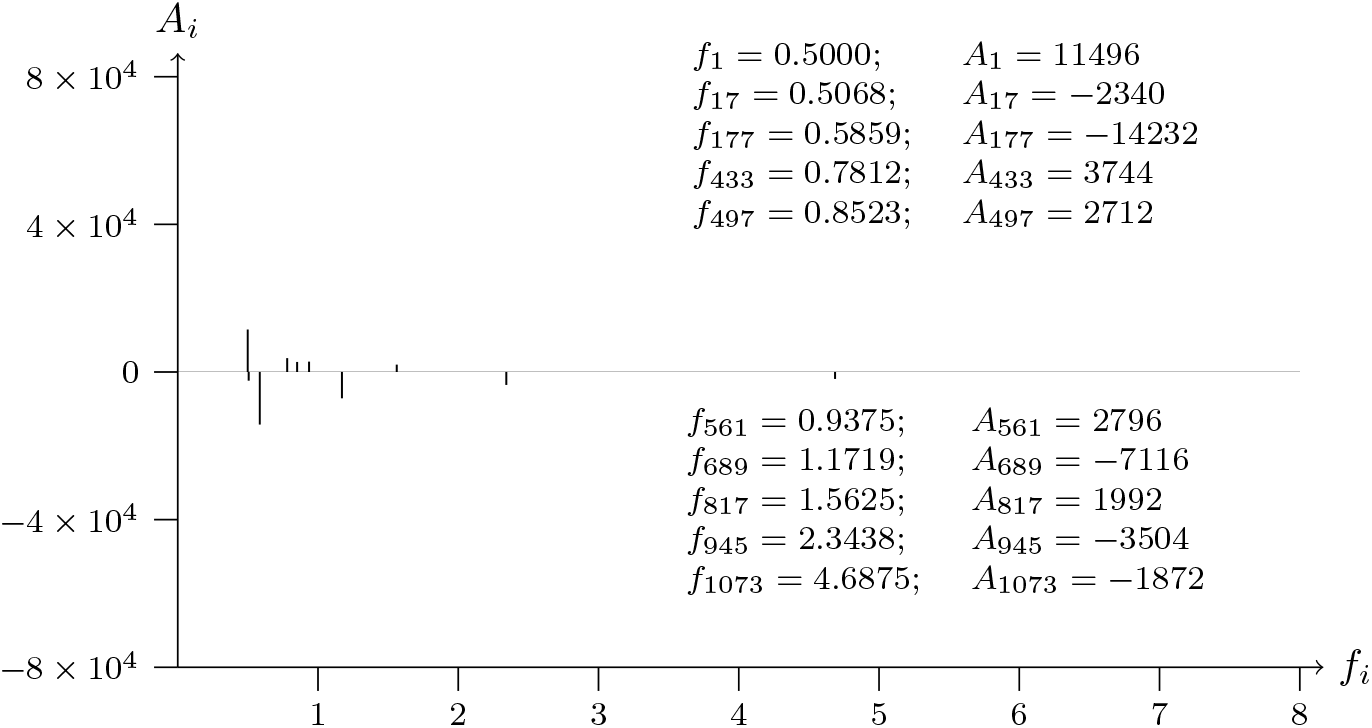
T2T-CHM13v2.0 chromosome 9; p telomere; analyzed data from 1 to 1200

**Figure 10b:**
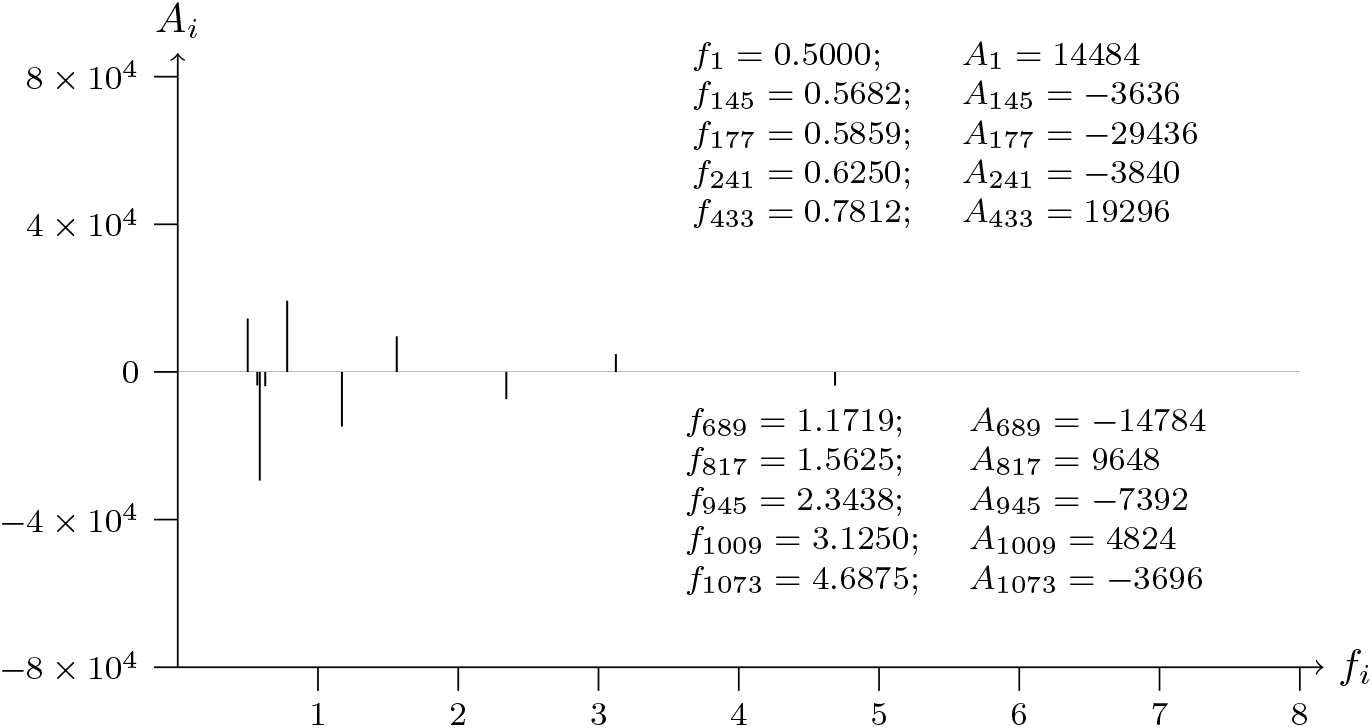
T2T-CHM13v2.0 chromosome 9; p telomere; analyzed data from 1200 to 1

**Figure 11a:**
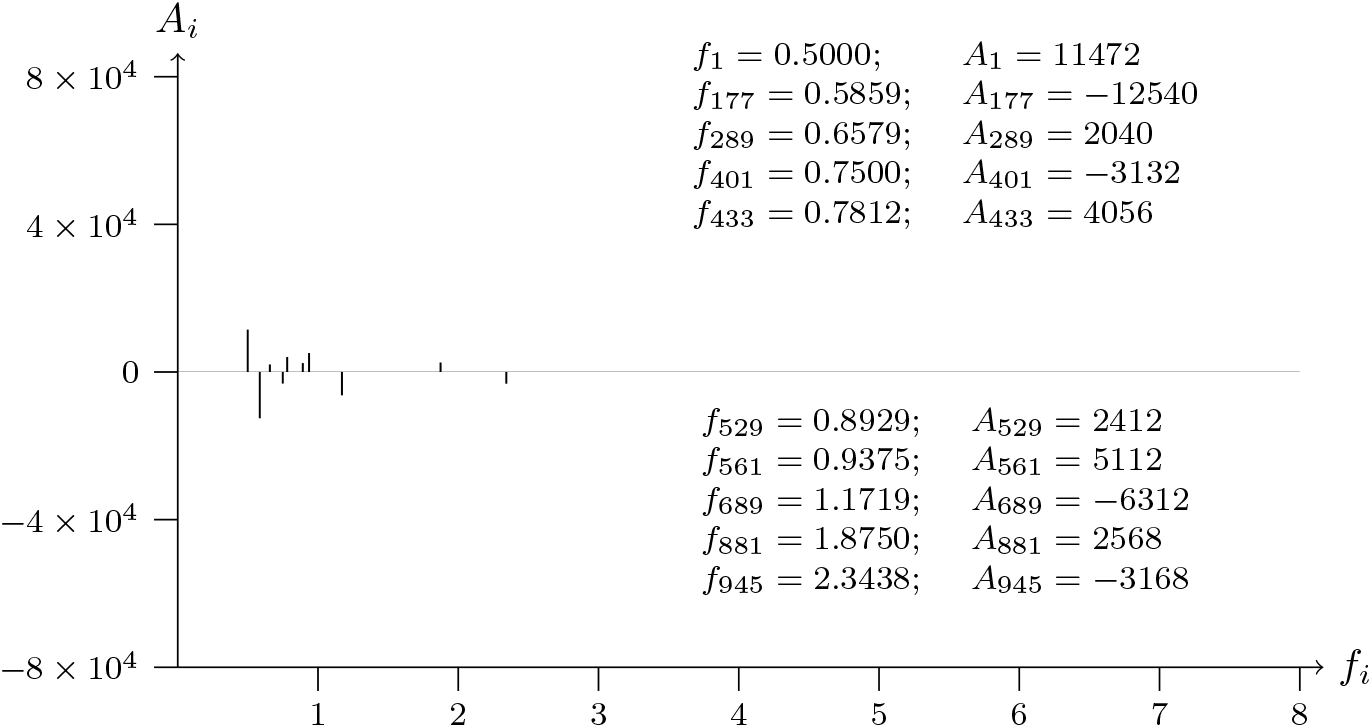
T2T-CHM13v2.0 chromosome 10; p telomere; analyzed data from 1 to 1200

**Figure 11b:**
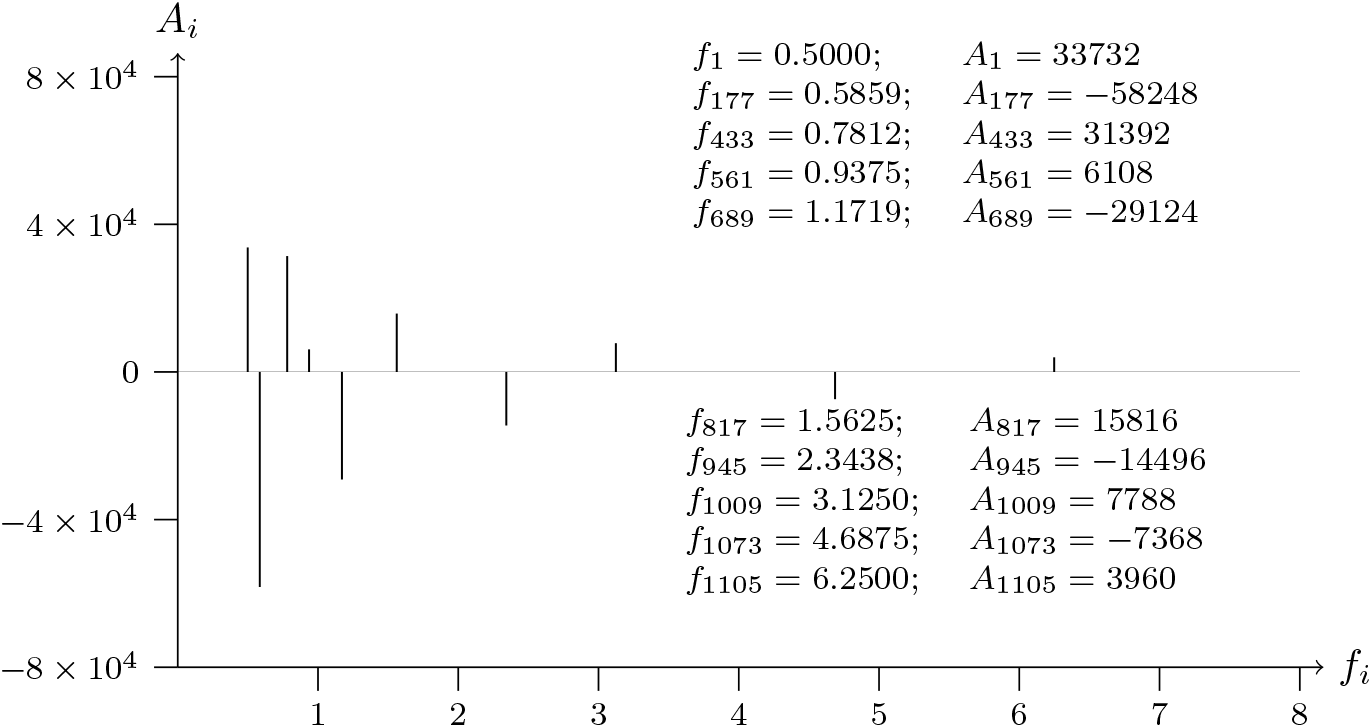
T2T-CHM13v2.0 chromosome 10; p telomere; analyzed data from 1200 to 1

**Figure 12a:**
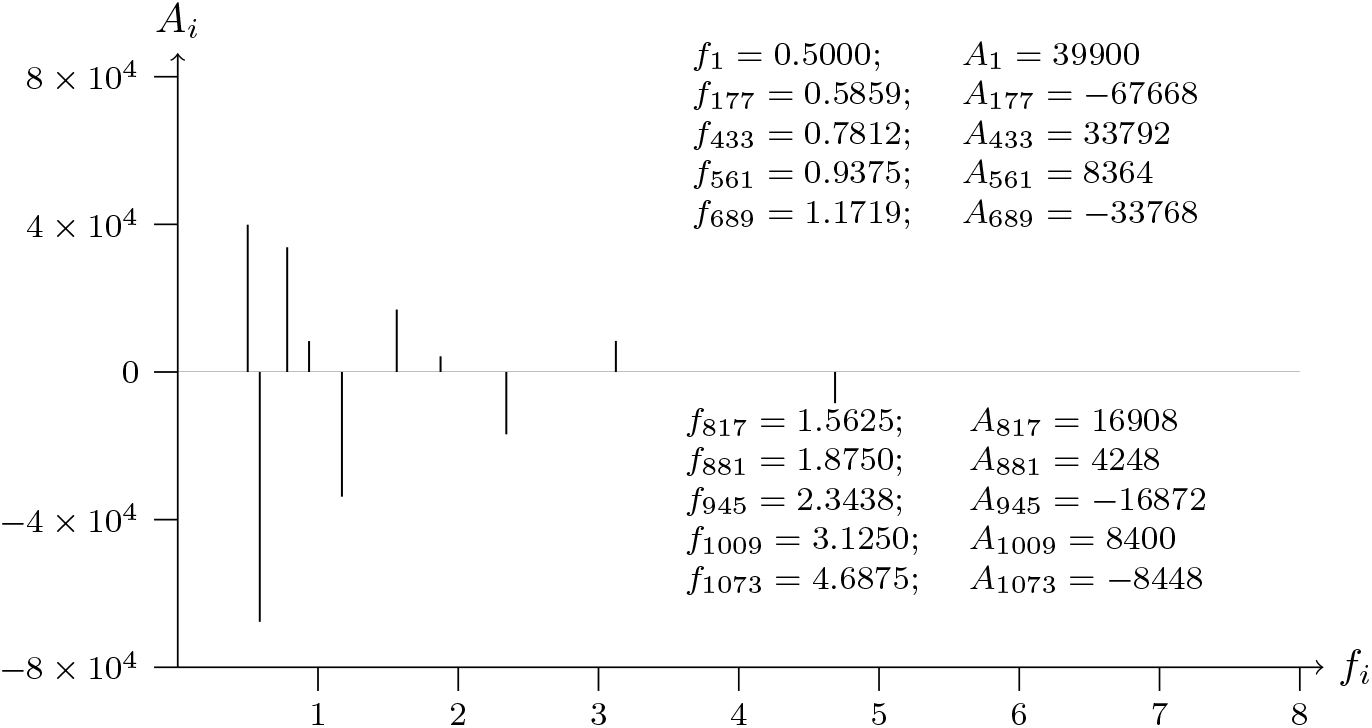
T2T-CHM13v2.0 chromosome 11; p telomere; analyzed data from 1 to 1200

**Figure 12b:**
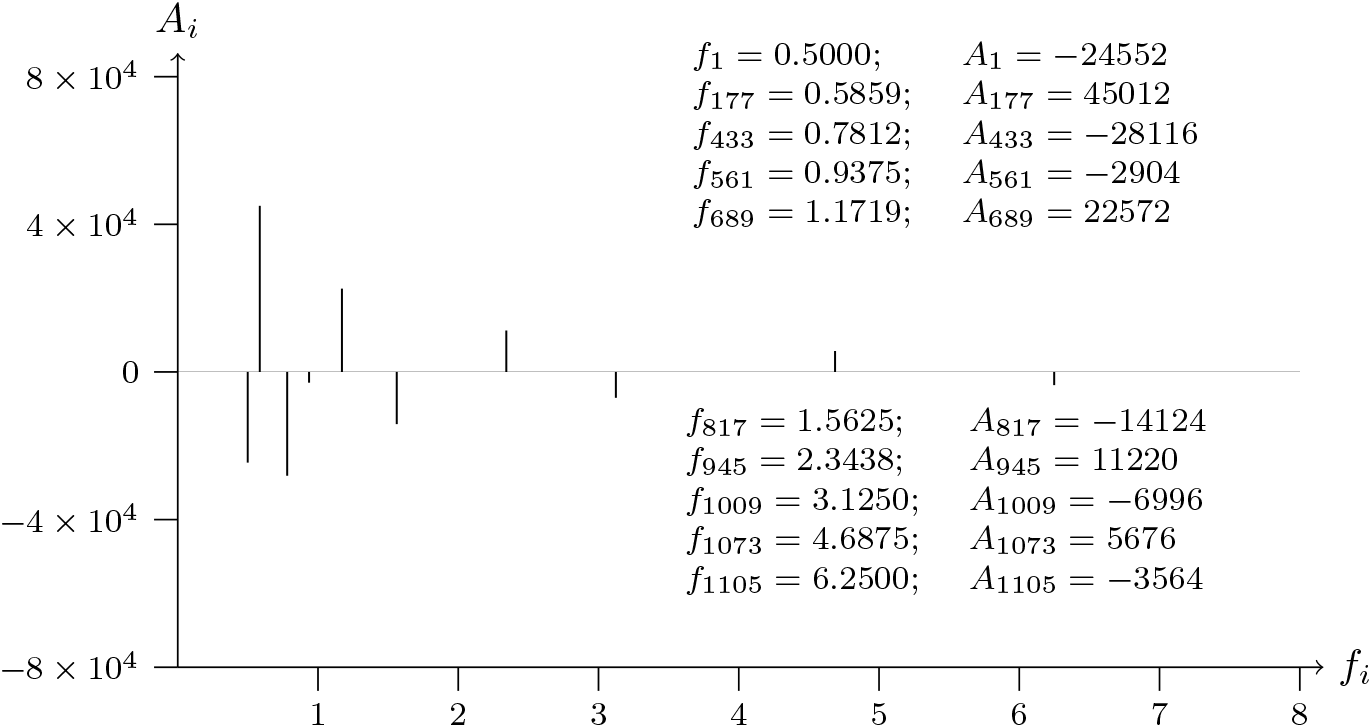
T2T-CHM13v2.0 chromosome 11; p telomere; analyzed data from 1200 to 1

**Figure 13a:**
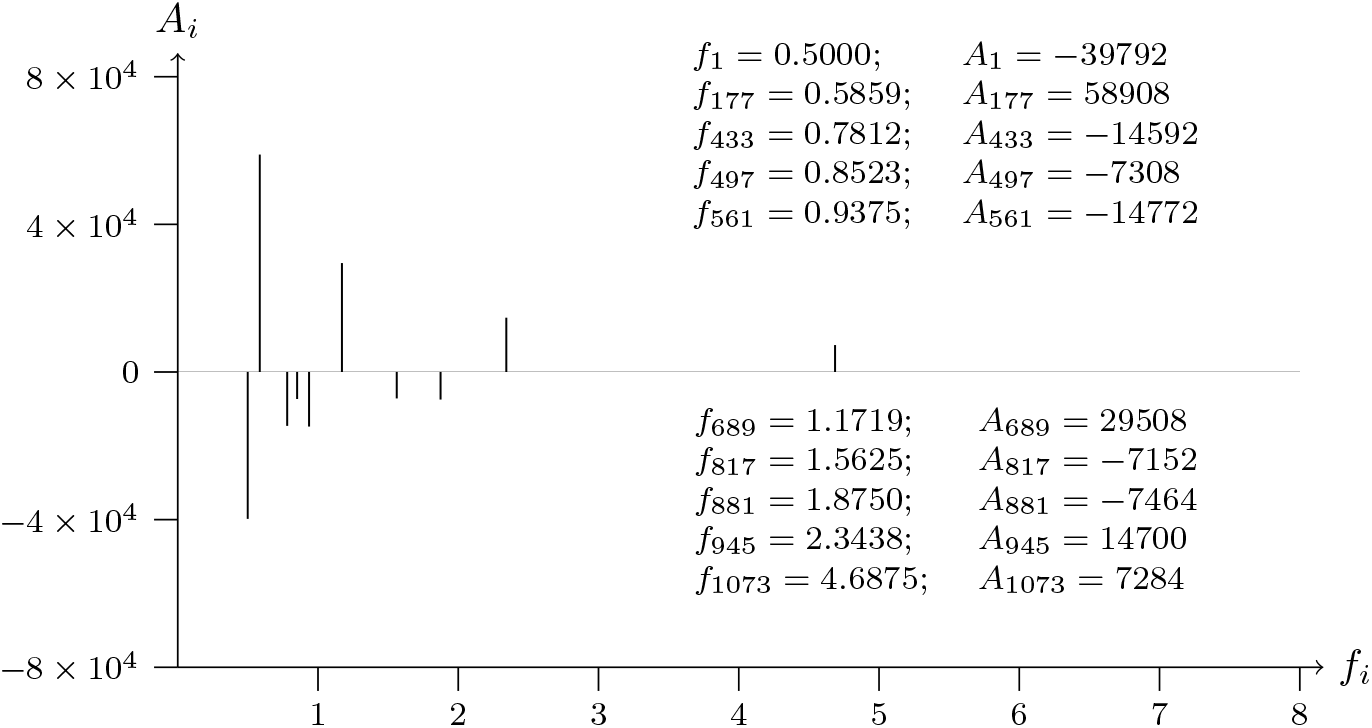
T2T-CHM13v2.0 chromosome 12; p telomere; analyzed data from 1 to 1200

**Figure 13b:**
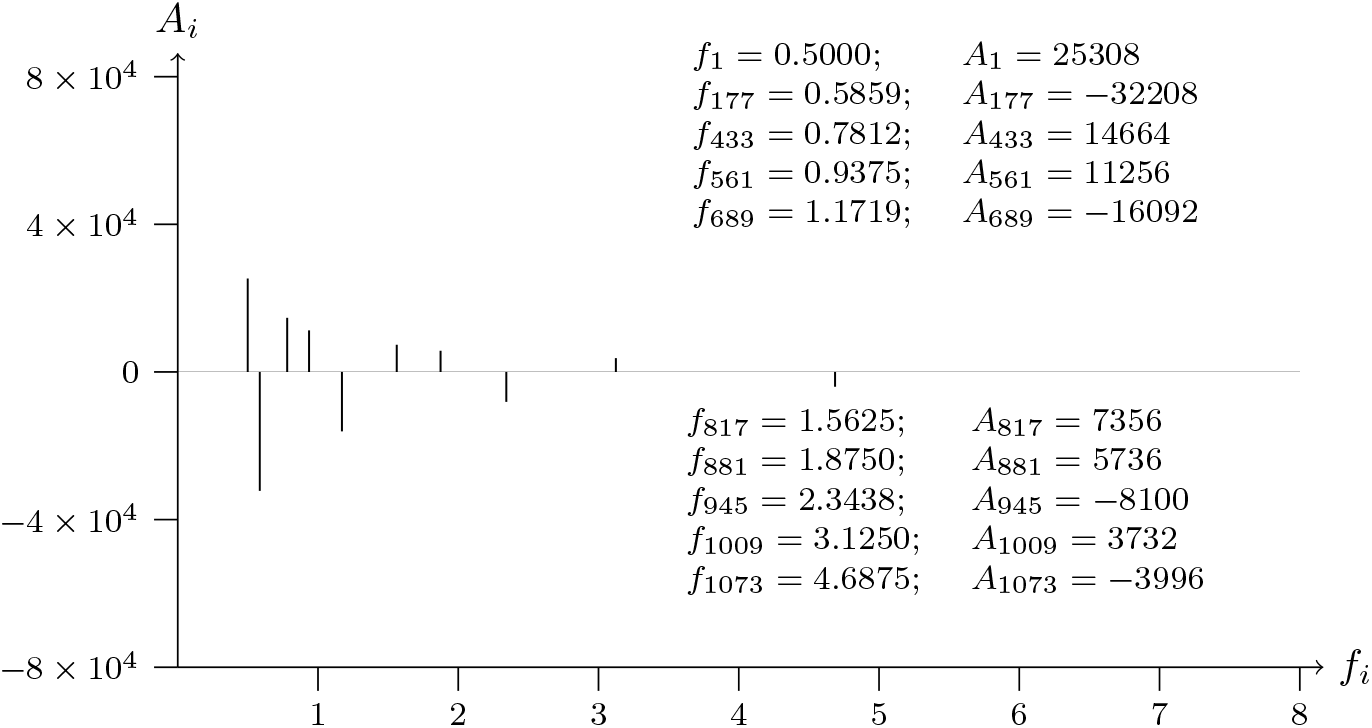
T2T-CHM13v2.0 chromosome 12; p telomere; analyzed data from 1200 to 1

**Figure 14a:**
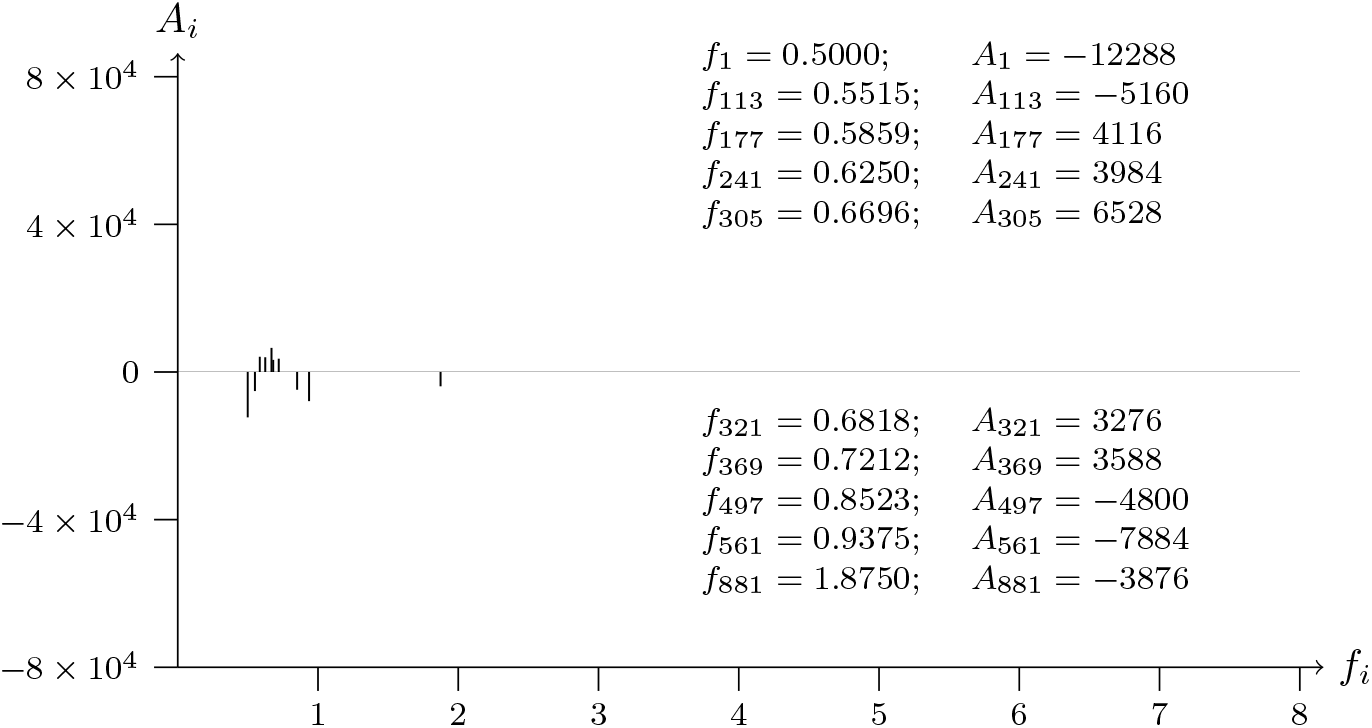
T2T-CHM13v2.0 chromosome 13; p telomere; analyzed data from 1 to 1200

**Figure 14b:**
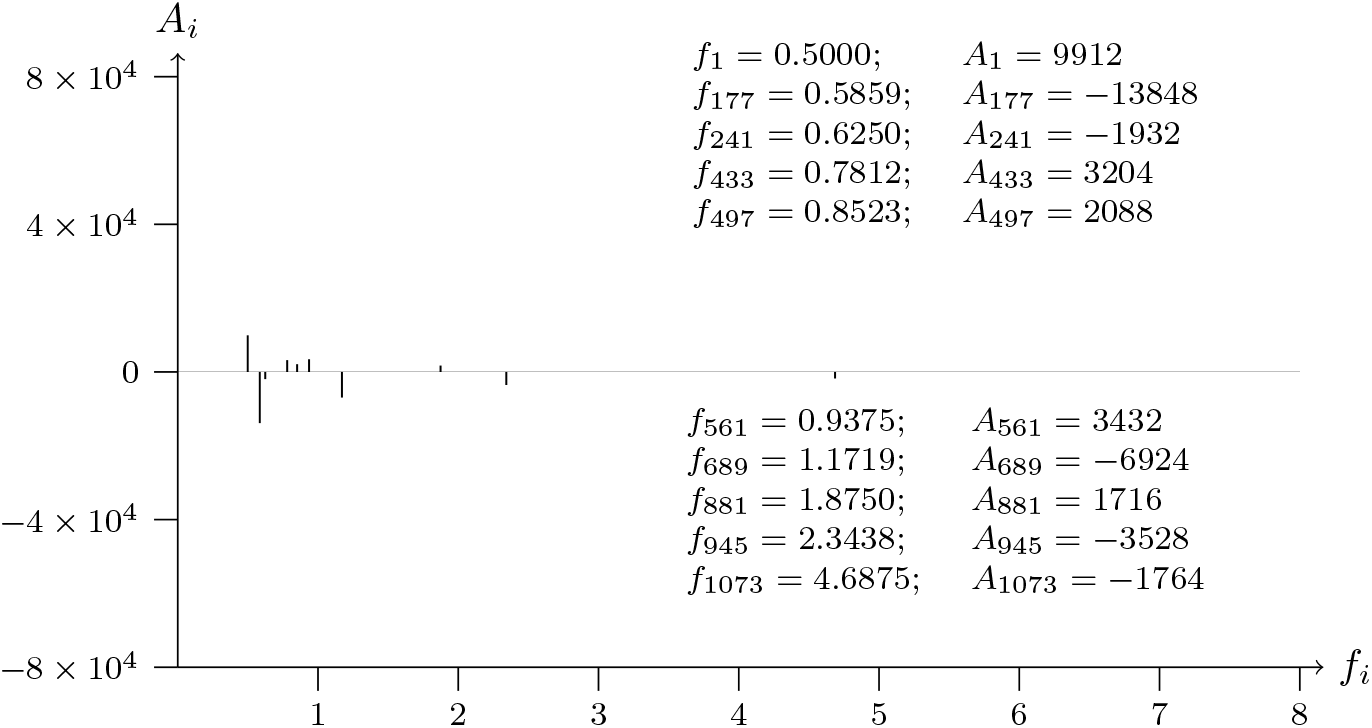
T2T-CHM13v2.0 chromosome 13; p telomere; analyzed data from 1200 to 1

**Figure 15a:**
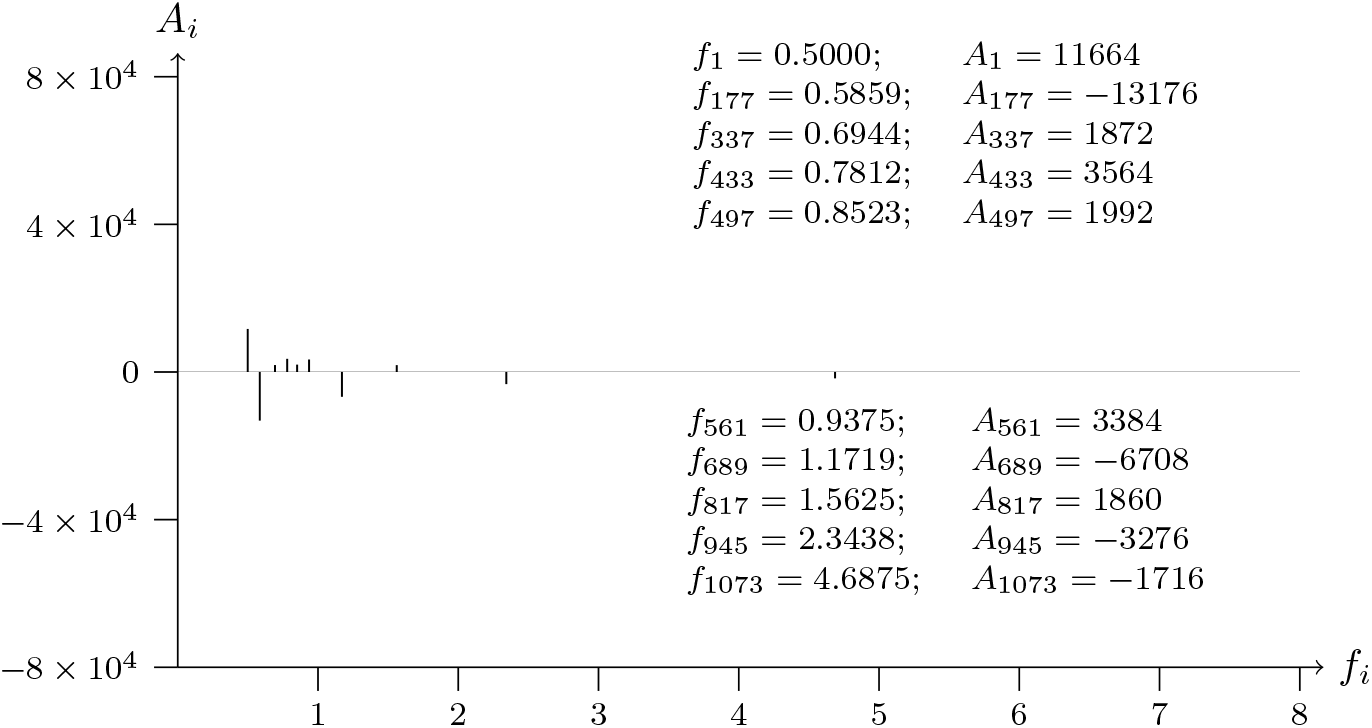
T2T-CHM13v2.0 chromosome 14; p telomere; analyzed data from 1 to 1200

**Figure 15b:**
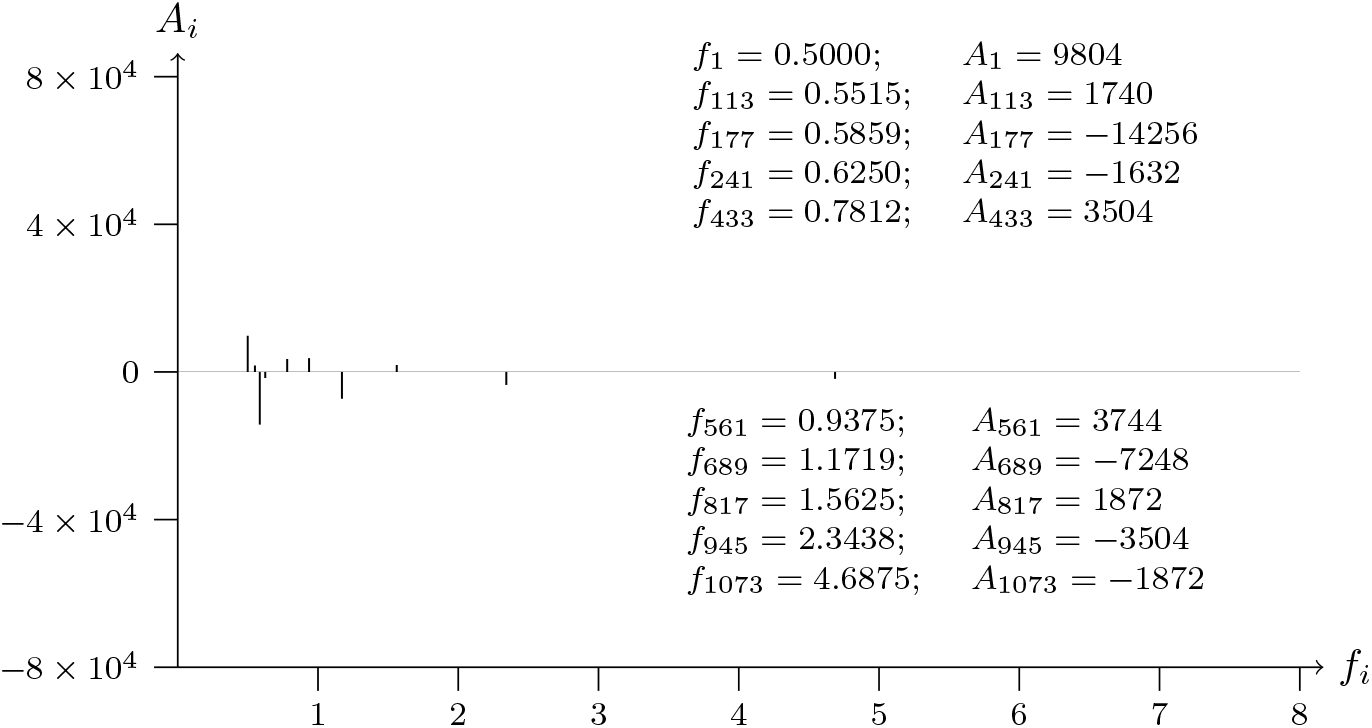
T2T-CHM13v2.0 chromosome 14; p telomere; analyzed data from 1200 to 1

**Figure 16a:**
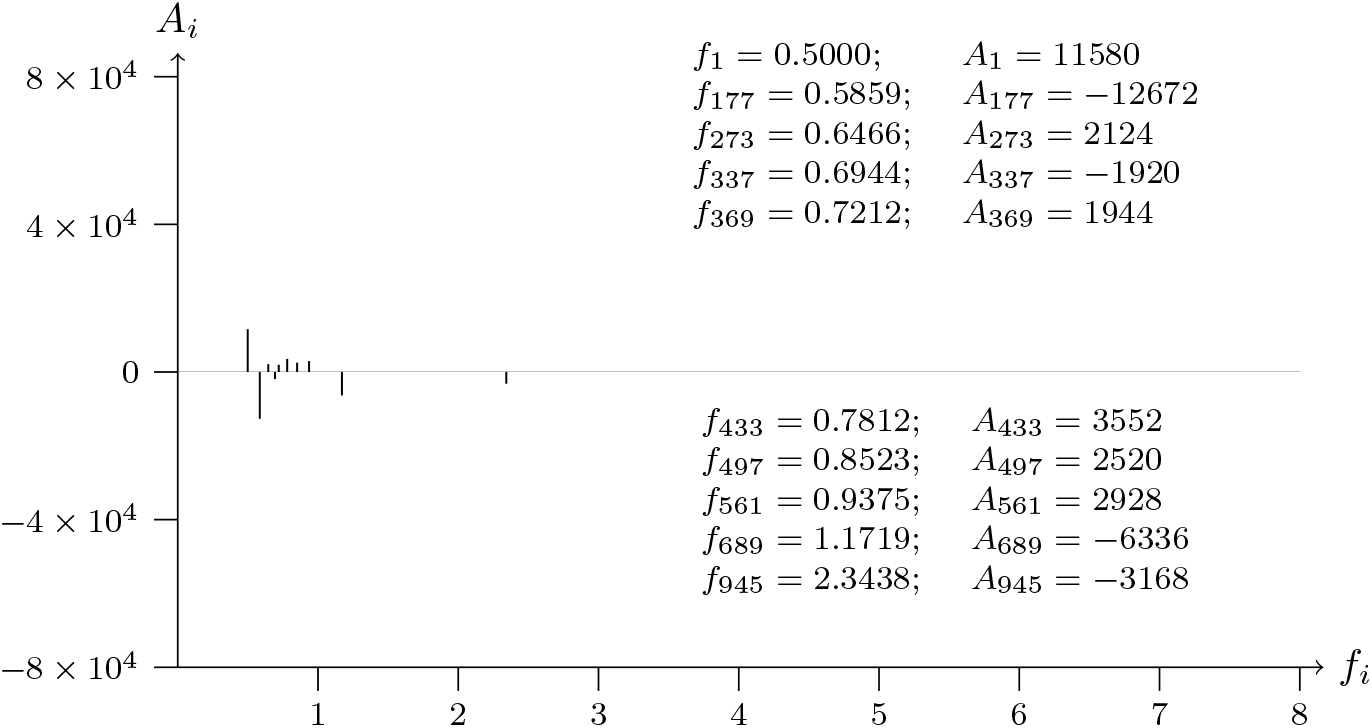
T2T-CHM13v2.0 chromosome 15; p telomere; analyzed data from 1 to 1200

**Figure 16b:**
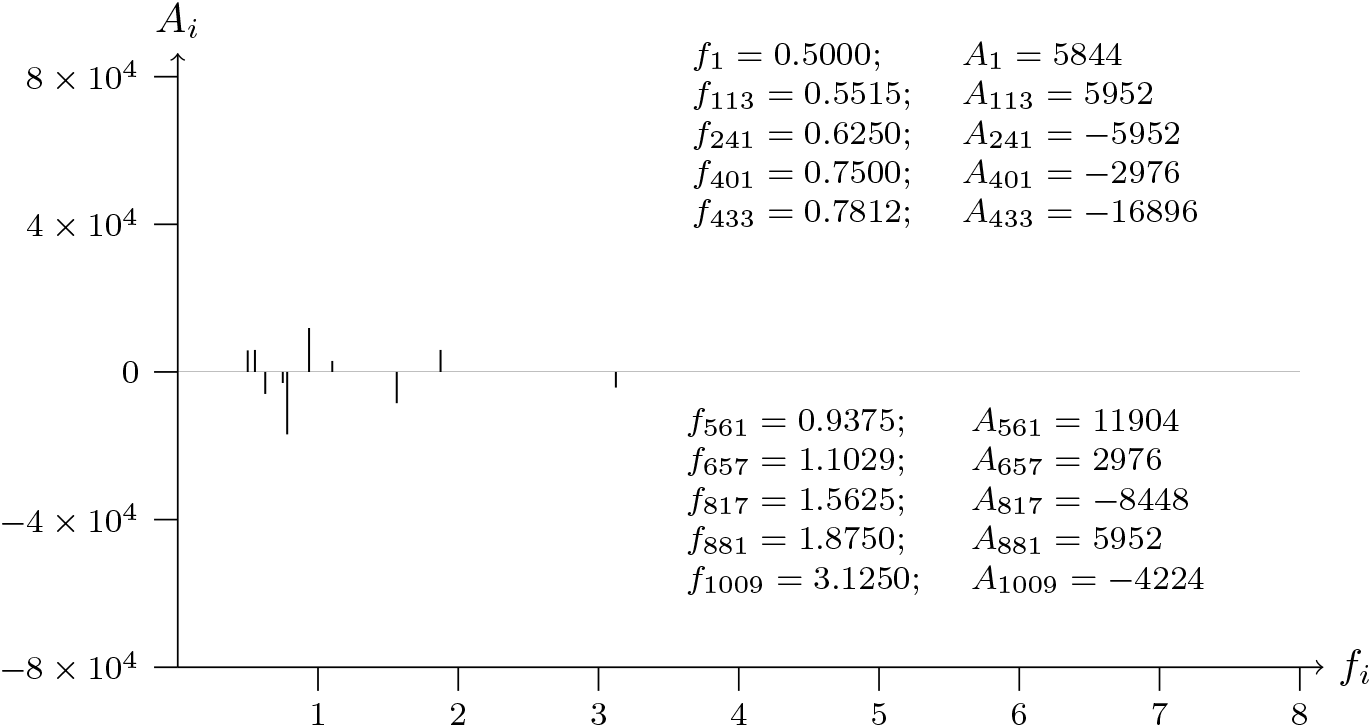
T2T-CHM13v2.0 chromosome 15; p telomere; analyzed data from 1200 to 1

**Figure 17a:**
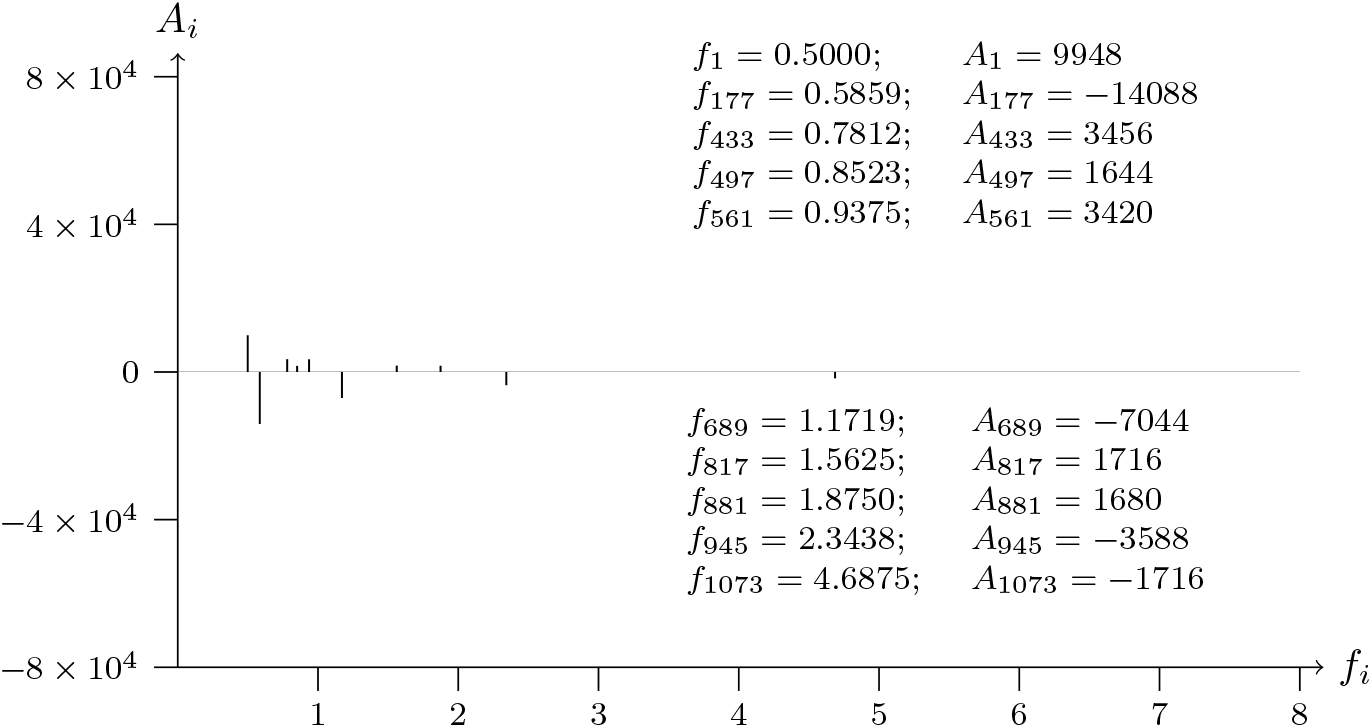
T2T-CHM13v2.0 chromosome 16; p telomere; analyzed data from 1 to 1200

**Figure 17b:**
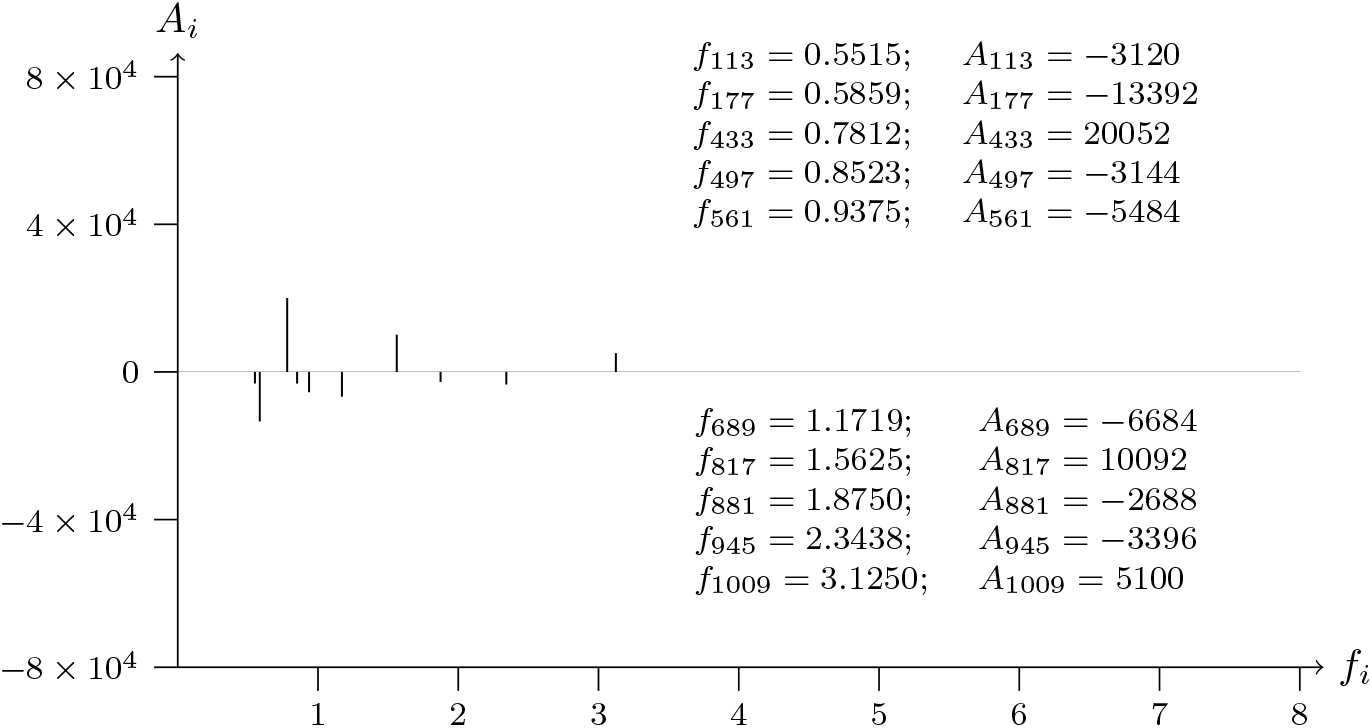
T2T-CHM13v2.0 chromosome 16; p telomere; analyzed data from 1200 to 1

**Figure 18a:**
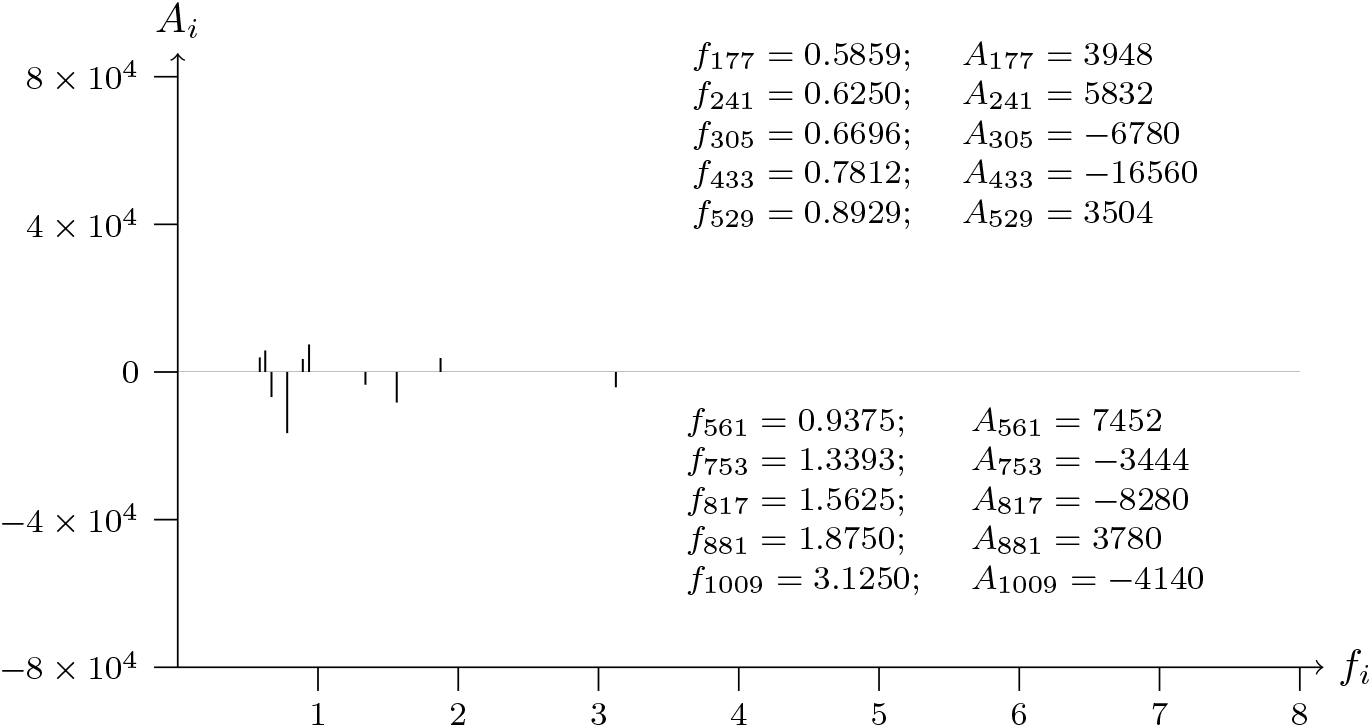
T2T-CHM13v2.0 chromosome 17; p telomere; analyzed data from 1 to 1200

**Figure 18b:**
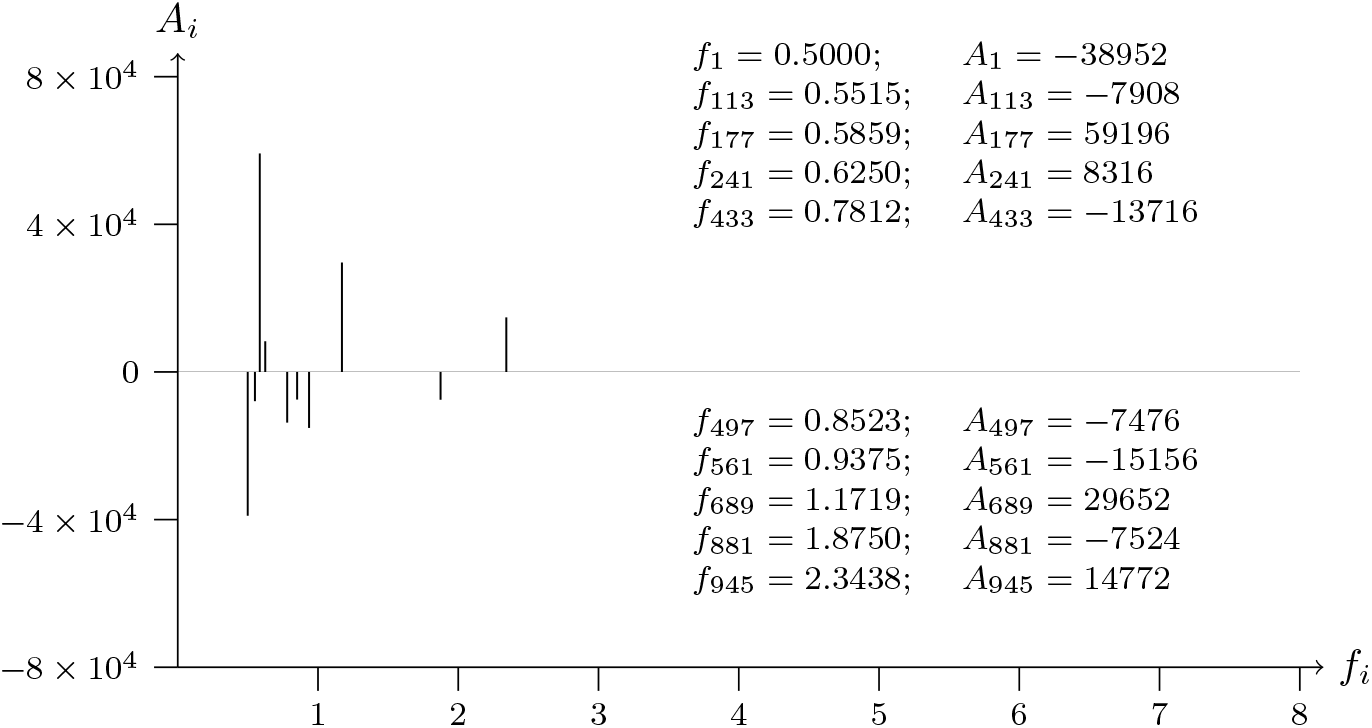
T2T-CHM13v2.0 chromosome 17; p telomere; analyzed data from 1200 to 1

**Figure 19a:**
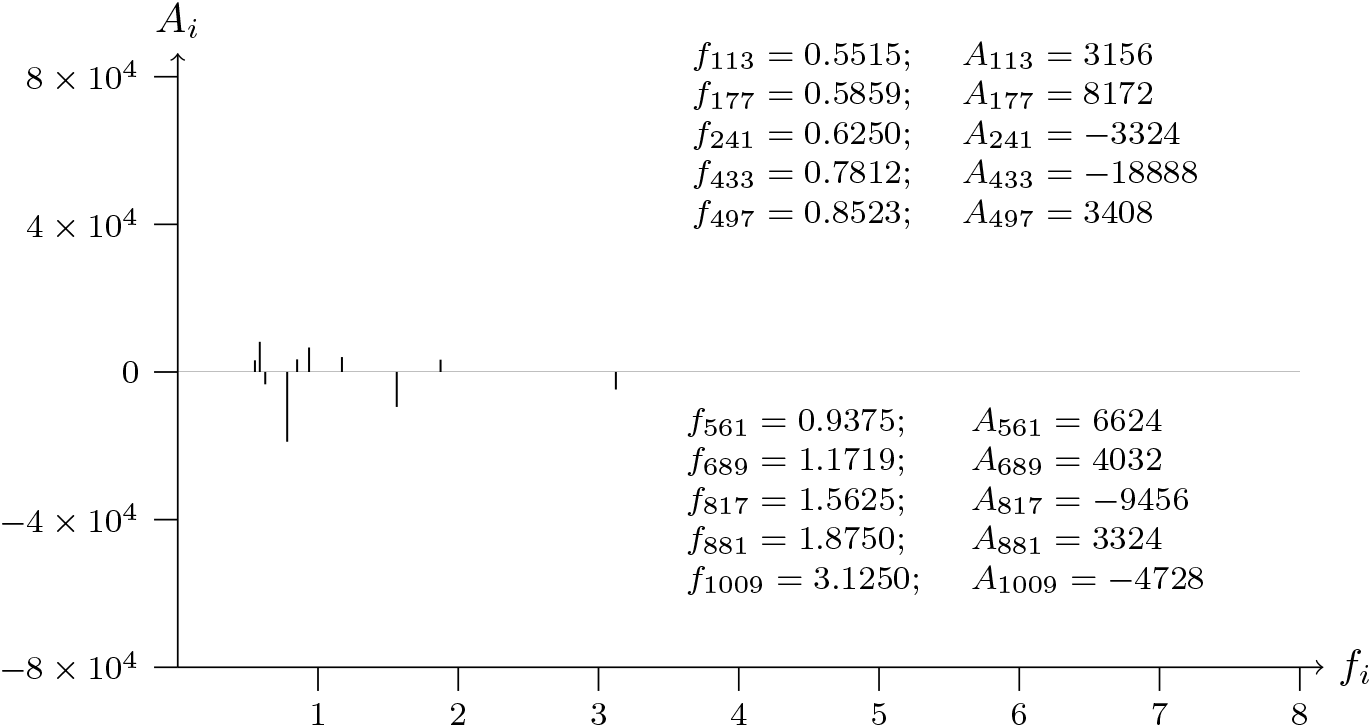
T2T-CHM13v2.0 chromosome 18; p telomere; analyzed data from 1 to 1200

**Figure 19b:**
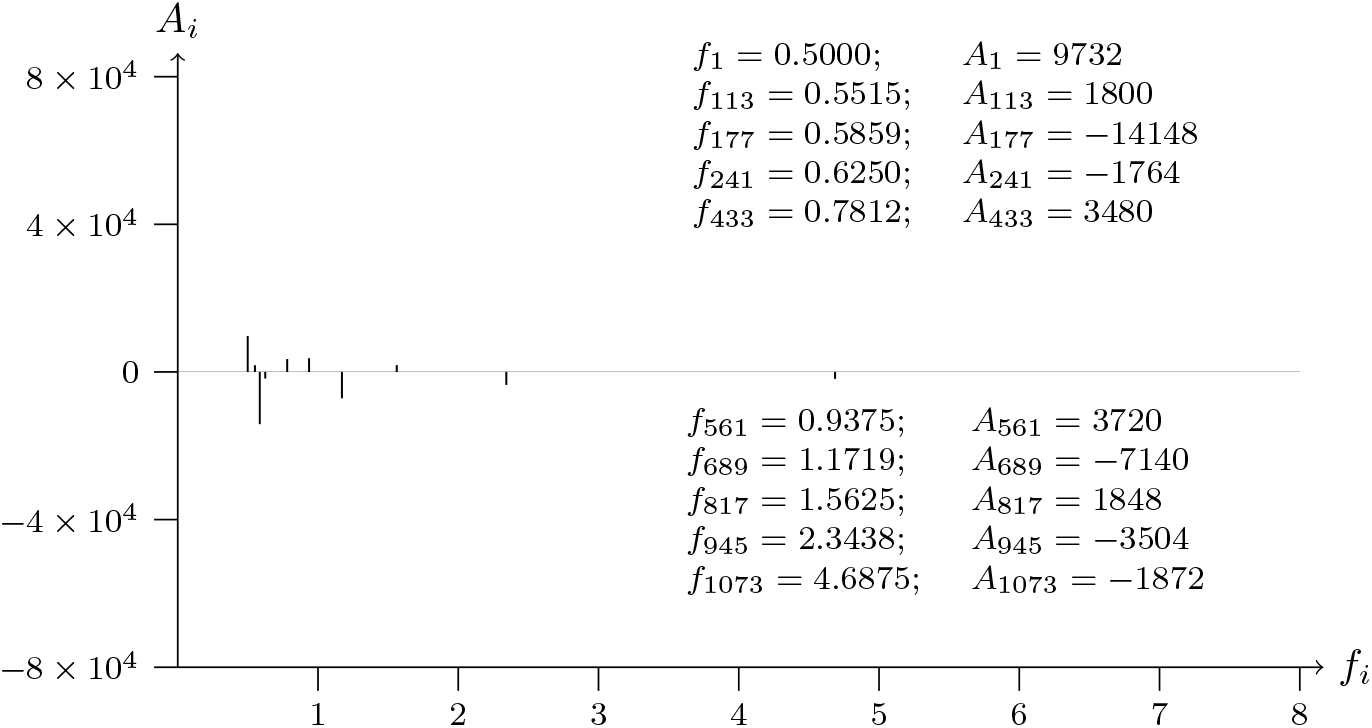
T2T-CHM13v2.0 chromosome 18; p telomere; analyzed data from 1200 to 1

**Figure 20a:**
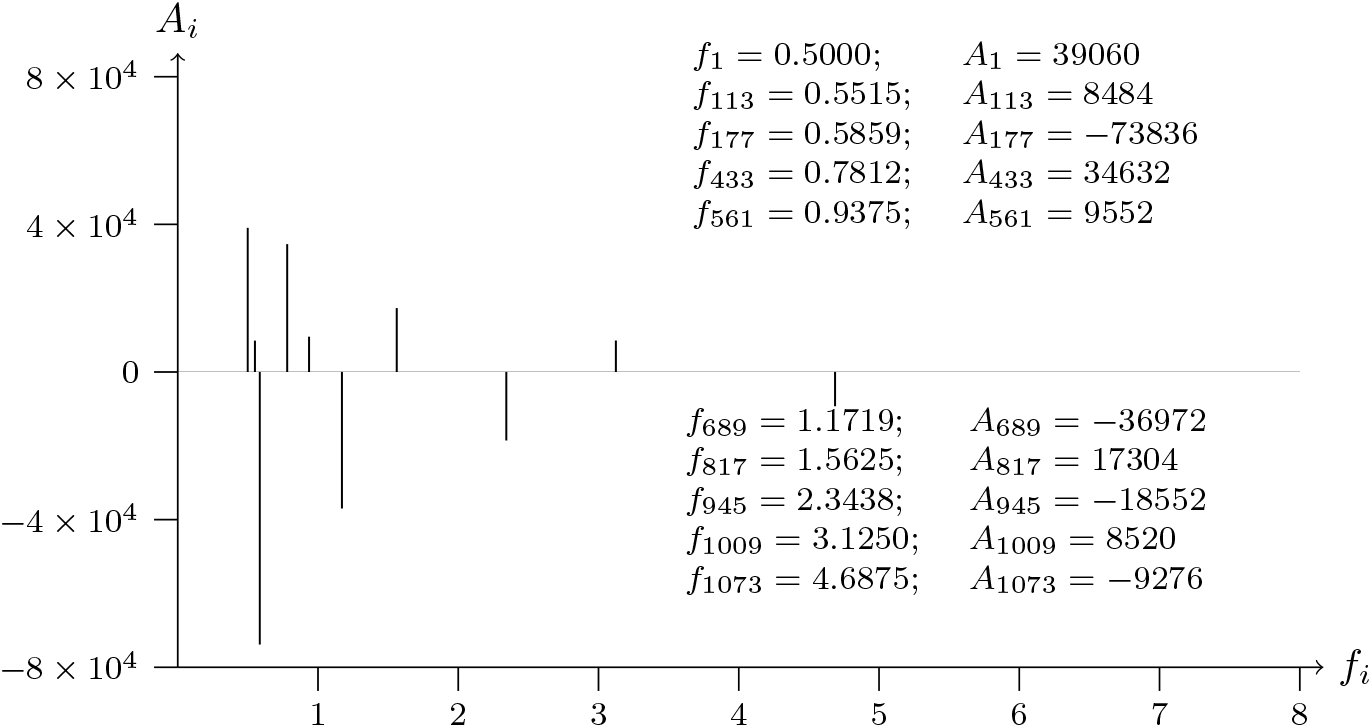
T2T-CHM13v2.0 chromosome 19; p telomere; analyzed data from 1 to 1200

**Figure 20b:**
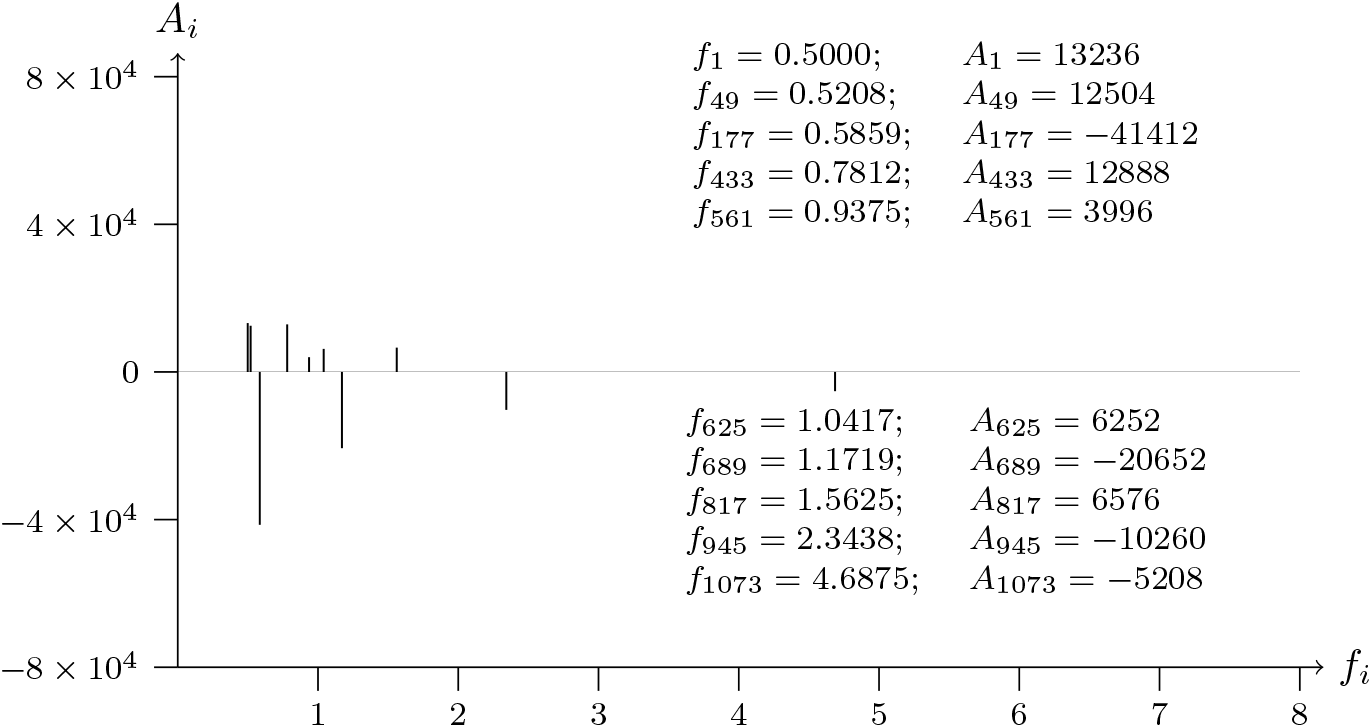
T2T-CHM13v2.0 chromosome 19; p telomere; analyzed data from 1200 to 1

**Figure 21a:**
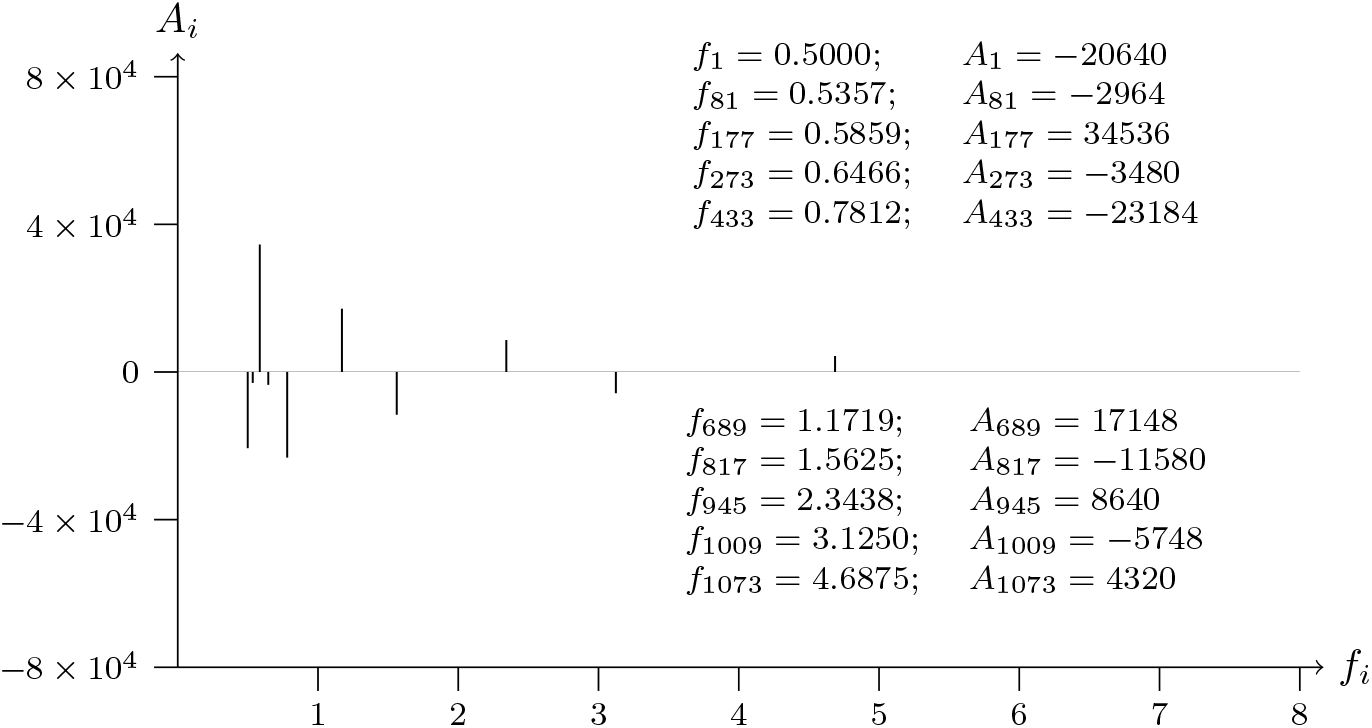
T2T-CHM13v2.0 chromosome 20; p telomere; analyzed data from 1 to 1200

**Figure 21b:**
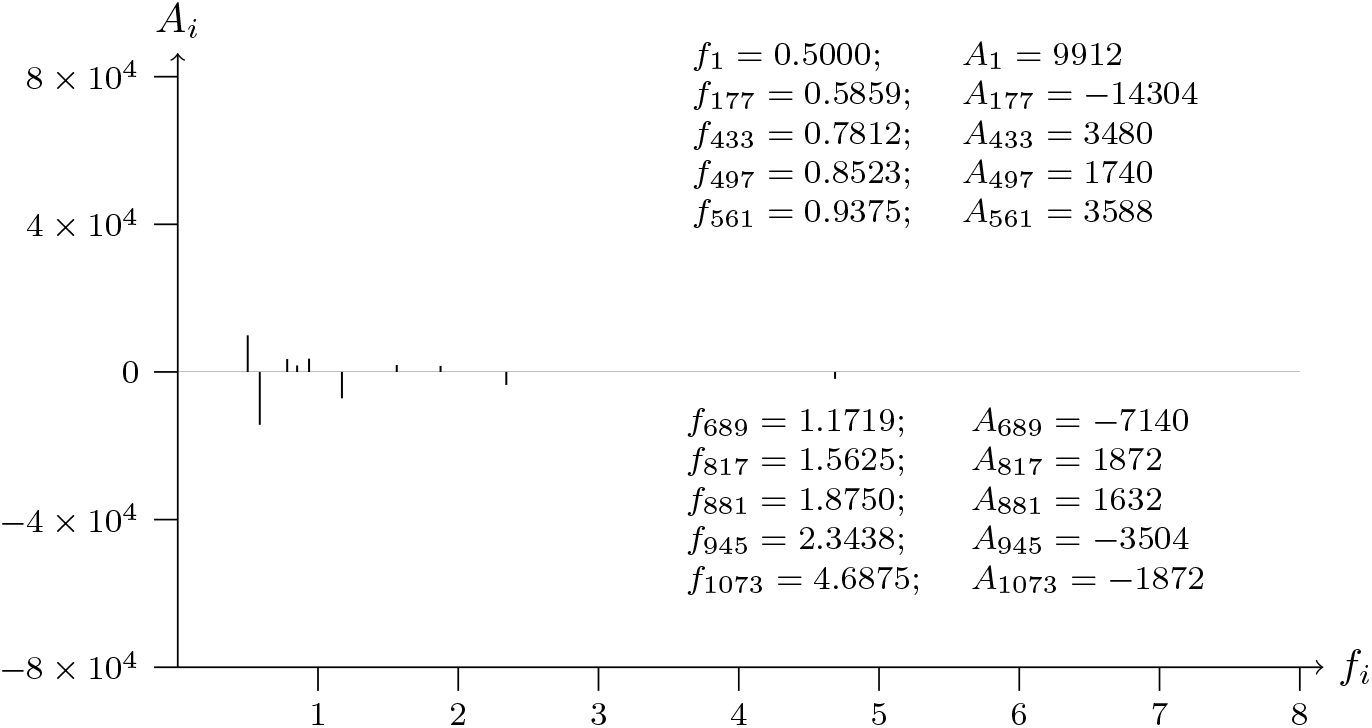
T2T-CHM13v2.0 chromosome 20; p telomere; analyzed data from 1200 to 1

**Figure 22a:**
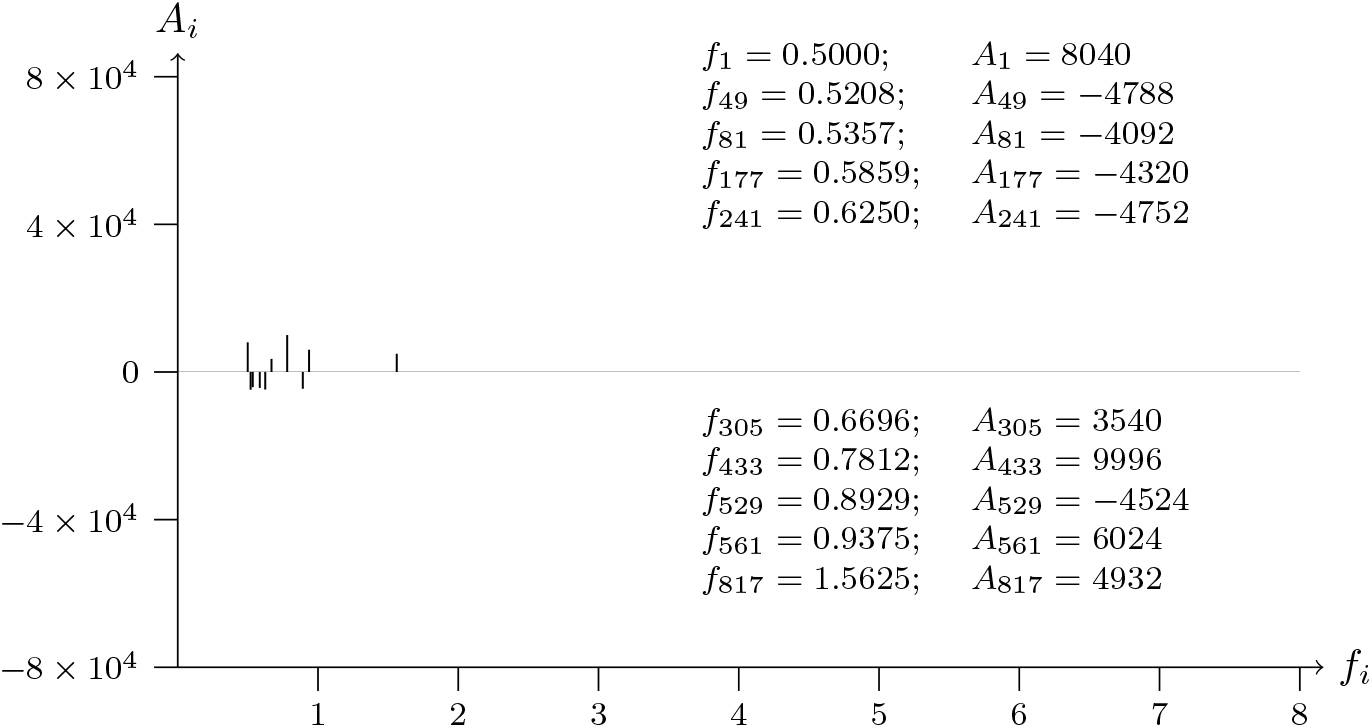
T2T-CHM13v2.0 chromosome 21; p telomere; analyzed data from 1 to 1200

**Figure 22b:**
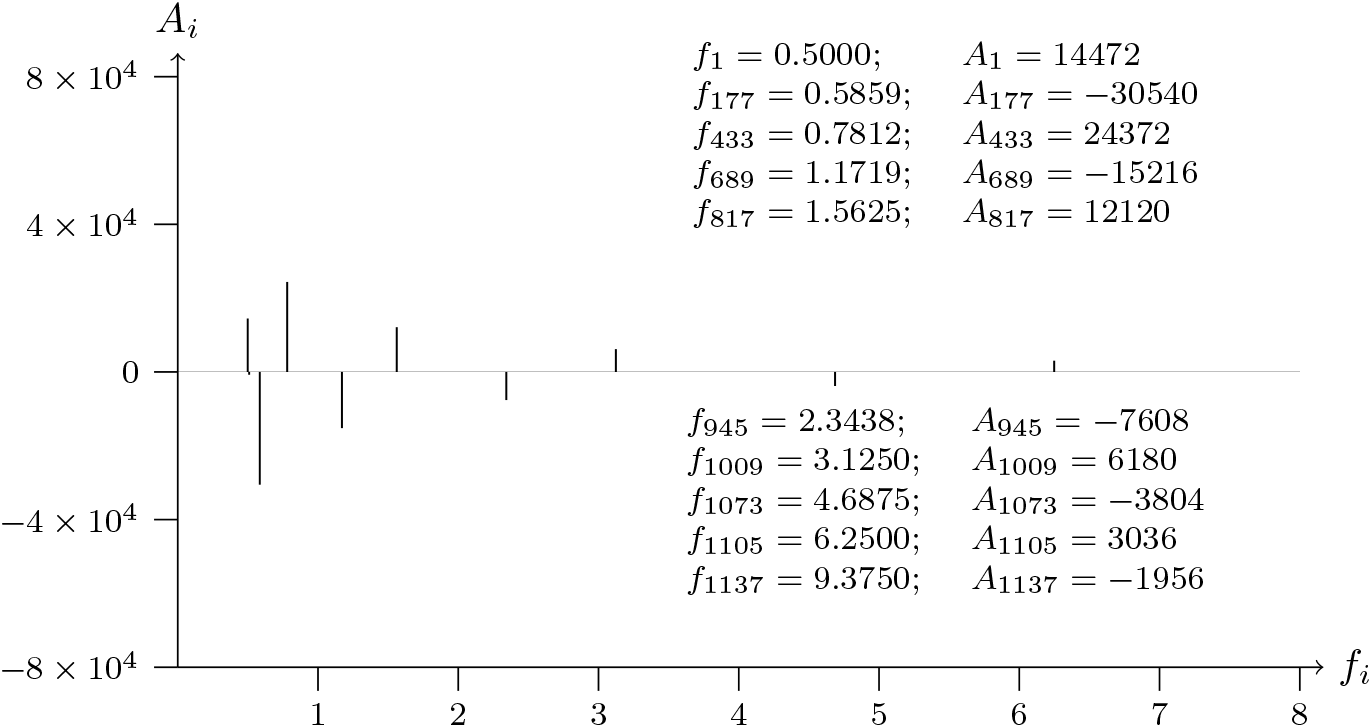
T2T-CHM13v2.0 chromosome 21; p telomere; analyzed data from 1200 to 1

**Figure 23a:**
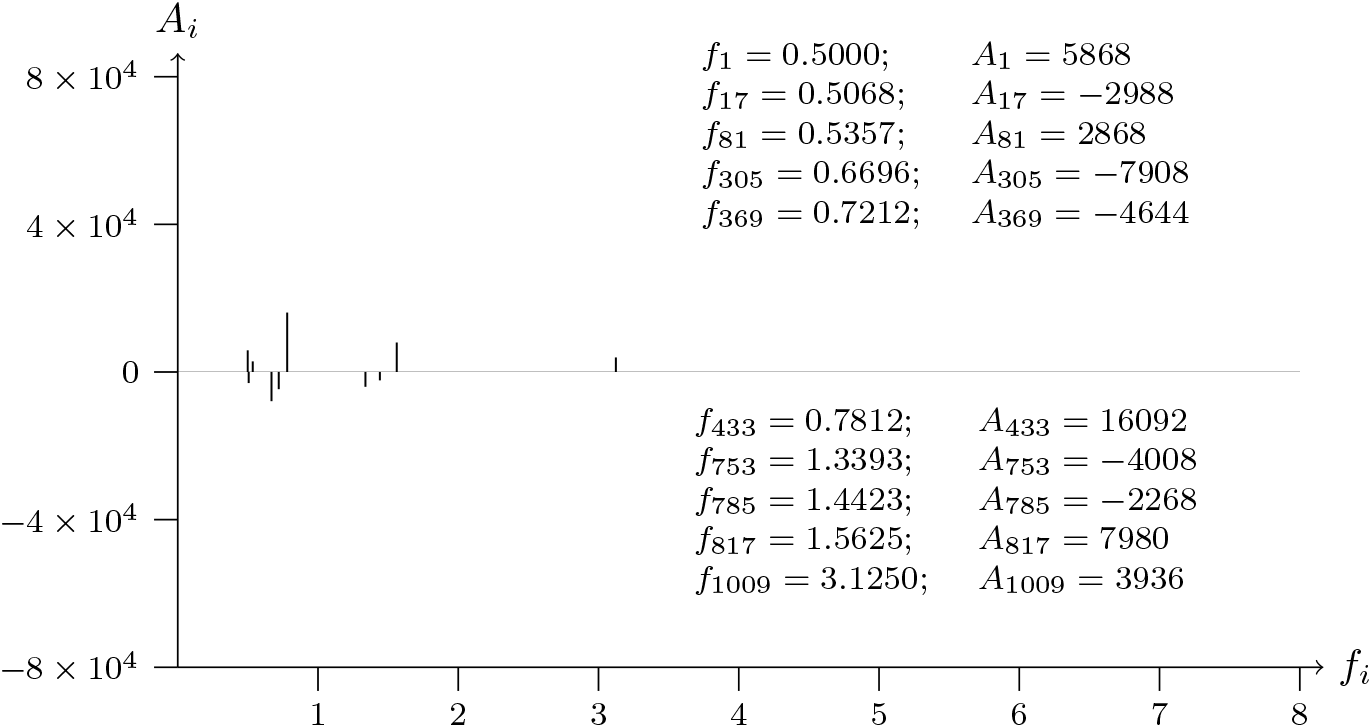
T2T-CHM13v2.0 chromosome 22; p telomere; analyzed data from 1 to 1200

**Figure 23b:**
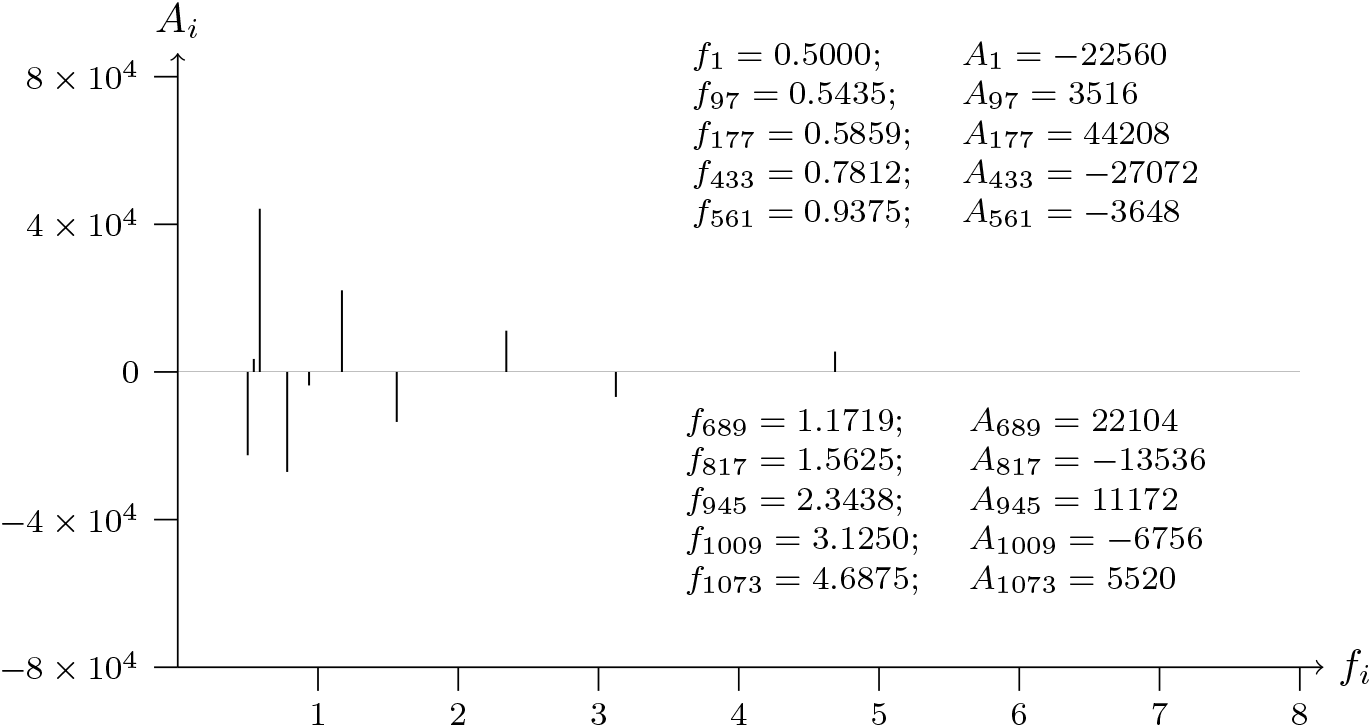
T2T-CHM13v2.0 chromosome 22; p telomere; analyzed data from 1200 to 1

**Figure 24a:**
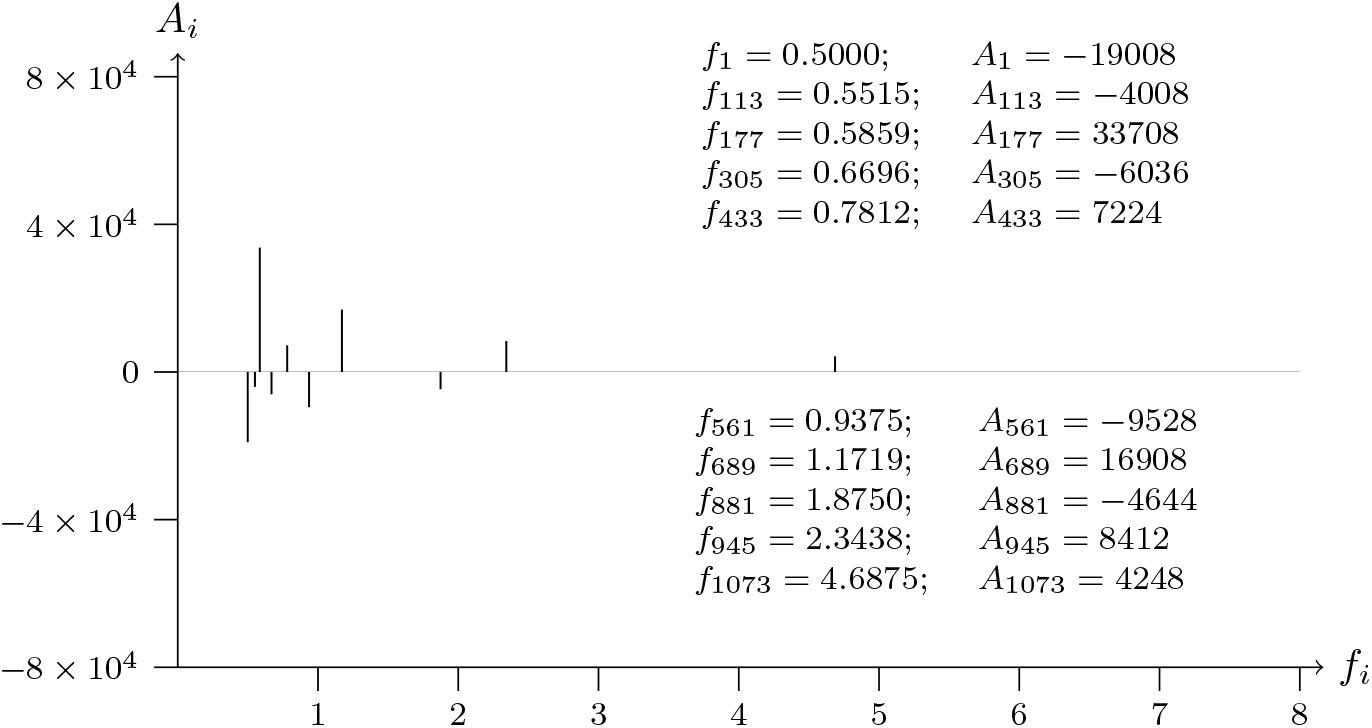
T2T-CHM13v2.0 chromosome X; p telomere; analyzed data from 1 to 1200

**Figure 24b:**
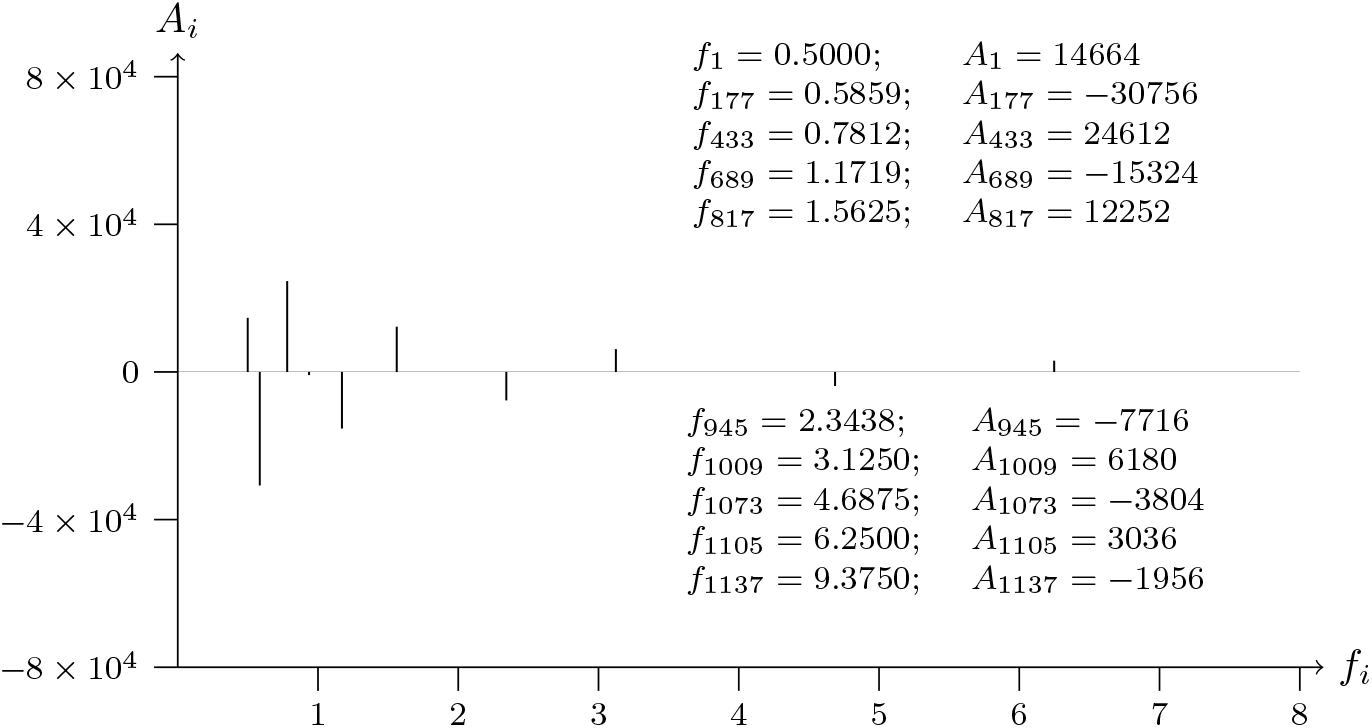
T2T-CHM13v2.0 chromosome X; p telomere; analyzed data from 1200 to 1

**Figure 25a:**
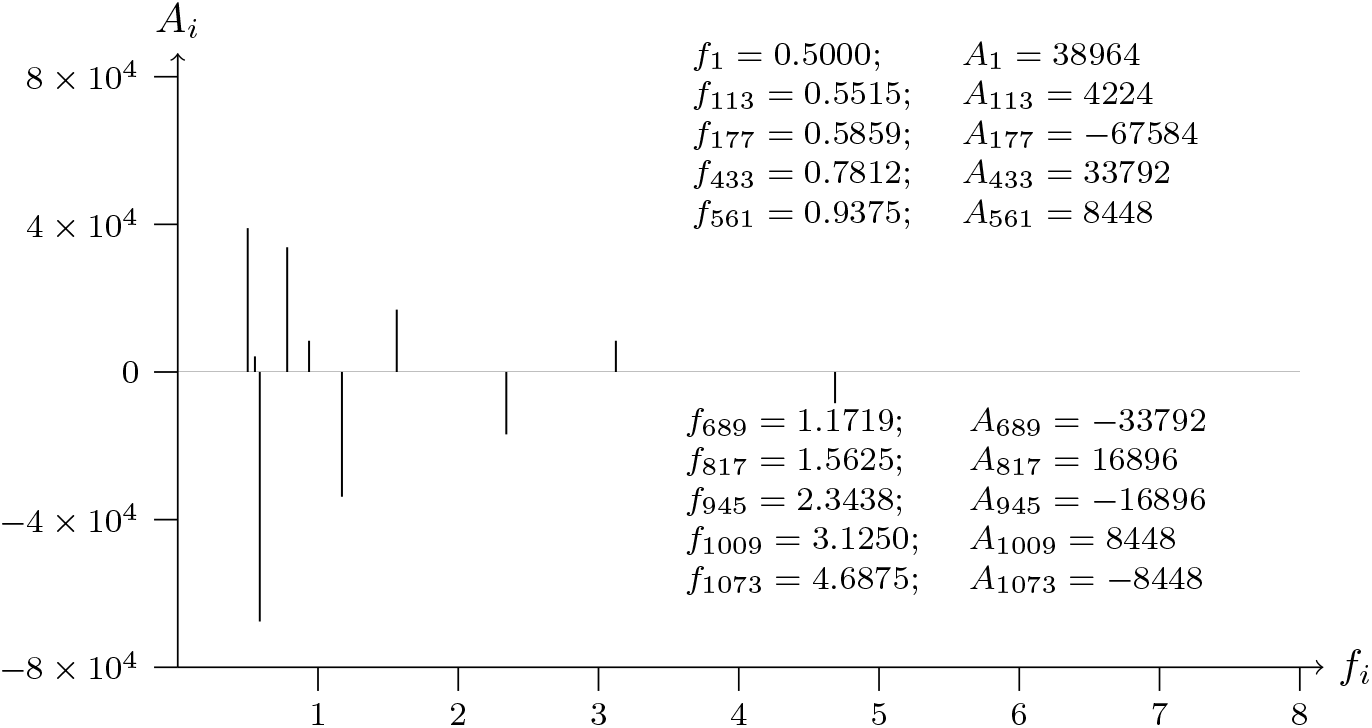
T2T-CHM13v2.0 chromosome Y; p telomere; analyzed data from 1 to 1200

**Figure 25b:**
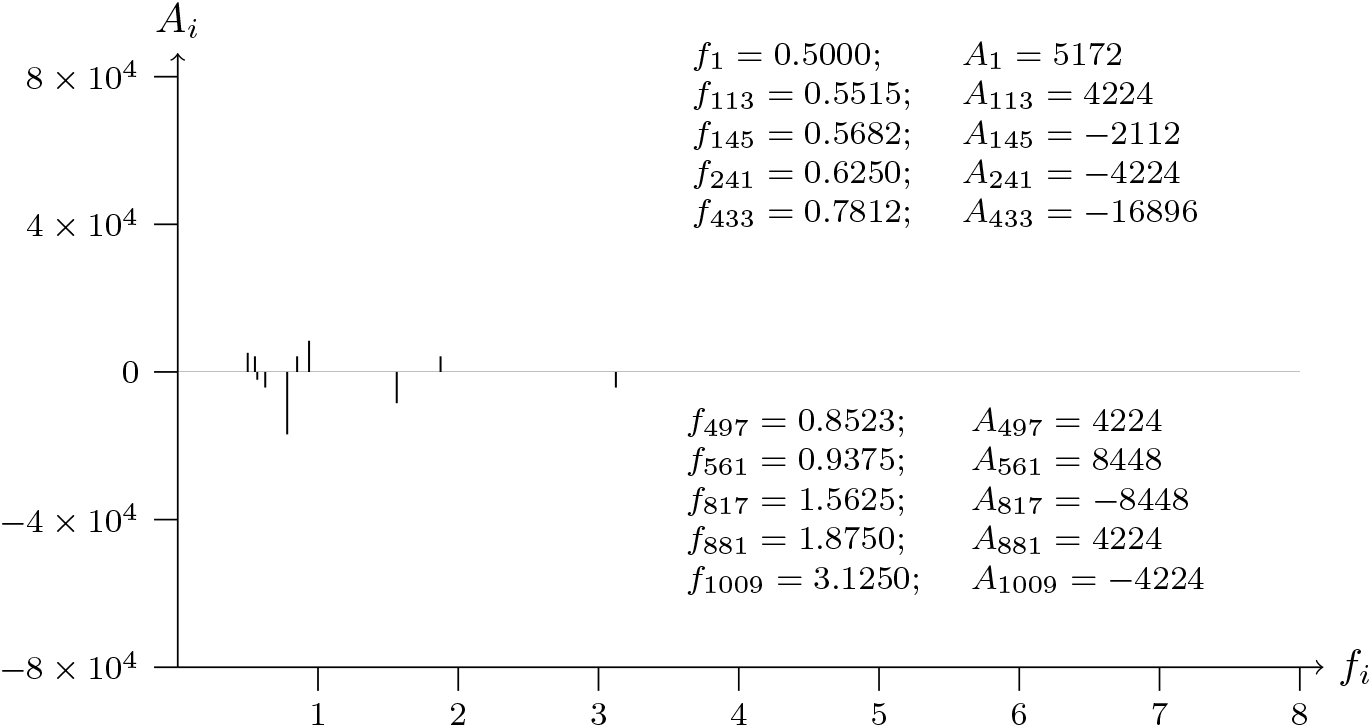
T2T-CHM13v2.0 chromosome Y; p telomere; analyzed data from 1200 to 1

### 3.1 SWT analysis of chromosome 1

**Figure 26a:**
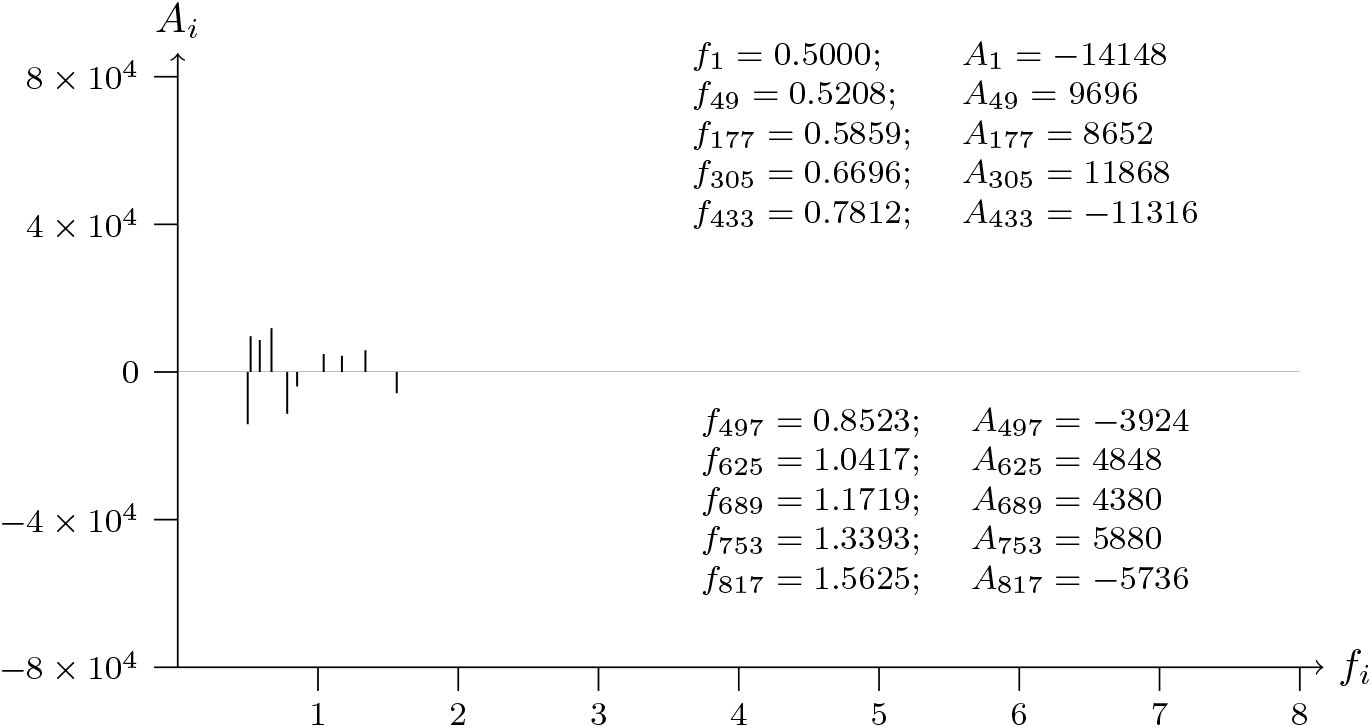
T2T-CHM13v2.0; chromosome 1; p centromere; analyzed data from 0 to 1199

**Figure 26b:**
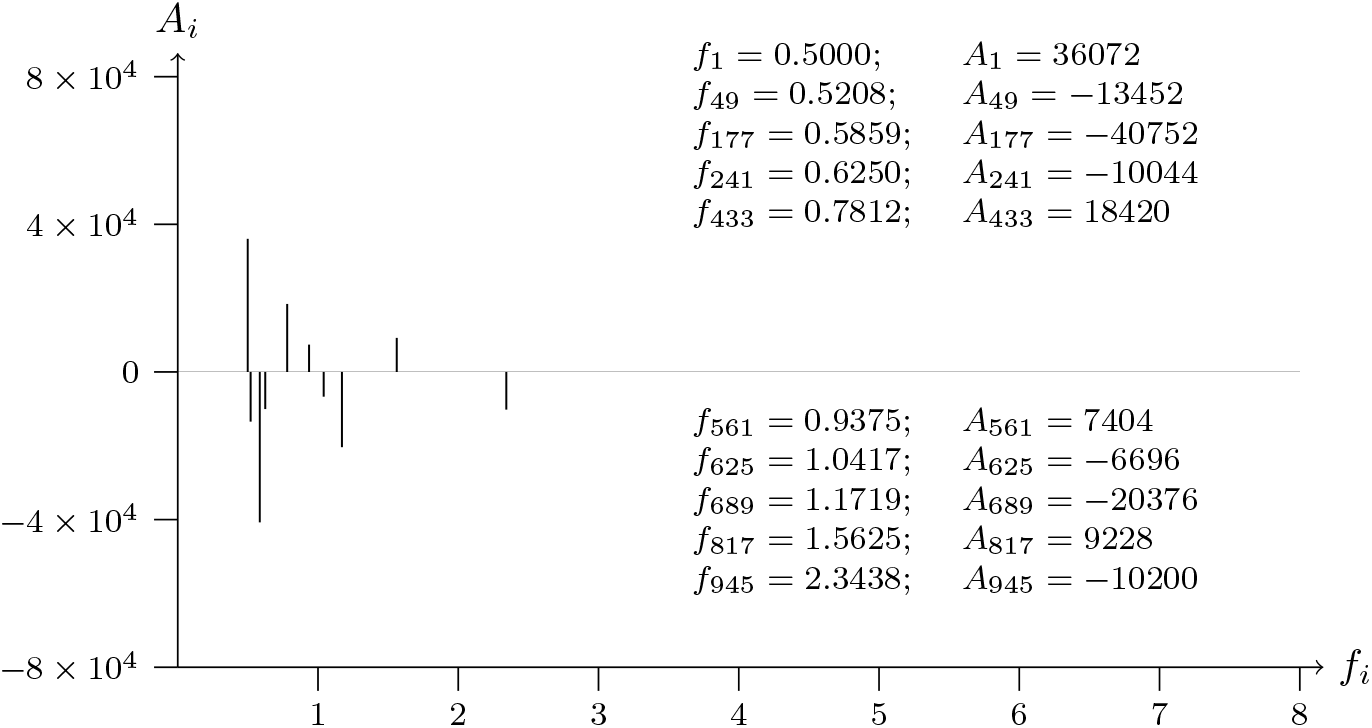
T2T-CHM13v2.0; chromosome 1; p centromere; analyzed data from 0 to 1199

**Figure 27a:**
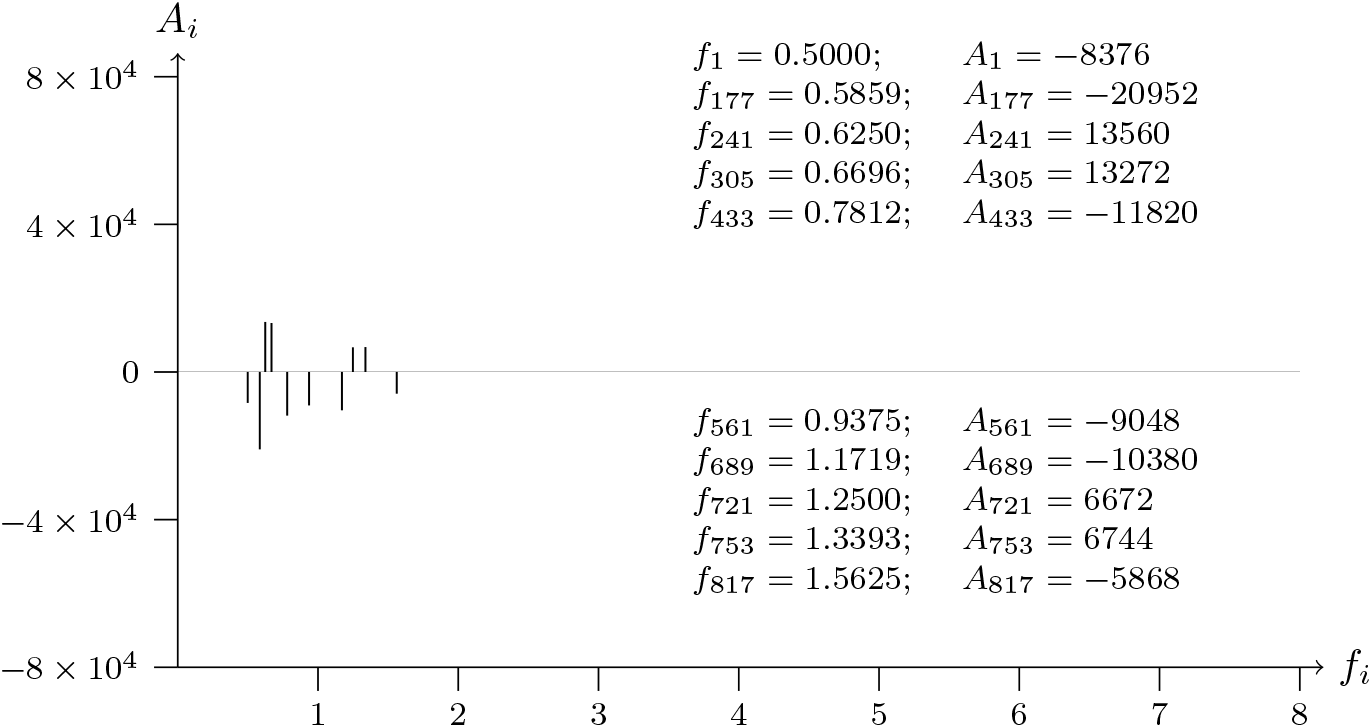
T2T-CHM13v2.0; chromosome 1; p centromere; analyzed data from 1126109 to 1127308

**Figure 27b:**
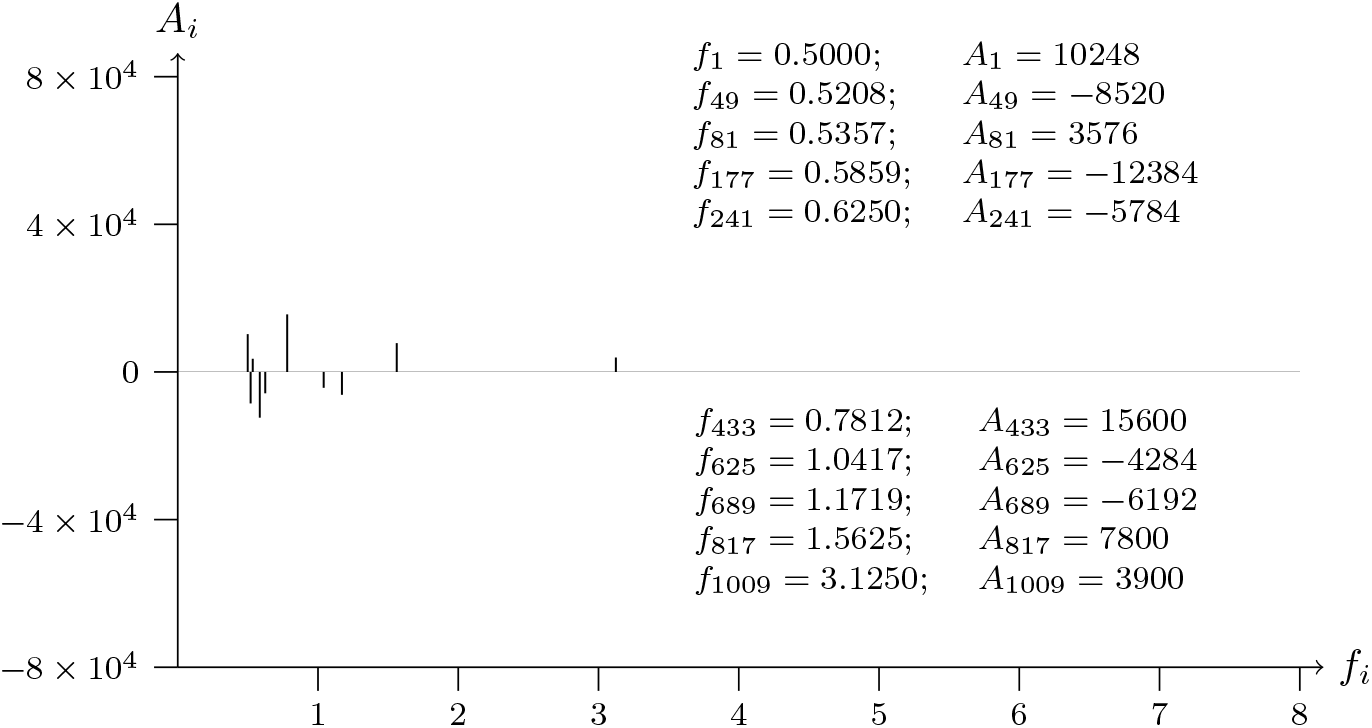
T2T-CHM13v2.0; chromosome 1; p centromere; analyzed data from 1126109 to 1127308

**Figure 28a:**
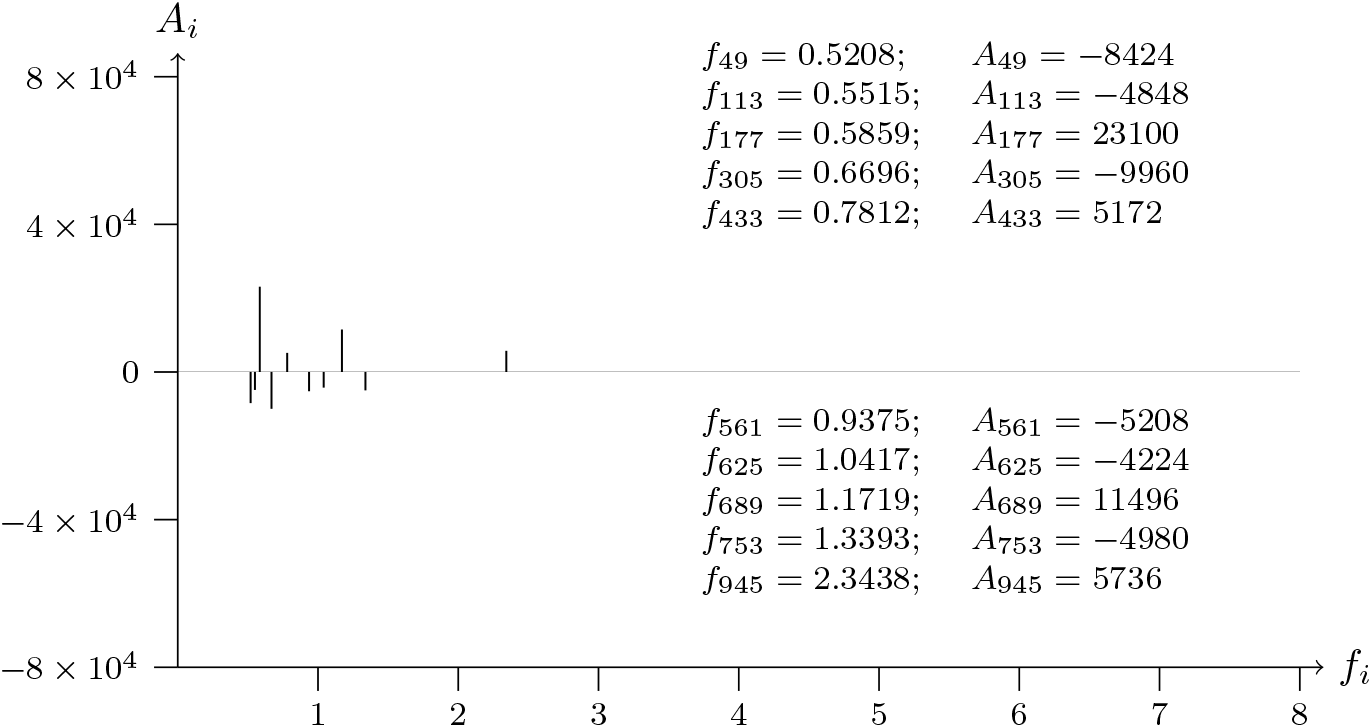
T2T-CHM13v2.0; chromosome 1; p centromere; analyzed data from 2251019 to 2252219

**Figure 28b:**
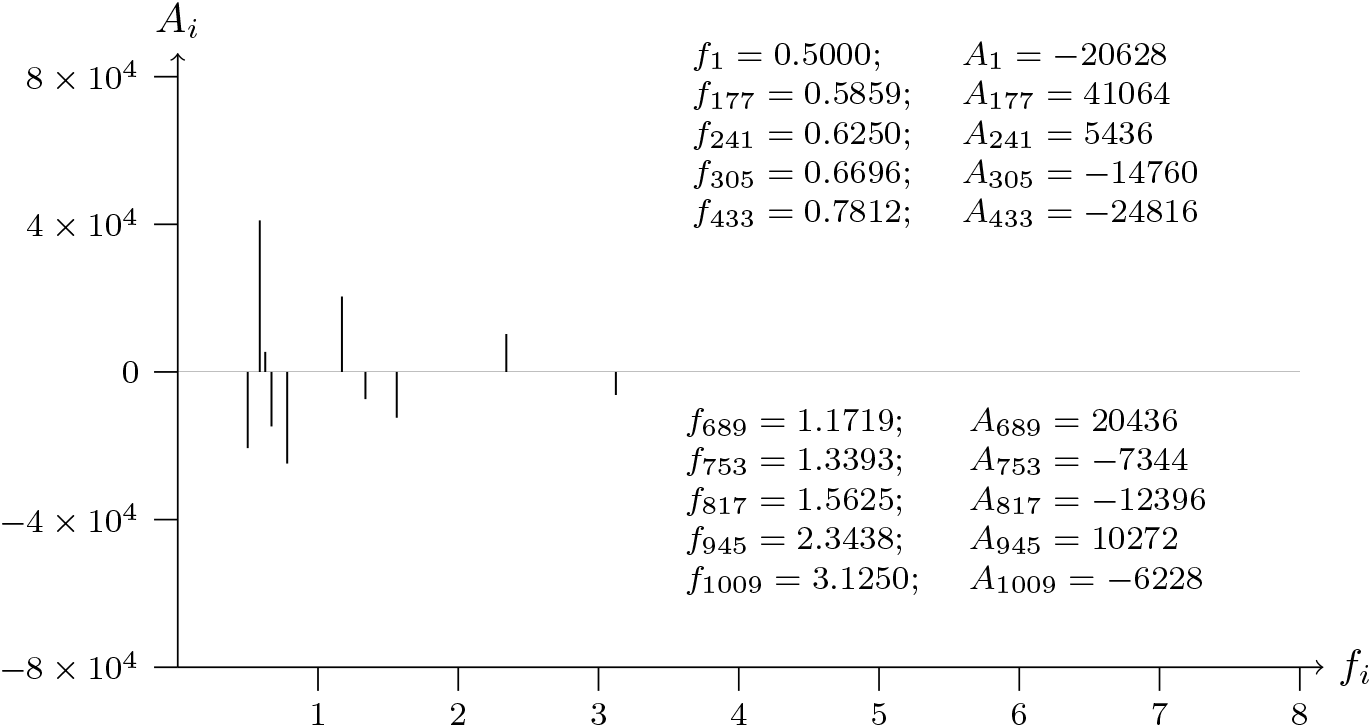
T2T-CHM13v2.0; chromosome 1; p centromere; analyzed data from 2251019 to 2252219

**Figure 29a:**
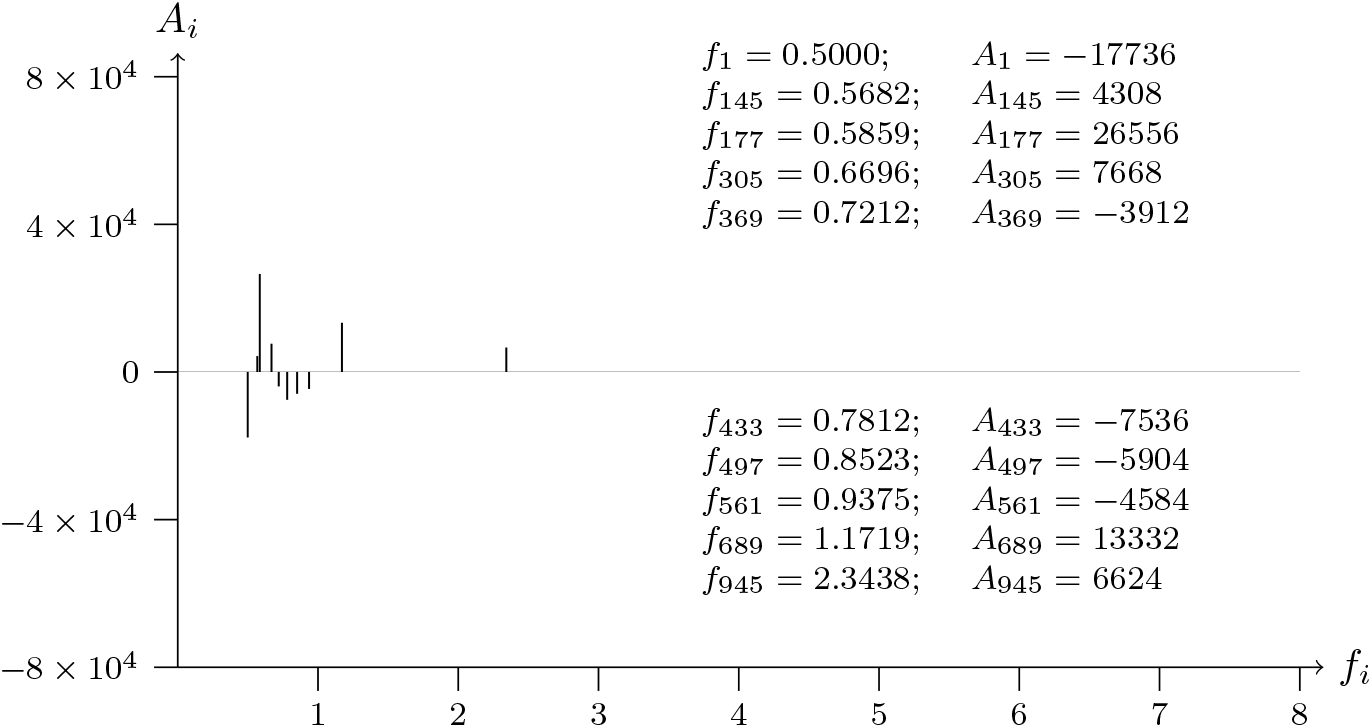
T2T-CHM13v2.0; chromosome 1; q centromere; analyzed data from 0 to 1199

**Figure 29b:**
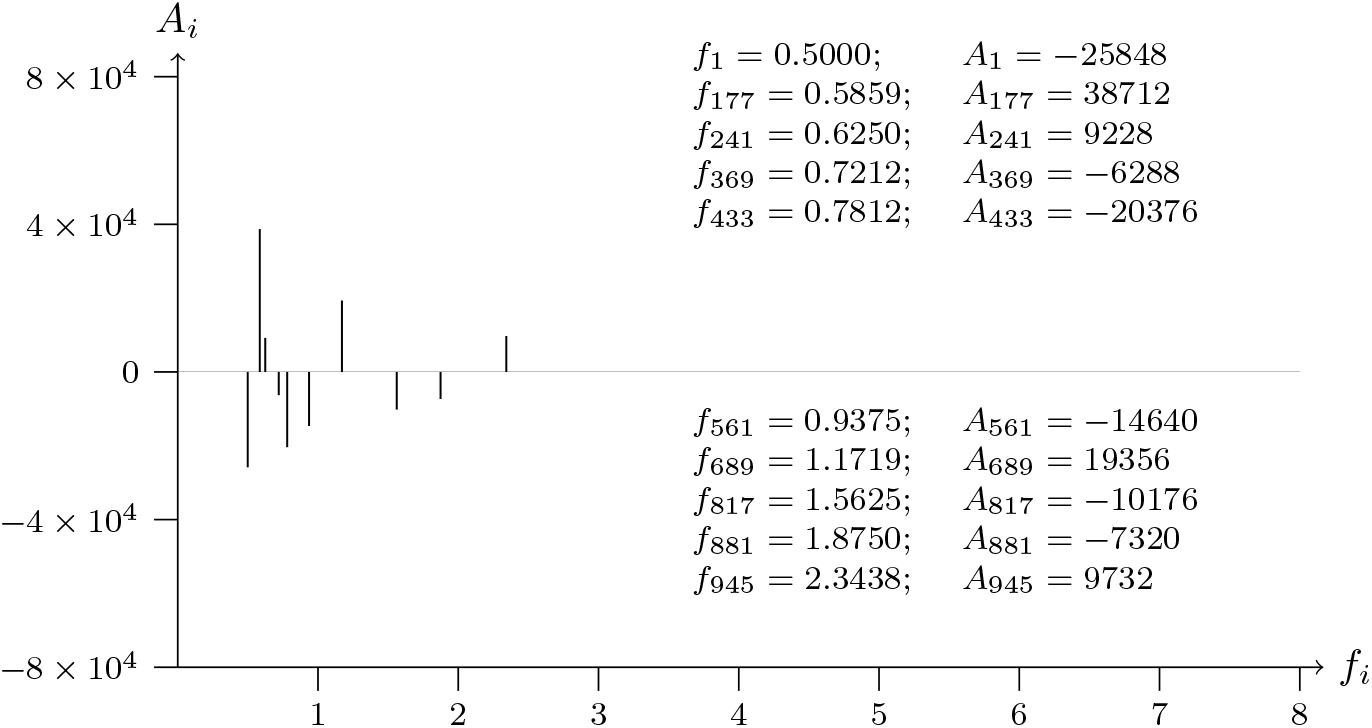
T2T-CHM13v2.0; chromosome 1; q centromere; analyzed data from 0 to 1199

**Figure 30a:**
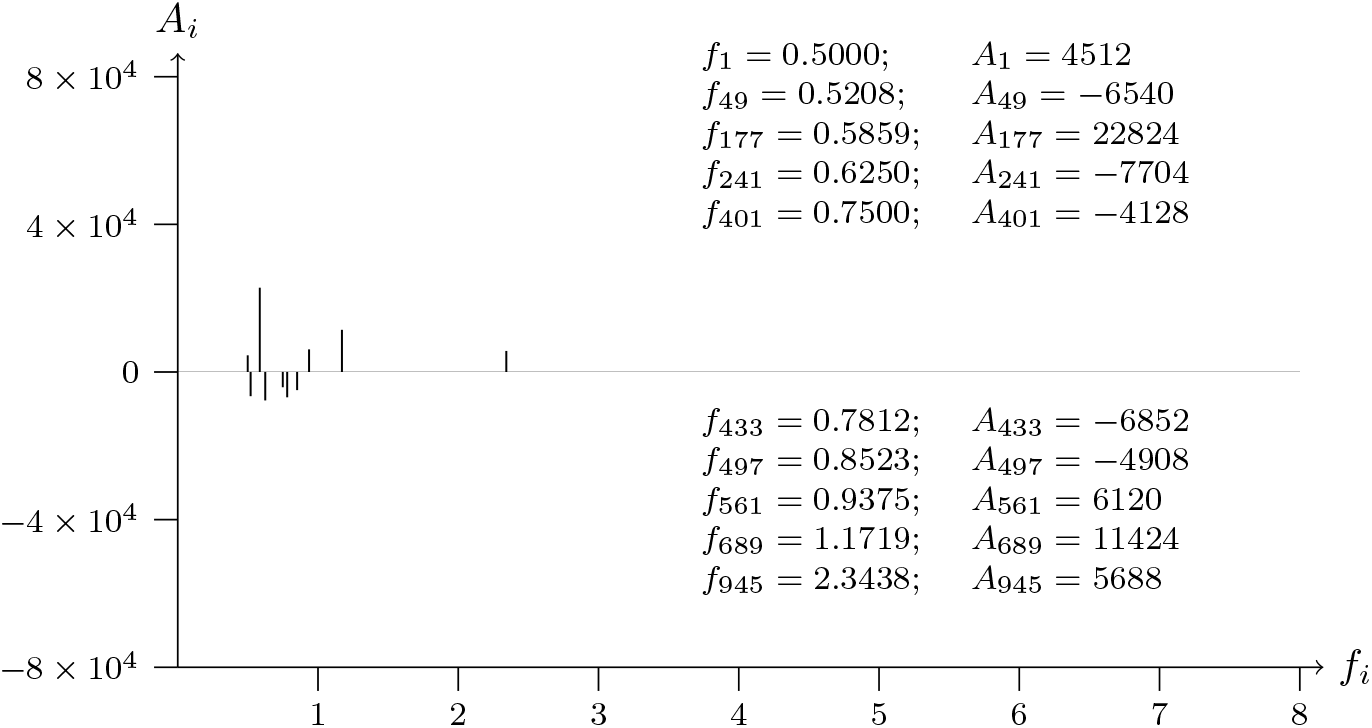
T2T-CHM13v2.0; chromosome 1; q centromere; analyzed data from 1126110 to 1127309

**Figure 30b:**
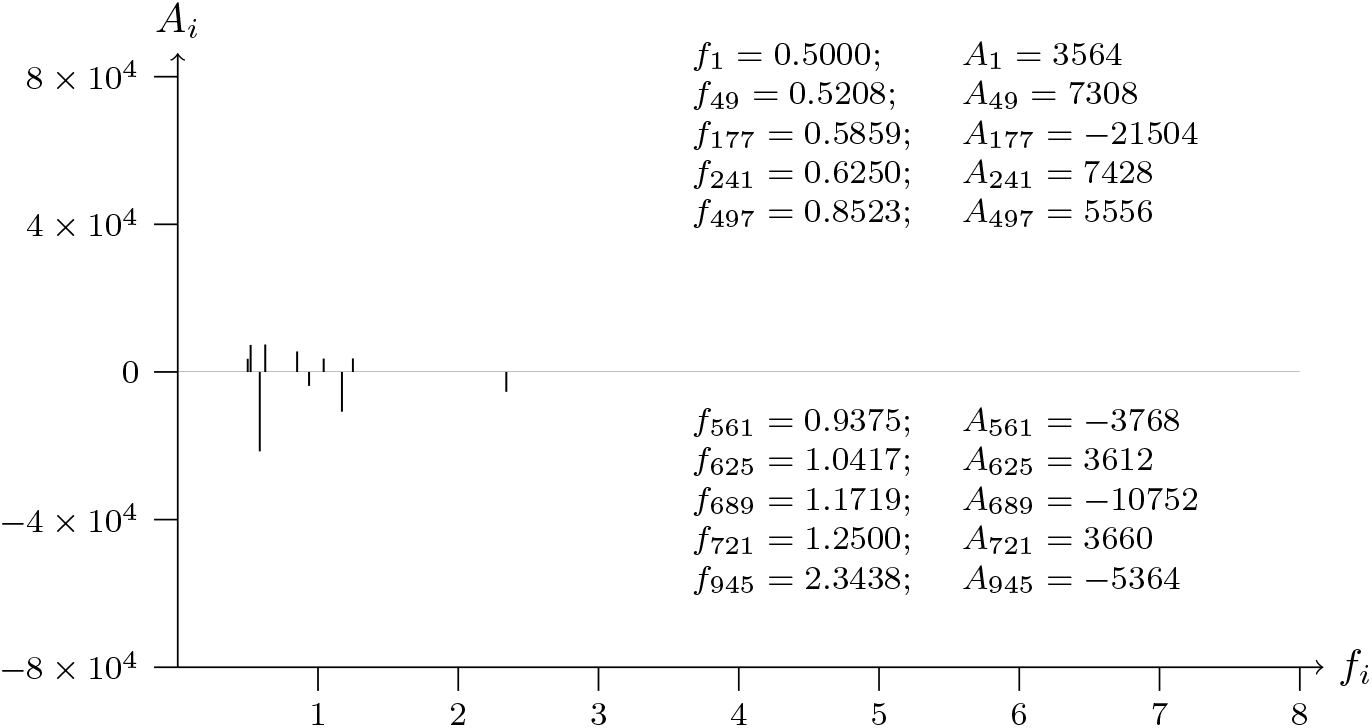
T2T-CHM13v2.0; chromosome 1; q centromere; analyzed data from 1126110 to 1127309

**Figure 31a:**
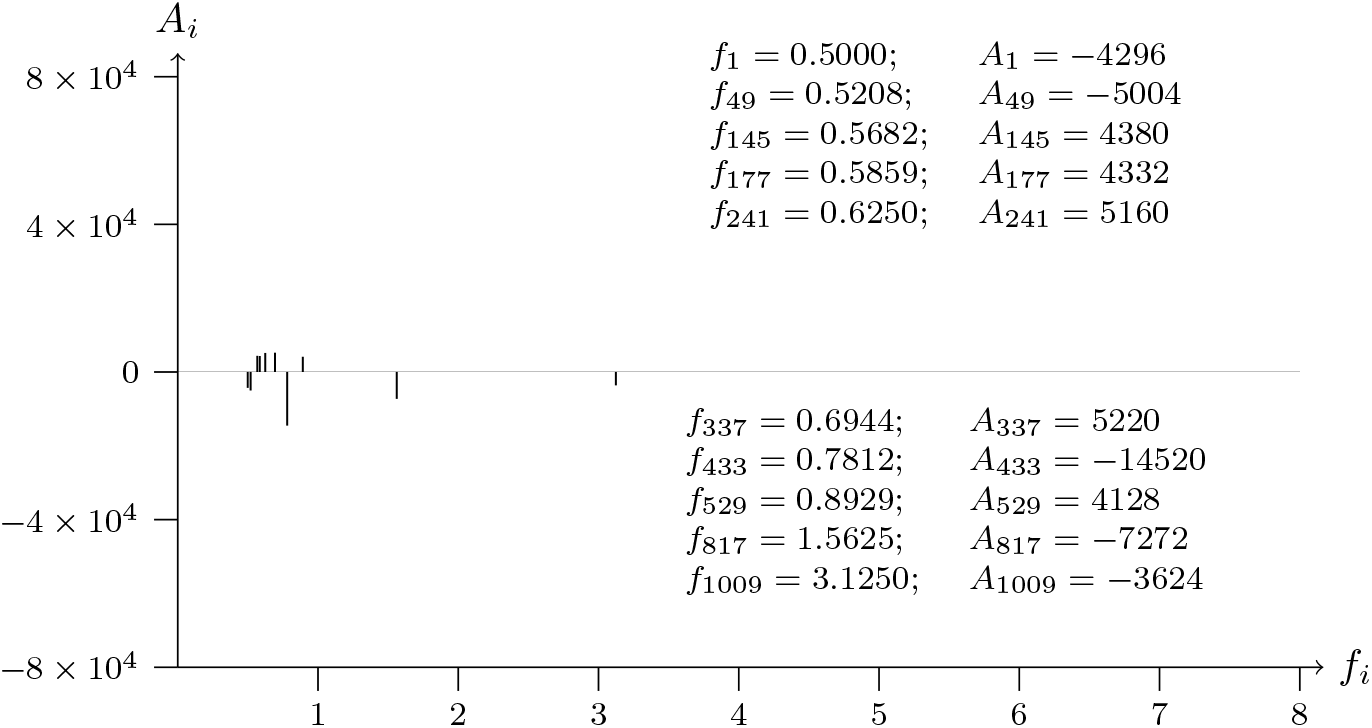
T2T-CHM13v2.0; chromosome 1; q centromere; analyzed data from 2251020 to 2252220

**Figure 31b:**
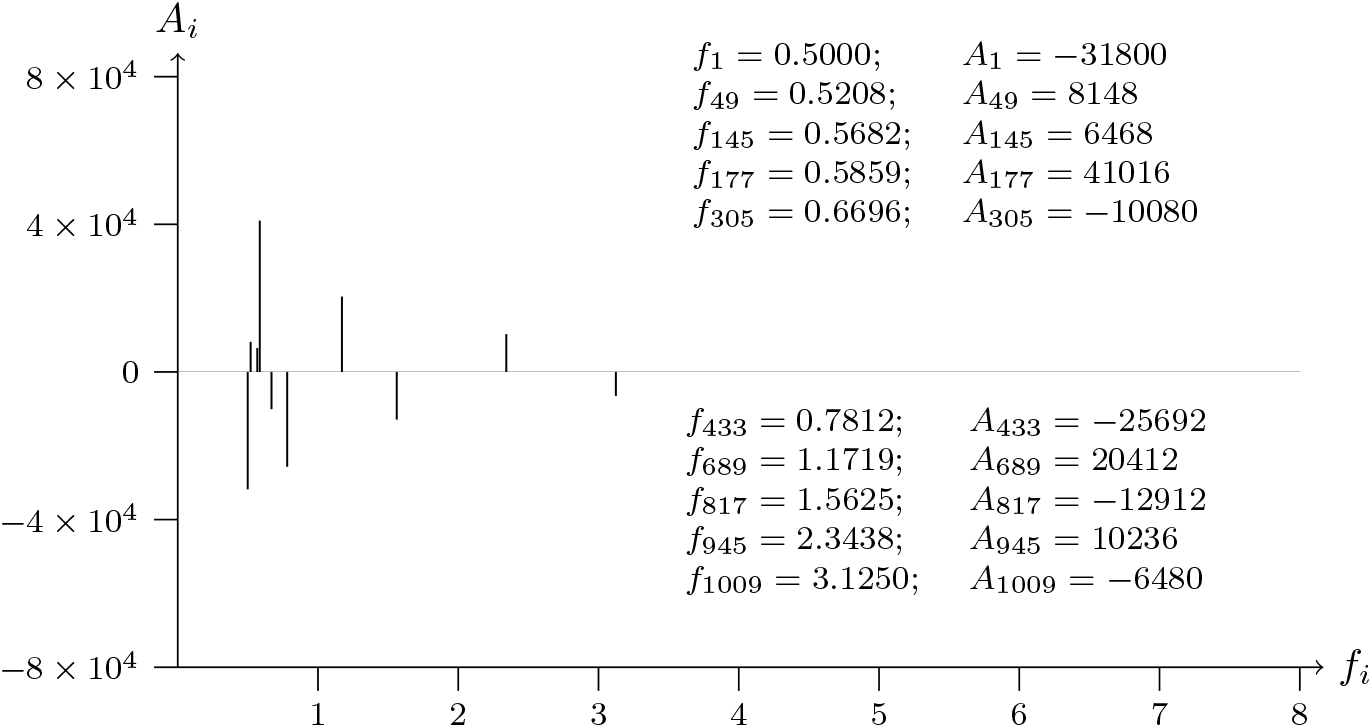
T2T-CHM13v2.0; chromosome 1; q centromere; analyzed data from 2251020 to 2252220

**Figure 32a:**
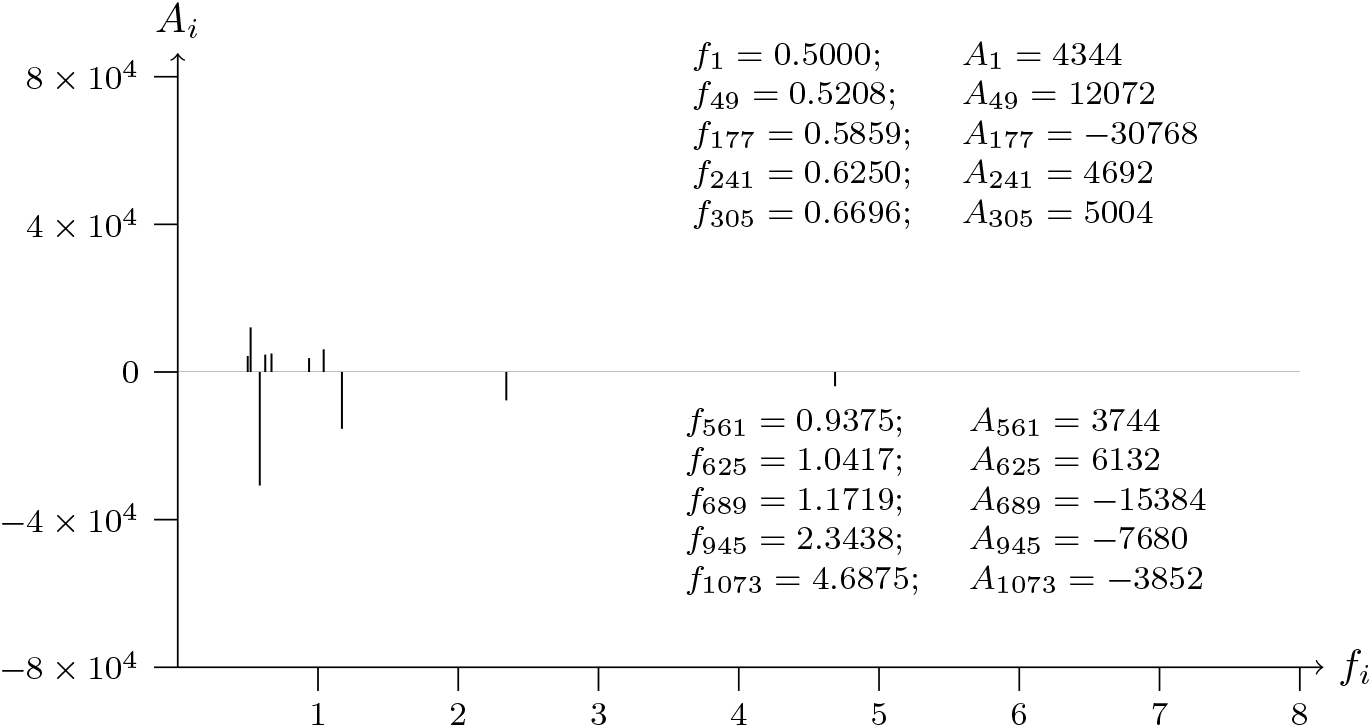
T2T-CHM13v2.0; chromosome 1; p telomere; analyzed data from 0 to 1199

**Figure 32b:**
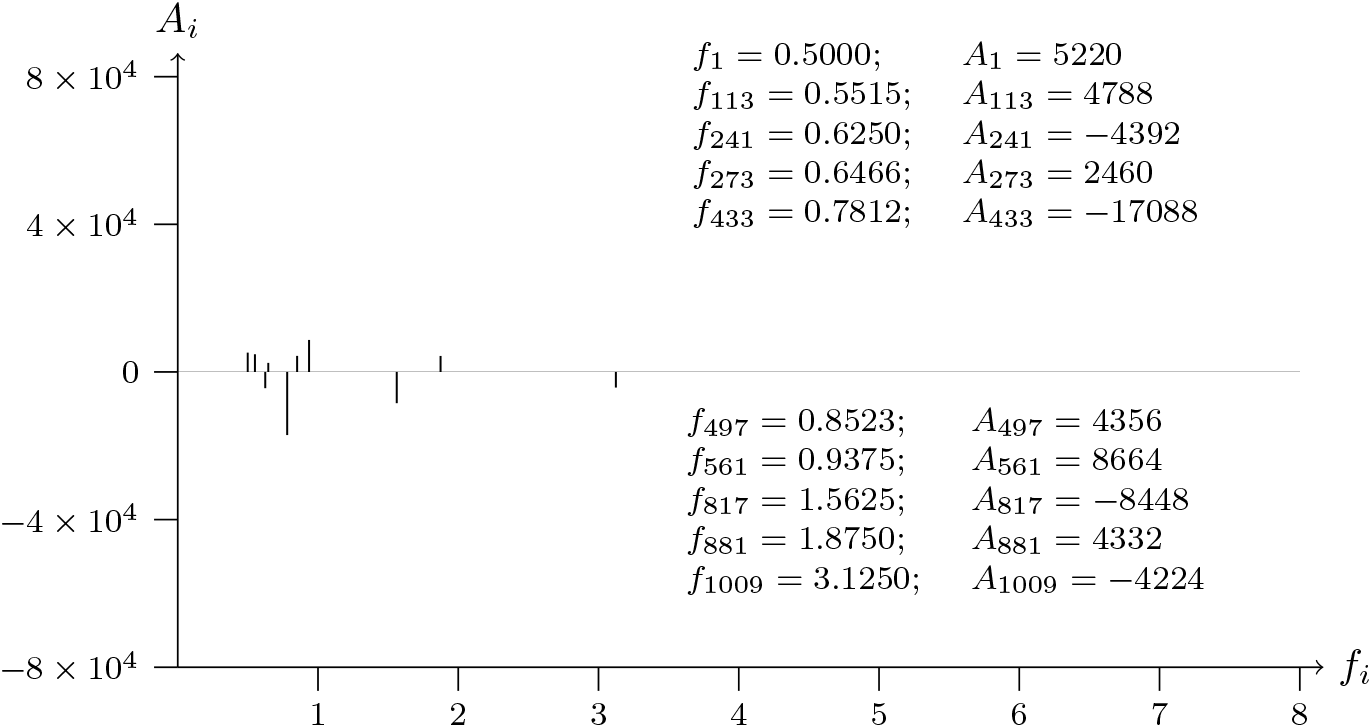
T2T-CHM13v2.0; chromosome 1; p telomere; analyzed data from 0 to 1199

**Figure 33a:**
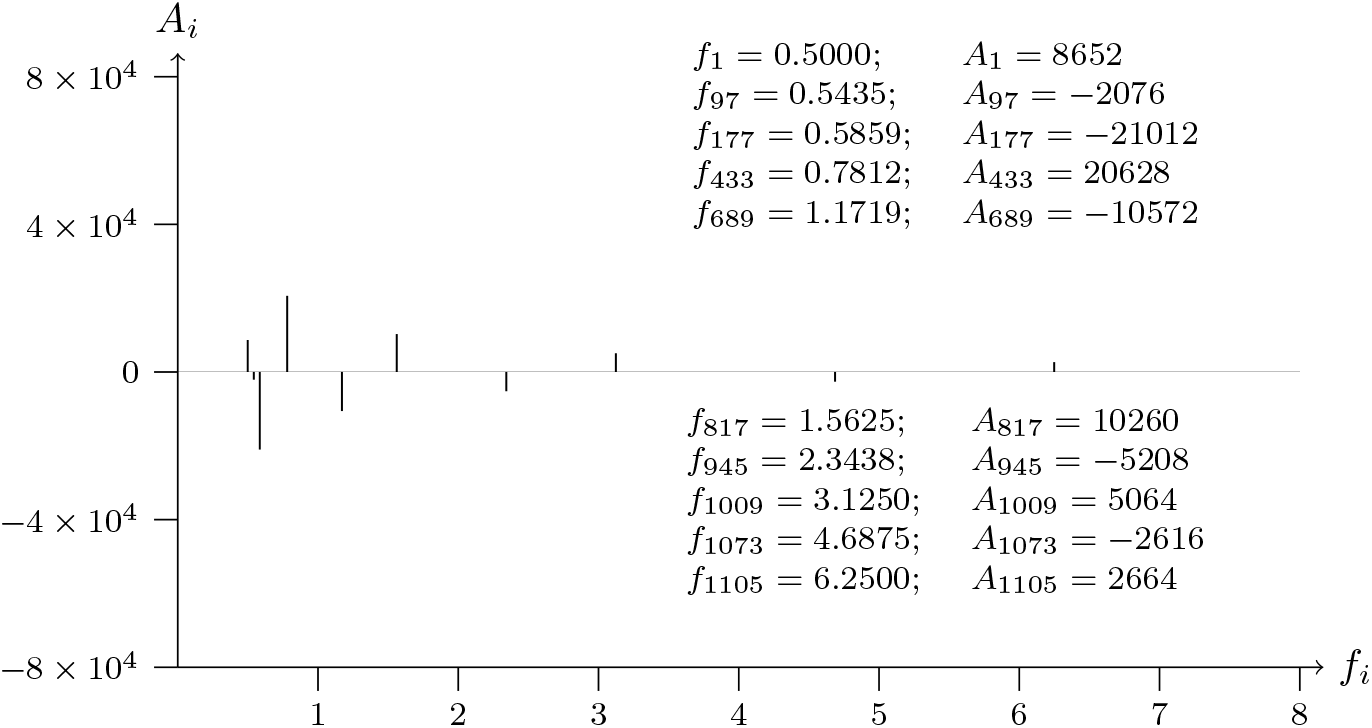
T2T-CHM13v2.0; chromosome 1; p telomere; analyzed data from 1500 to 2699

**Figure 33b:**
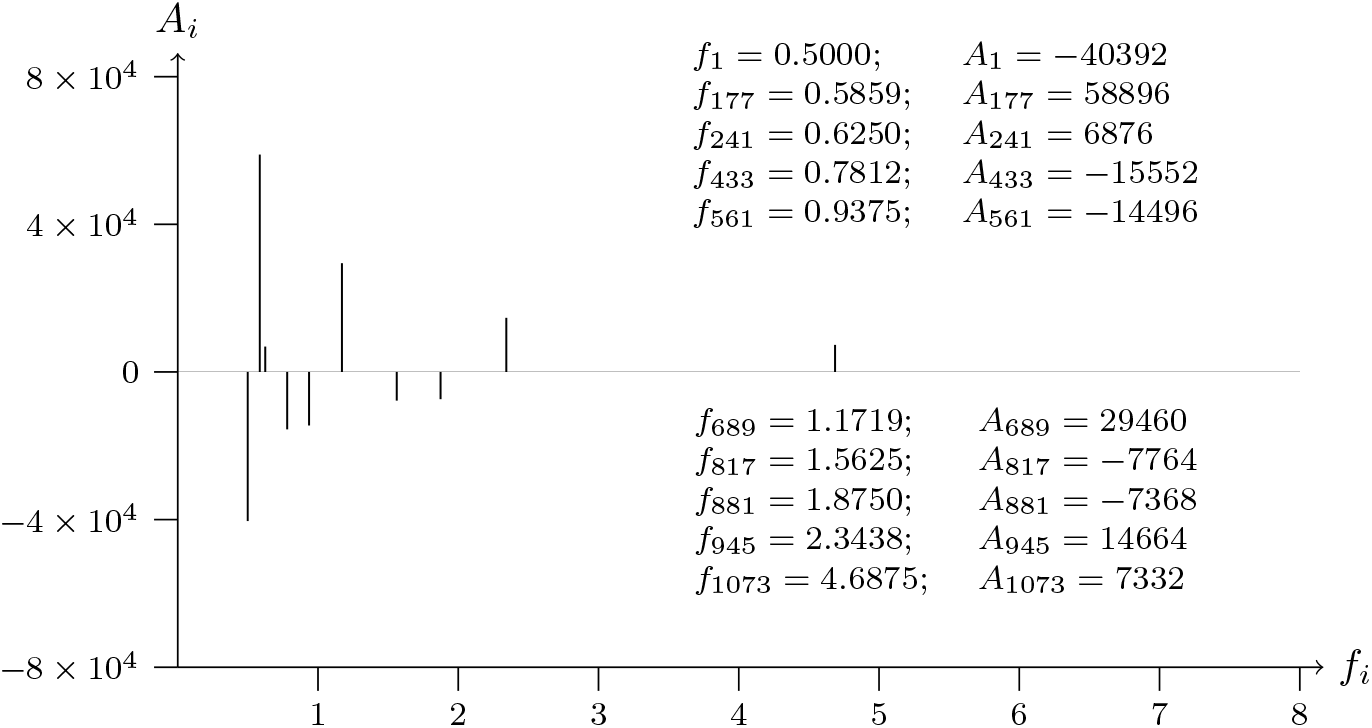
T2T-CHM13v2.0; chromosome 1; p telomere; analyzed data from 1500 to 2699

**Figure 34a:**
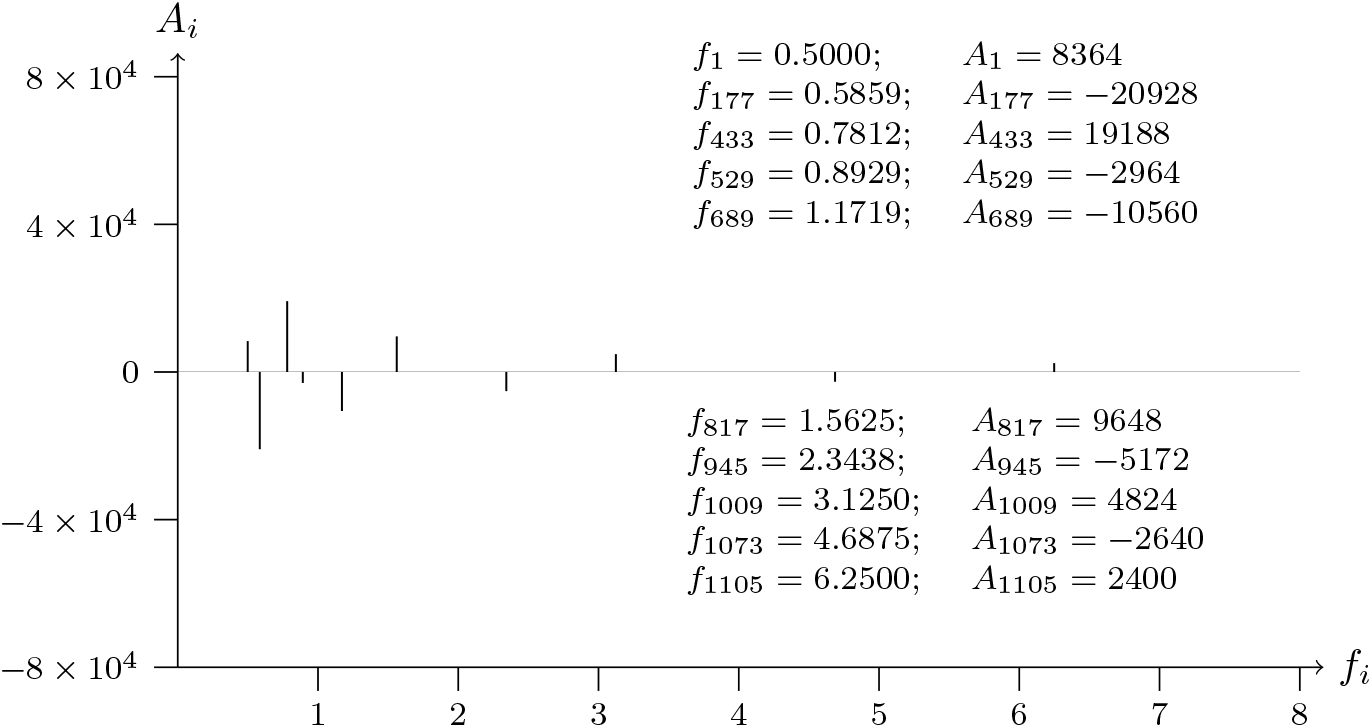
T2T-CHM13v2.0; chromosome 1; p telomere; analyzed data from 1800 to 3000

**Figure 34b:**
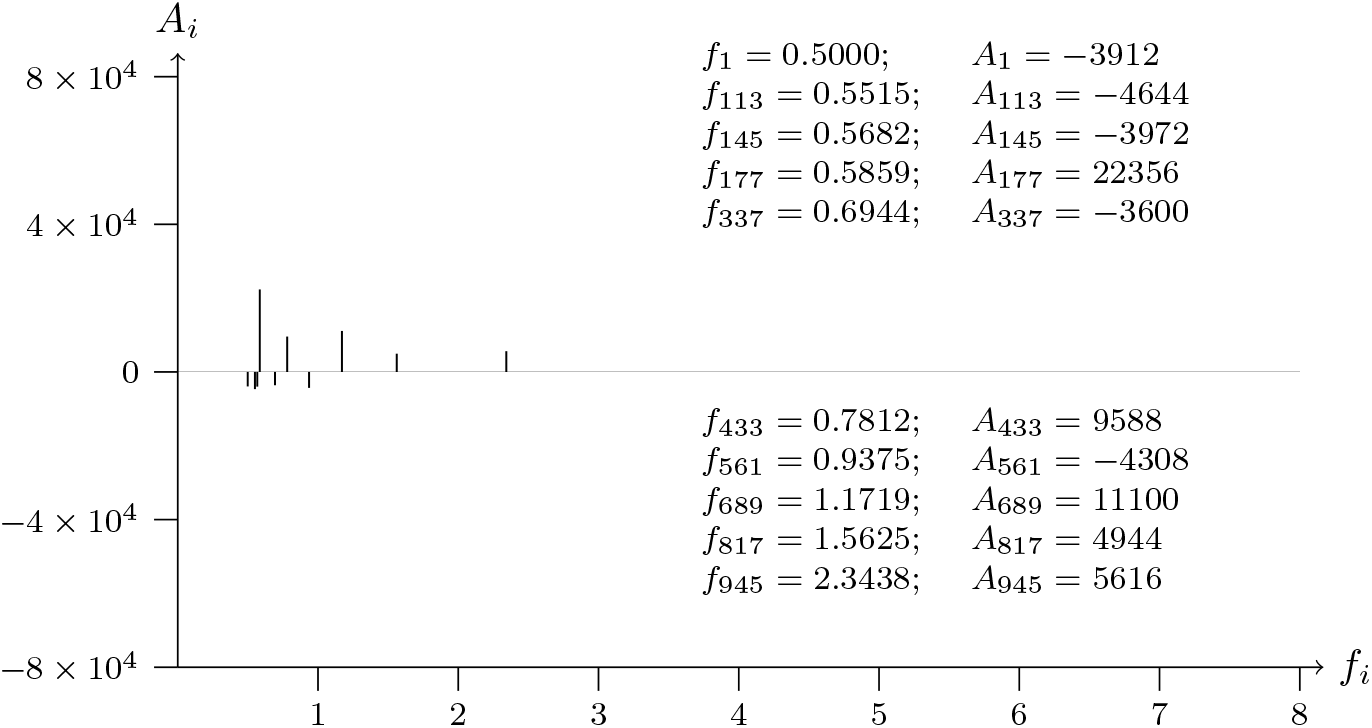
T2T-CHM13v2.0; chromosome 1; p telomere; analyzed data from 1800 to 3000

**Figure 35a:**
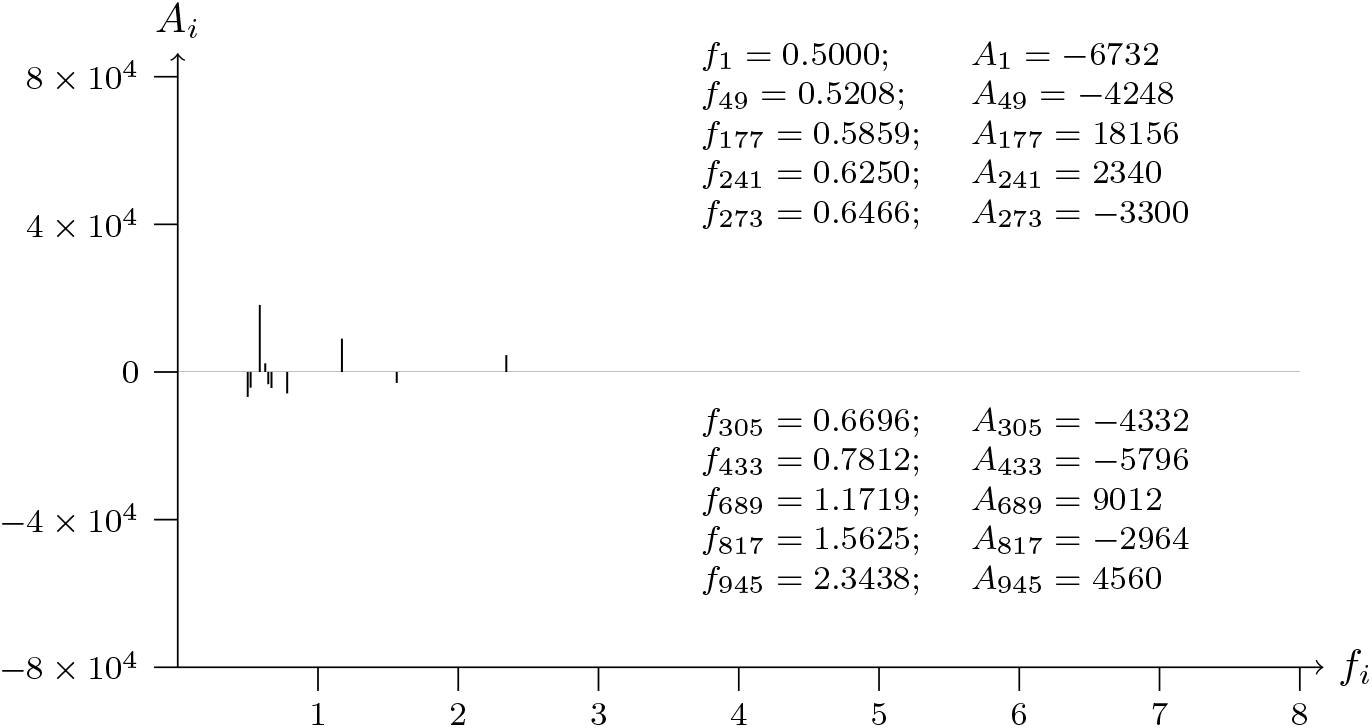
T2T-CHM13v2.0; chromosome 1; q telomere; analyzed data from 0 to 1199

**Figure 35b:**
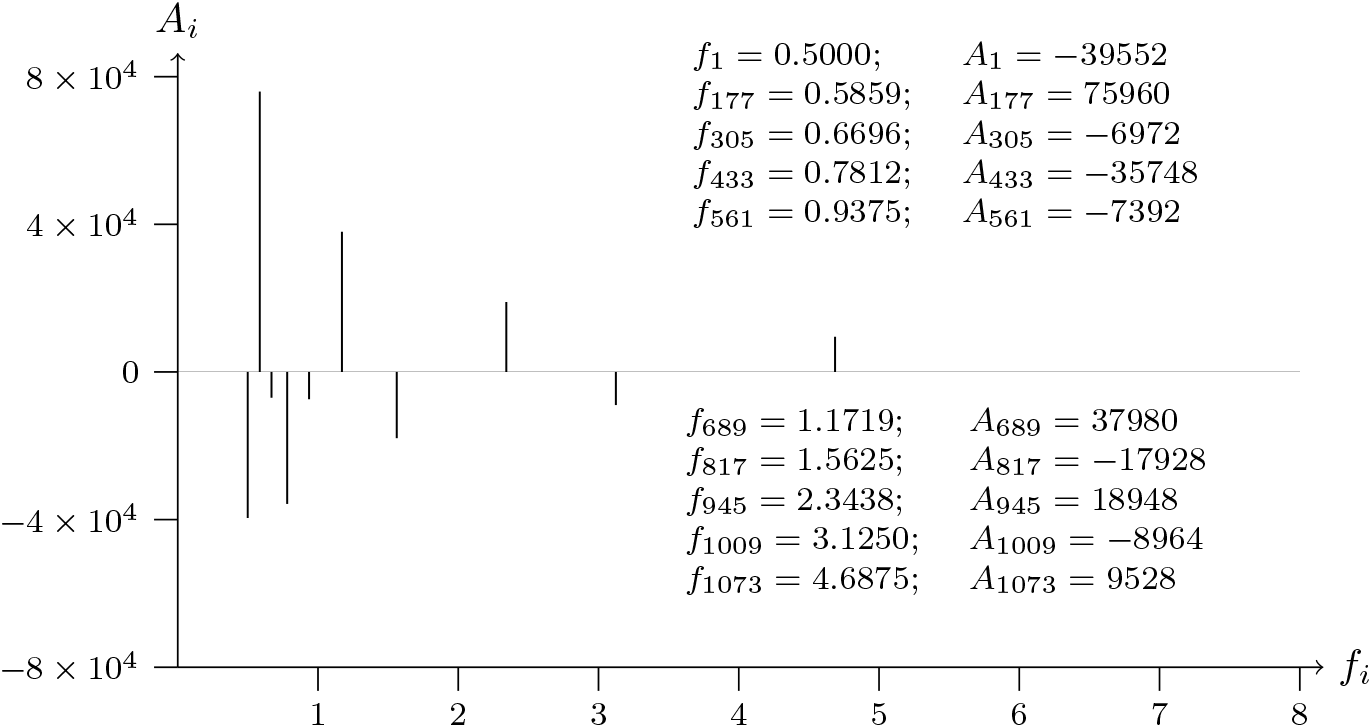
T2T-CHM13v2.0; chromosome 1; q telomere; analyzed data from 0 to 1199

**Figure 36a:**
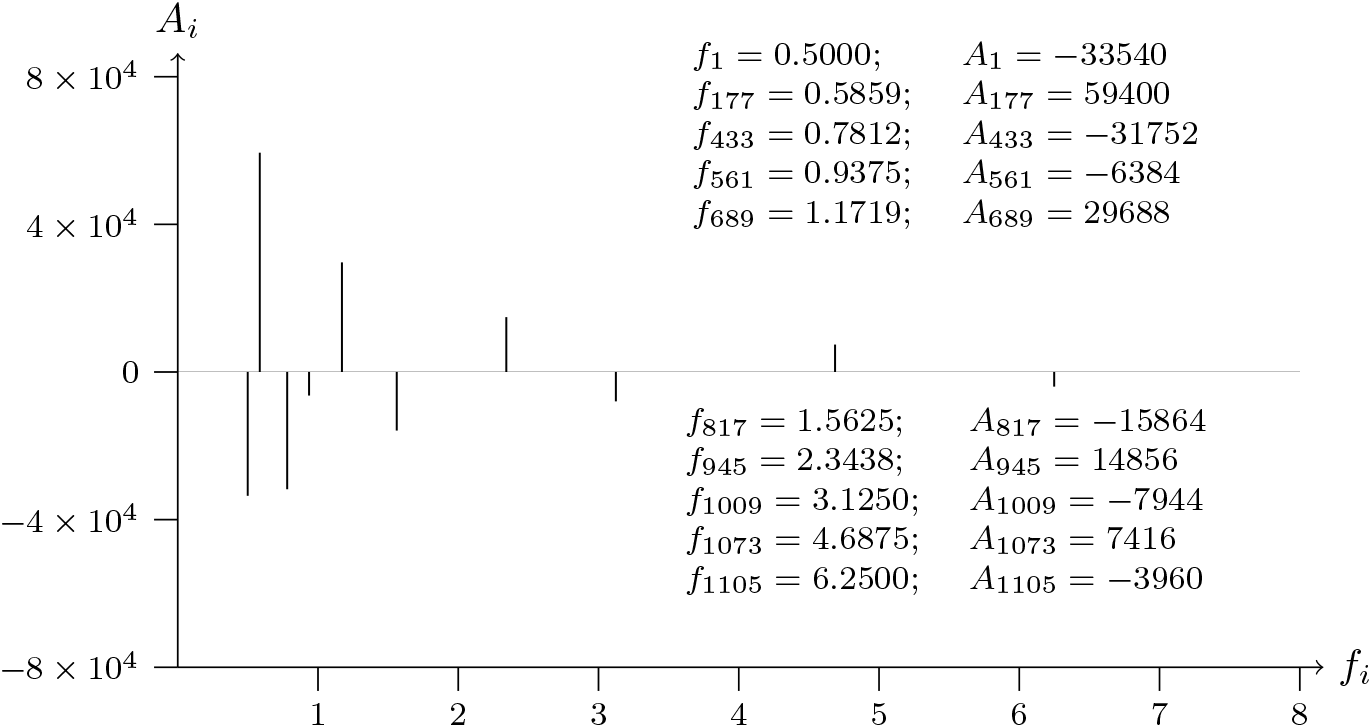
T2T-CHM13v2.0; chromosome 1; q telomere; analyzed data from 1664 to 2863

**Figure 36b:**
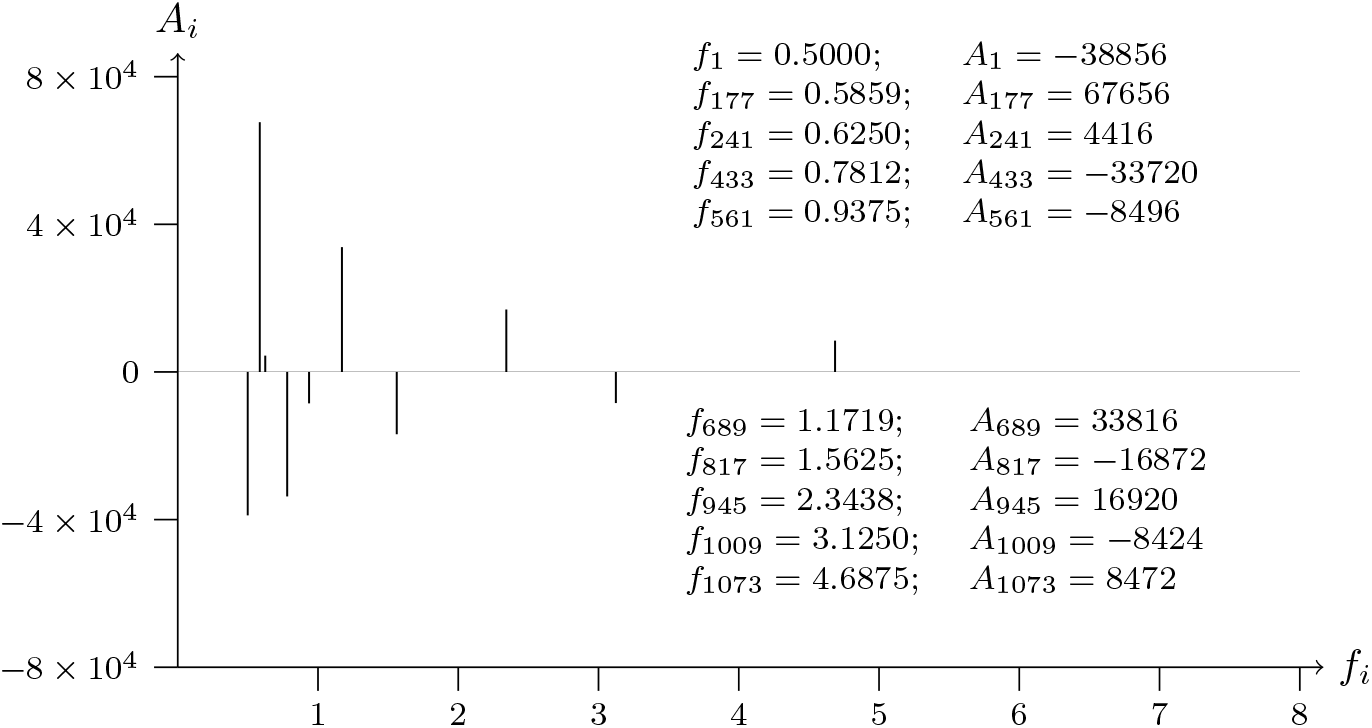
T2T-CHM13v2.0; chromosome 1; q telomere; analyzed data from 1664 to 2863

**Figure 37a:**
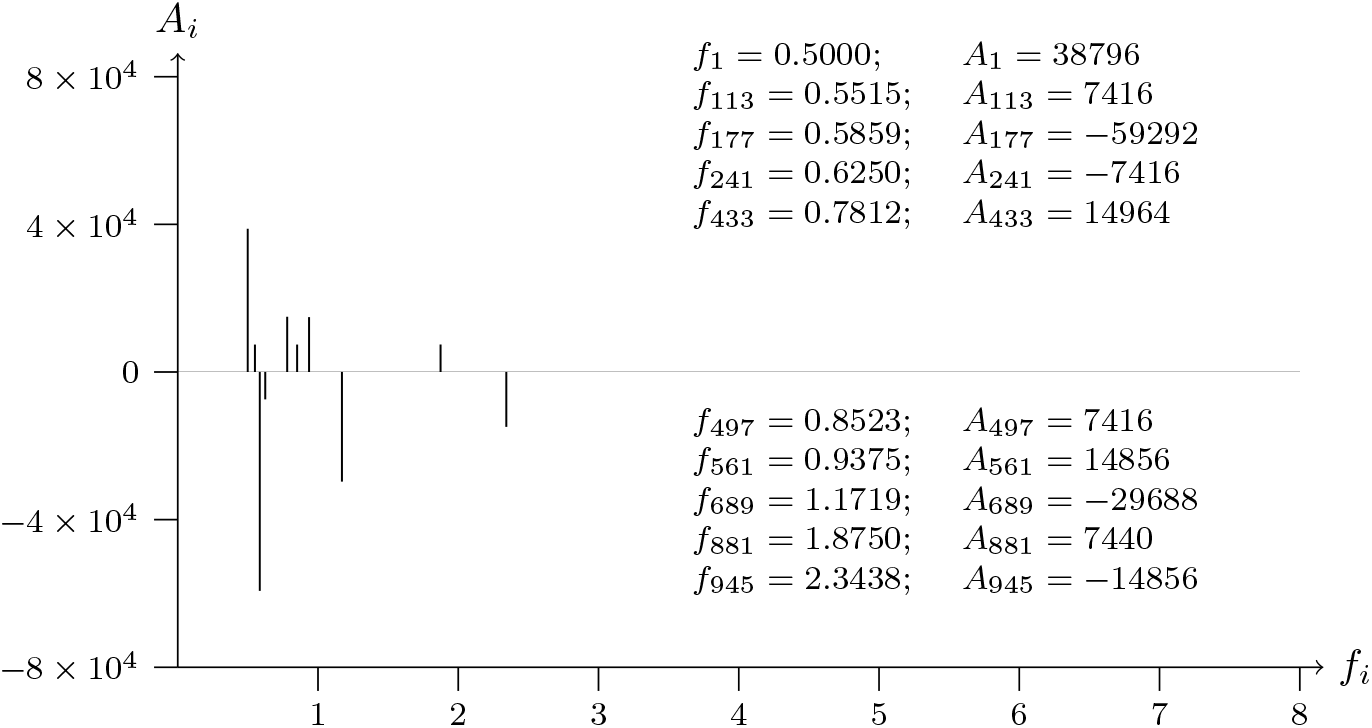
T2T-CHM13v2.0; chromosome 1; q telomere; analyzed data from 2128 to 3328

**Figure 37b:**
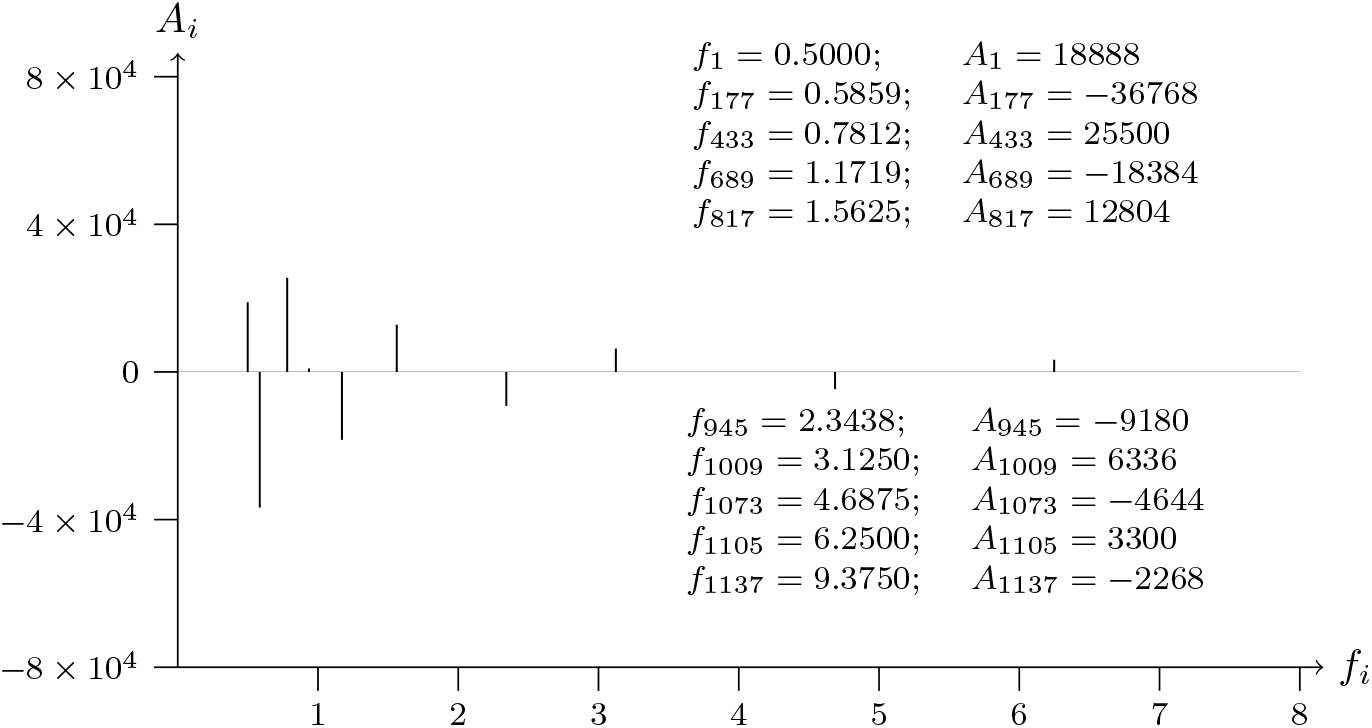
T2T-CHM13v2.0; chromosome 1; q telomere; analyzed data from 2128 to 3328

## 4 Biological analysis SWT for genes

**Figure 38a:**
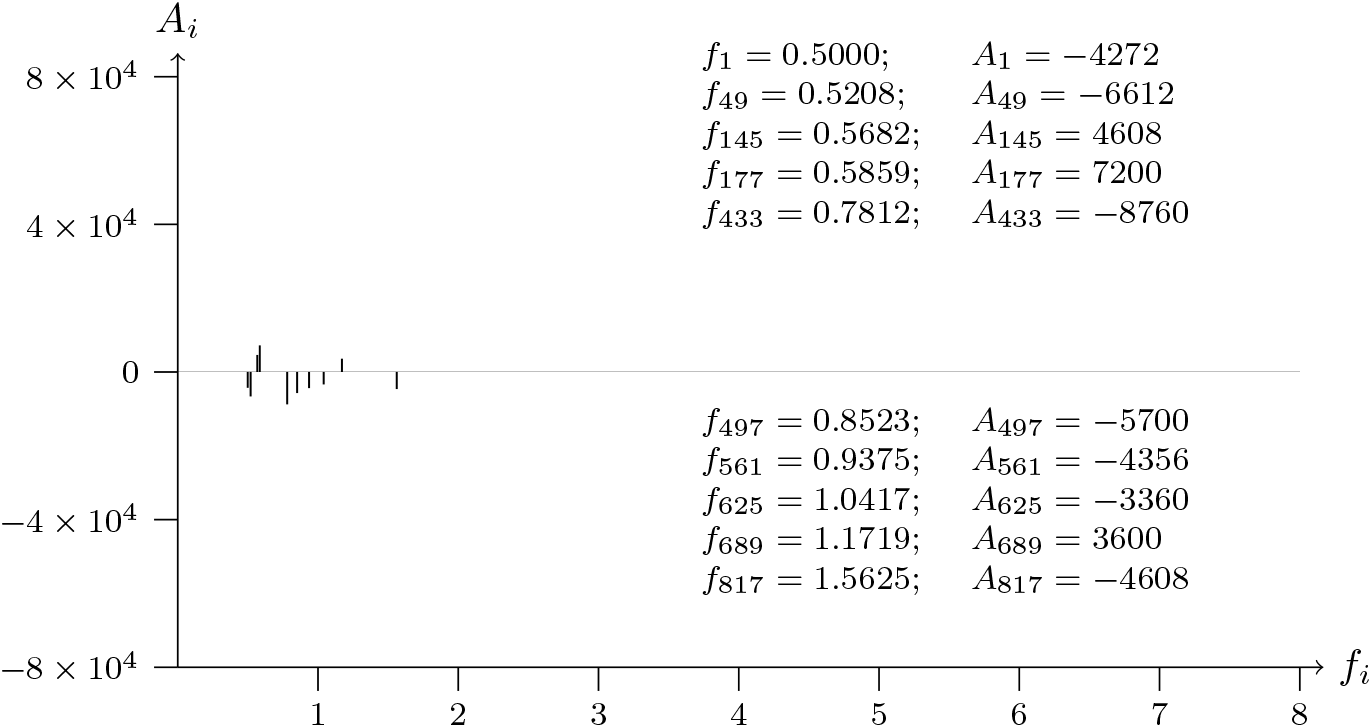
T2T-CHM13v2.0; chromosome 1; RUSC1 (oncogene) exon-NM 001105203.2-2 analyzed data from 154460228 to 15446142811111

**Figure 38b:**
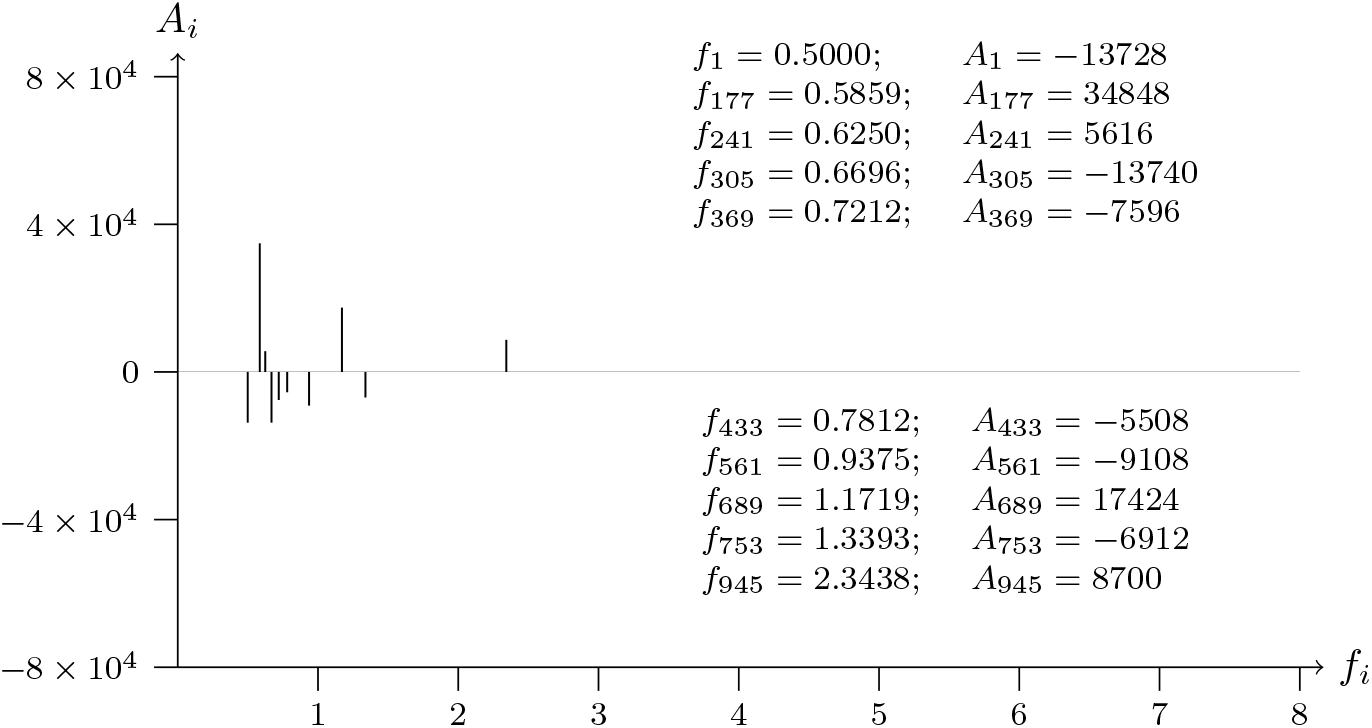
T2T-CHM13v2.0; chromosome 1; RUSC1 (oncogene) exon-NM 001105203.2-2 analyzed data from 154461428 to 154460228

**Figure 39a:**
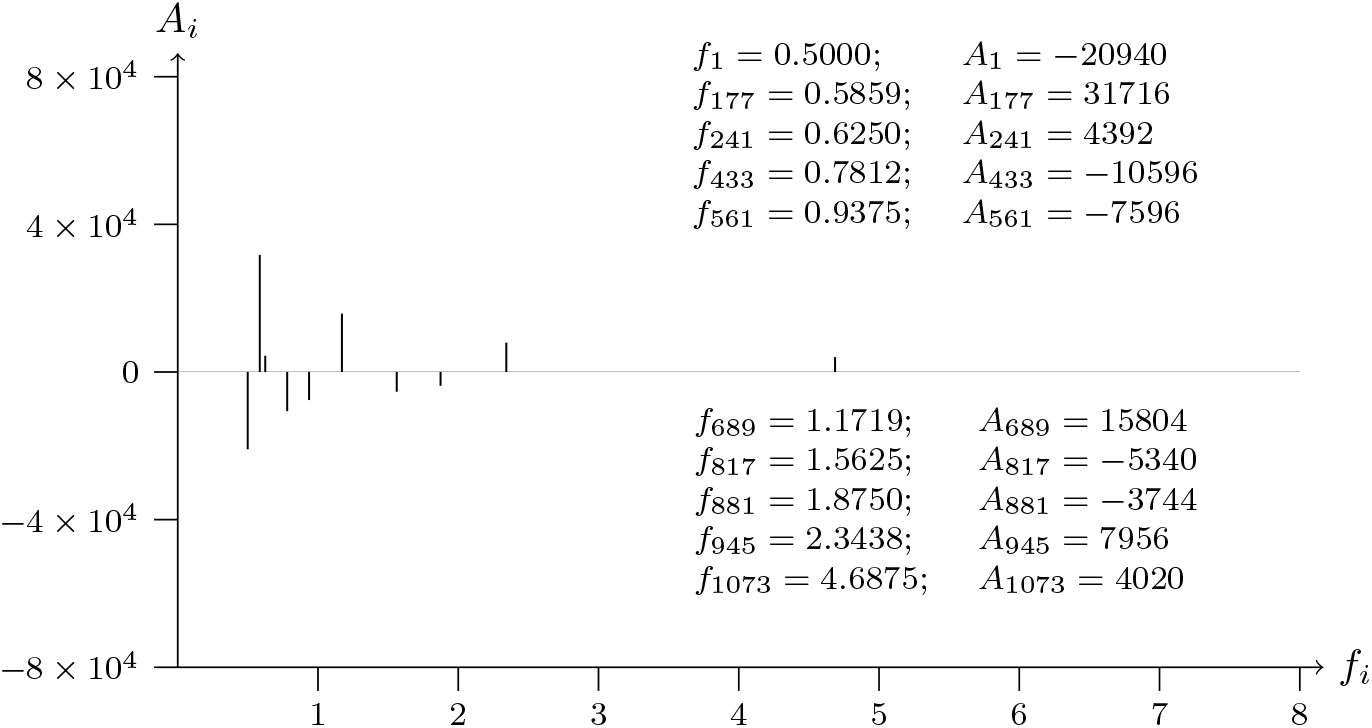
T2T-CHM13v2.0; chromosome 1; RUSC1 (oncogene) intron-NM 001105203.2-2 analyzed data from 154461670 to 154462870

**Figure 39b:**
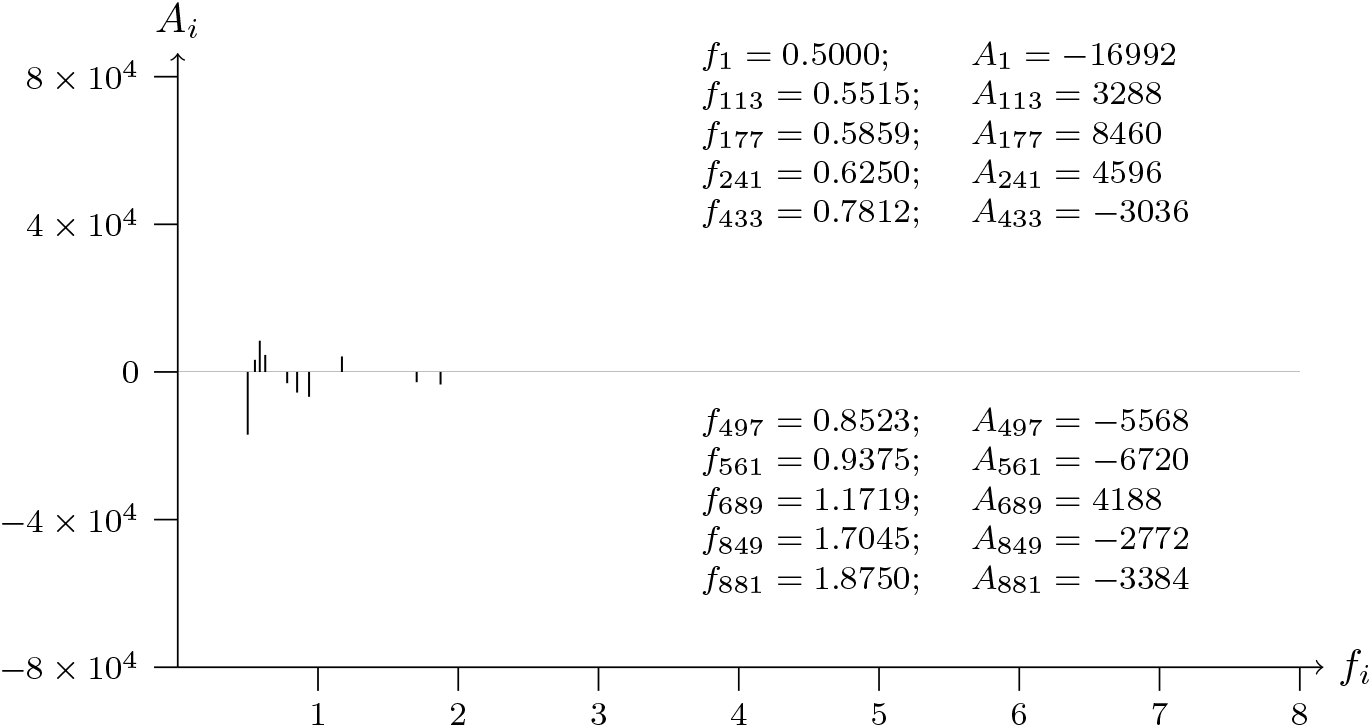
T2T-CHM13v2.0; chromosome 1; RUSC1 (oncogene) intron-NM 001105203.2-2 analyzed data from 154462870 to 154461670

**Figure 40a:**
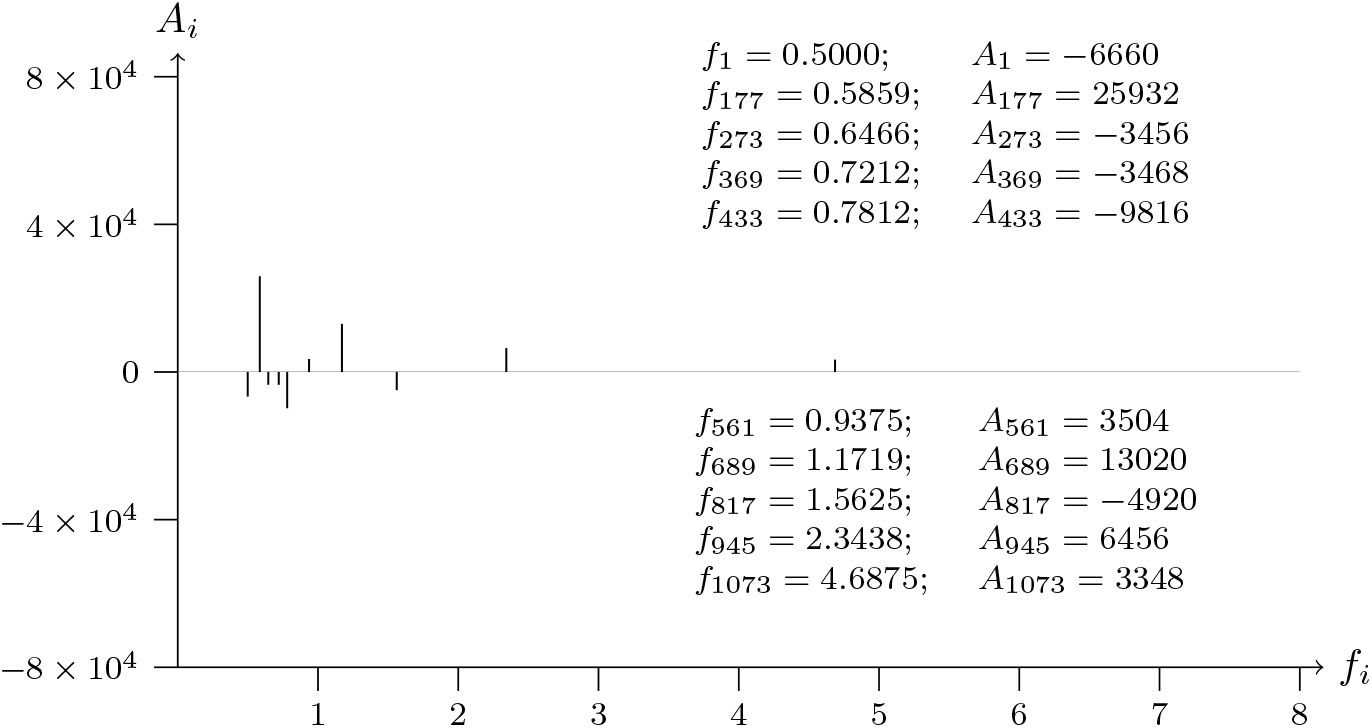
T2T-CHM13v2.0; chromosome 1; RUSC1 (oncogene) intron-NM 001105203.2-9 analyzed data from 154466815 to 154468015

**Figure 40b:**
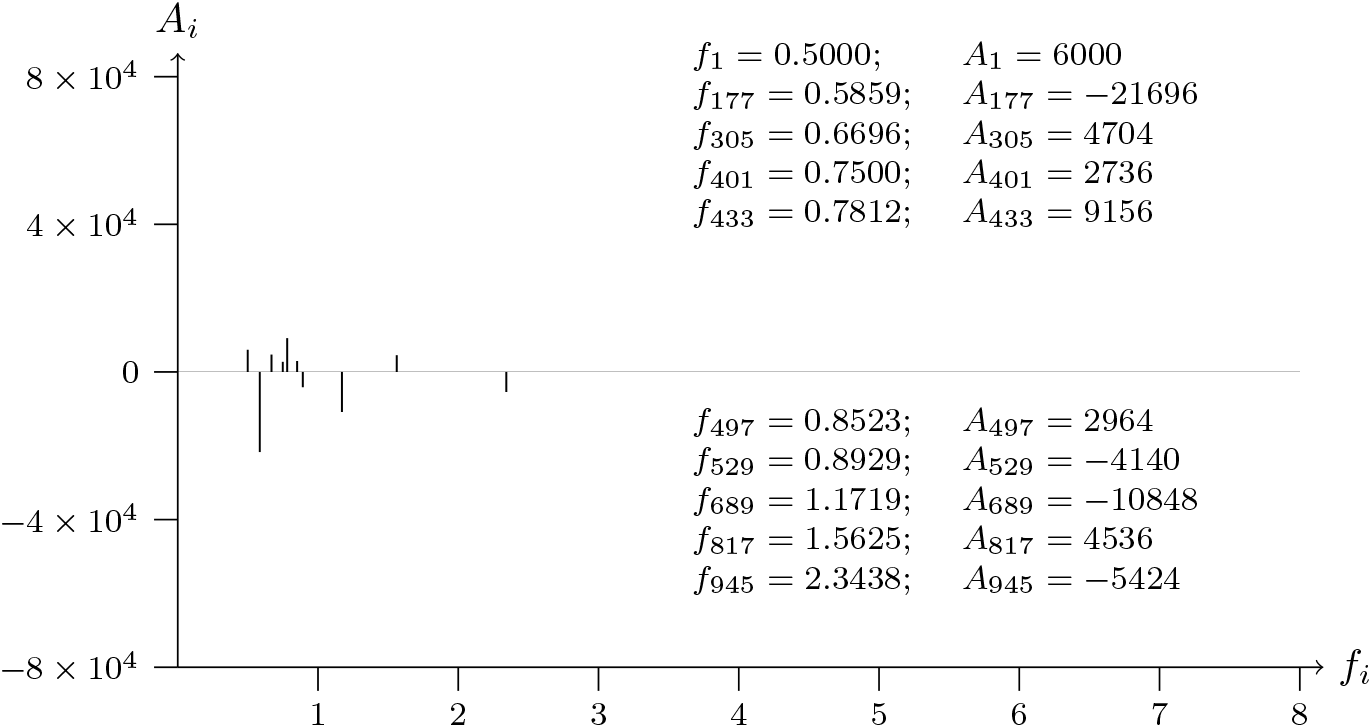
T2T-CHM13v2.0; chromosome 1; RUSC1 (oncogene) intron-NM 001105203.2-9 analyzed data from 154468015 to 154466815

**Figure 41a:**
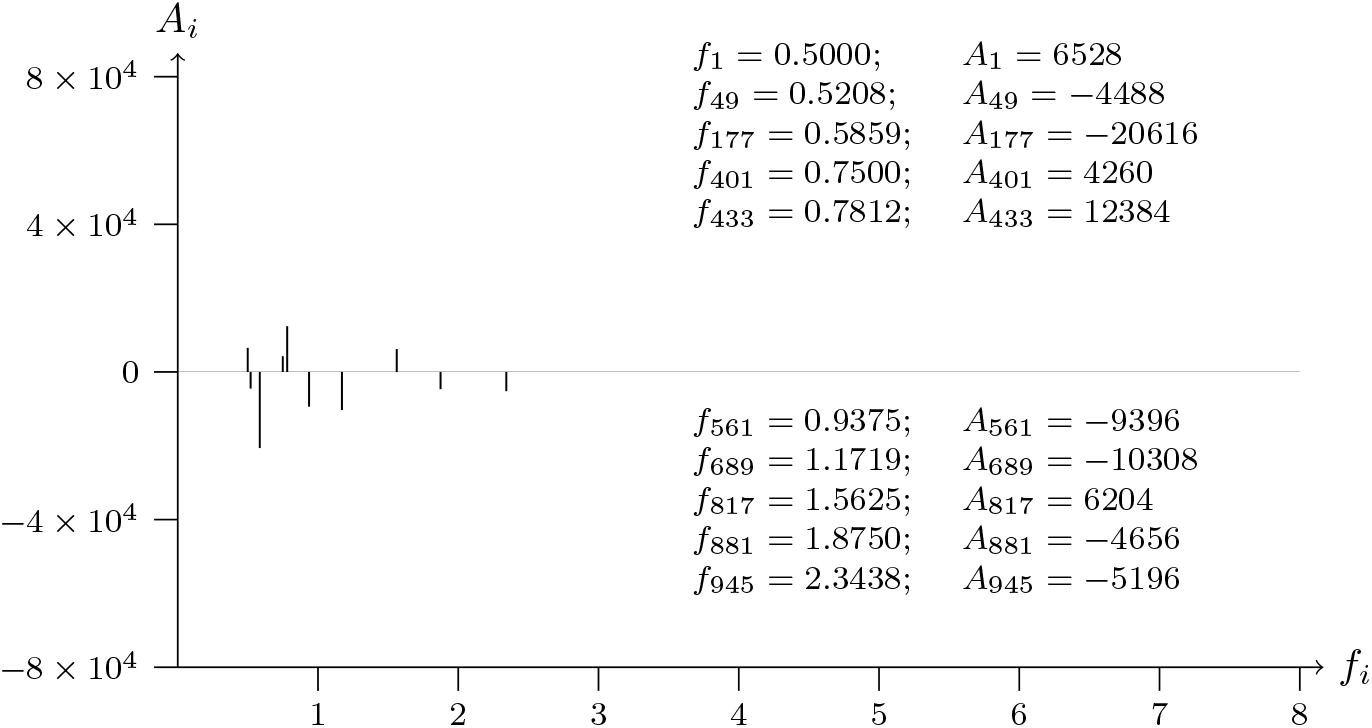
T2T-CHM13v2.0; chromosome 1; GPR161 exon-NM 001349635.1-5 analyzed data from 167431007 to 167432207

**Figure 41b:**
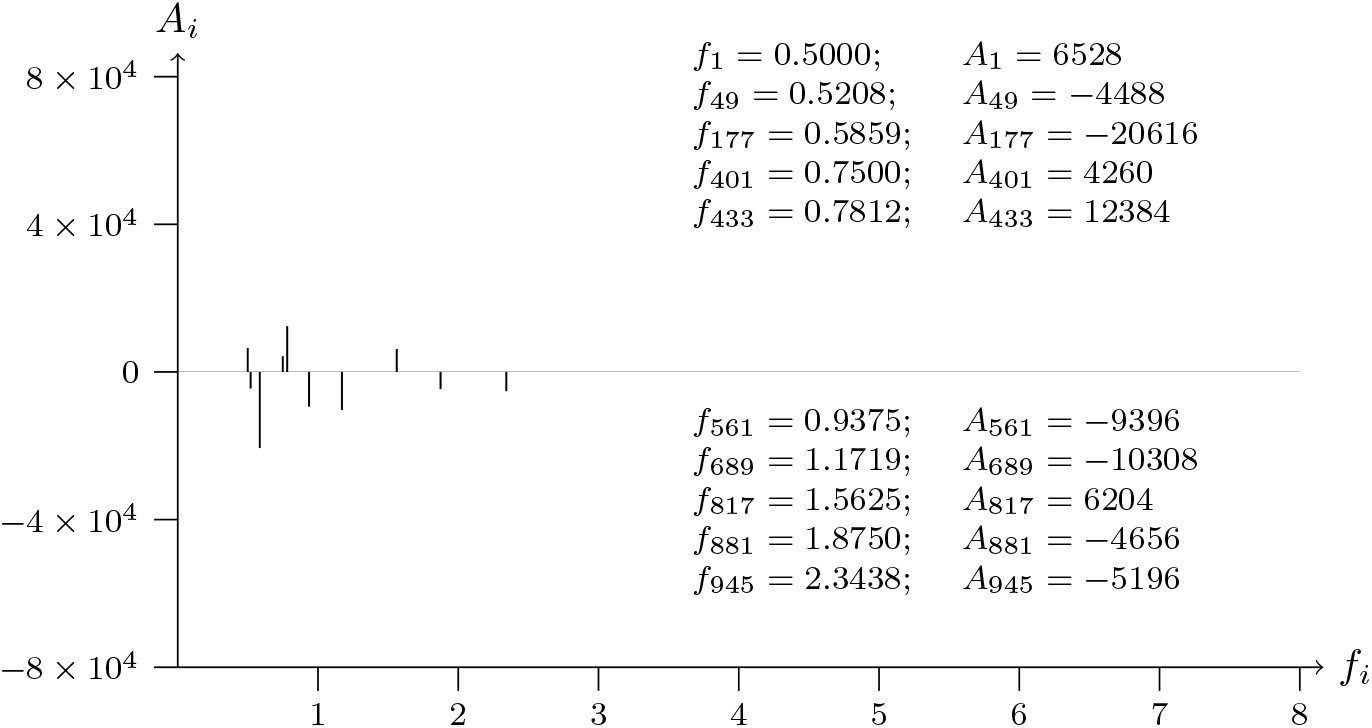
T2T-CHM13v2.0; chromosome 1; GPR161 exon-NM 001349635.1-5 analyzed data from 167432207 to 167431007

**Figure 42a:**
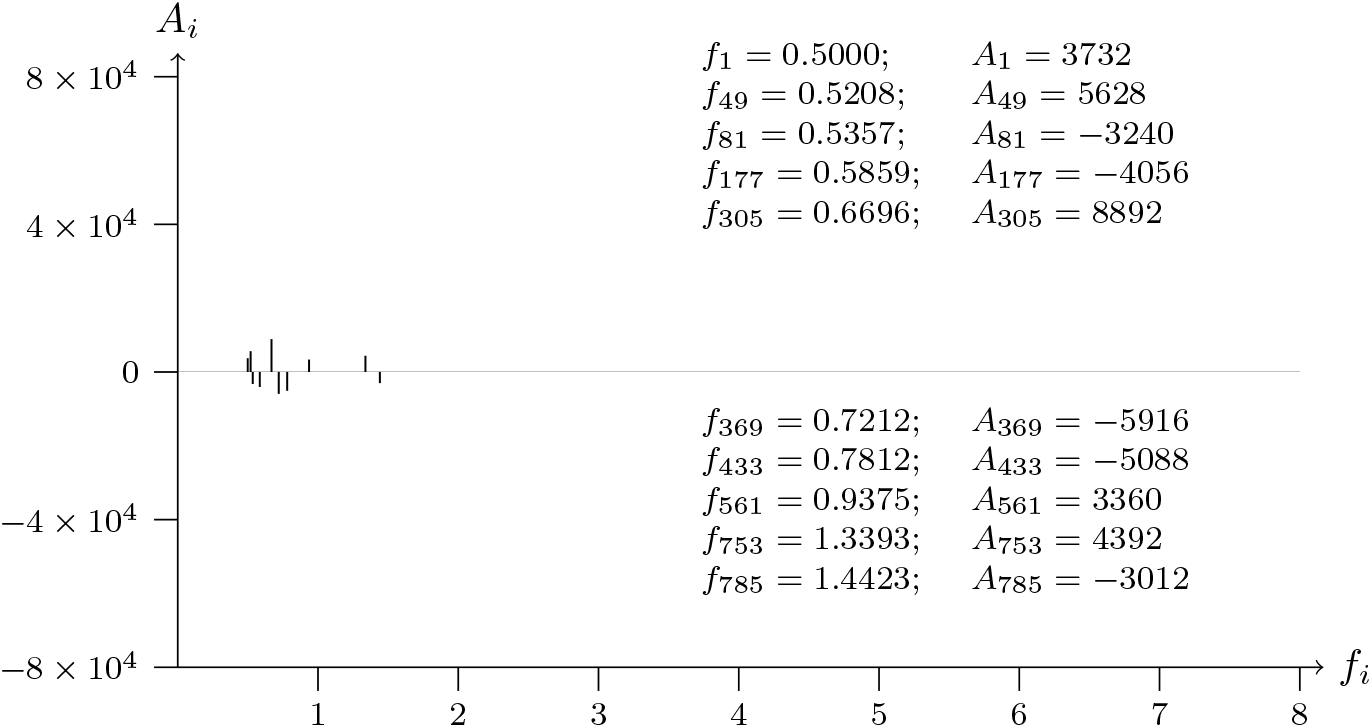
T2T-CHM13v2.0; chromosome 17; BRCA1 (tumor supressor) exon-NM 001408458.1-22 analyzed data from 43902857 to 43904057

**Figure 42b:**
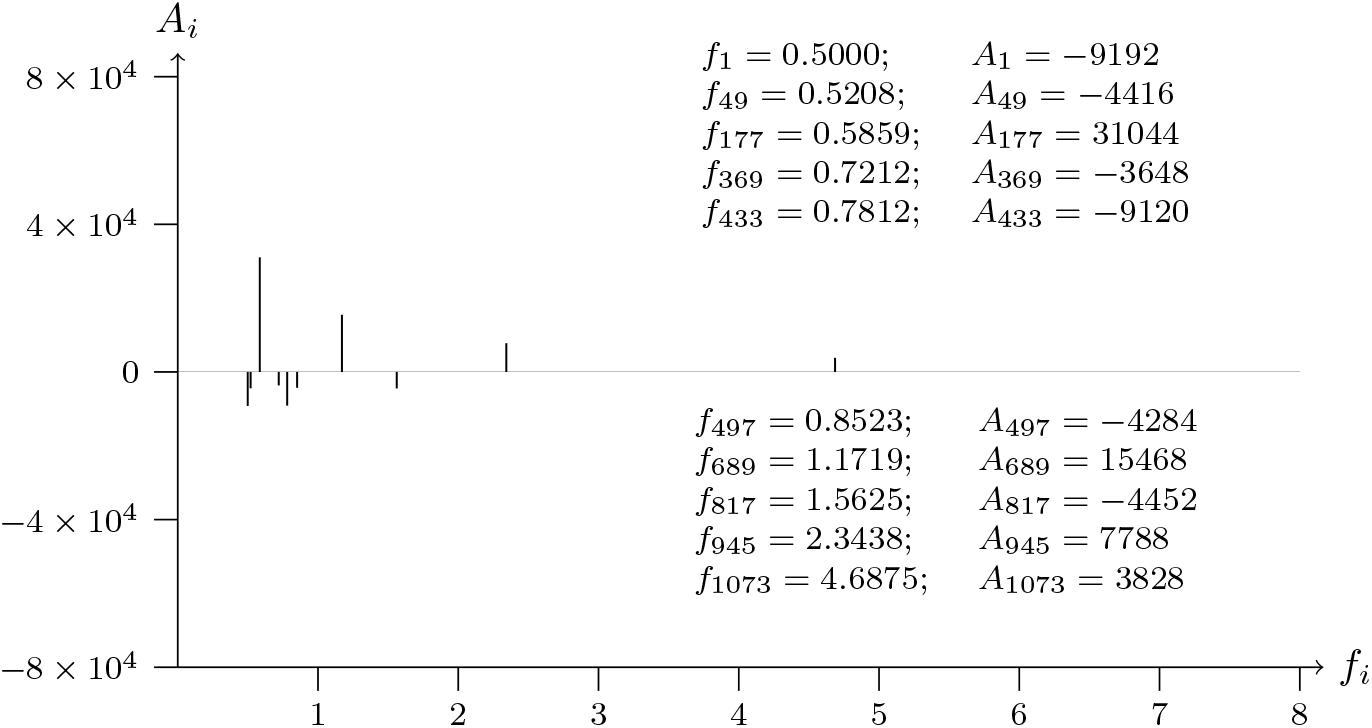
T2T-CHM13v2.0; chromosome 17; BRCA1 (tumor supressor) exon-NM 001408458.1-22 analyzed data from 43904057 to 43902857

**Figure 43a:**
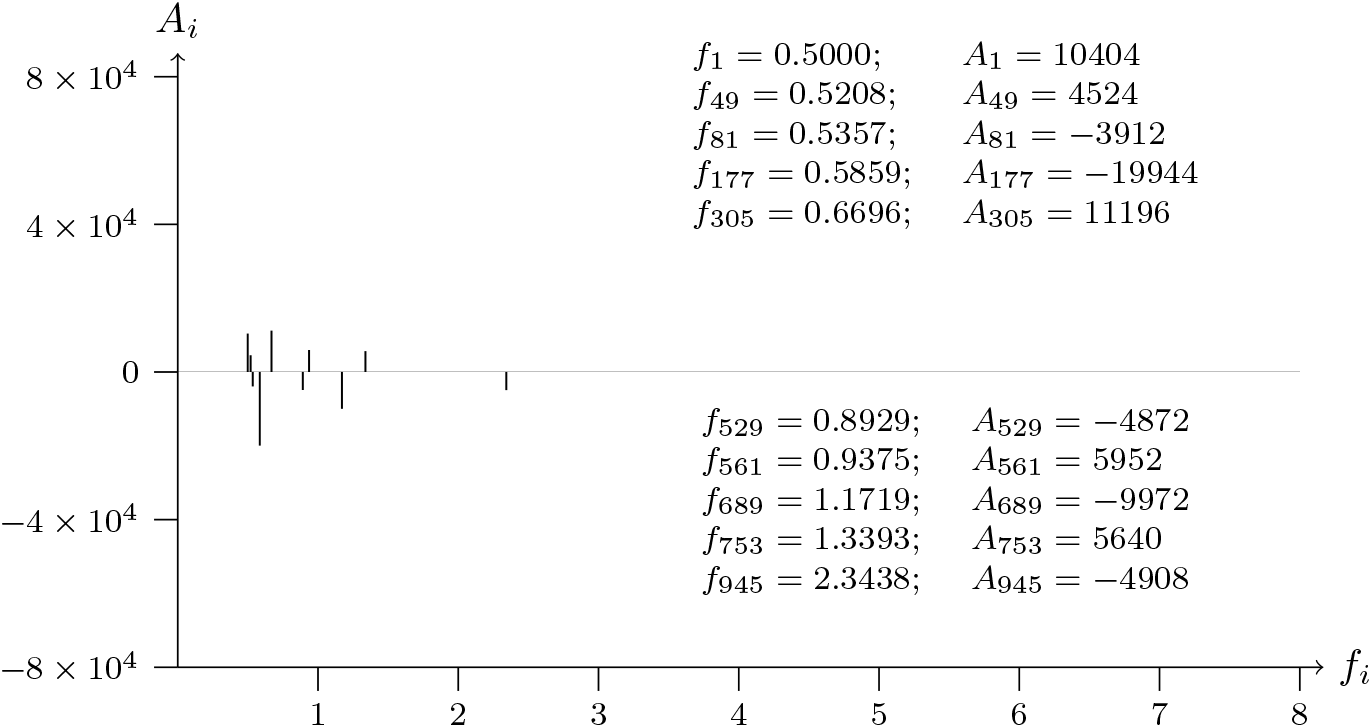
T2T-CHM13v2.0; chromosome 17; BRCA1 (tumor supressor) intron-NM 001408458.1-22 analyzed data from 43904362 to 43905562

**Figure 43b:**
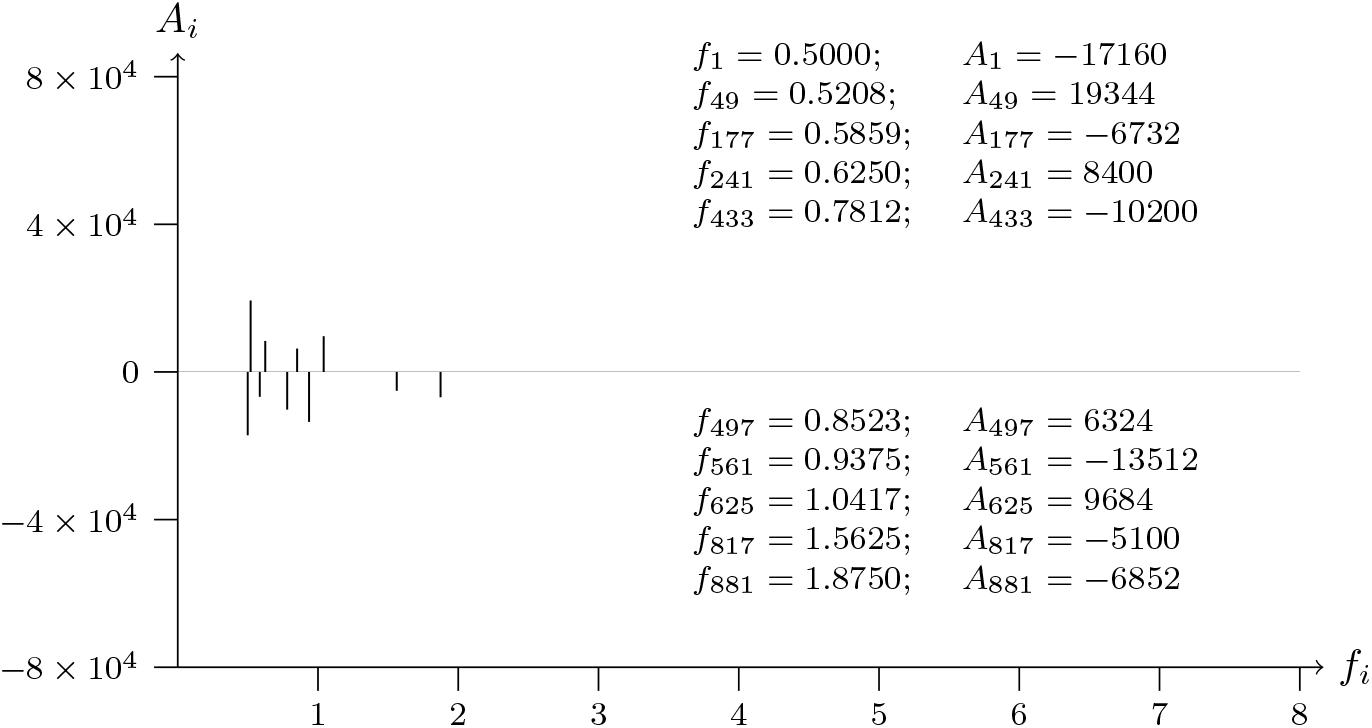
T2T-CHM13v2.0; chromosome 17; BRCA1 (tumor supressor) intron-NM 001408458.1-22 analyzed data from 43905562 to 43904362

**Figure 44a:**
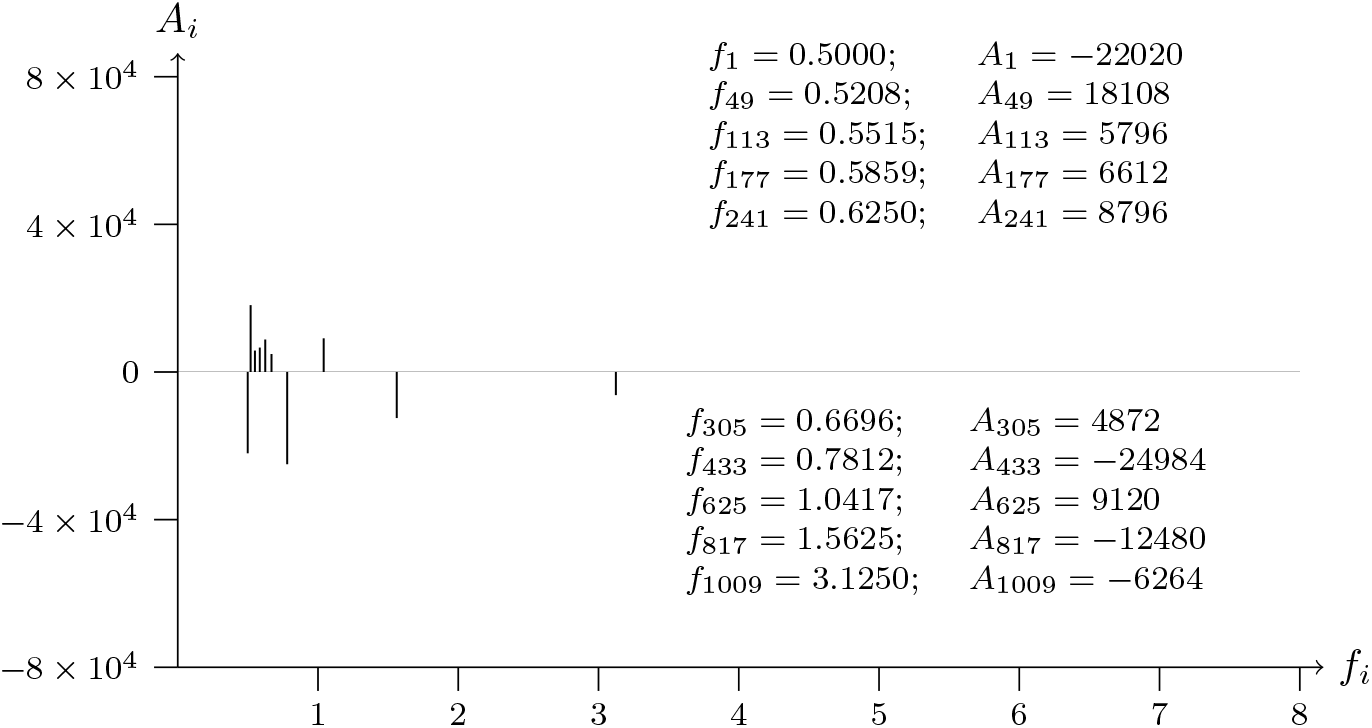
T2T-CHM13v2.0; chromosome 13; BRCA2 (tumor supressor) exon-NM 000059.4-11 analyzed data from 31553507 to 31554707

**Figure 44b:**
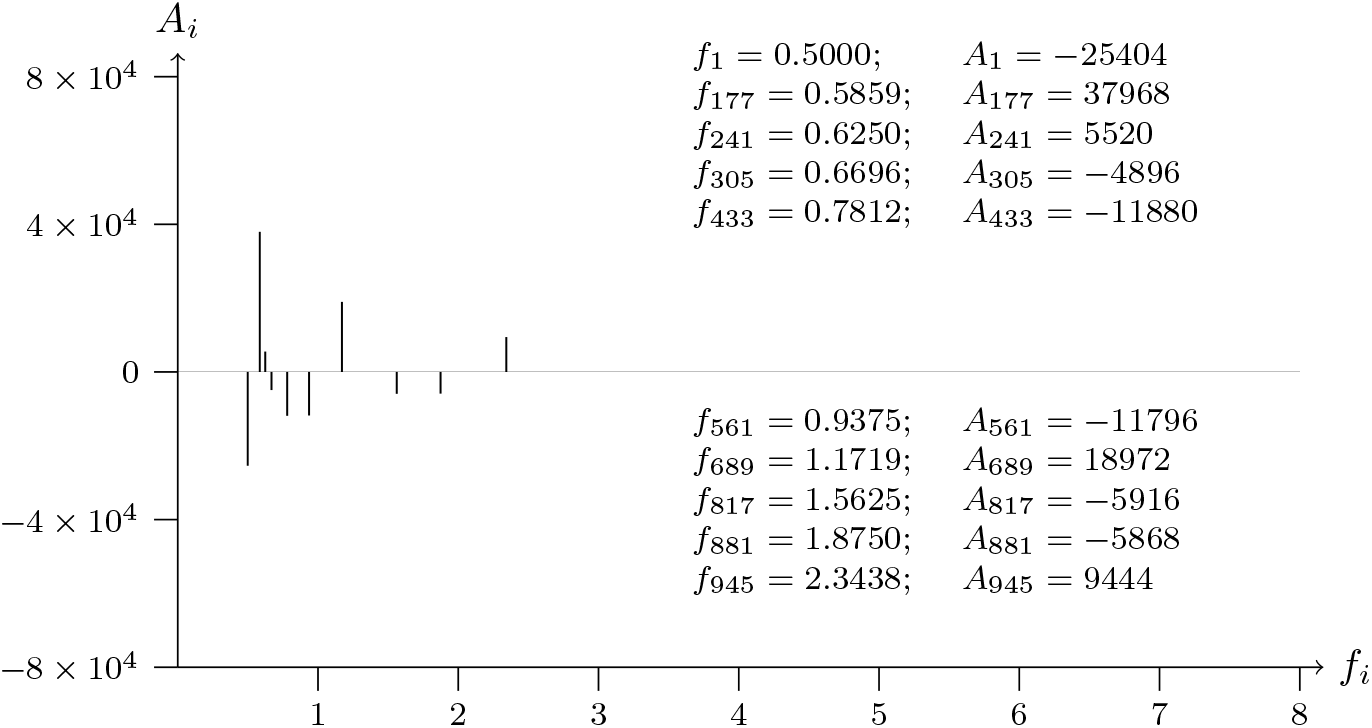
T2T-CHM13v2.0; chromosome 13; BRCA2 (tumor supressor) exon-NM 000059.4-11 analyzed data from 31554707 to 31553507

**Figure 45a:**
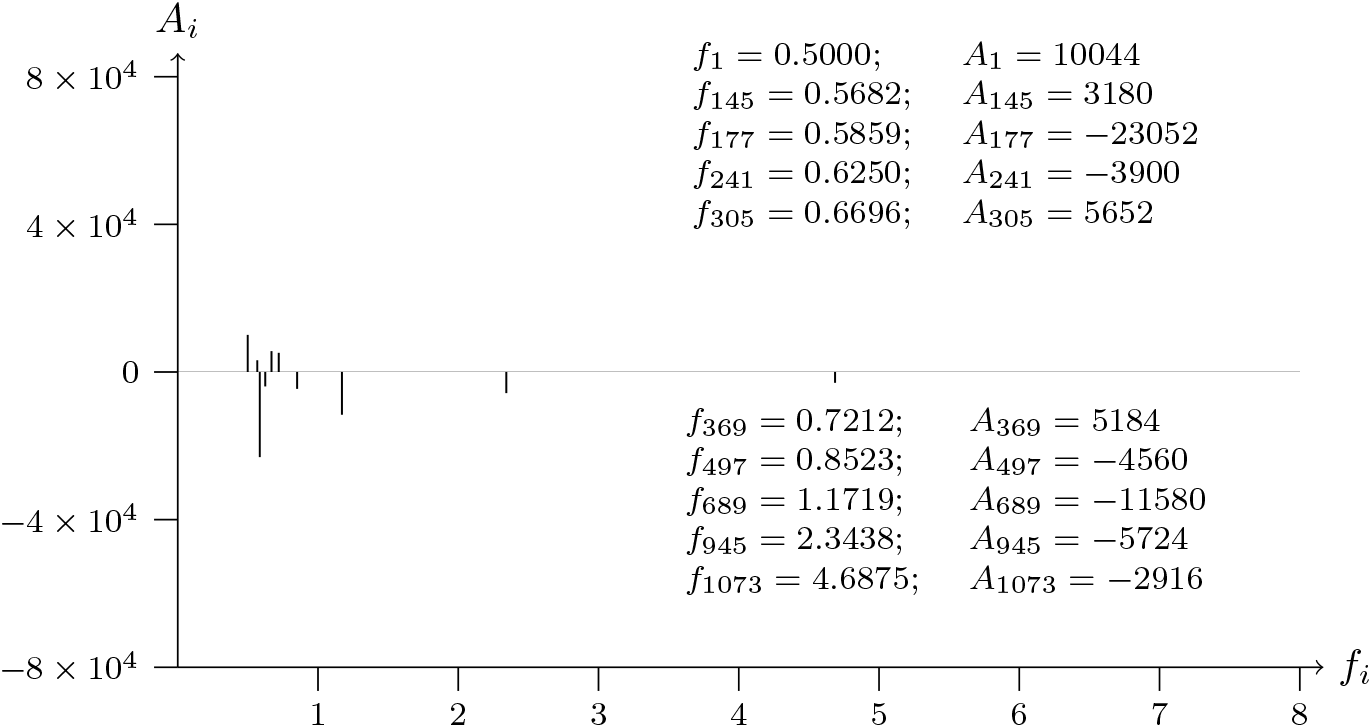
T2T-CHM13v2.0; chromosome 13; BRCA2 (tumor supressor) exon-NM 000059.4-27 analyzed data from 31615404 to 31616604

**Figure 45b:**
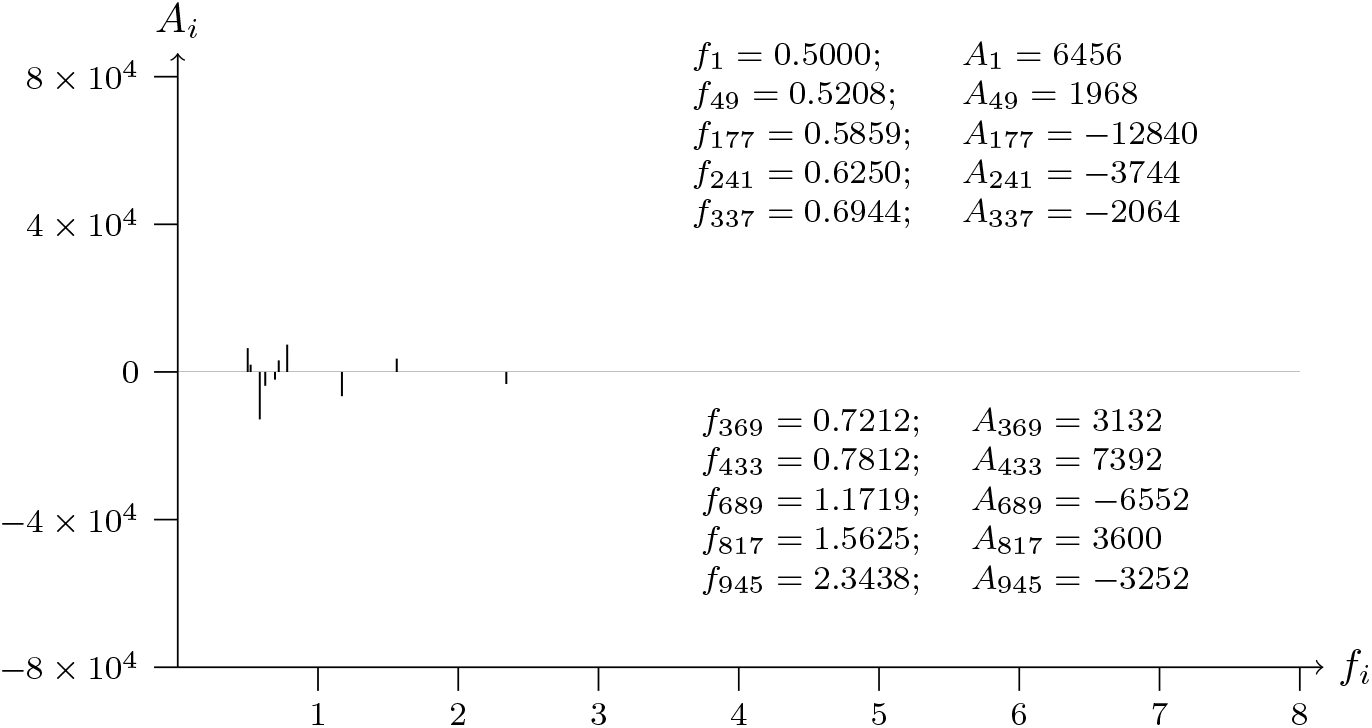
T2T-CHM13v2.0; chromosome 13; BRCA2 (tumor supressor) exon-NM 000059.4-27 analyzed data from 31616604 to 31615404

**Figure 46a:**
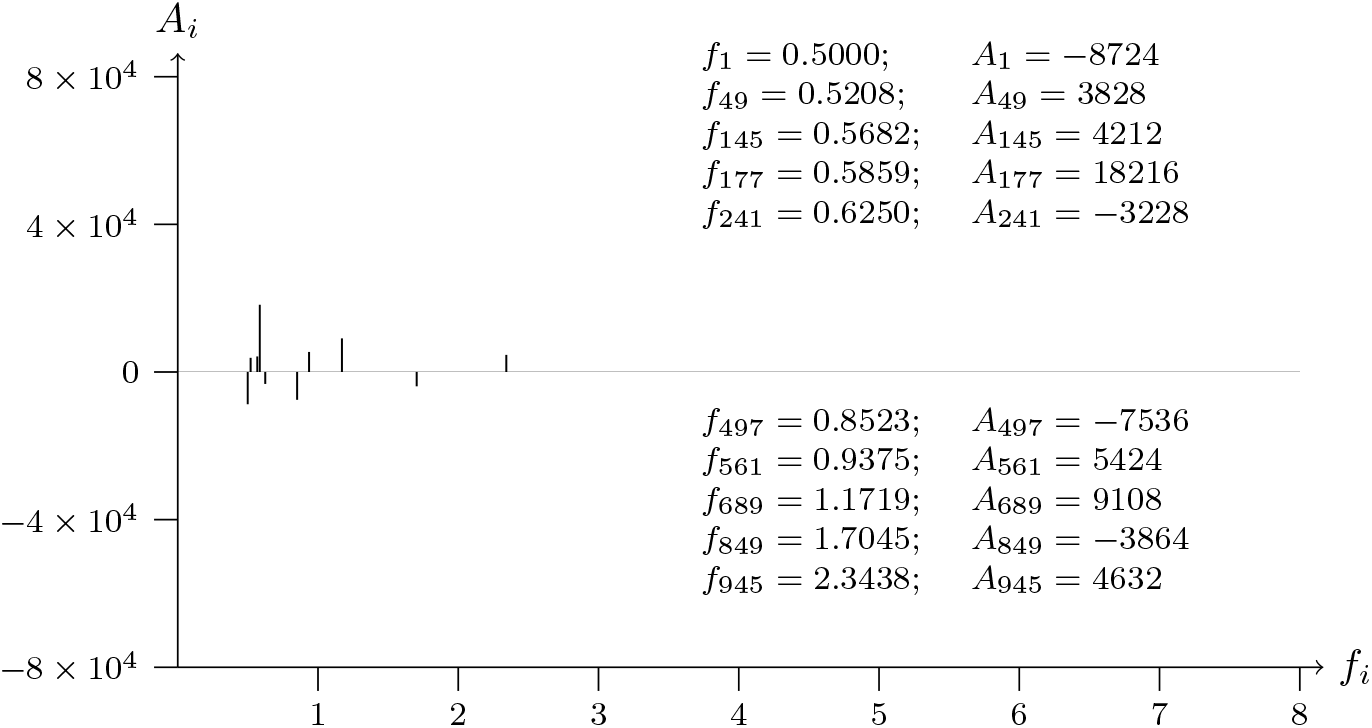
T2T-CHM13v2.0; chromosome 7; EGFR (oncogene) exon-NM 005228.5-28 analyzed data from 55365684 to 55366884

**Figure 46b:**
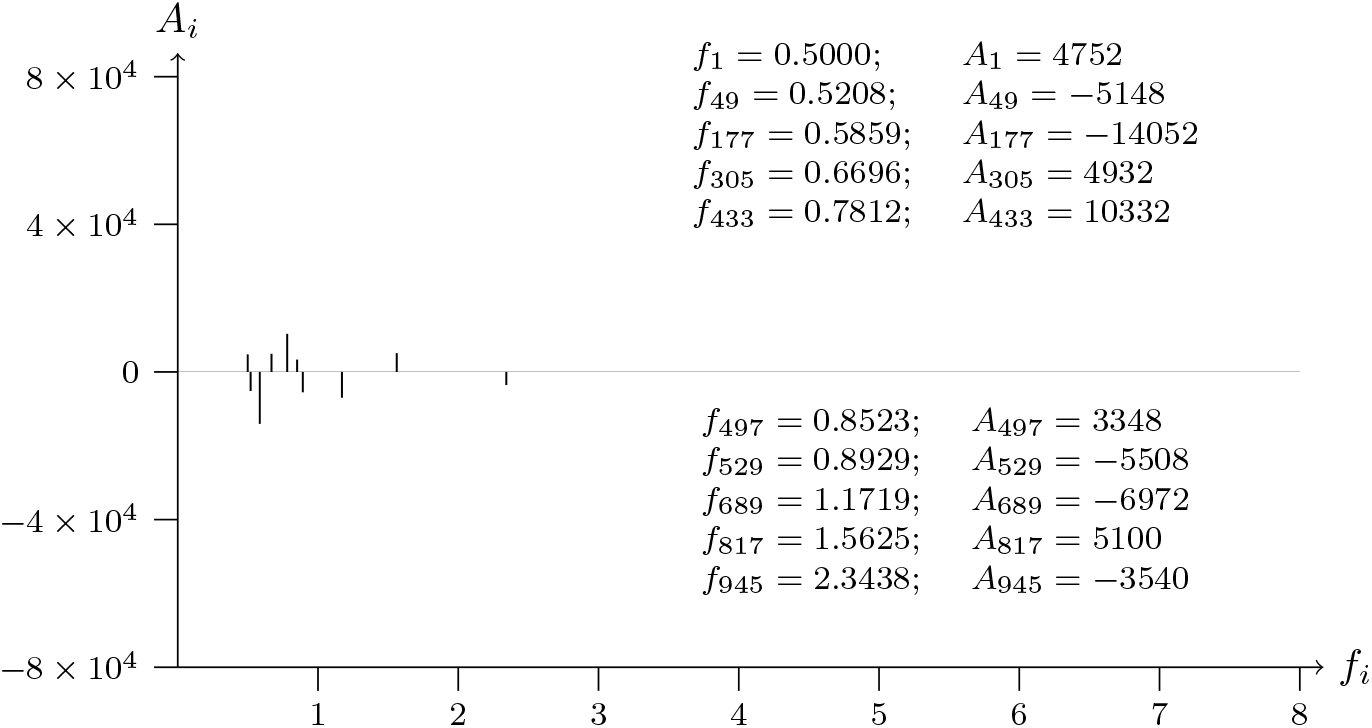
T2T-CHM13v2.0; chromosome 7; EGFR (oncogene) exon-NM 005228.5-28 analyzed data from 55366884 to 55365684

**Figure 47a:**
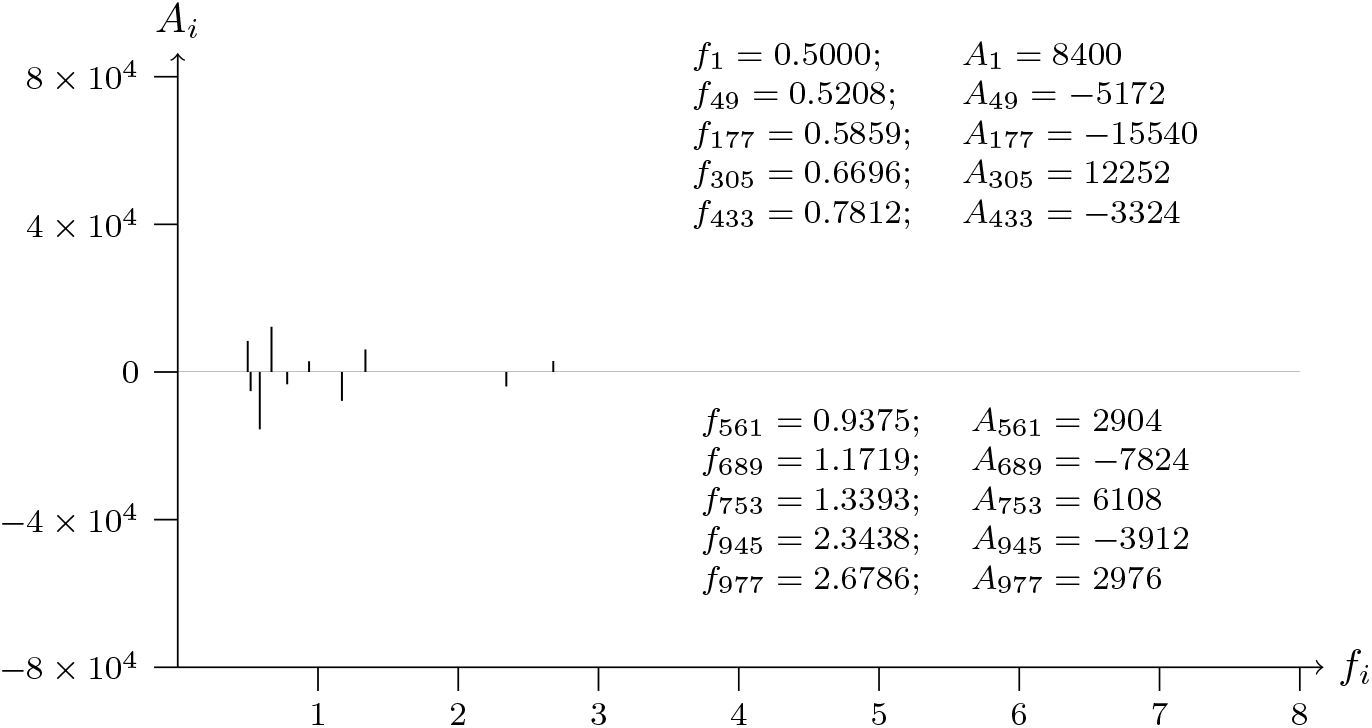
T2T-CHM13v2.0; chromosome 1; ELK4 (oncogene) exon-NM 001973.4-5 analyzed data from 204872578 to 204873778

**Figure 47b:**
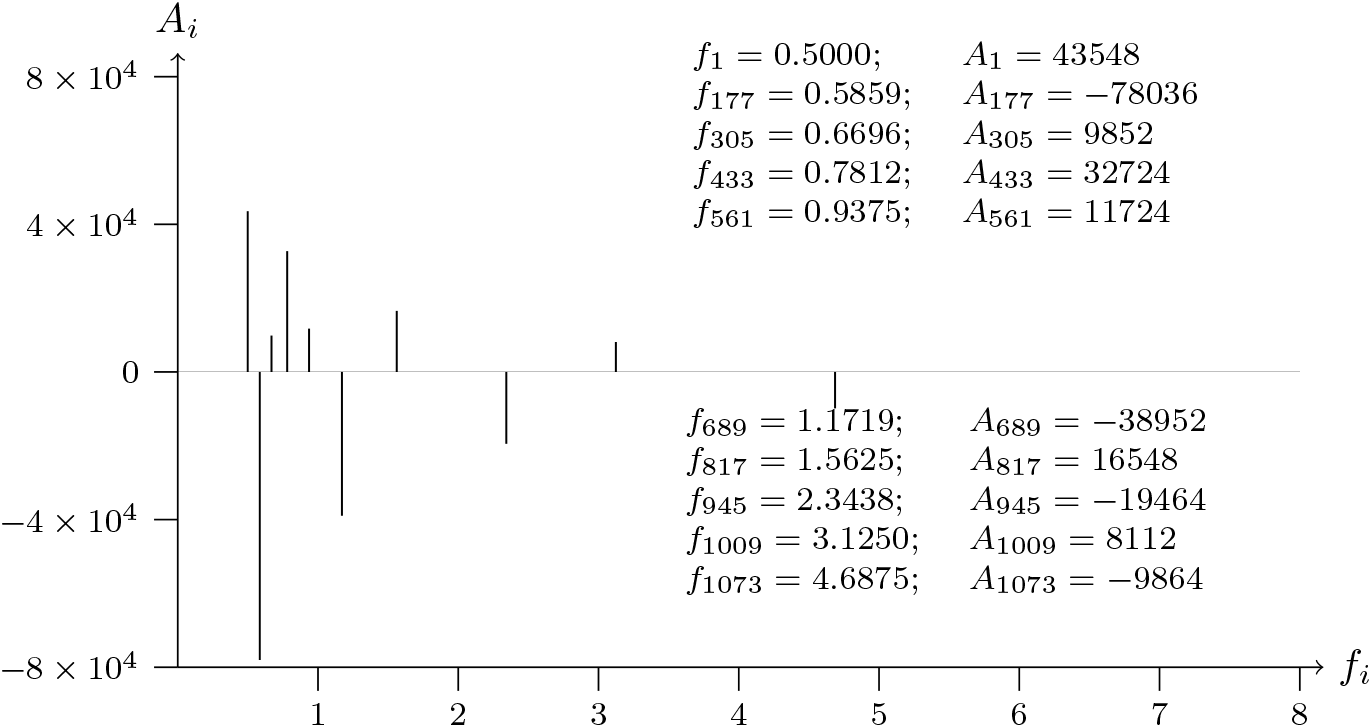
T2T-CHM13v2.0; chromosome 1; ELK4 (oncogene) exon-NM 001973.4-5 analyzed data from 204873778 to 204872578

**Figure 48a:**
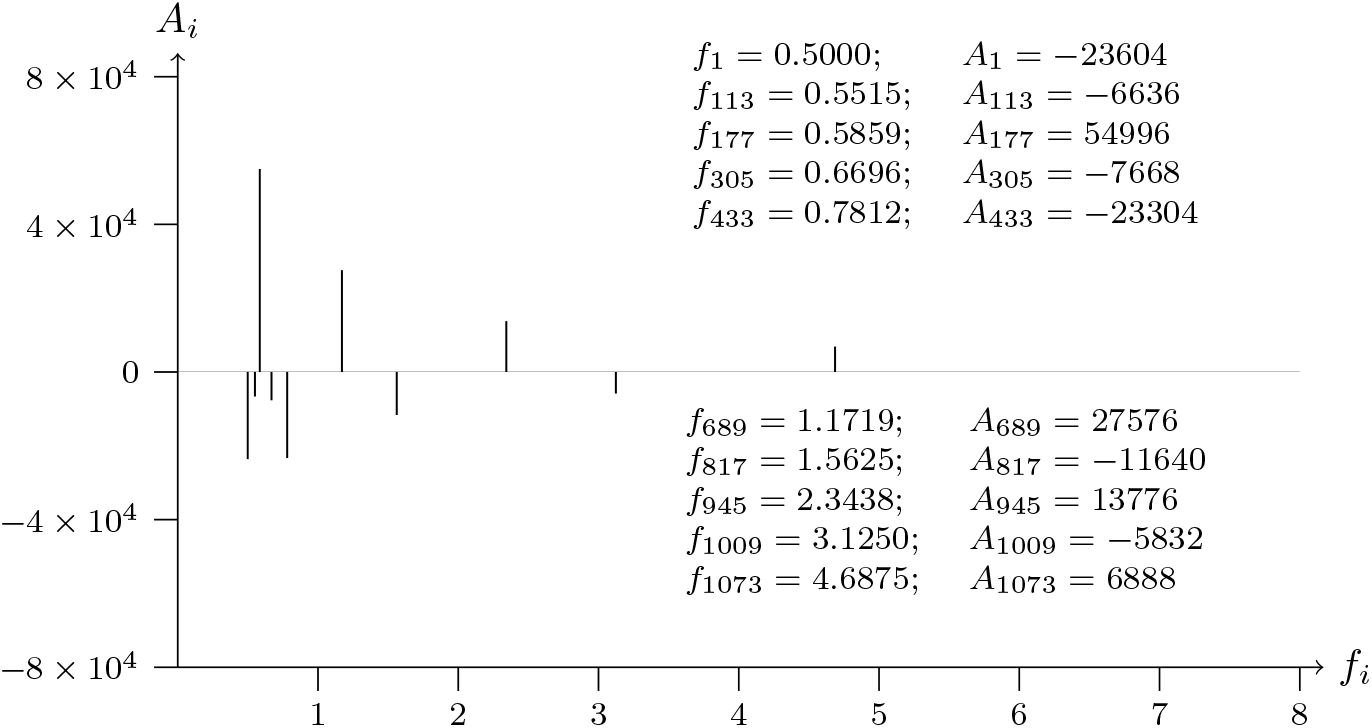
T2T-CHM13v2.0; chromosome 2; REL (oncogene) exon-NM 002908.4-11 analyzed data from 60927604 to 60928804

**Figure 48b:**
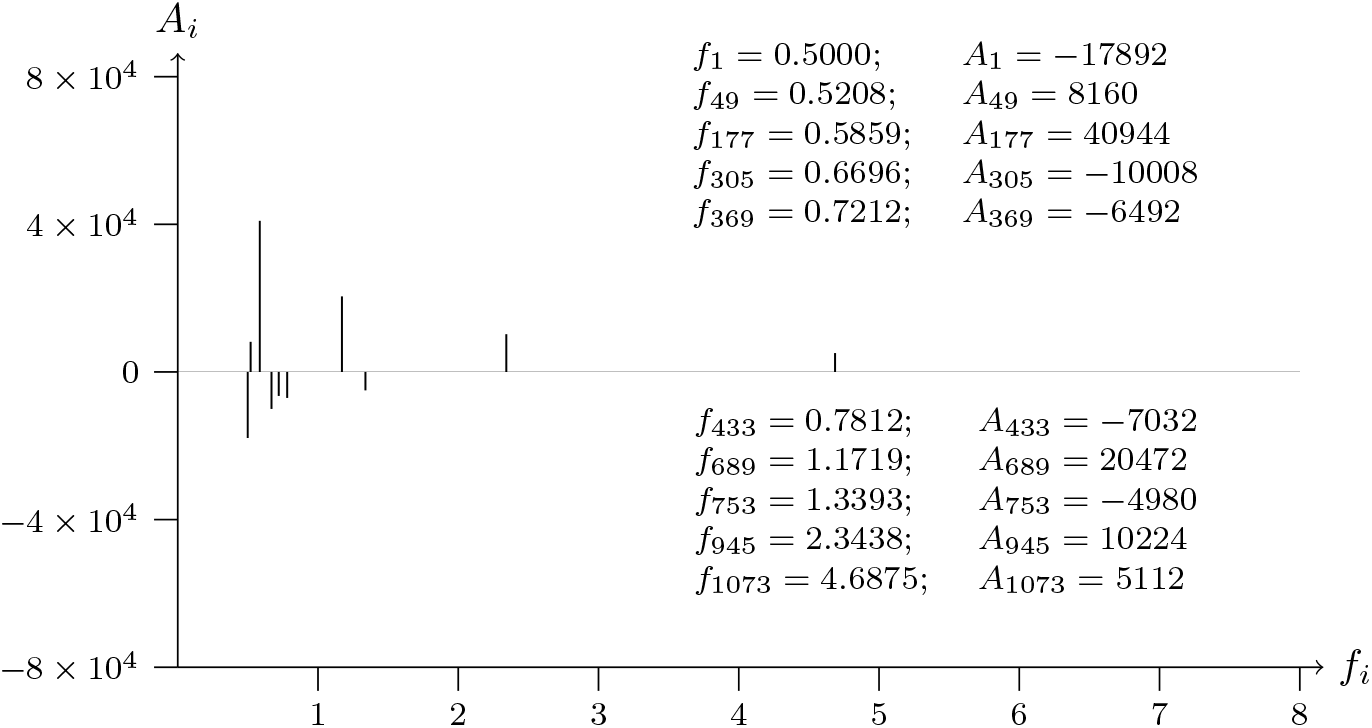
T2T-CHM13v2.0; chromosome 2; REL (oncogene) exon-NM 002908.4-11 analyzed data from 60928804 to 60927604

**Figure 49a:**
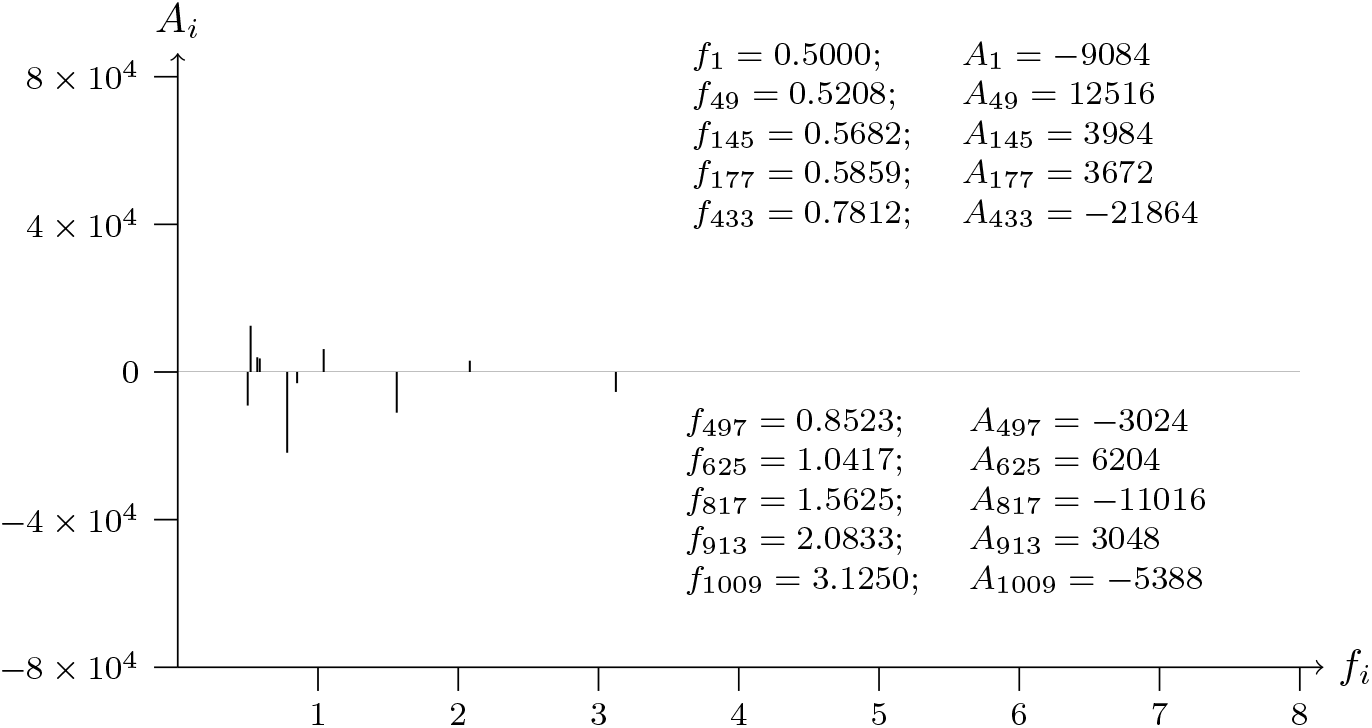
T2T-CHM13v2.0; chromosome 12; VEZT exon-NM 001352092.2-13 analyzed data from 95280935 to 95282135

**Figure 49b:**
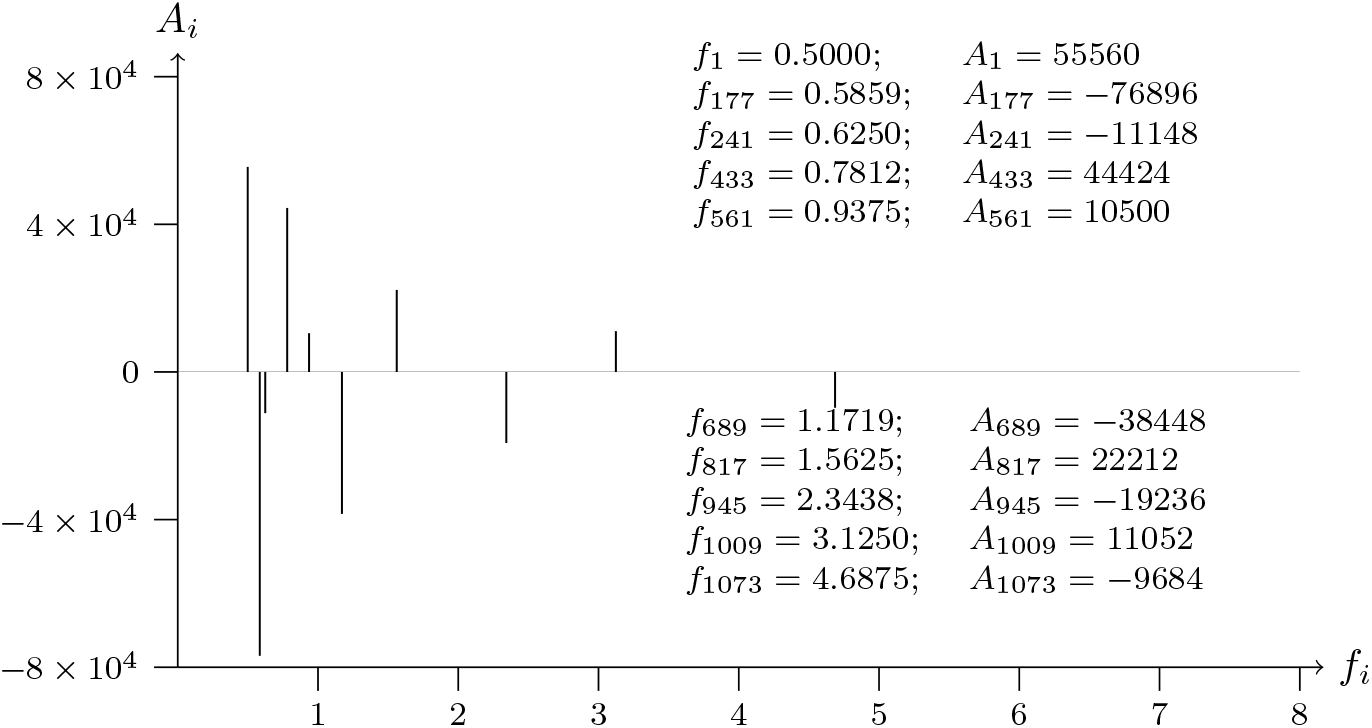
T2T-CHM13v2.0; chromosome 12; VEZT exon-NM 001352092.2-13 analyzed data from 95282135 to 95280935

**Figure 50a:**
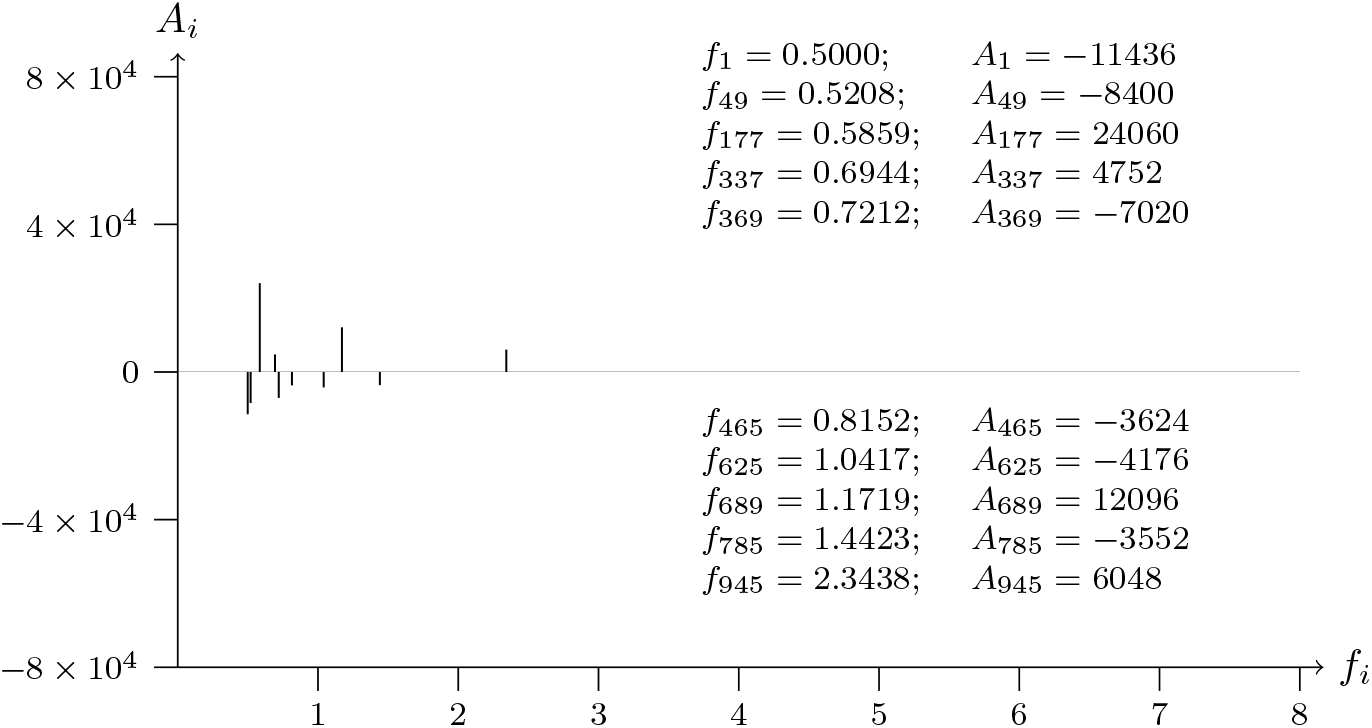
T2T-CHM13v2.0; chromosome 12; VEZT intron-NM 001352092.2-1 analyzed data from 95199140 to 95200340

**Figure 50b:**
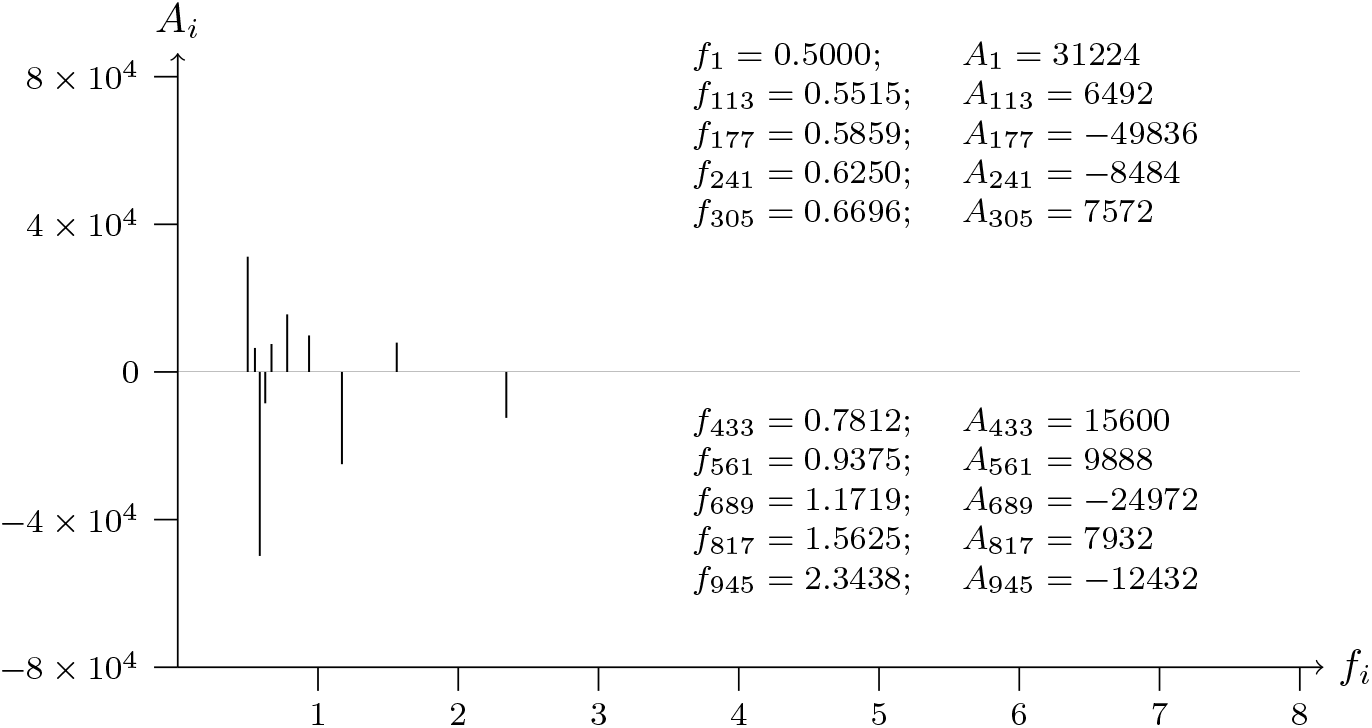
T2T-CHM13v2.0; chromosome 12; VEZT intron-NM 001352092.2-1 analyzed data from 95200340 to 95199140

**Figure 51a:**
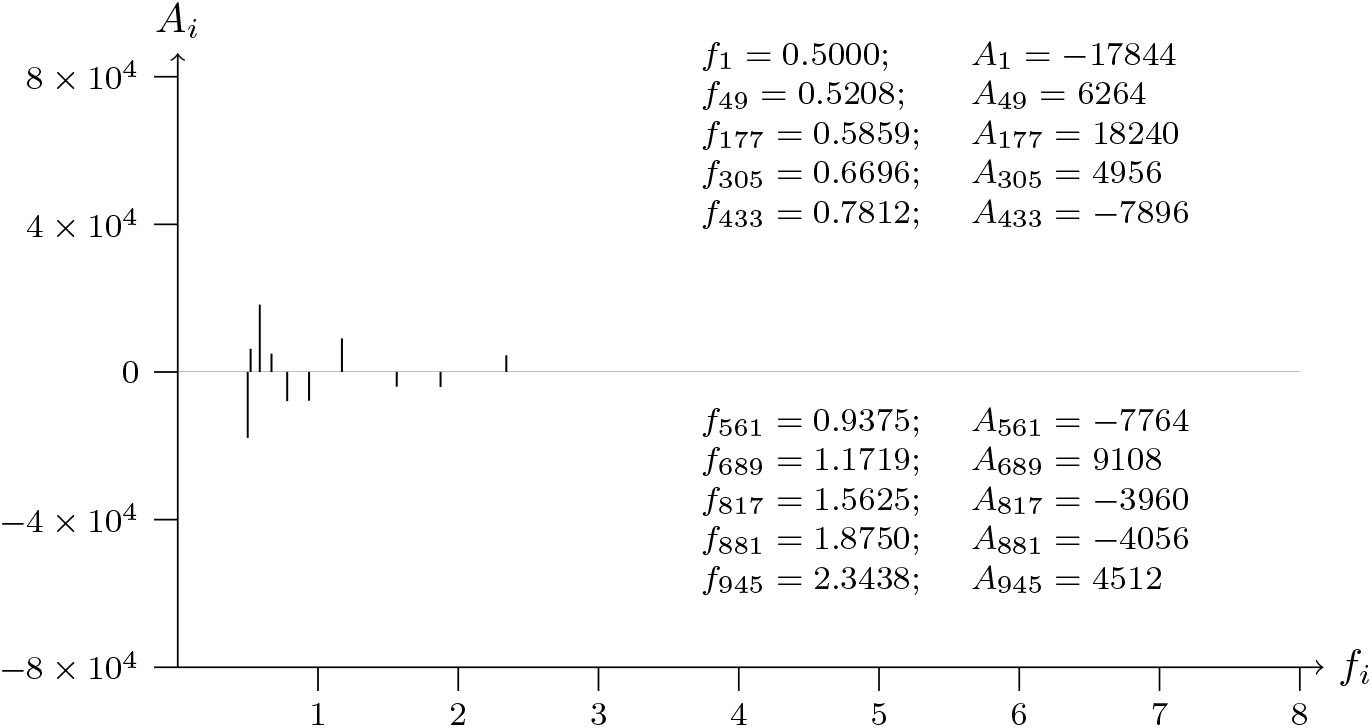
T2T-CHM13v2.0; chromosome 12; VEZT intron-NM 001352092.2-2 analyzed data from 95232839 to 95234039

**Figure 51b:**
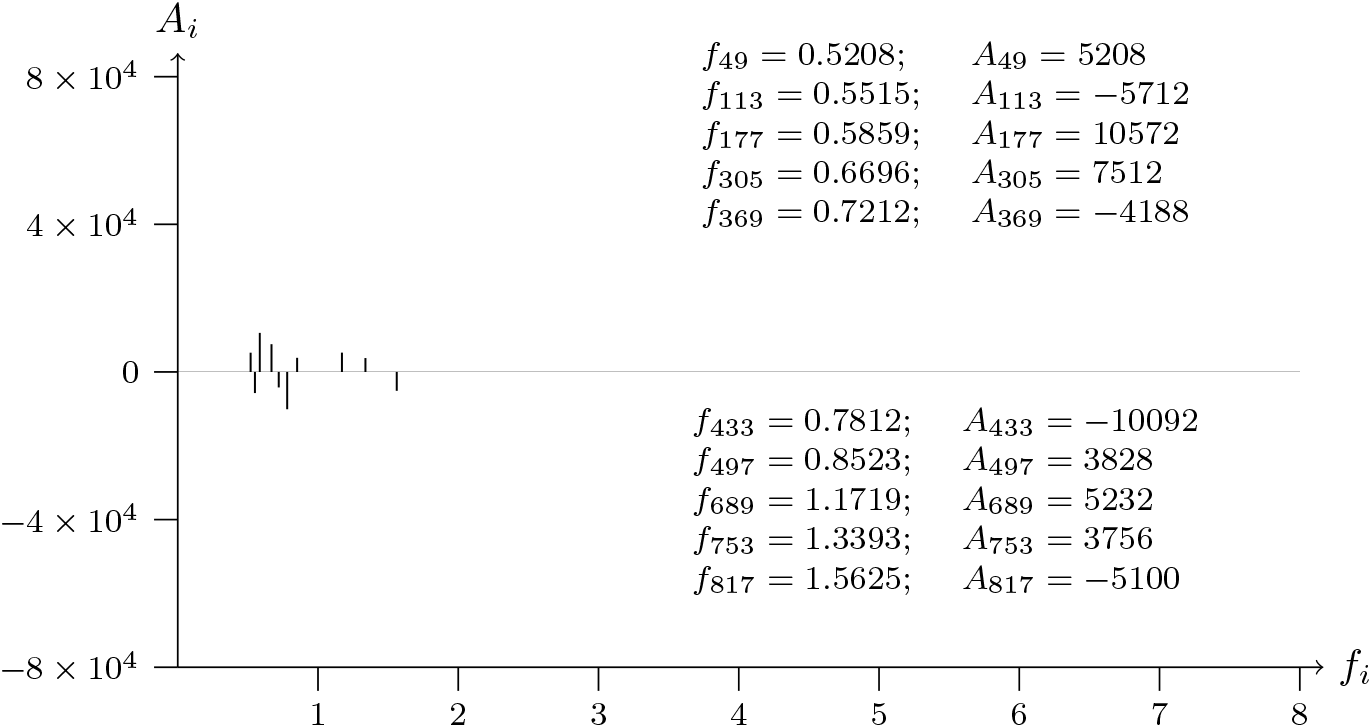
T2T-CHM13v2.0; chromosome 12; VEZT intron-NM 001352092.2-2 analyzed data from 95234039 to 95232839

**Figure 52a:**
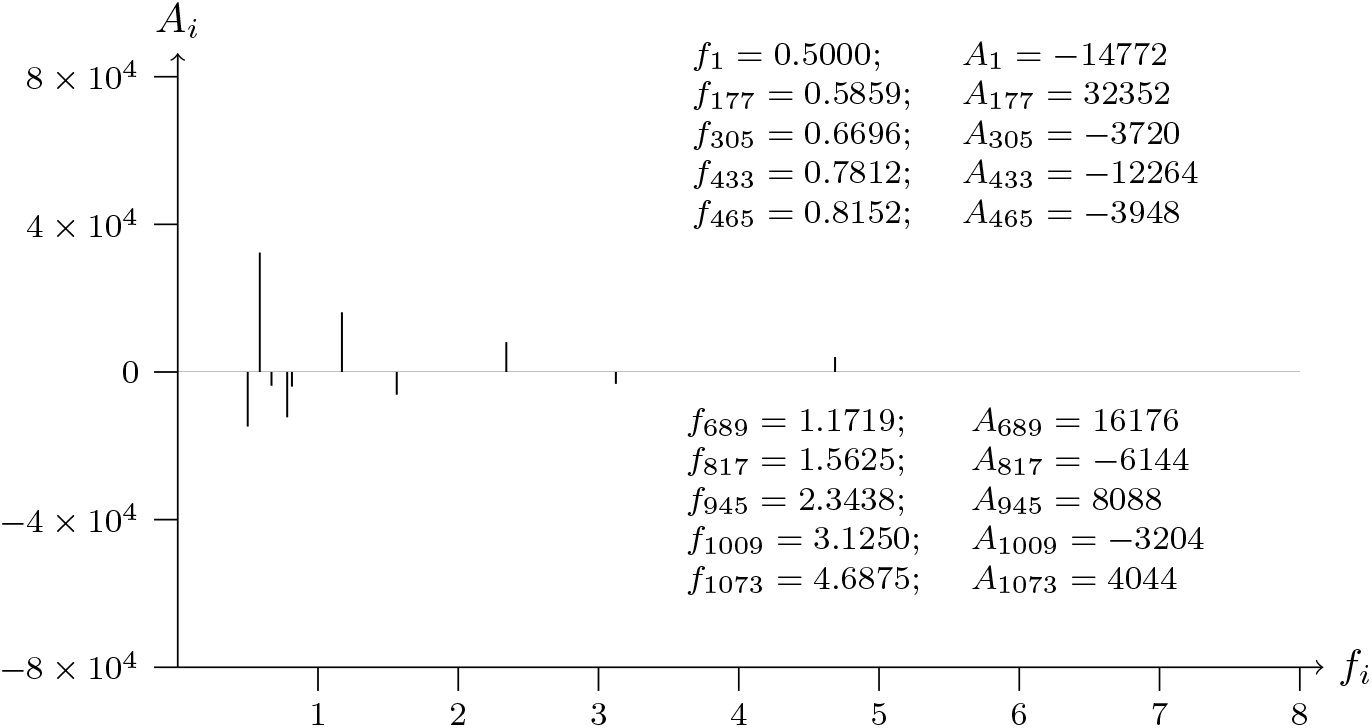
T2T-CHM13v2.0; chromosome 12; VEZT intron-NM 001352092.2-4 analyzed data from 95238006 to 95239206

**Figure 52b:**
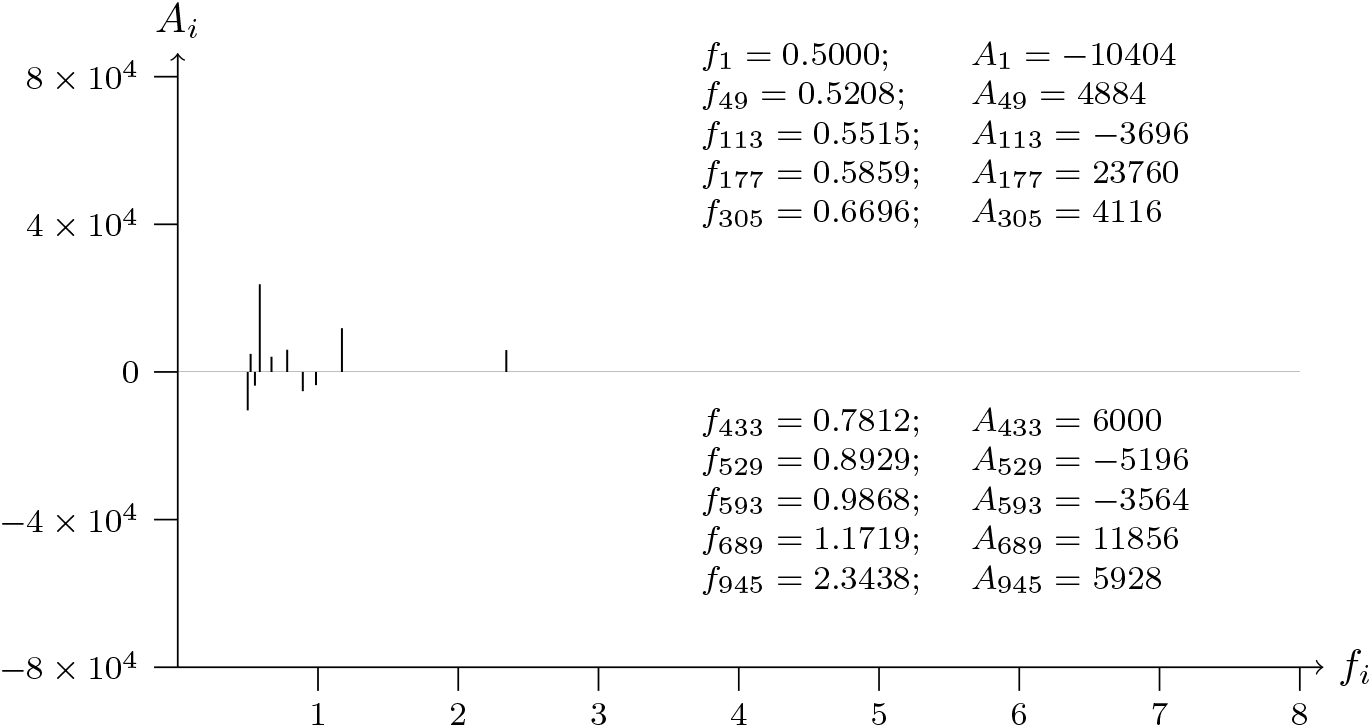
T2T-CHM13v2.0; chromosome 12; VEZT intron-NM 001352092.2-4 analyzed data from 95239206 to 95238006

**Figure 53a:**
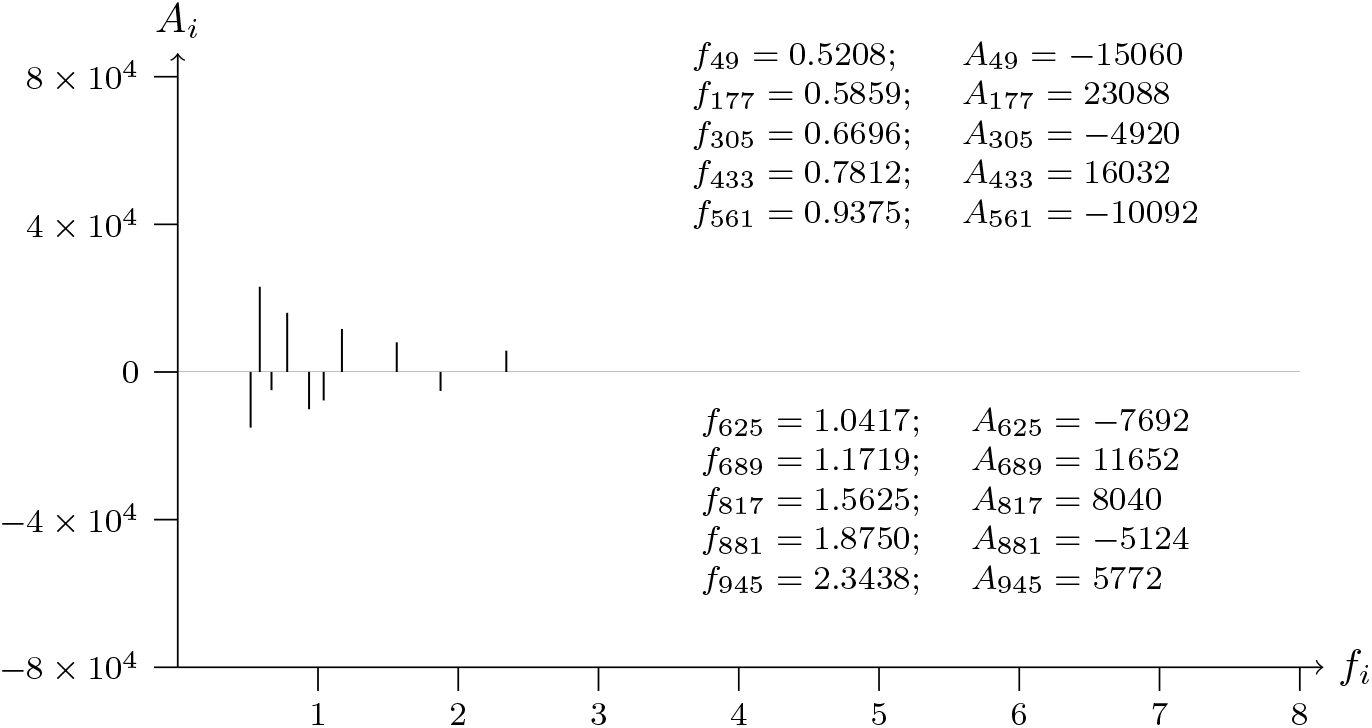
T2T-CHM13v2.0; chromosome 12; VEZT intron-NM 001352092.2-5 analyzed data from 95243850 to 95245050

**Figure 53b:**
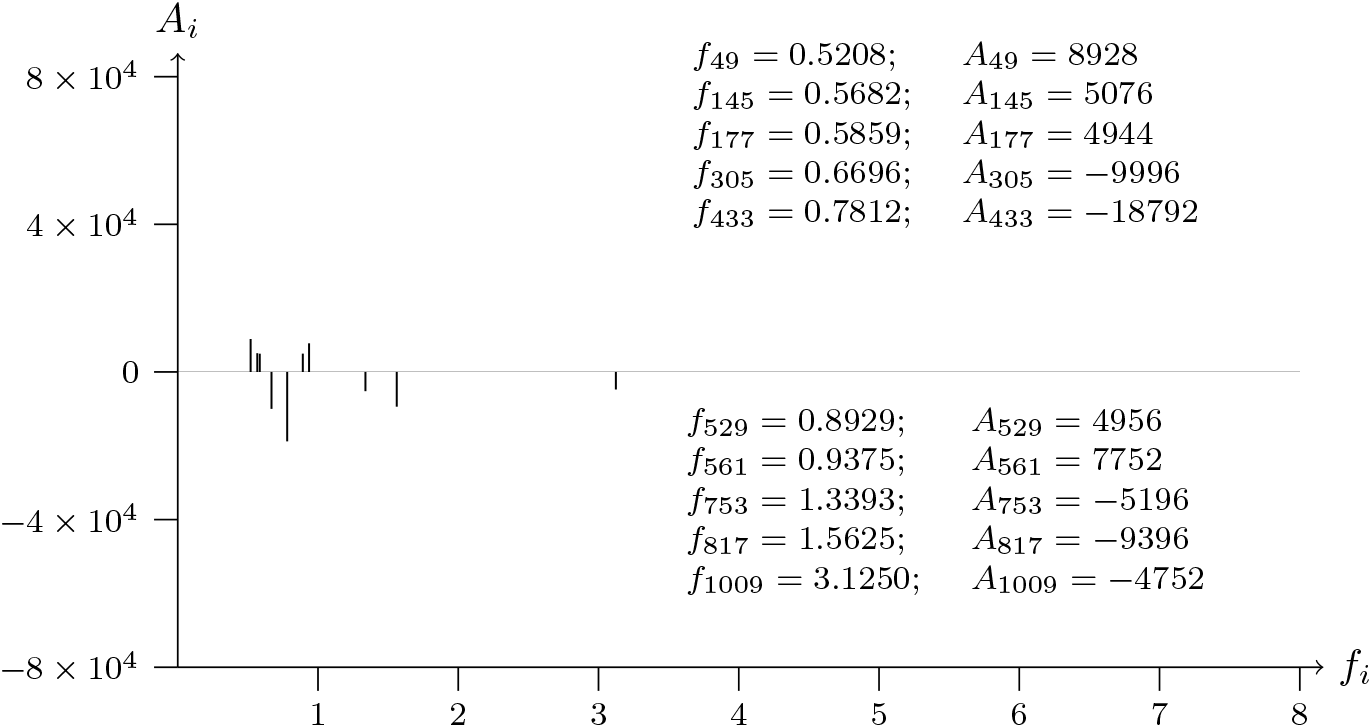
T2T-CHM13v2.0; chromosome 12; VEZT intron-NM 001352092.2-5 analyzed data from 95245050 to 95243850

**Figure 54a:**
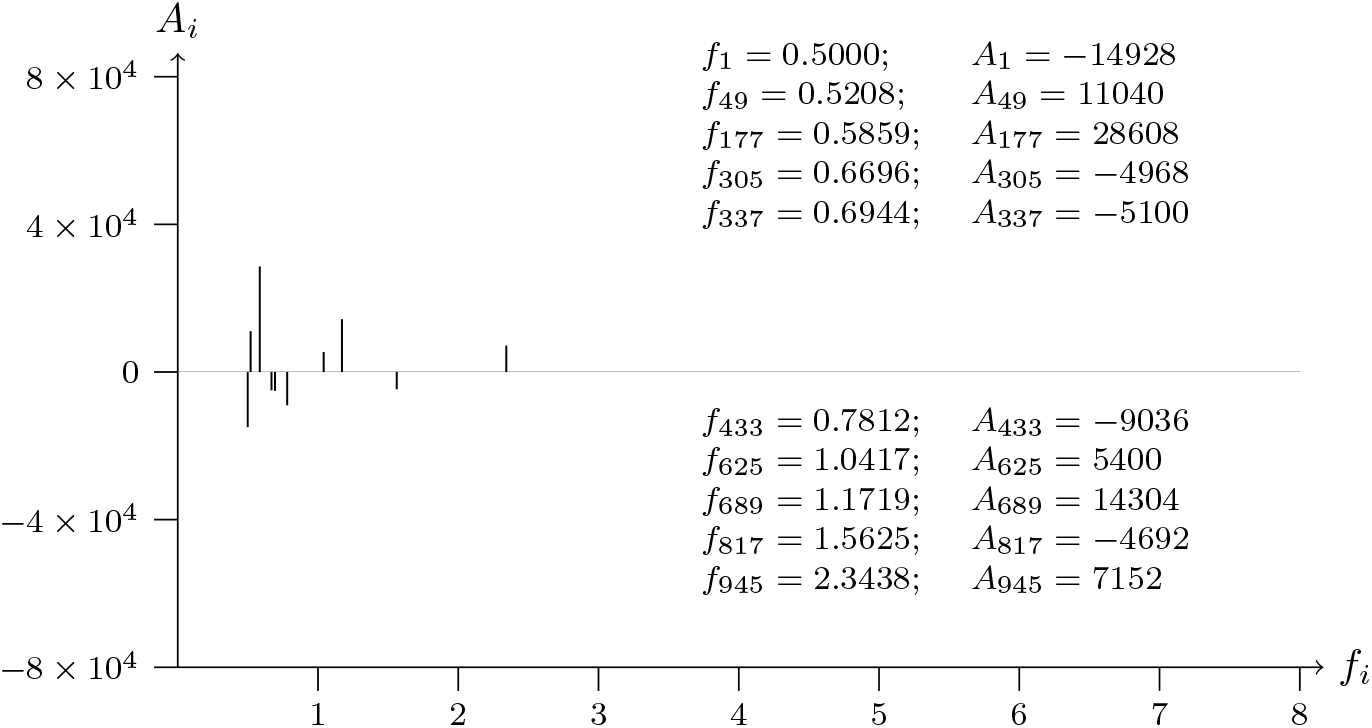
T2T-CHM13v2.0; chromosome 12; VEZT intron-NM 001352092.2-6 analyzed data from 95247401 to 95248601

**Figure 54b:**
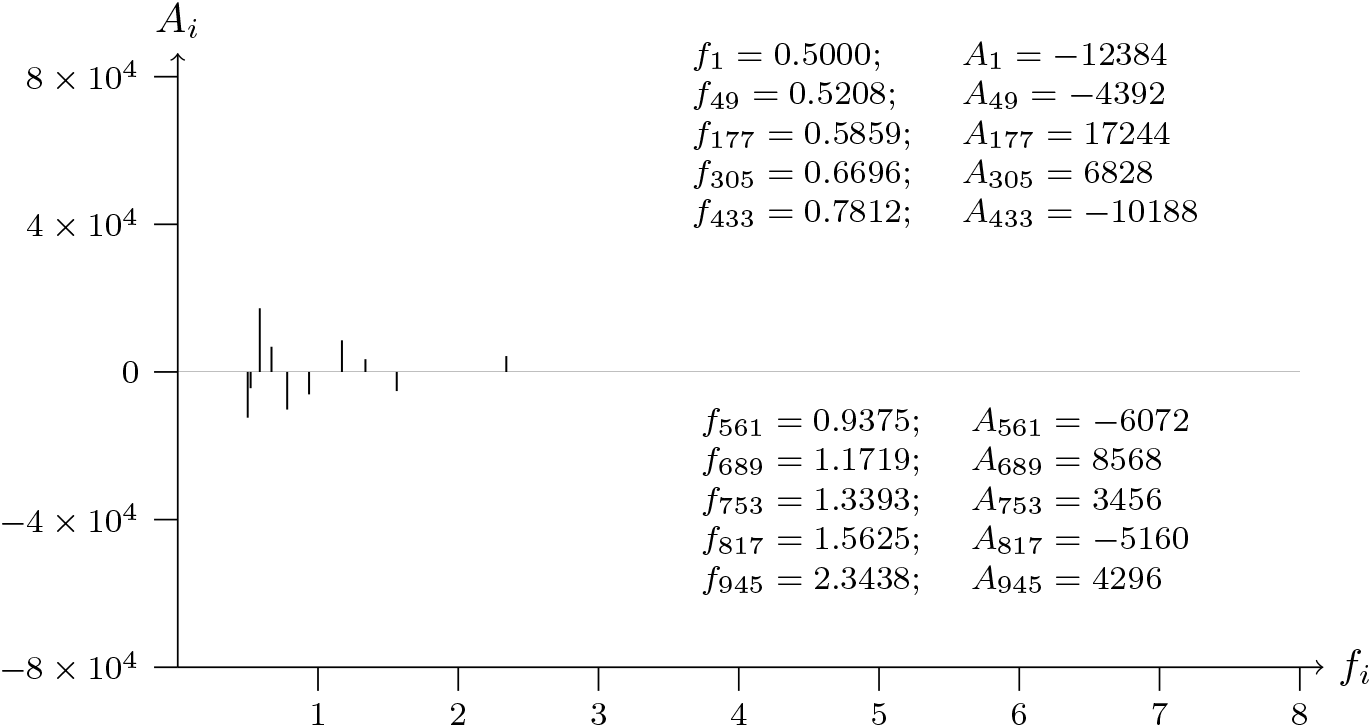
T2T-CHM13v2.0; chromosome 12; VEZT intron-NM 001352092.2-6 analyzed data from 95248601 to 95247401

**Figure 55a:**
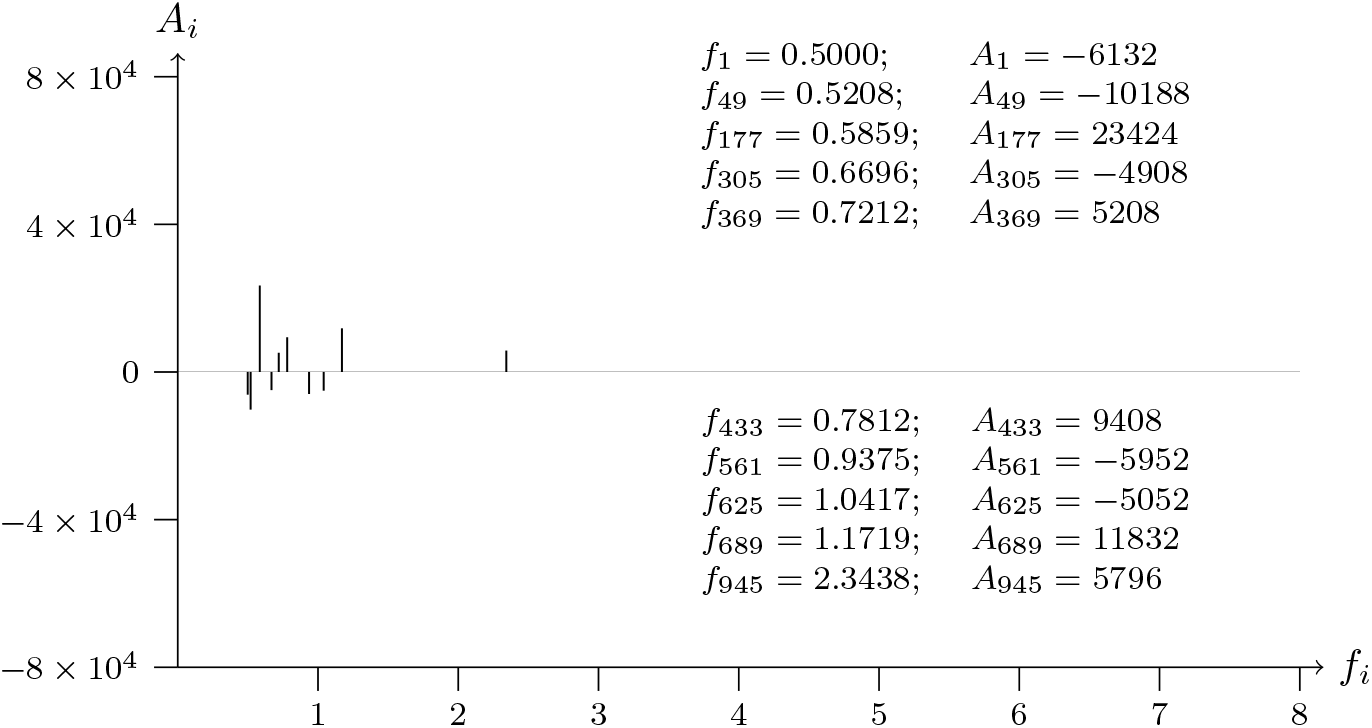
T2T-CHM13v2.0; chromosome 12; VEZT intron-NM 001352092.2-7 analyzed data from 95250957 to 952521571

**Figure 55b:**
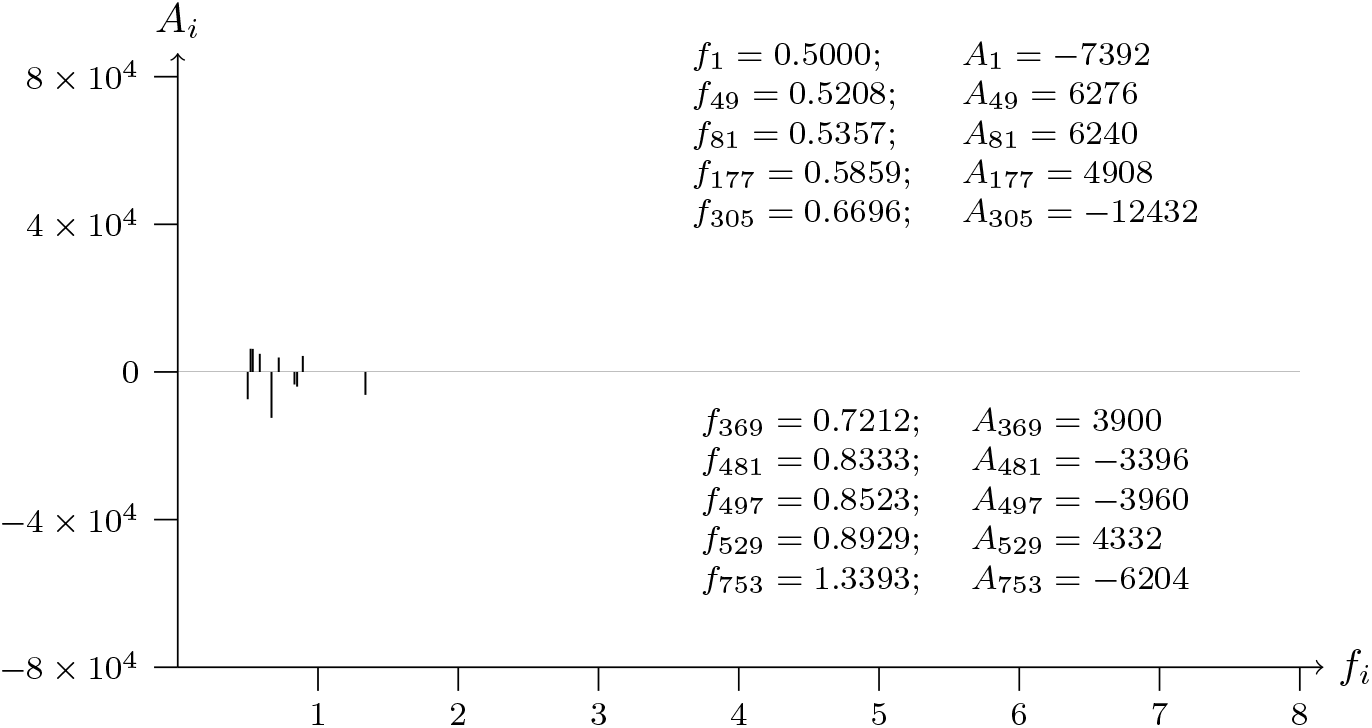
T2T-CHM13v2.0; chromosome 12; VEZT intron-NM 001352092.2-7 analyzed data from 95252157 to 95250957

**Figure 56a:**
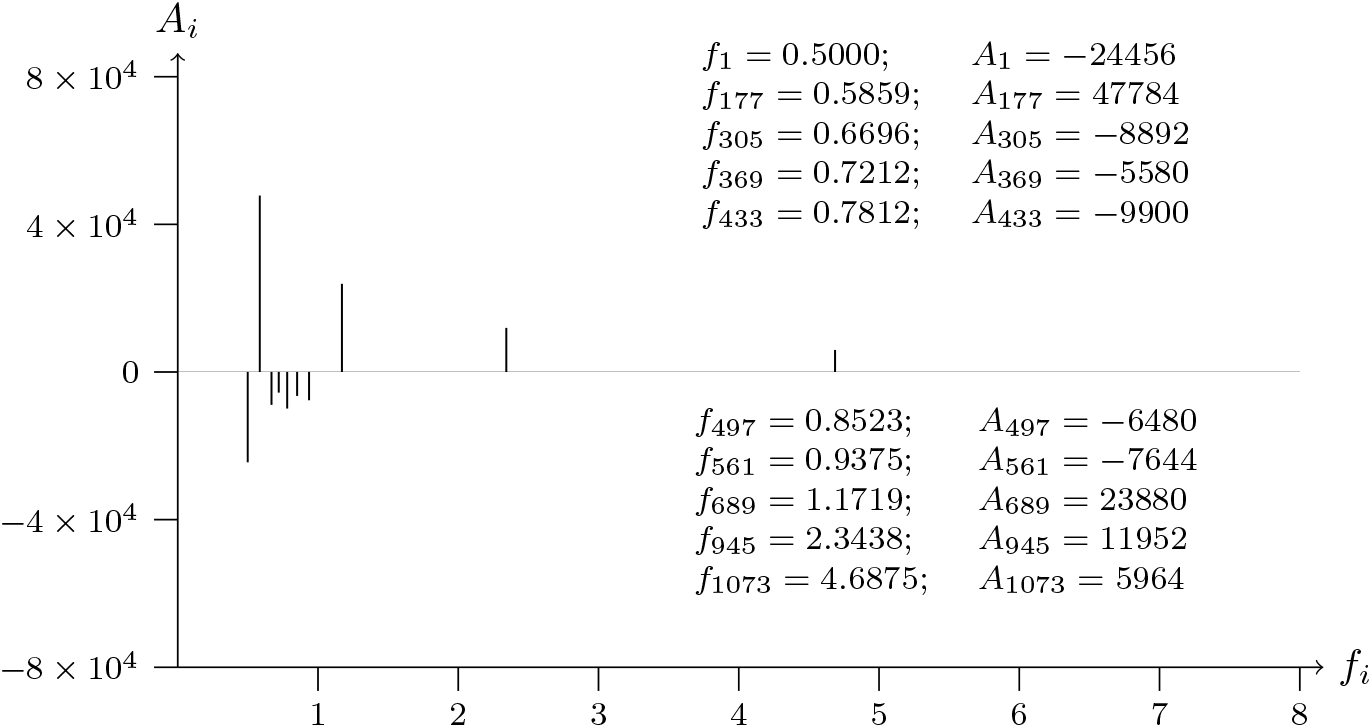
T2T-CHM13v2.0; chromosome 12; VEZT intron-NM 001352092.2-8 analyzed data from 95255658 to 95256858

**Figure 56b:**
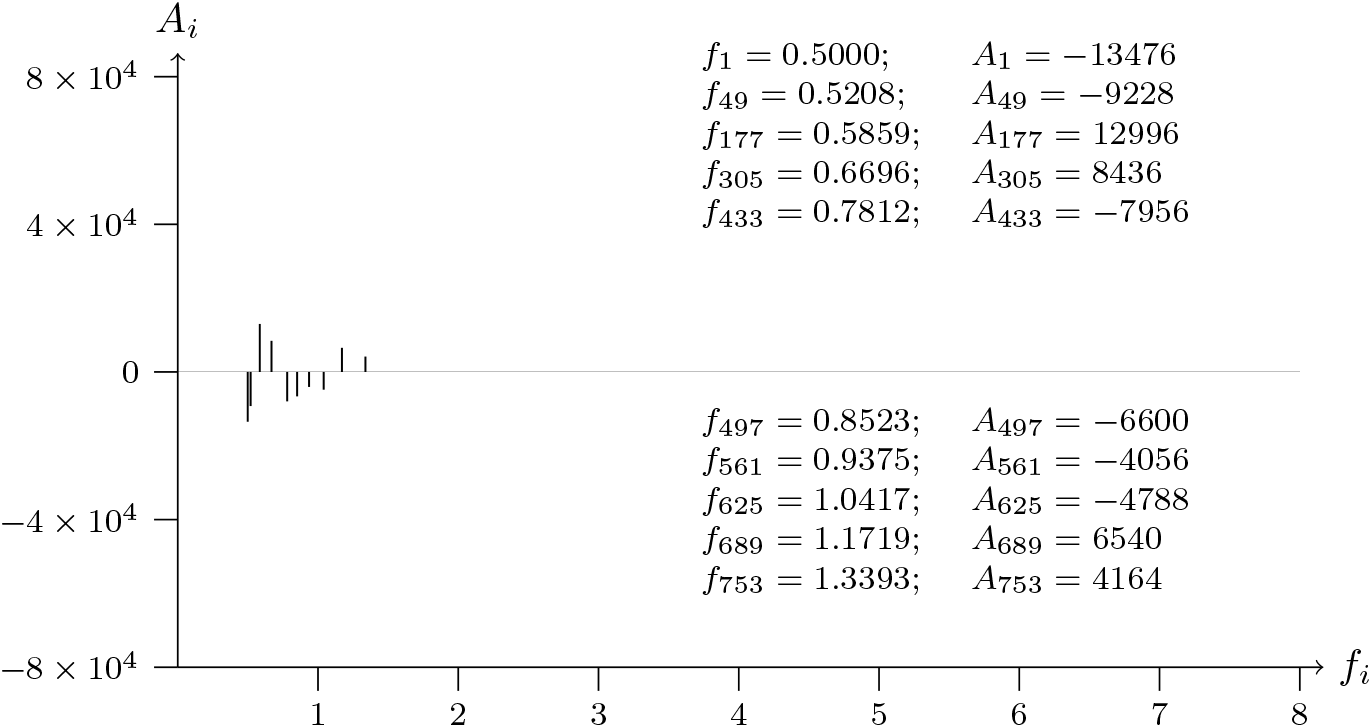
T2T-CHM13v2.0; chromosome 12; VEZT intron-NM 001352092.2-8 analyzed data from 95256858 to 95255658

**Figure 57a:**
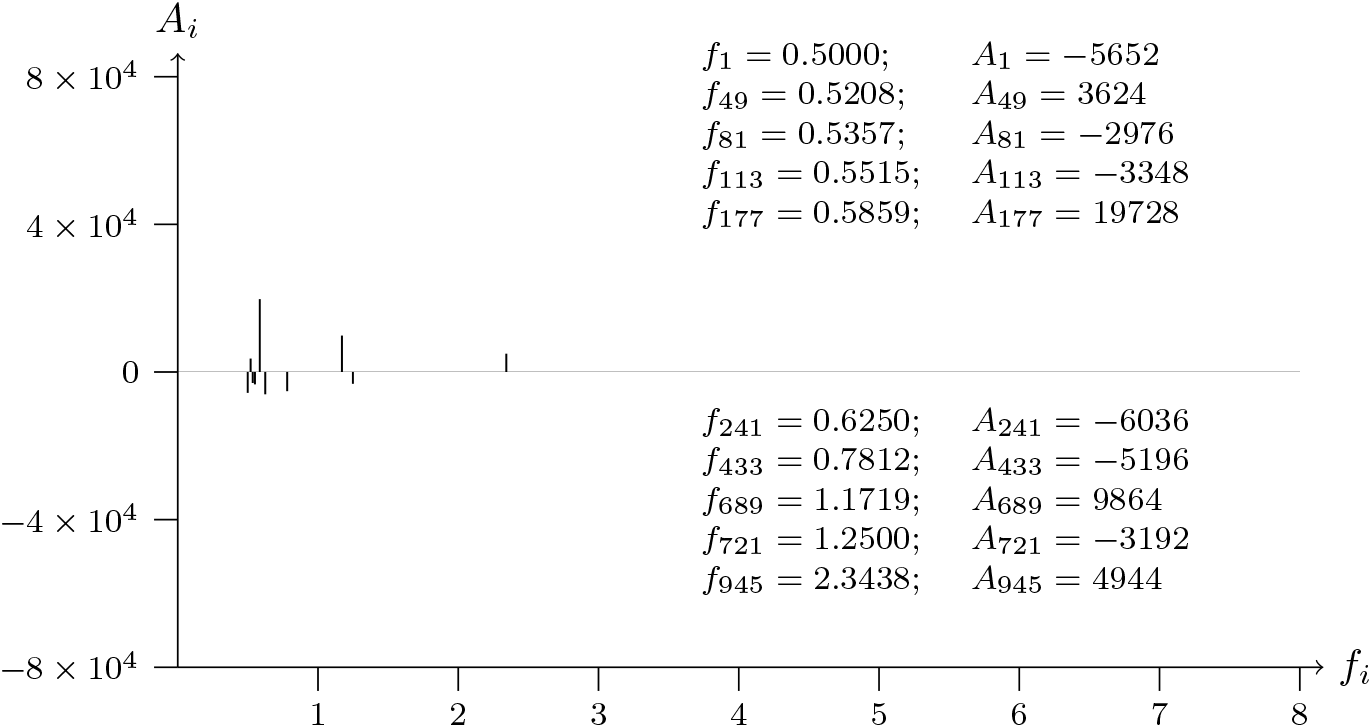
T2T-CHM13v2.0; chromosome 12; VEZT intron-NM 001352092.2-9 analyzed data from 95263413 to 95264613

**Figure 57b:**
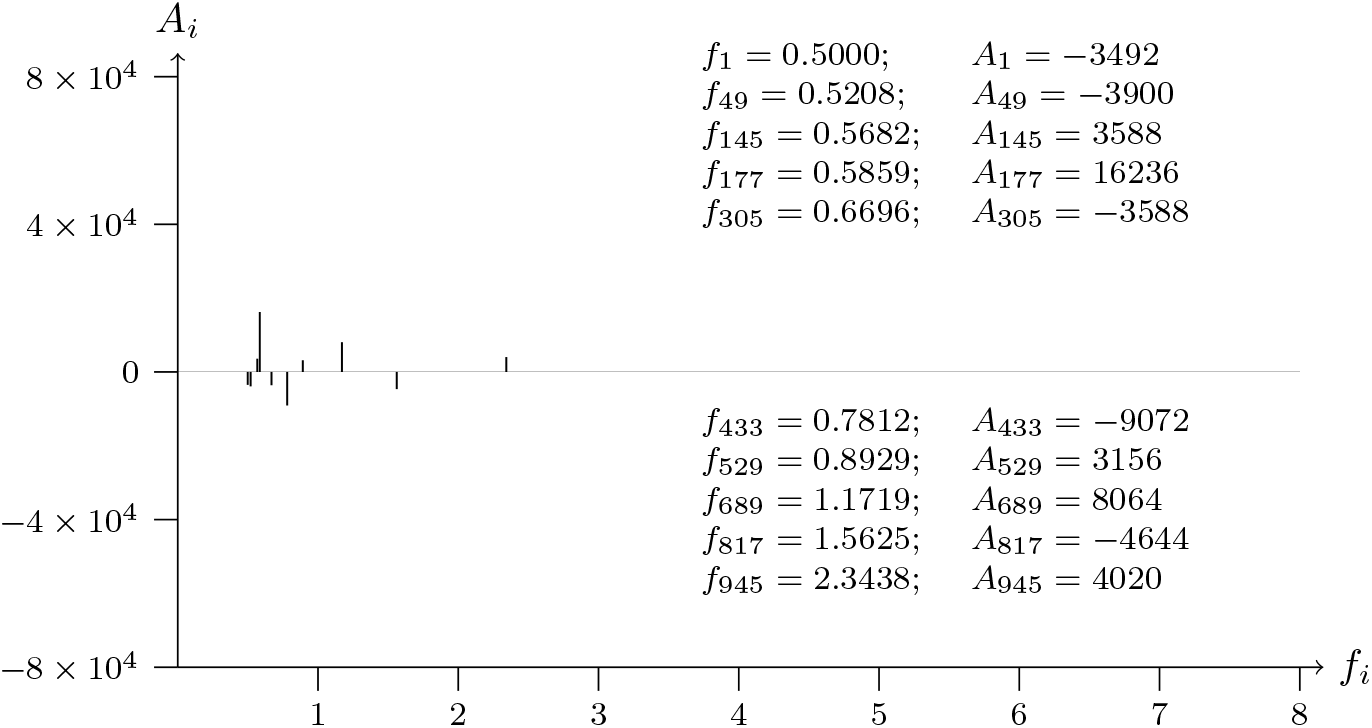
T2T-CHM13v2.0; chromosome 12; VEZT intron-NM 001352092.2-9 analyzed data from 95264613 to 95263413

**Figure 58a:**
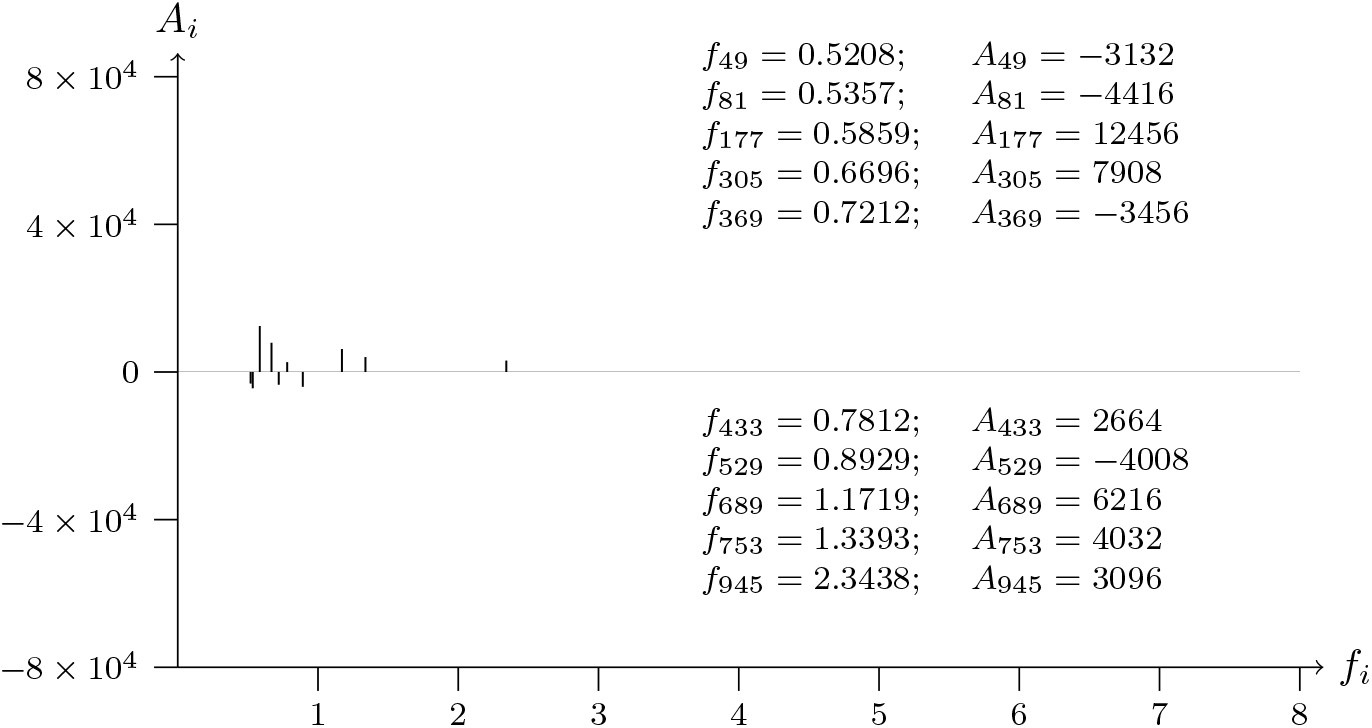
T2T-CHM13v2.0; chromosome 12; VEZT intron-NM 001352092.2-10 analyzed data from 95268628 to 95269828

**Figure 58b:**
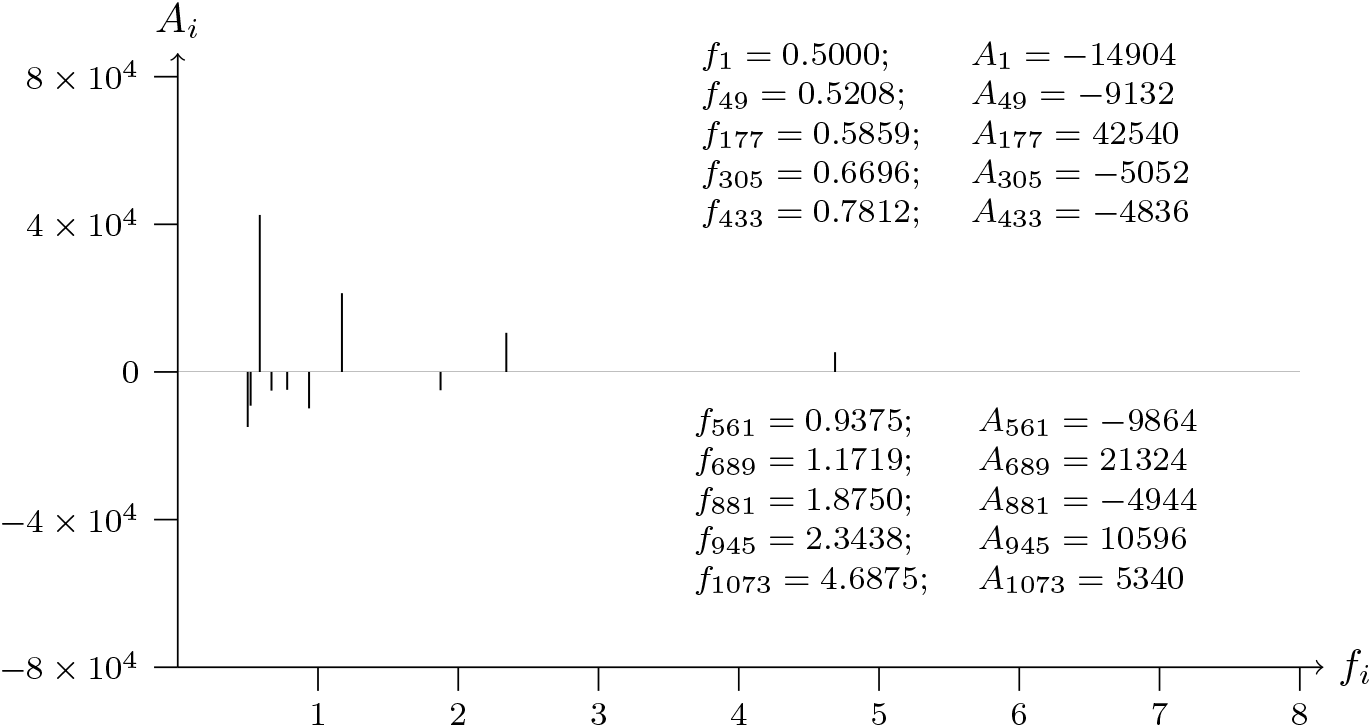
T2T-CHM13v2.0; chromosome 12; VEZT intron-NM 001352092.2-10 analyzed data from 95269828 to 95268628

**Figure 59a:**
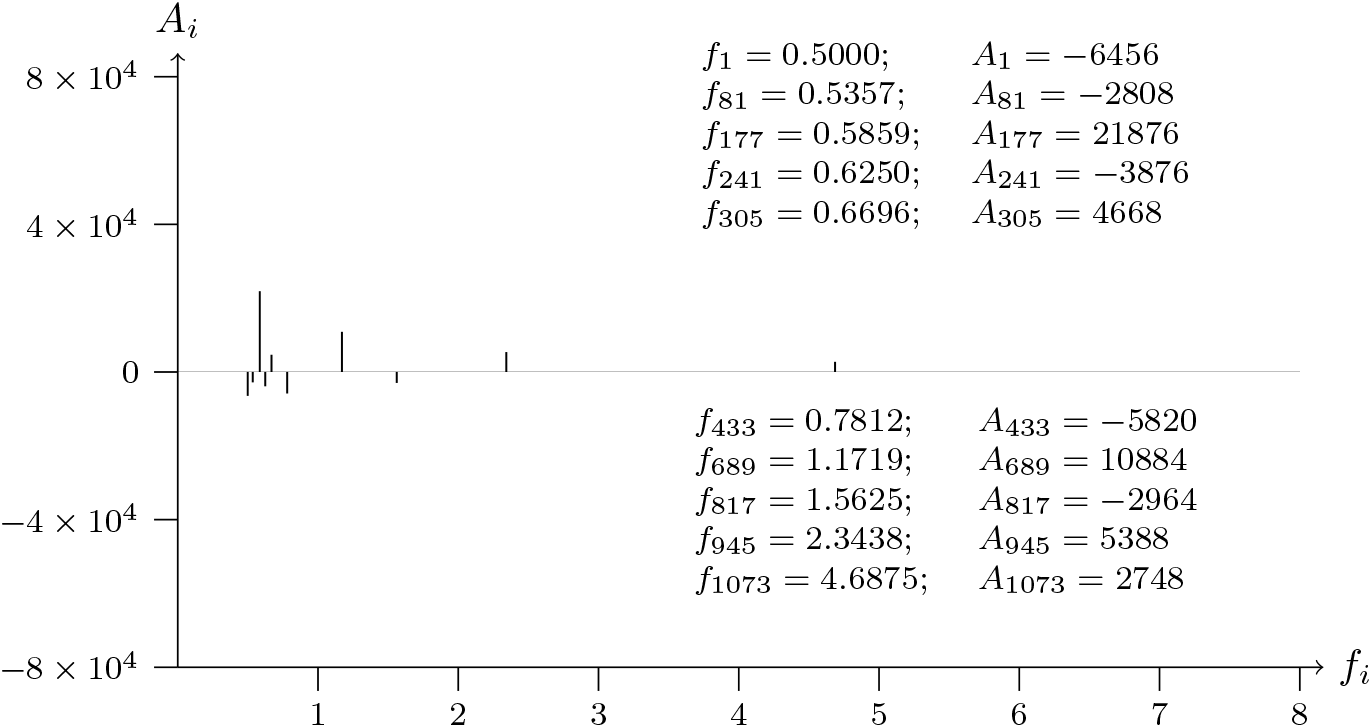
T2T-CHM13v2.0; chromosome 12; VEZT intron-NM 001352092.2-11 analyzed data from 95275142 to 95276342

**Figure 59b:**
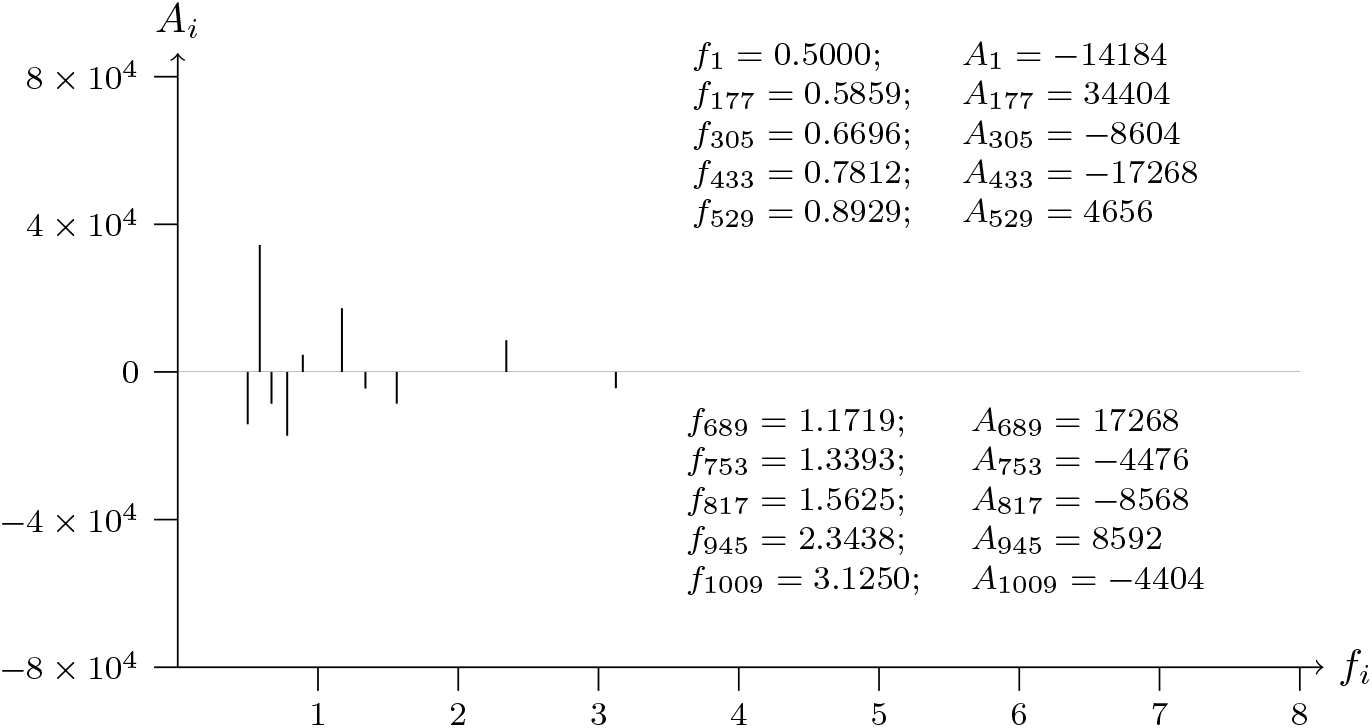
T2T-CHM13v2.0; chromosome 12; VEZT intron-NM 001352092.2-11 analyzed data from 95276342 to 95275142

**Figure 60a:**
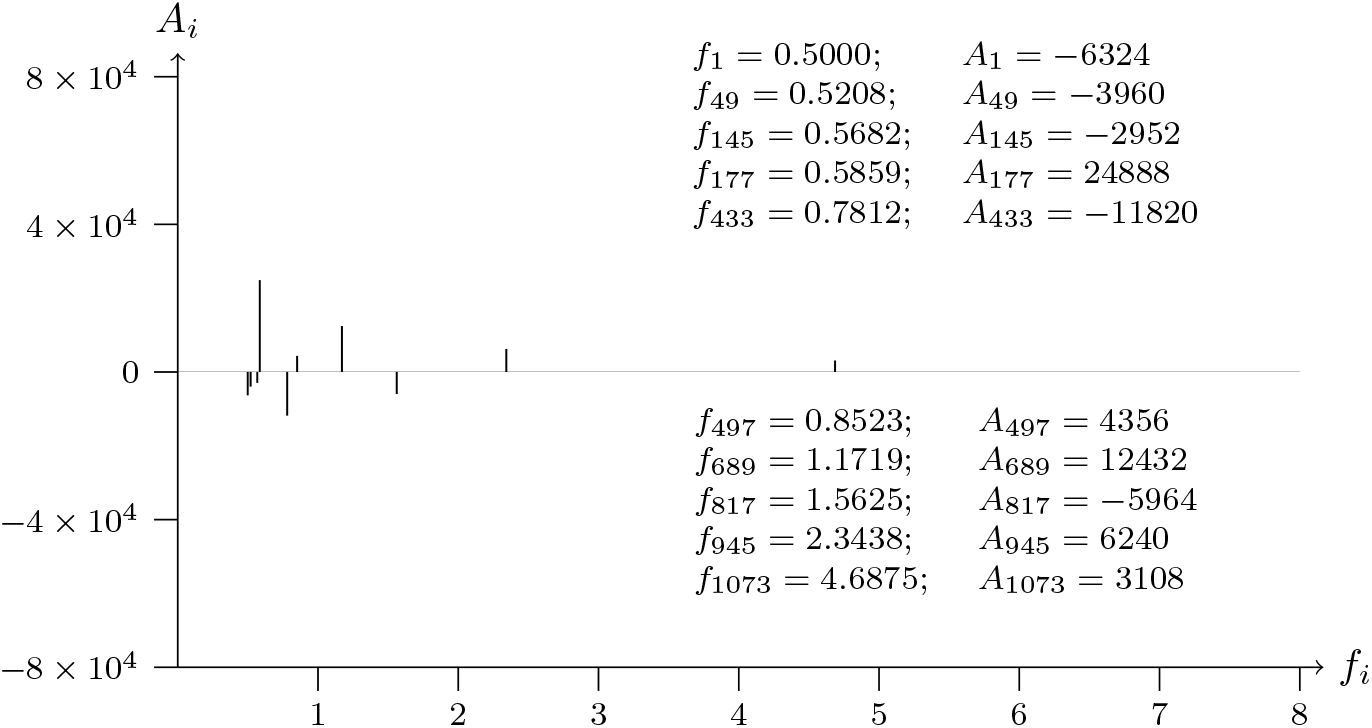
T2T-CHM13v2.0; chromosome 12; VEZT intron-NM 001352092.2-12 analyzed data from 95277028 to 95278228

**Figure 60b:**
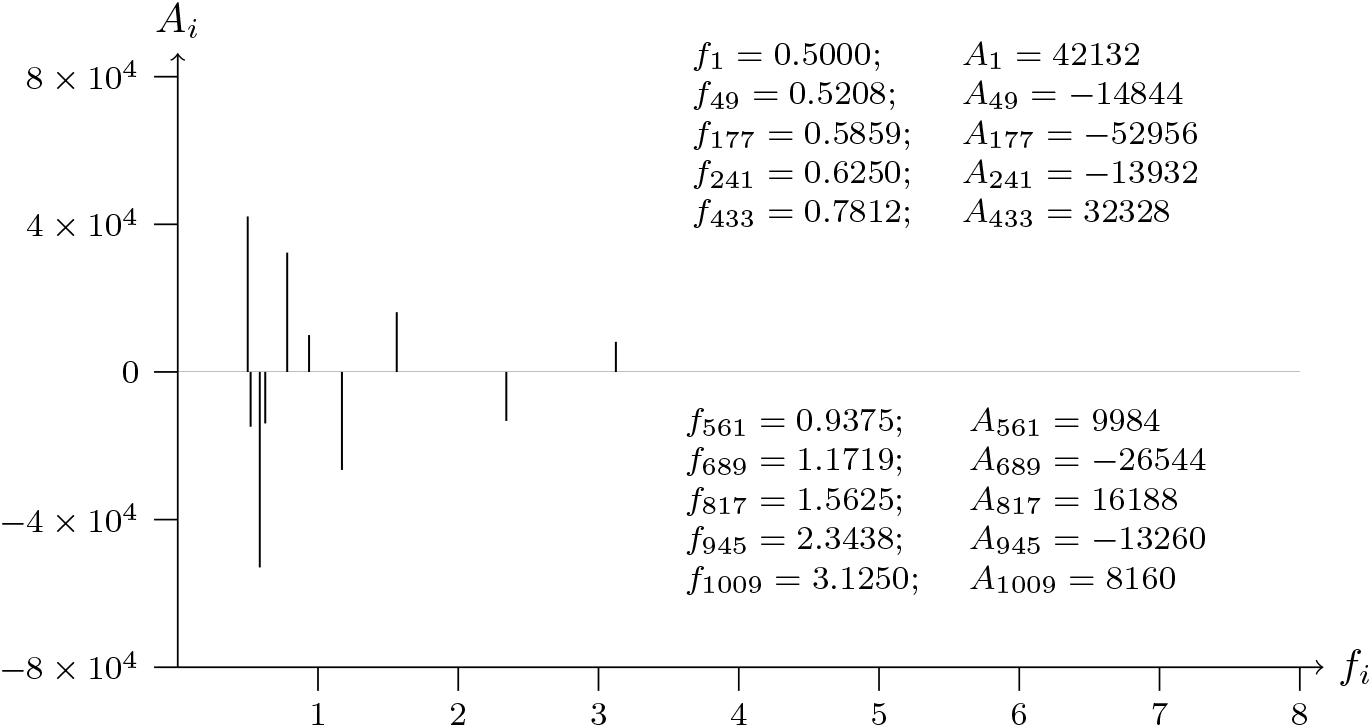
T2T-CHM13v2.0; chromosome 12; VEZT intron-NM 001352092.2-

**Figure 61a:**
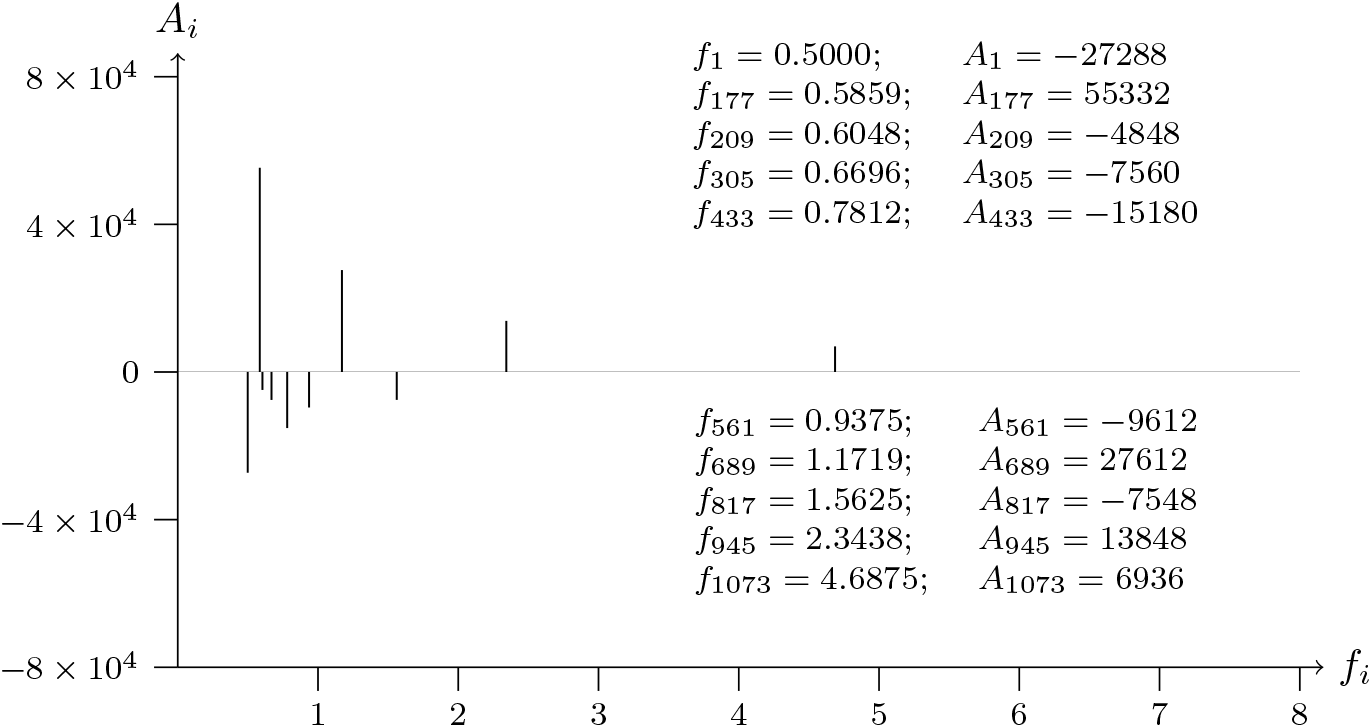
T2T-CHM13v2.0; chromosome 12; TAOK3 exon-NM 016281.4-21 analyzed data from 118137047 to 118138247

**Figure 61b:**
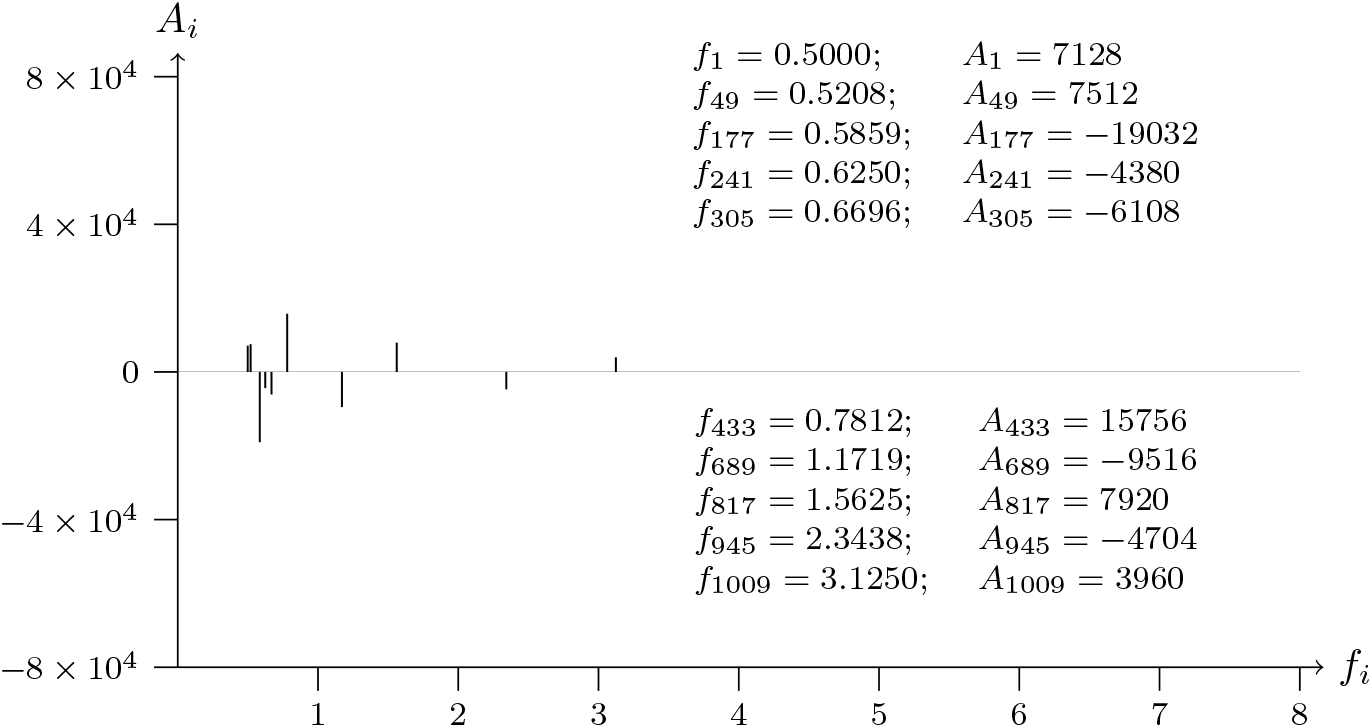
T2T-CHM13v2.0; chromosome 12; TAOK3 exon-NM 016281.4-21

**Figure 62a:**
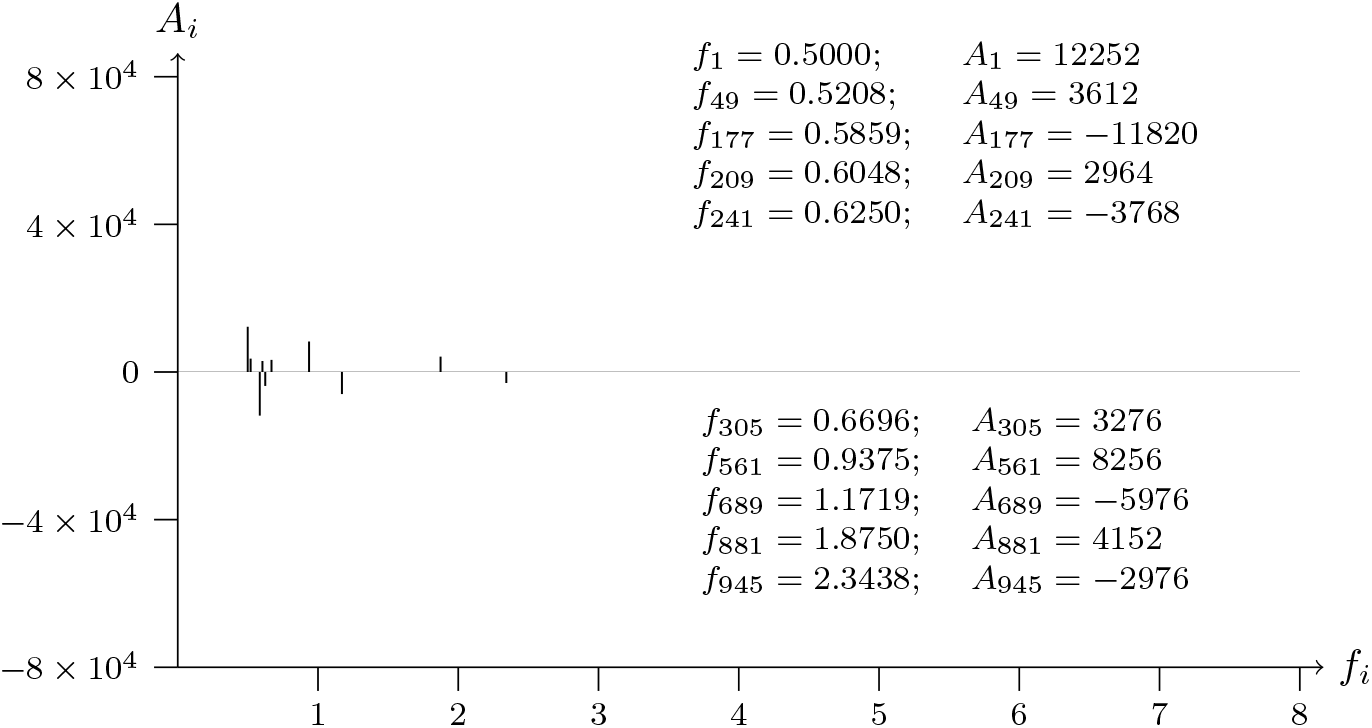
T2T-CHM13v2.0; chromosome 12; SMUG1 exon-NM 001351259.2-2 analyzed data from 54145927 to 54147127

**Figure 62b:**
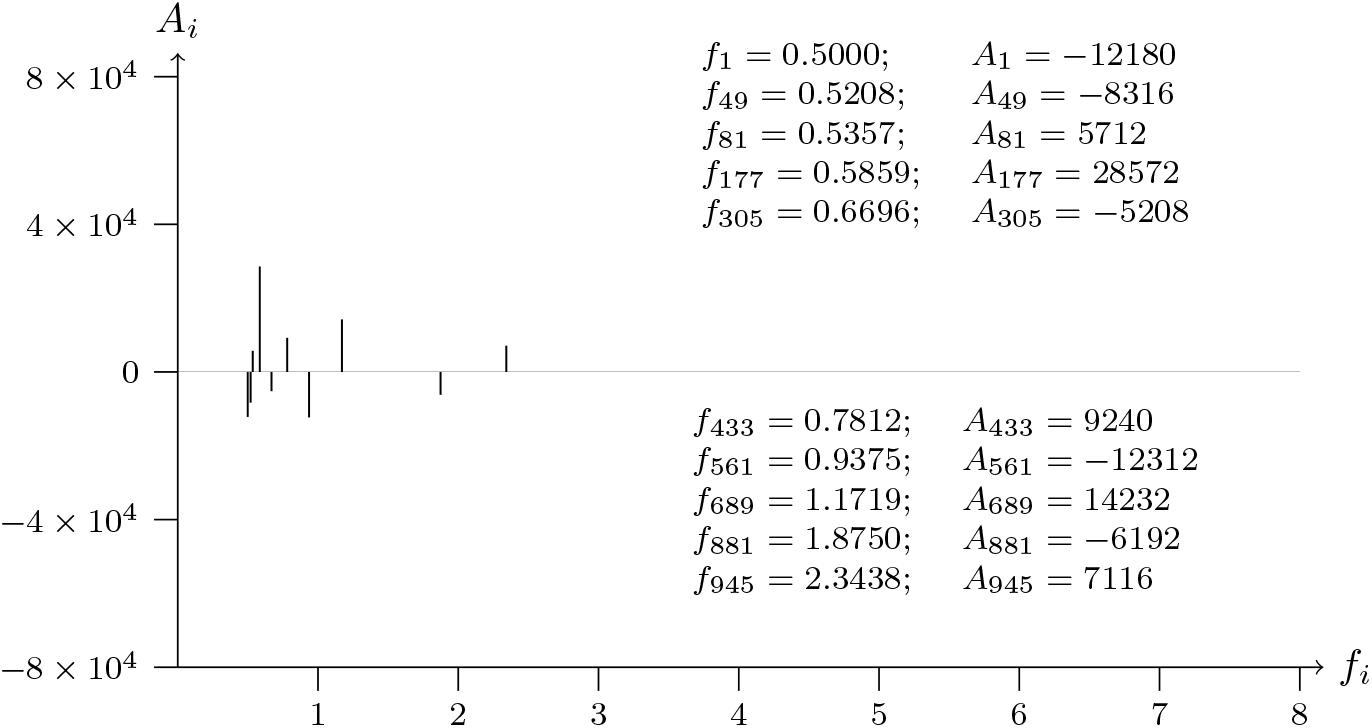
T2T-CHM13v2.0; chromosome 12; SMUG1 exon-NM 001351259.2-

**Figure 63a:**
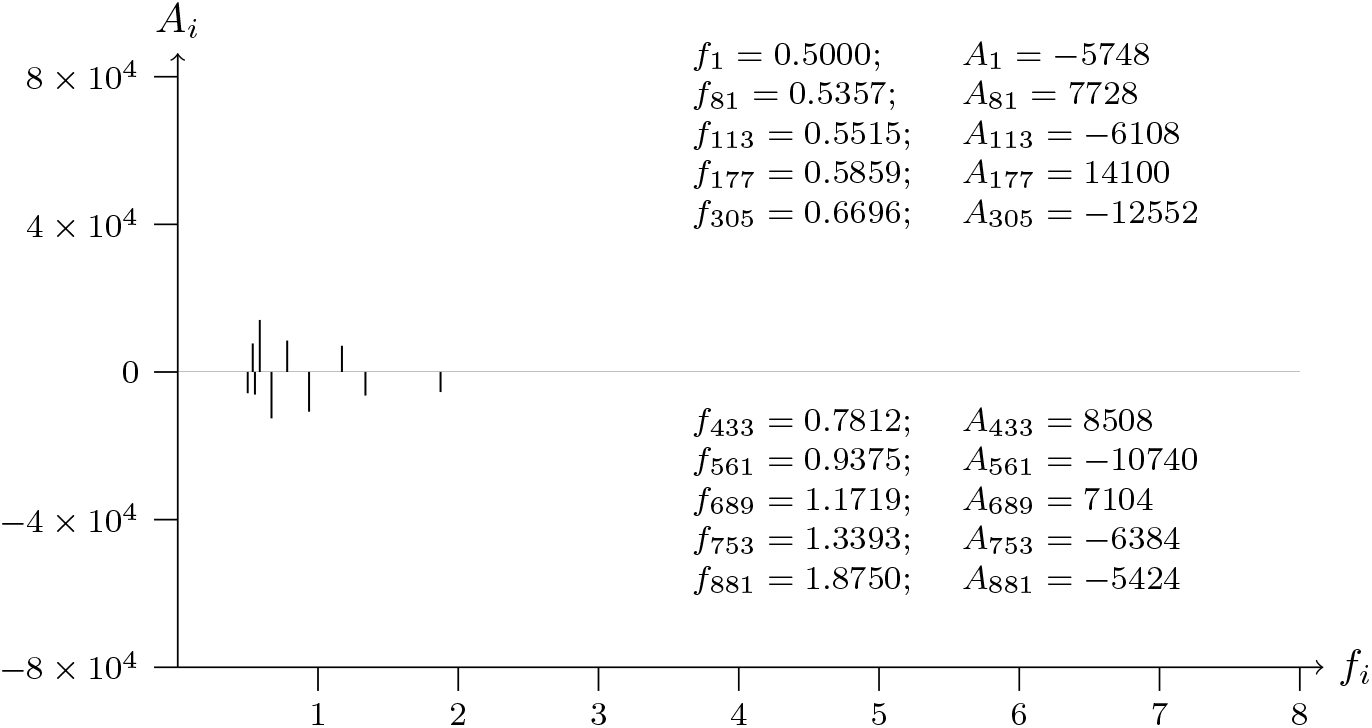
T2T-CHM13v2.0; chromosome 4; MXD4 exon-NM 006454.3-6 analyzed data from 2246204 to 2247404

**Figure 63b:**
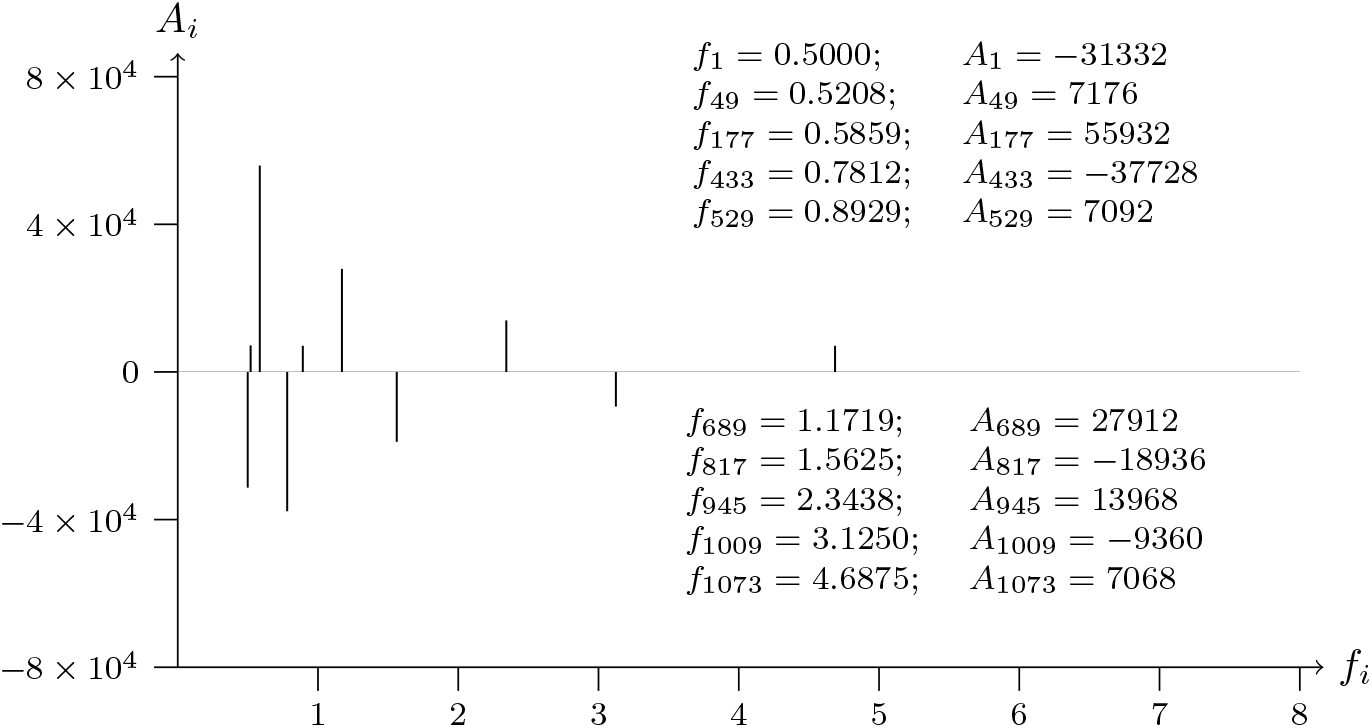
T2T-CHM13v2.0; chromosome 4; MXD4 exon-NM 006454.3-6 analyzed data from 2247404 to 2246204

**Figure 64a:**
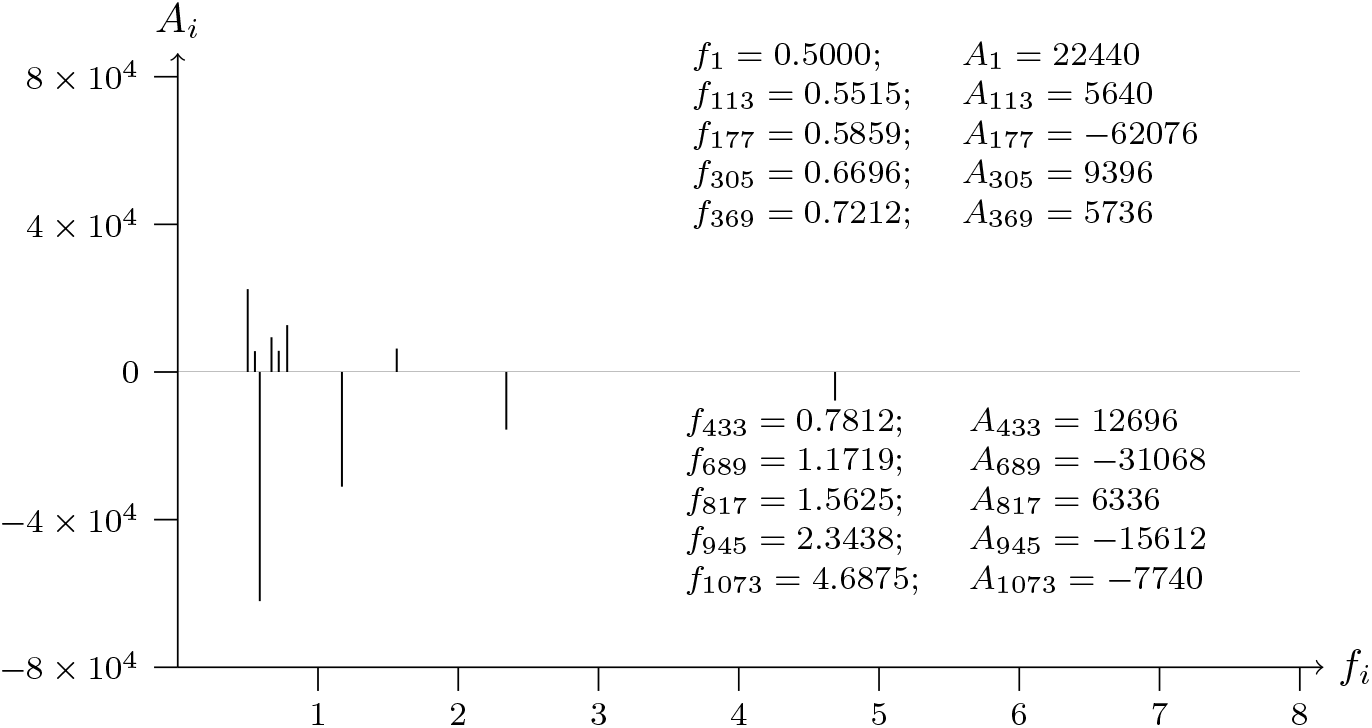
T2T-CHM13v2.0; chromosome 4; AFAP1 exon-NM 001371091.1-18 analyzed data from 7730695 to 7731895

**Figure 64b:**
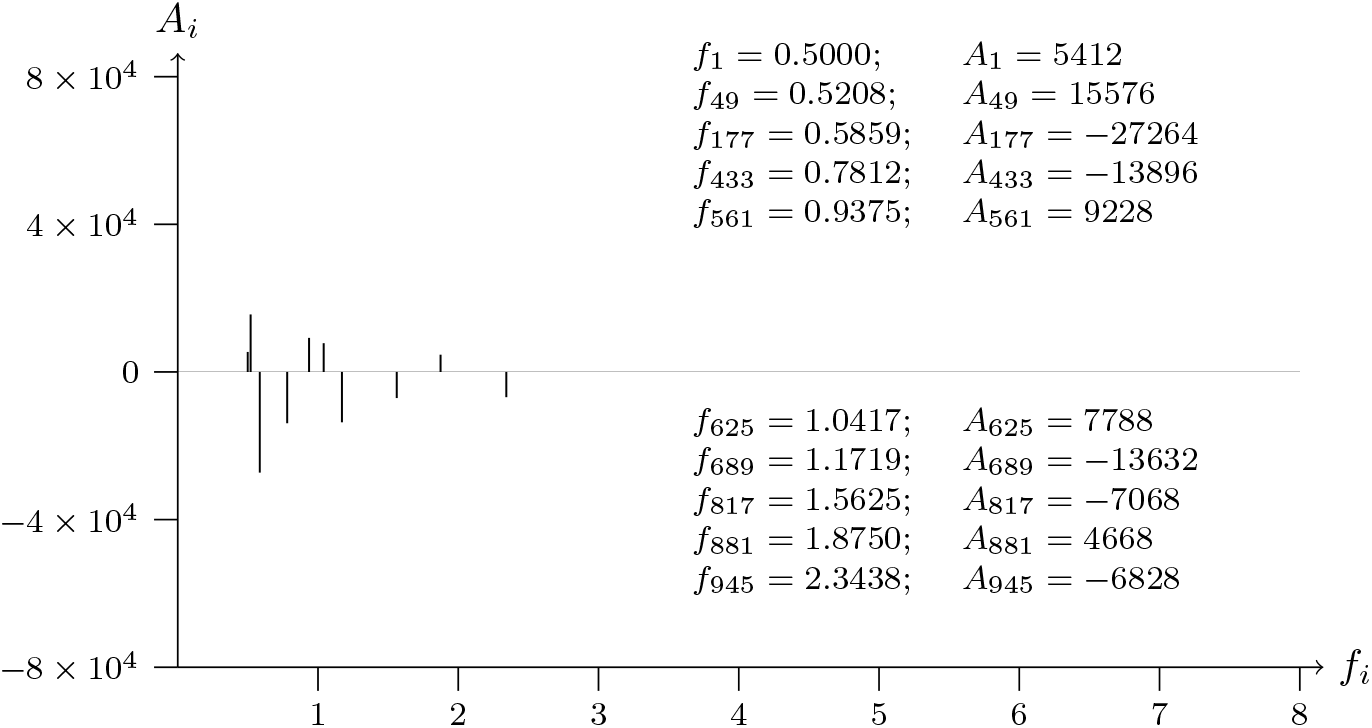
T2T-CHM13v2.0; chromosome 4; AFAP1 exon-NM 001371091.1-18 analyzed data from 7731895 to 7730695

**Figure 65a:**
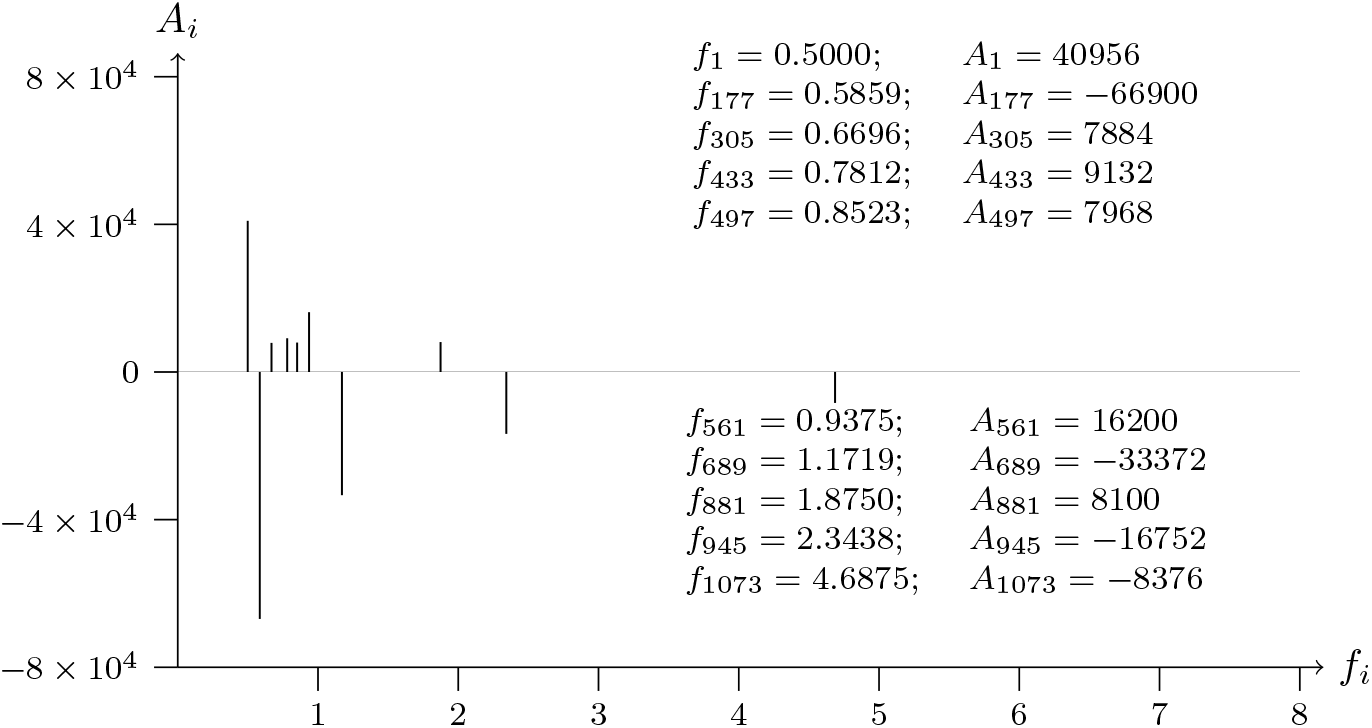
T2T-CHM13v2.0; chromosome 19; SIGLEC10 exon-NM 001171161.2-9 analyzed data from 54498651 to 54499851

**Figure 65b:**
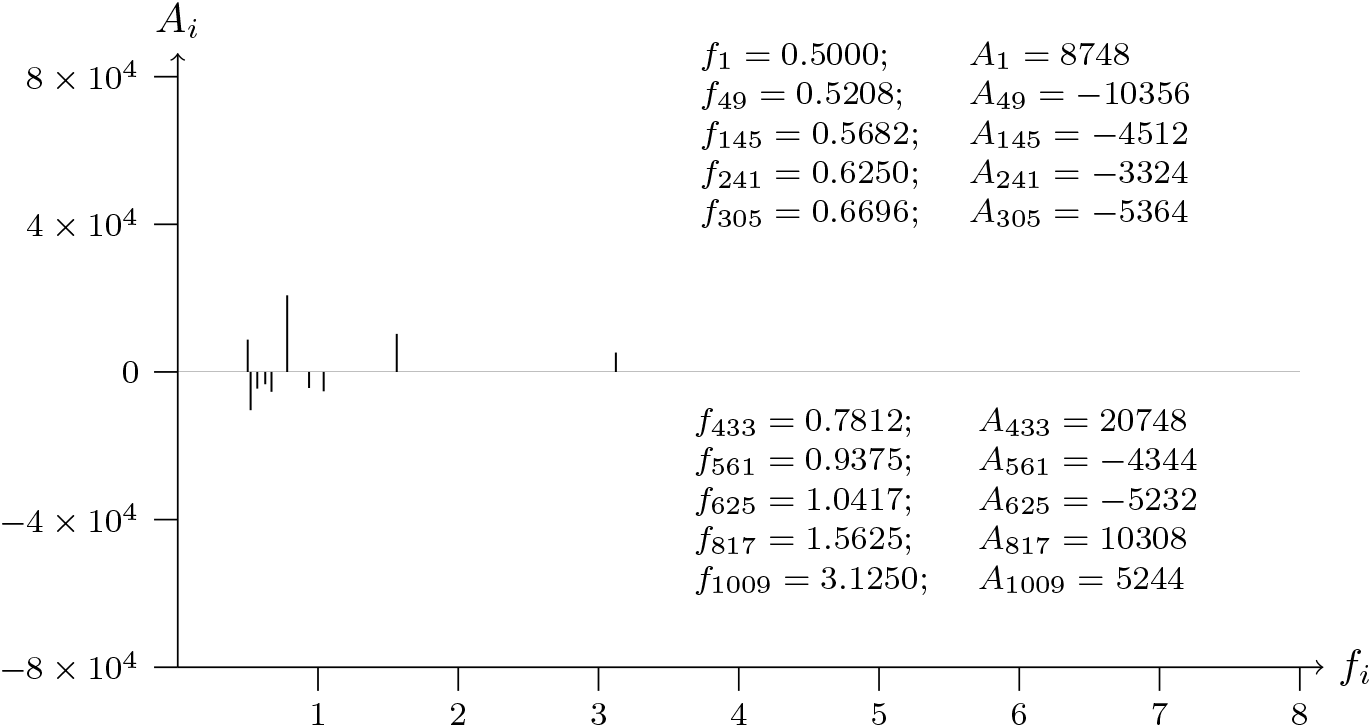
T2T-CHM13v2.0; chromosome 19; SIGLEC10 exon-NM 001171161.2-9 analyzed data from 54499851 to 54498651

**Figure 66a:**
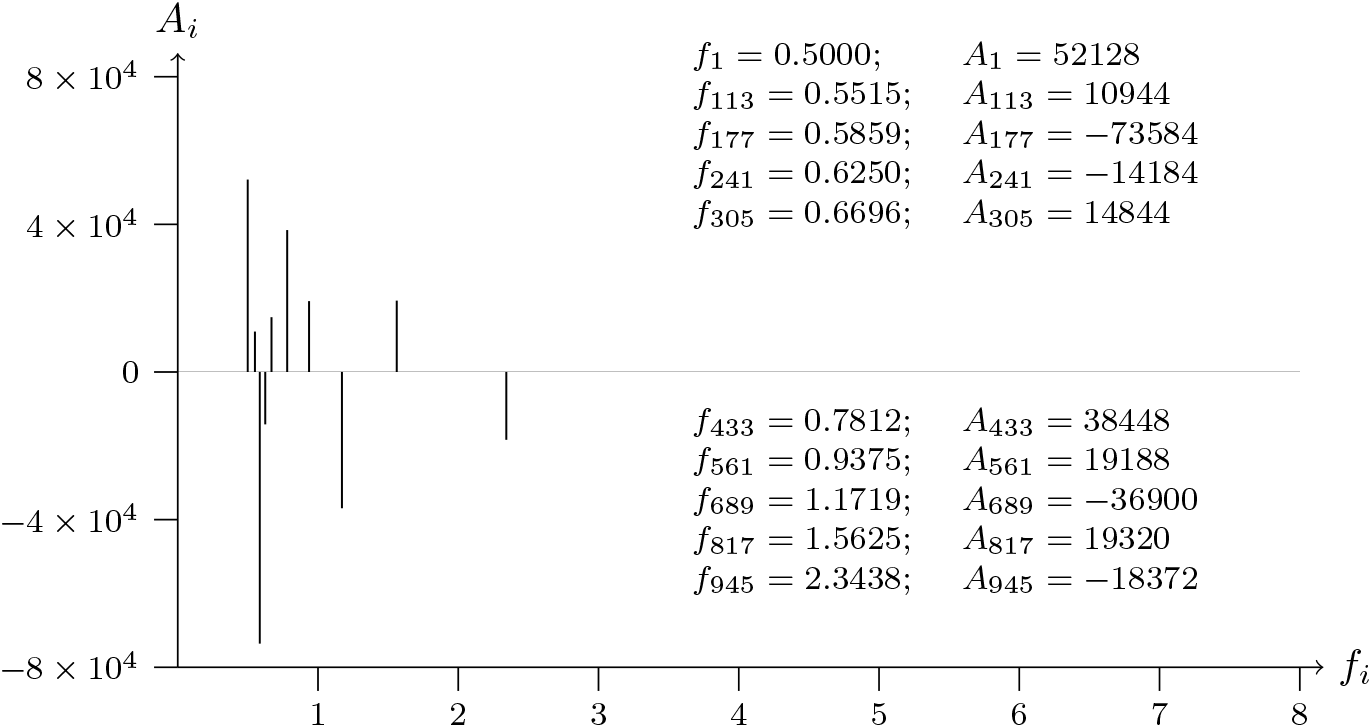
T2T-CHM13v2.0; chromosome 19; CEACAM1 exon-NM 001184816.2-7 analyzed data from 45326710 to 45327910

**Figure 66b:**
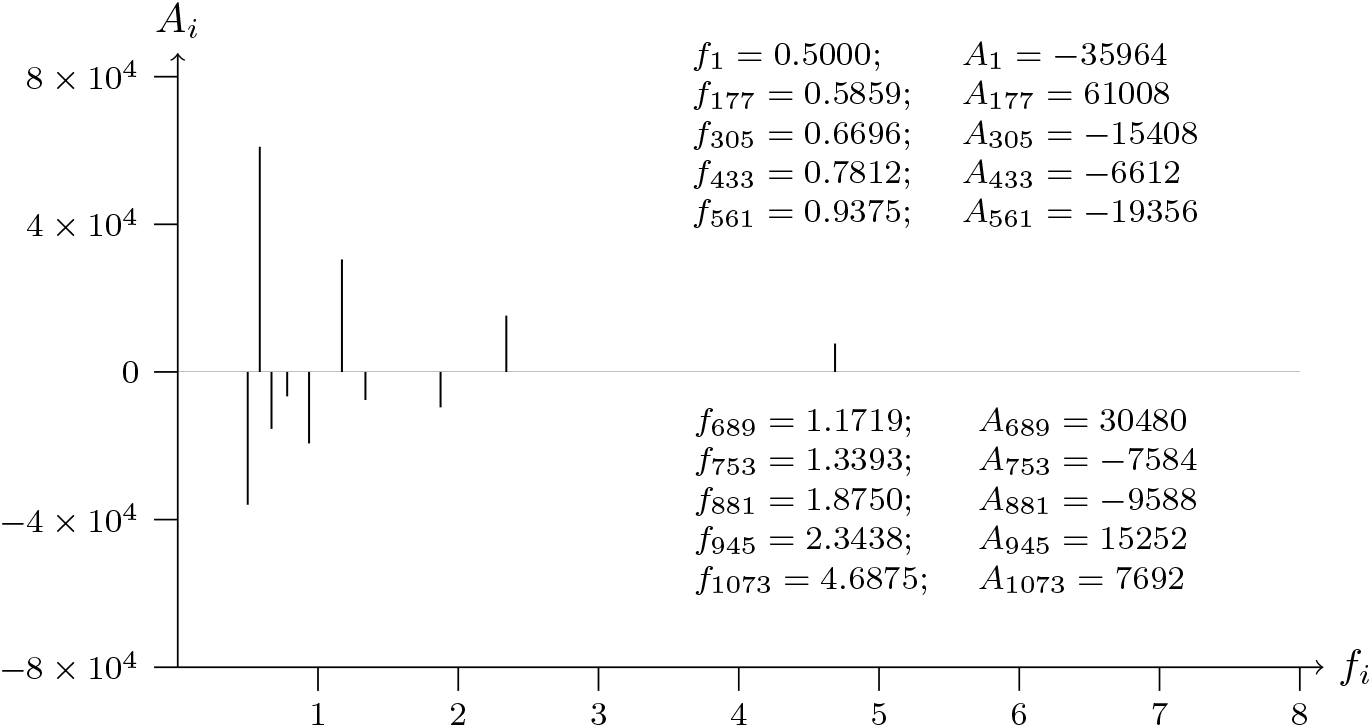
T2T-CHM13v2.0; chromosome 19; CEACAM1 exon-NM 001184816.2-7 analyzed data from 45327910 to 45326710

**Figure 67a:**
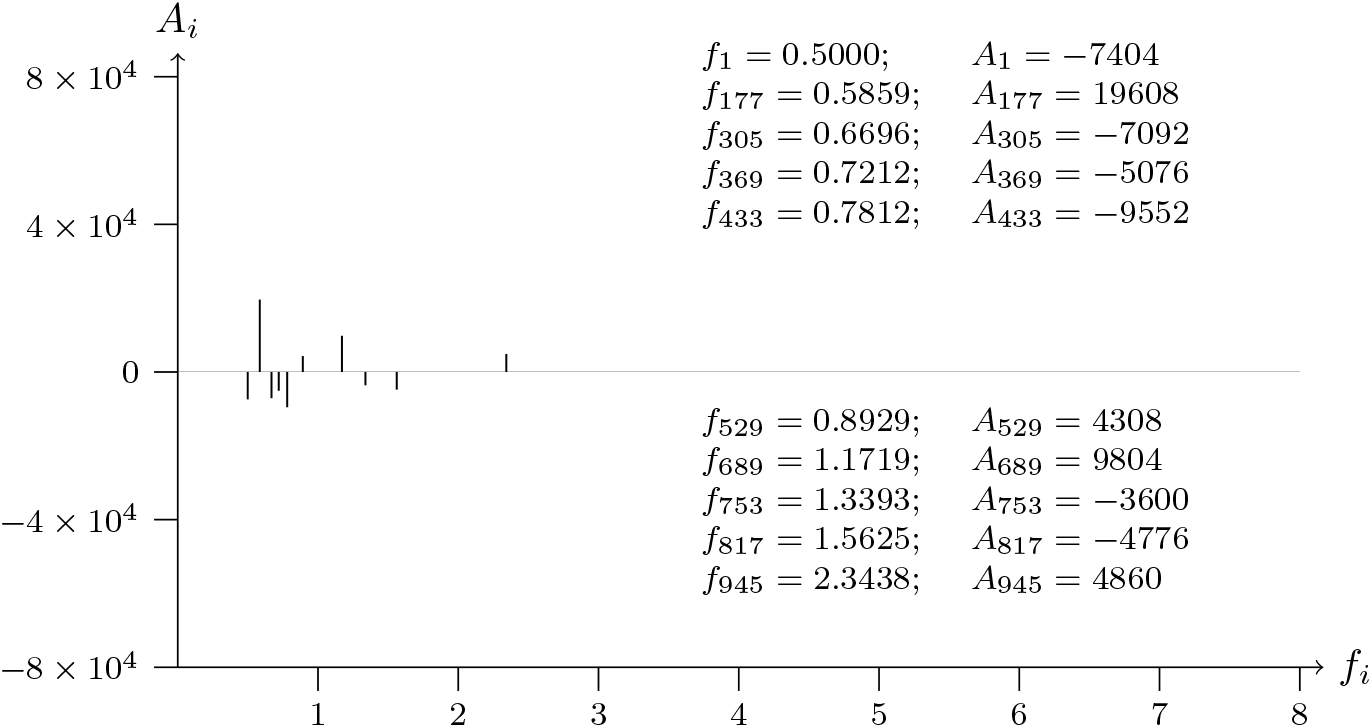
T2T-CHM13v2.0; chromosome 16; ADGRG1 exon-NM 001370440.1-15 analyzed data from 63458652 to 63459852

**Figure 67b:**
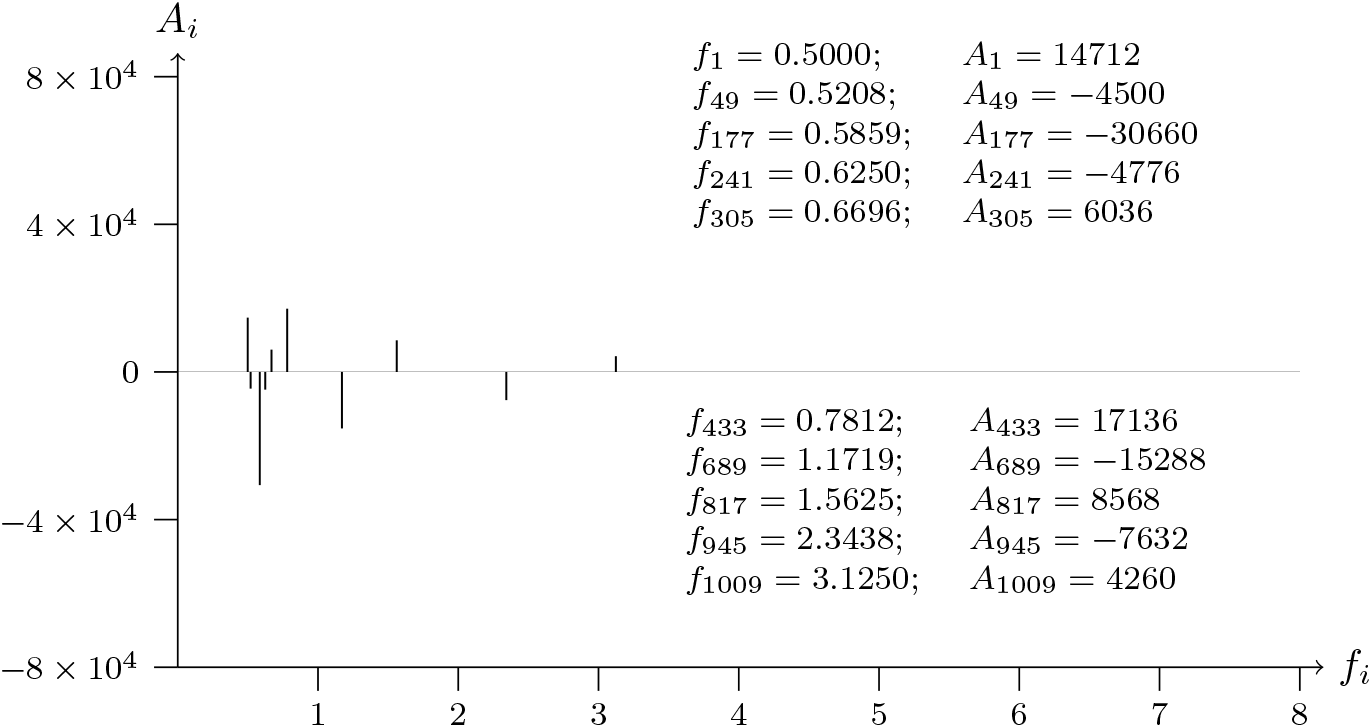
T2T-CHM13v2.0; chromosome 16; ADGRG1 exon-NM 001370440.1-15 analyzed data from 63459852 to 63458652

**Figure 68a:**
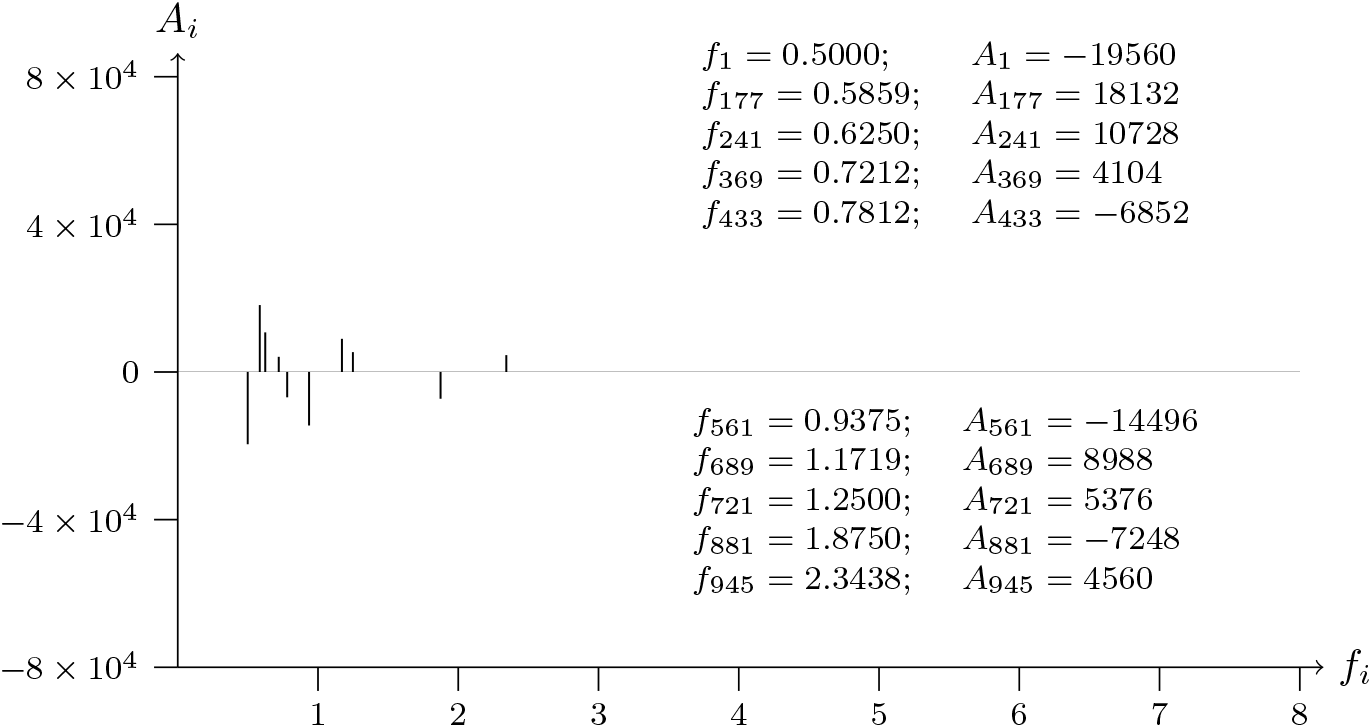
T2T-CHM13v2.0; chromosome 16; KNOP1 exon-NM 001348534.2-4 analyzed data from 19632739 to 19633939

**Figure 68b:**
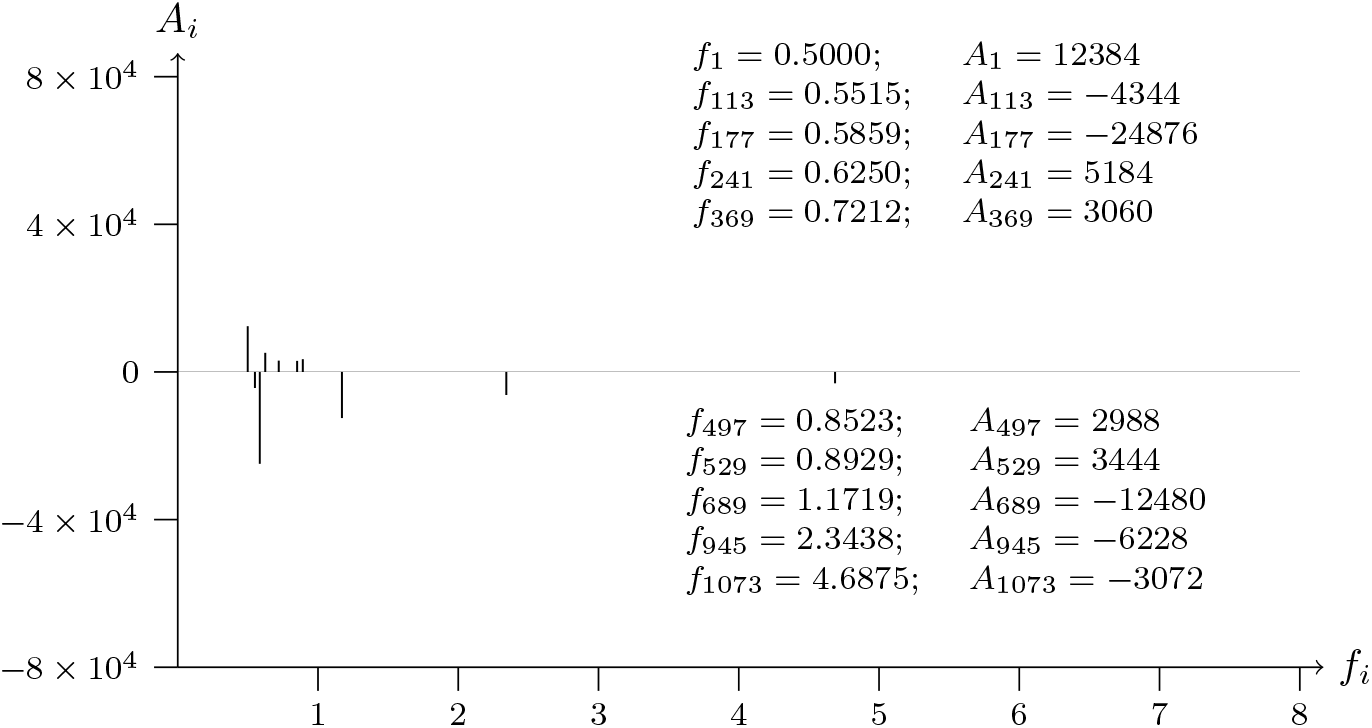
T2T-CHM13v2.0; chromosome 16; KNOP1 exon-NM 001348534.2-4 analyzed data from 19633939 to 19632739

## 5 Preliminary Observations on Some of the Results Obtained

If instead of considering the values of the *A_i_* and the *f_i_* corresponding to the 10 greatest amplitudes of the 1200 *A_i_*, for *i* = 1, 2, 3*, …,* 1200, consideration is given to the *A_i_* and the *f_i_* corresponding to the 40 greatest amplitudes of those *A_i_*, then the 40 *A_i_*and the corresponding *f_i_*will be known as the “prominent” ones. This is what was done when the following preliminary observations were carried out on some of the results obtained.

Attention was given to 77 samples from diverse parts of the human genome [131] (T2T-CHM13v2.0, downloaded from the National Center for Biotechnology Information Genome Database) of 1200 bases of DNA material [131]. Each sample of 1200 bases was analyzed in the same order as it is provided in the downloaded material [132].

Only the 40 most “prominent” *A_i_* and their respective frequencies *f_i_* were considered for each sample.

The information about the names and locations of genes, exons and introns has been extracted from secondary databases from the same datasource [132] (using the alignment file provided by the server). The listing of cancer-related genes (oncogenes and tumor suppressors) has also been obtained from NCBI [133].

Therefore, regarding the most common frequencies corresponding to those “prominent” *A_i_*of the 77 samples, we can observe that:

1. Certain frequencies are present in 100% of the 77 samples, such as *f*_177_ = 0.585938 and *f*_689_ = 1.171875. This means that the presence of “prominent” *A_i_* for these frequencies is related to genomic data in general.
2. Other frequencies are present in most cases (with no regard to any classification); for example, *f*_433_ = 0.781250 is “prominent” in 80% of the 77 cases analyzed.
3. Some frequencies distinguish specific subsets of cases:

a. *f*_1_ = 0.5 is “prominent” in 80% of the telomeres analyzed and it is always “prominent” (100%) in the remaining samples.
b. *f*_241_ = 0.625 is closely related to exons, because it is “prominent” in 80% of the exons, with no distinction as to the type of exon (whether it is a normal exon or a cancer-related exon).
c. *f*_625_ = 1.041667 is closely related to oncogenes and tumor suppression genes, since it is “prominent” in 80% of the exons and introns related to those genes in the subset analyzed.
d. *f*_81_ = 0.535714 and *f*_137_ = 9.375 can be interpreted as markers for introns, due to their appearance in 80% of the introns (with no distinction as to the type of the corresponding gene).
e. *f*_305_ = 0.669634 is not present in telomeres; thus it can be used to distinguish between telomeric data and non-telomeric data.
f. *f*_817_ = 1.5625 and *f*_945_ = 2.34375 appear to group non-coding data (telomeres, centromeres, introns); that occurs in 90% of samples analyzed.
g. *f*_369_ = 0.721154 is “prominent” in 80% of the samples of telomeres and introns, but that frequency does not appear to be significant in centromeres.
h. *f*_49_ = 0.520833 is present in all the cancer-related exons (these exons are part of oncogenes and tumor suppressor genes); and in 80% of the introns and telomeres.

## 6 Discussion and Perspectives

The SWM has certain characteristics of interest with respect to other possible methods for the analysis of numerical sequences. Thus, for example, a) it can be applied both for overall and local analysis; b) it has no “effect” such as that of the Gibbs phenomenon which would require corrections to ensure the quality of the approximations achieved to the numerical sequences analyzed; c) it depends on the order in which the elements constituting each numerical sequence analyzed are presented, and not simply on the statistical distribution of those elements; and d) the complete results of each analysis make it possible to reconstruct, unambiguously, the sequence of bases of DNA analyzed.

The preliminary observations specified in section 4 suggest certain patterns that are detected by the SWM. Experts in genetics and bioinformatics will determine whether these interesting regularities support the hypothesis formulated in the introduction of this article – in which case using data analysis techniques to go into greater depth on the significance of the results obtained would be justified – or whether this hypothesis should be discarded.

